# Remote homology search with hidden Potts models

**DOI:** 10.1101/2020.06.23.168153

**Authors:** Grey W. Wilburn, Sean R. Eddy

## Abstract

Most methods for biological sequence homology search and alignment work with primary sequence alone, neglecting higher-order correlations. Recently, statistical physics models called Potts models have been used to infer all-by-all pairwise correlations between sites in deep multiple sequence alignments, and these pairwise couplings have improved 3D structure predictions. Here we extend the use of Potts models from structure prediction to sequence alignment and homology search by developing what we call a hidden Potts model (HPM) that merges a Potts emission process to a generative probability model of insertion and deletion. Because an HPM is incompatible with efficient dynamic programming alignment algorithms, we develop an approximate algorithm based on importance sampling, using simpler probabilistic models as proposal distributions. We test an HPM implementation on RNA structure homology search benchmarks, where we can compare directly to exact alignment methods that capture nested RNA base-pairing correlations (stochastic context-free grammars). HPMs perform promisingly in these proof of principle experiments.

**Author summary:** Computational homology search and alignment tools are used to infer the functions and evolutionary histories of biological sequences. Most widely used tools for sequence homology searches, such as BLAST and HMMER, rely on primary sequence conservation alone. It should be possible to make more powerful search tools by also considering higher-order covariation patterns induced by 3D structure conservation. Recent advances in 3D protein structure prediction have used a class of statistical physics models called Potts models to infer pairwise correlation structure in multiple sequence alignments. However, Potts models assume alignments are given and cannot build new alignments, limiting their use in homology search. We have extended Potts models to include a probability model of insertion and deletion so they can be applied to sequence alignment and remote homology search using a new model we call a hidden Potts model (HPM). Tests of our prototype HPM software show promising results in initial benchmarking experiments, though more work will be needed to use HPMs in practical tools.

## Introduction

An important task in bioinformatics is determining whether a new sequence of unknown biological function is evolutionarily related, or homologous, to other known sequences or families of sequences. Critical to the concept of homology is alignment: homology tools create multiple sequence alignments (MSAs) in which evolutionarily related positions are aligned in columns by inferring patterns of sequence conservation induced by complex evolutionary constraints maintaining the structure and function of the sequence [1].

Homology search tools are used in a wide range of biological problems, but it is common for these tools to fail to identify distantly related sequences; many genes evolve quickly enough that homologs may exist yet be undetectable [2]. More powerful and sensitive homology search tools are therefore needed.

One possible way to improve the sensitivity of homology search and alignment is to develop new methods that successfully capture patterns of residue correlation induced by 3D structural constraints. State-of-the-art homology search methods do not model certain important elements of structure-induced conservation. Methods such as BLAST and HMMER, the latter of which uses profile hidden Markov models (pHMMs), align and score sequences using primary sequence conservation alone [3, 4, 5]. Specific methods for RNA homology search, such as the software package Infernal, use a class of probabilistic models called profile stochastic context free grammars (pSCFGs) to infer primary *and* secondary structure conservation in RNA [6, 7]. However, Infernal is limited to nested, disjoint pairs of nucleotides, meaning it cannot capture complicated 3D RNA structural elements like pseudoknots and base triples, let alone complex correlation structure in protein MSAs.

An opportunity to build a more sensitive sequence homology search method has arisen via two key developments. First, there has been recent exciting progress in exploring pairwise correlations between columns in protein and RNA MSAs using Potts model (aka Markov random field) methods from statistical physics [8, 9, 10, 11, 12, 13, 14]. Previous work has applied Potts models to homology search and alignment problems with specific proteins, but not to biological sequences in general [15, 16, 17, 18, 19, 20, 21]. Potts models have also been used to study protein-protein interactions [22, 23, 24, 25], mutational effects [26, 27, 28, 29, 30], cellular morphogenesis [31], and collective neuron function [32]. Building upon previous methods that use pairwise sequence correlation to infer conserved base pairs in RNA structure and 3D structure in proteins [33, 34], a Potts model expresses the probability that a particular sequence belongs to a family represented by an MSA as a function of all possible characters (amino acids or nucleotides) at each position and all possible pairs of characters across all positions.

Second, very large datasets of aligned homologous protein and RNA sequences are available, such as Pfam for protein and Rfam for RNA [35, 36]. Deep MSAs allow for new methods that analyze more subtle patterns of sequence conservation.

Potts models have two drawbacks for applications in sequence homology search. First, Potts models assume *fixed length* sequences in *given* MSAs, treating insertions and deletions as an extra character (5th nucleotide or 21st amino acid). Homology search involves scoring *variable length* sequences and *inferring* alignments. Second, models such as pHMMs and pSCFGs use efficient dynamic programming algorithms that align and score sequences in polynomial time. Unfortunately, the all-by-all pairwise nature of Potts models makes them incompatible with dynamic programming algorithms and therefore impractical for homology search.

It would be useful to have a homology search tool that can both handle higher-order sequence correlation patterns, as a Potts model can, *and* efficiently align and score variable length sequences, as BLAST, HMMER, and Infernal are able to do. Inspired by early work using Potts models for protein alignment and the success of Potts models in structure prediction, we have developed a class of models called hidden Potts models (HPMs) that combines the coupled Potts emission process with a probabilistic insert-deletion model. In addition, we have developed an approximate algorithm that uses importance sampling to align and score variable length sequences to a parameterized HPM.

We have built a proof of principle HPM software implementation. To test the efficacy of HPMs in homology search and alignment, we compare our software to HMMER and Infernal in RNA remote homology benchmark tests based on trusted, hand-curated noncoding RNA alignments. RNA is an ideal testbed, as secondary and tertiary structure are largely dictated by stereotyped base-pairing interactions, making model parameterization interpretable. Also, pSCFGs capture nested pairwise correlations, a key part of RNA homology search, leading to a natural comparison to an existing approach that captures much of the pairwise residue correlation structure in conserved RNAs. As HPMs capture nested *and* non-nested pairwise correlations, we expect them to at least equal, if not outperform, pSCFGs in RNA remote homology search and alignment.

## Results

### Hidden Potts models emit variable length sequences

A generative homology model describes the probability of creating an unaligned sequence of observed characters, 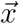. In the case of biological sequences, characters in 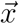 represent monomer units in a specific alphabet (20 amino acids for protein, 4 nucleotides for DNA and RNA).

A Potts model expresses the probability of a homologous sequence as a function of primary conservation *and* all possible pairwise correlations between all consensus sites in a biological sequence (i.e., consensus columns in a multiple sequence alignment). Going beyond a Potts model, a hidden Potts model consists of *M* sites called *match states* (squares in Fig 1), corresponding to well-conserved columns in an MSA, and intermediate insert states (diamonds) that handle variable-length insertions between match states. An HPM has a fixed number of match states, which restricts the model to *glocal* alignment (local to the sequence but global to the model).

**Figure 1:**
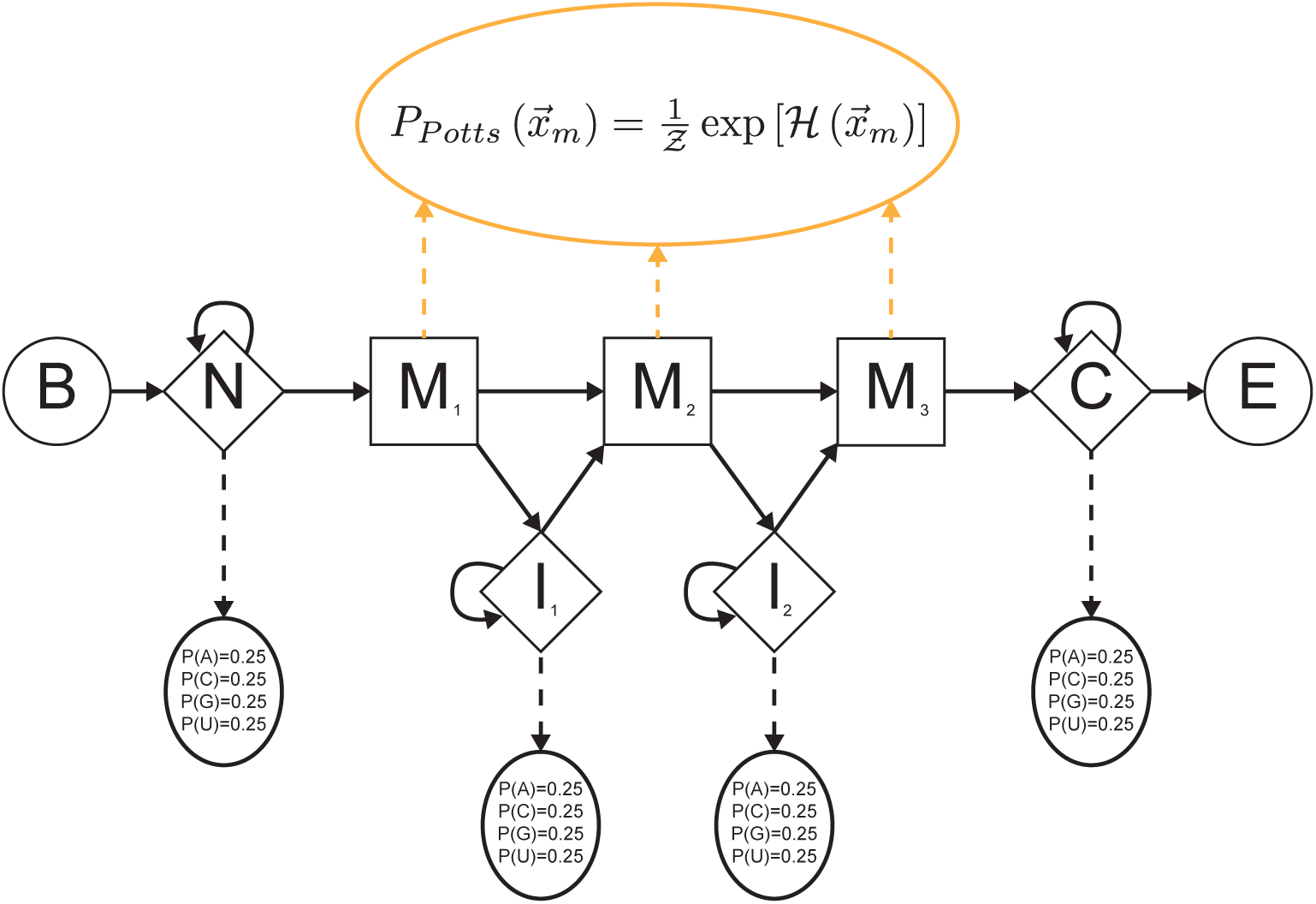
Hidden Potts model architecture. Squares are conserved match states and diamonds are insert states. No delete states exist. Silent begin and end states are represented by circles. An HPM is a hybrid between a Potts model and a pHMM: correlated character generation (including deletion “characters” rather than delete states) in match columns (consensus sites in an MSA) comes from a Potts distribution (dotted orange arrows), while transition probabilities linking states and site-independent character emissions in unaligned insert columns come from a pHMM (solid and dashed black arrows, respectively).

An HPM is a hybrid between a profile hidden Markov model (pHMM) [4] and a Potts model. Much like a pHMM, an HPM consists of an observed sequence 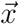 and a *hidden* state path 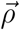 representing the alignment of 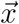 to the model. 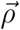 is a Markov chain, with the individual states mapping to columns in a multiple sequence alignment. Each state *emits* an observed character, with all emissions combining to form 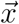.

Unlike a pHMM, HPM states do not emit characters independent of other states, and deletions relative to the model’s consensus are handled differently. HPM match states generate a set of characters with a single, correlated emission from a Potts model. As Potts models can only handle fixed length sequences, HPMs do not model the lack of a character in a consensus column with a special delete state, but rather as the emission of a “dummy” deletion character from a match state. The deletion character effectively serves as an extra character in the alphabet (21st amino acid or 5th nucleotide). As such, we divide 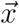 into two non-contiguous subsequences: 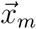, the characters emitted from match states (which may include deletion characters); and 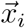, the characters generated by insert states (all of which represent monomers). A pHMM path, which we represent as 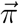 and can include delete states, is not identical to a corresponding HPM path: a single HPM path 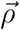 can map to up to 2^*M*^ possible paths under a pHMM, representing the possible distribution of deletions in consensus columns.

Mathematically, 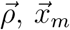, and 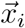 are nuisance variables that must be marginalized over to obtain 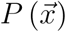. The joint probability of 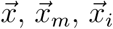, and 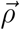 under an HPM can be factorized into two terms.

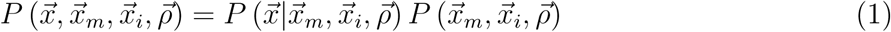

Given 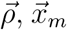 and 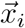 can be dealigned to produce a unique, unaligned sequence 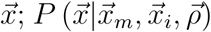 is 1 if 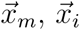, and 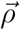 can be combined to produce unaligned sequence 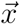 and 0 otherwise.

The joint probability of 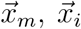, and 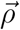 can further be factored into three terms: 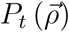, a product of transition probabilities linking states to one another (solid black arrows in Fig 1); 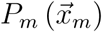, describing the Potts emission of characters from match states (dashed orange arrows); and 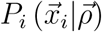, a product of independent emission of characters from insert states (dashed black arrows).

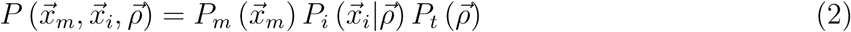

The entire match sequence 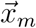 is generated by one multi-character emission from the Potts distribution. The Potts distribution assumes the probability of generating a sequence from match states is given by an exponential Boltzmann distribution:

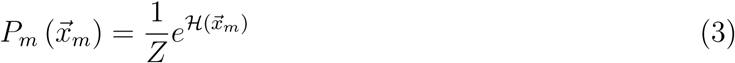

Here 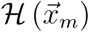 is referred to as the *Hamiltonian*, while *Z* is the *partition function*, a normalization constant. The Hamiltonian is given by:

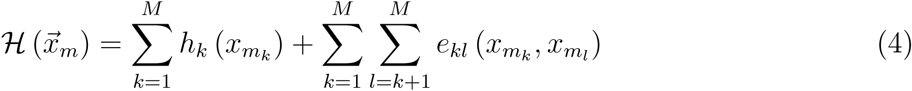

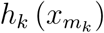 represents the *single position preference* for character 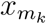 in match state *m*_*k*_. 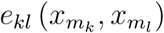 corresponds to the *statistical coupling* between character preferences at separate match columns *k* and *l*. These are the Potts model parameters that we estimate from an input MSA.

The insert emission probability factorizes into independent terms for each character in 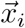.

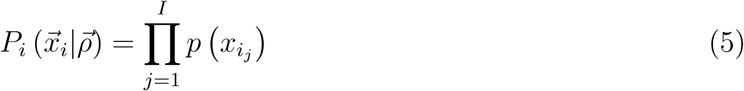

The probability of a state path 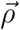 is given by:

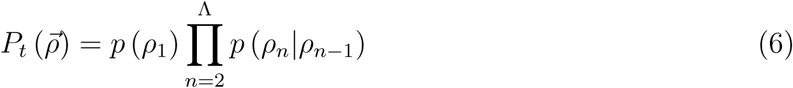

Here, Λ is the total number of states in 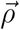; with the inclusion of deletion characters, Λ is at least as large as the total number of non-gap characters in the observed sequence, *L*.

HPMs are generative models capable of producing any sequence, while the inclusion of possible self-transitions within insert states allows for the generation of sequences of any length. The steps to generate a sequence are:

1. Choose the path 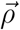 by performing a random walk along the state path following 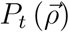.
2. Choose insert state characters by independently drawing from 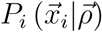.
3. Generate match column characters from a single emission from 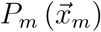 (the Potts distribution).
4. Dealign the sequence by combining 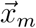 and 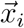 and removing deletion characters. This corresponds to choosing the unaligned sequence 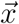 for which 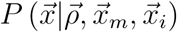 is 1.

### Training an HPM

Our hidden Potts model design contains two independent pieces, the pHMM-like and Potts-like parts of the model, that we train separately. We train the pHMM-like elements (*P*_*t*_, *P*_*i*_) using the HMMER software package program `hmmbuild` [5]. We train Potts models used for *P*_*m*_ with the Gremlin structure prediction software [11], which uses an approximate pseudolikelihood maximization method to estimate the *h*_*k*_ and *e*_*kl*_ Potts model terms [37]. Both programs take the same MSA as input. We combine the two models with our own code (`hpmbuild`) to produce a fully parameterized HPM.

A Potts model produced by Gremlin is not normalized, as the partition function *Z* is not calculated. Therefore, we are only able to calculate the probability of a given 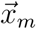 under a Potts model up to *Z*.

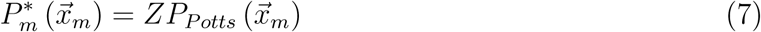

Thus, we only know the marginal probability of a sequence under an HPM up to a factor of Z.

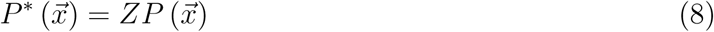

Implications of using unnormalized probability distributions are discussed below.

### Importance sampling alignment algorithm

In order to use HPMs in remote homology search, we need algorithms to efficiently align and score sequences of varying length with a parameterized HPM. Profile HMMs and profile SCFGs use dynamic programming algorithms to optimally align and score sequences in polynomial time. However, dynamic programming algorithms do not work in models like HPMs with non-nested correlation terms (*e*_*kl*_’s).

Aligning a sequence consists of finding the best possible combination of 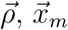, and 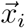 for a given sequence under an HPM.

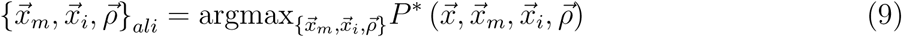

Scoring a sequence entails summing the unnormalized joint probabilities for a given sequence 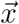 and the set of *all possible* combinations of 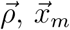, and 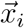 to find the unnormalized probability that the HPM generated the sequence, 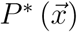.

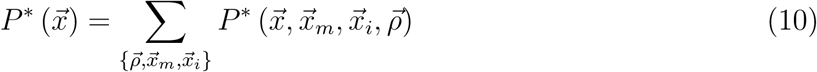

Evaluating all possible combinations of 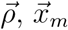, and 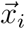 by brute force enumeration is computationally intractable, so we need an approximate method to align and score sequences efficiently. One could imagine using Monte Carlo integration, randomly sampling a finite number of combinations from 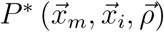 and approximating the marginalization sum above to obtain 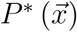. However, the space of possible combinations is enormous, and few of the sampled combinations would yield non-negligible probabilities 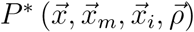.

To more efficiently score and align sequences, we use a related method, *importance sampling*, illustrated in Fig 2 and described in detail in S1 Appendix. Under importance sampling, instead of sampling 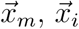, and 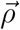 from the prior distribution 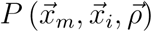, we instead sample a smaller number of paths from a different distribution, the “proposal” distribution, 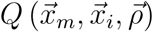, in which the probability mass for all three variables is much more concentrated.

**Figure 2:**
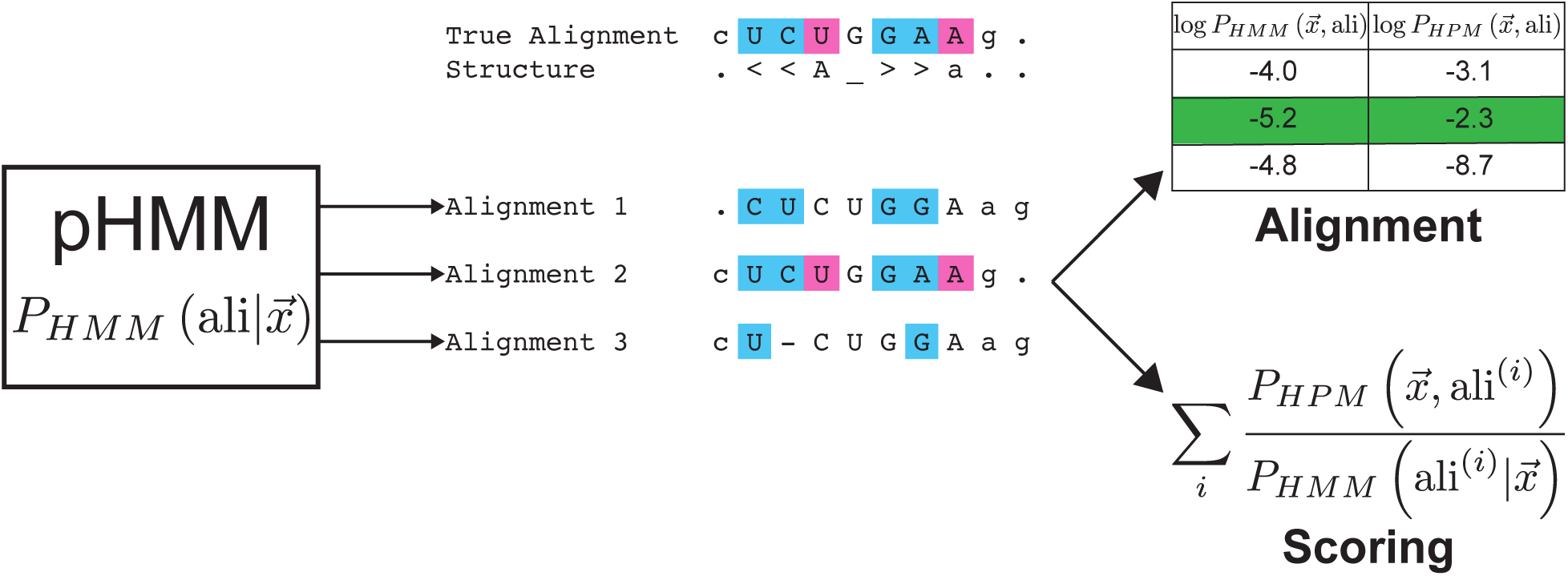
Importance sampling alignment algorithm schematic. Toy example of aligning an RNA sequence CUCUGGAAG to models of its sequence/structure consensus, where the true structural alignment has two nested base pairs (brackets in structure line) and one pseudoknot (‘Aa’). Suboptimal alignments of the sequence are sampled probabilistically using a pHMM. A pHMM does not capture residue correlations due to base-pairing, so only some proposed alignments satisfy the expected consensus nested (cyan) or pseudoknotted (pink) base pairs. The proposed alignments are re-scored and re-ranked under the HPM, which does capture correlation structure; the correct alignment with the highest probability under the HPM is identified (green), and the sequence’s total probability 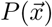 is obtained by importance-weighted summation over the sampled alignments.

Using importance sampling, for *R* samples, 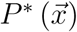 is approximated by:

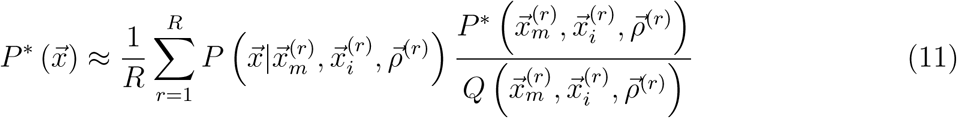

The approximate alignment is the sampled combination of 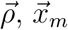, and 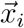 that maximizes the numerator of the importance sampling sum.

For a proposal distribution, we use the posterior probability of an HPM path, match sequence, and insert sequence given an unaligned sequence under a pHMM, 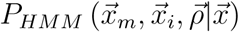. We show in S1 Appendix that 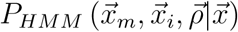 is equivalent to 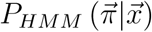, where 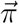 is a pHMM-style path with delete states. The 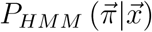 posterior distribution over all possible alignments of a sequence to a pHMM can be sampled exactly and efficiently by stochastic tracebacks of the Forward dynamic programming matrix [38, 1], requiring one *𝒪* (*ML*) time calculation of the Forward matrix followed by an *𝒪* (*L*) time stochastic traceback per sampled alignment. Re-scoring each sampled alignment with the HPM takes *𝒪* (*M* ^2^ + *L*) time, where the most expensive step is evaluating all-vs-all pairwise *e*_*kl*_ terms in the Potts emission of 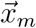. verall, for a proposed sample of *R* alignments, our approach takes *𝒪* (*ML* + *RL* + *RM* ^2^) time, typically dominated by the *RM* ^2^ term.

Intuitively, our approach assumes that a pHMM has the alignment reasonably well constrained, though not correct in detail, which is our experience with alignments of remotely homologous sequences that fall near detection thresholds. Because the pHMM neglects residue correlations, the alignment that best satisfies conserved correlation patterns is often not the optimal pHMM alignment. So long as this alignment is probable enough to be present in a large stochastic sample from the pHMM’s distribution over alignment uncertainty, then rescoring that sample with the more complex HPM will find it.

We have taken 10^6^ samples from 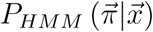 per sequence when aligning and scoring, which takes roughly 1 minute per sequence for typical *M* and *L*. Our tests indicate that the importance sampling approximation for 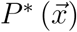 generally converges by 10^6^ samples, and the optimal alignment is generally stable (see S1 Appendix for more information).

To score homology searches, we use a standard log odds score, in which we compare 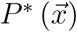 to the probability that 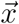 was generated by a null model, 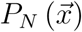 [1].

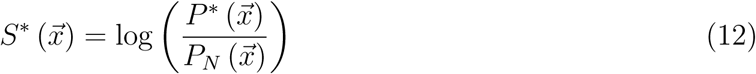

Ideally, we would use a normalized HPM distribution 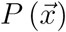, which requires knowing the partition function, to produce a normalized log odds score 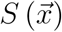. Calculating *Z* exactly using all possible match state subsequences is intractable. As *Z* is a constant for a particular model, scores can be compared relatively within a single database search with a given query HPM, but not qualitatively across different query HPMs (with different unknown *Z*’s). For future work, statistical significance will have to be evaluated empirically. We can still use the importance sampling alignment algorithm to find a maximum likelihood alignment of a target sequence to a query HPM or to calculate the posterior probabilities of alignments of target sequences to an HPM, tasks for which *Z* cancels.

### HPMs perform promisingly

To test the performance of HPMs in remote homology search and alignment, we designed benchmark experiments using a few deep, well-curated RNA alignments. We chose to test on RNA alignments for three reasons. First, secondary and tertiary structure in RNA is largely dictated by nucleotide base-pairing, which makes the *e*_*kl*_ terms in the HPM easily interpretable. Second, pSCFG methods already use RNA secondary structure conservation patterns, so we can compare the performance of HPMs not just to sequence-only models like pHMMs, but also to existing models that already capture most RNA pairwise correlation structure. Finally, because the residue correlation structure in RNAs largely consists of disjoint base pairs, we can make some back-of-the-envelope observations about how much additional information content an HPM will capture compared to a pSCFG or a pHMM. This estimation serves as a gauge of how much we might be able to improve homology search and alignment, as discussed next.

### Information content of RNA multiple sequence alignments

We can get a sense of how much additional statistical signal is capturable by HPMs compared to pSCFGs and pHMMs from the Shannon relative entropies (information content) of single MSA columns and column pairs. The relative entropy of a residue emission probability distribution for a single position [39] is roughly the expected score per position in a pHMM. The difference between the relative entropy of the joint emission distribution for two positions and the sum of their individual relative entropies is the *mutual information*, the extra information (in bits) gained by treating the two positions as a correlated pair. We define secondary structure information content as the summed mutual information of all nested base pairs in an RNA consensus structure; this is a measure of how much statistical signal a pSCFG gains over a pHMM for an RNA alignment. Similarly, we can define “tertiary structure” information content as the summed mutual information of all other disjoint base pairs in an RNA consensus structure, such as pseudoknots, not included in a nested pSCFG description. This is not a complete picture of the statistical signal capturable by an HPM, because the measure neglects non-disjoint pairs (e.g. base triples) and other higher-order correlations that an HPM can also capture. However, RNA pseudoknots are one of the most important features we aim to capture in more powerful sequence homology search and alignment models.

As shown in Fig 3, most of the information content in Rfam RNA MSAs results from primary and secondary structure conservation. Tertiary structure information content contributes to a lesser degree. These results indicate that HPMs have the potential to only slightly gain sensitivity over a pSCFGs in RNA homology search. However, when trying to improve the sensitivity of homology search, even small increases in signal are potentially useful.

**Figure 3:**
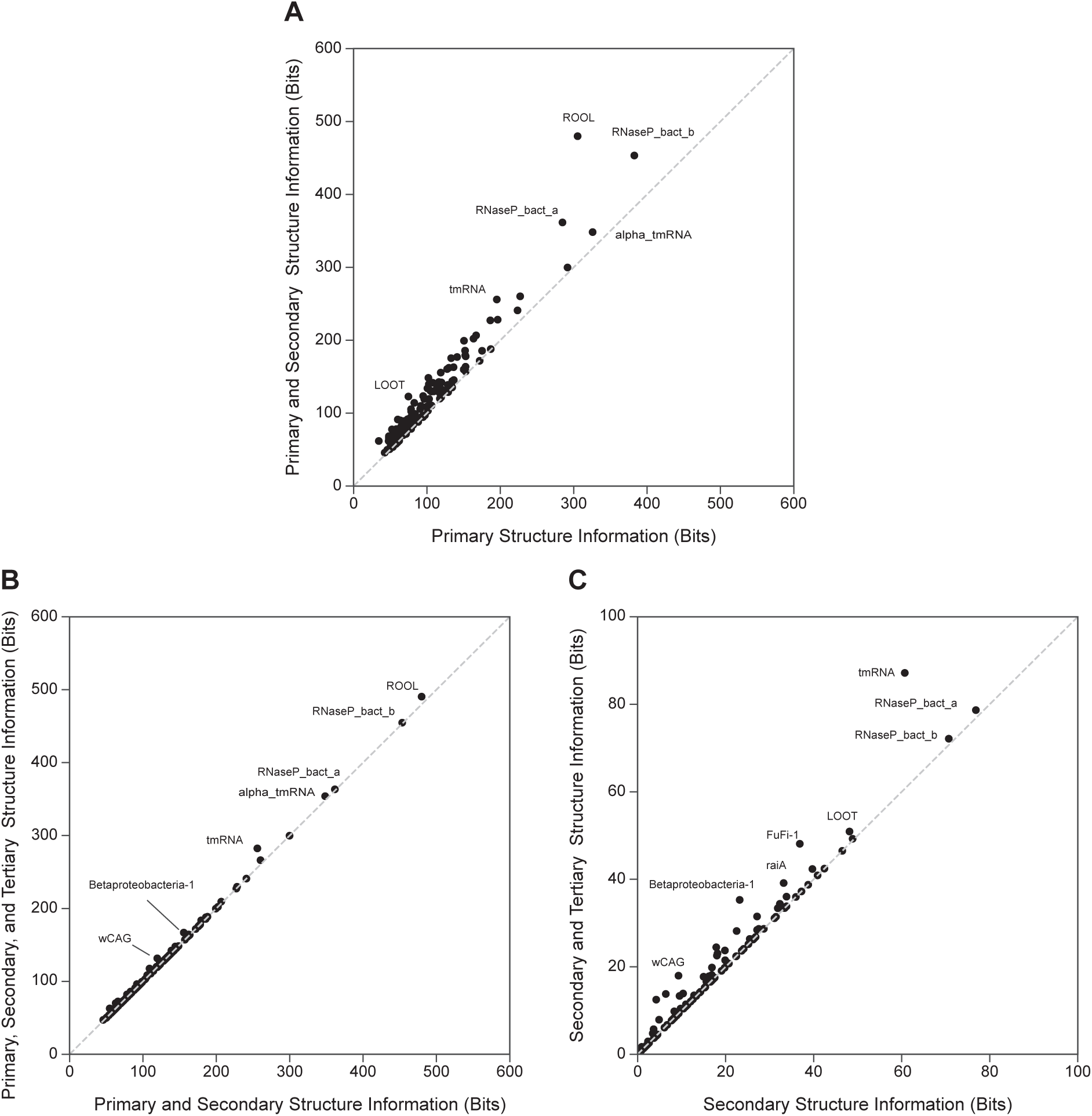
Additional information contained in pairwise covariation in RNA MSAs. For each of the 127 Rfam 14.1 seed MSAs with more than 100 sequences [36], we infer a predicted consensus structure (including nested and non-nested base pairs) using CaCoFold and R-scape [40, 41]. (A) Primary information content (X axis) versus the sum of primary and secondary information content (Y axis). (B) Sum of primary and secondary information content (X axis) versus the information content from all three levels of structure (Y axis) (C) Secondary information content (X axis) versus the sum of information content from secondary and tertiary structure (Y axis). Not included in (C) is ROOL (RF03087), with 305.3 bits of primary information, 174.5 bits of secondary information, and 10.7 bits of tertiary information.

### Benchmark procedure

We choose to use a small number of well-studied sequence families of known 3D structure, vidual alignment results in detail. We use three RNA alignments for these experiments: an alignment of 1415 transfer RNA (tRNA) sequences [42], a 1446-sequence Twister type P1 ribozyme MSA [43], and the 433-sequence SAM riboswitch seed alignment from Rfam family RF00162 [36]. Summary statistics for each benchmark dataset are shown in table 1. Each training alignment contains at most a small amount of information content resulting from pairwise covariation induced by conserved tertiary structure.

**Table 1:** Benchmark dataset statistics and training alignment information content. Hand curated MSAs are split into training and test sets based on [44]. For each training MSA, information content in the primary sequence (in bits) is calculated [39], while information in secondary structure (nested base pairs) and tertiary structure (all other disjoint pairwise interactions between sites) is estimated using mutual information [6]. Each family’s consensus structure is inferred using CaCoFold and R-scape on the the training alignment [40, 41]. Though R-Scape identifies no tertiary structure using the SAM riboswitch training alignment, a four-base pair pseudoknot has been observed experimentally [45]. This lack of pseudoknot detection is a characteristic of our SAM training alignment; R-scape predicts the pseudoknot when analyzing the RF00162 seed alignment.

| Family | $N_{train}$ | $N_{test}$ | Consensus Length | 1° info (bits) | 2° info (bits) | 3° info (bits) |
| --- | --- | --- | --- | --- | --- | --- |
| tRNA | 1357 | 30 | 72 | 40.3 | 26.7 | 1.2 |
| Twister | 1005 | 40 | 65 | 57.5 | 4.5 | 1.3 |
| SAM | 192 | 8 | 108 | 121.8 | 11.1 | 0.0 |

To create benchmark datasets, we divide each reference MSA into an in-clade training set and a remotely homologous test set, as we have done in previous work on RNA homology search [44]. To create a challenging test and simulate a search for very distantly related sequences, we cluster sequences by single linkage clustering and then split the clusters so that the following conditions are satisfied:

- No test sequence is more than 60 % identical to any training sequence.
- No two test sequences are more than 70 % identical.

For scoring benchmarks, we add 200,000 randomly-generated, non-homologous decoy sequences to the test set. We use synthetic sequences in order to avoid penalizing a method that identifies remote, previously unknown evolutionary relationships. Decoys are created with characters drawn i.i.d from the nucleotide composition of the positive test sequences, with the length of each decoy matching a randomly-selected positive test sequence. We score all of the test and decoy sequences with homology models built using the training MSAs. We measure scoring effectiveness by creating a ROC plot that shows true positive rate, or sensitivity, as a function of false positive rate.

To test the effectiveness of HPMs in remote homology alignment, we measure how accurately each model aligns remote homologs relative to the original alignment. We define accuracy to be the fraction of residues in consensus columns aligned correctly across all test sequences. We do not measure alignment accuracy in insert columns, as pHMMs, pSCFGs, and HPMs do not align characters in insert states.

### Scoring and alignment benchmark results

The chief motivation in using HPMs for homology search is the potential for conserved patterns of pairwise evolution to be captured by the all-by-all pairwise *e*_*kl*_ terms. Are the *e*_*kl*_’s useful for homology search? To answer this question, we compare the performance of default HPMs with all *e*_*kl*_’s included (red curves in Fig 4) against HPMs for which the *e*_*kl*_ terms are restricted to 0 in training (“no *e*_*kl*_” HPMs, black curves). The HPM outperforms the no *e*_*kl*_ HPM in both the tRNA and Twister ribozyme benchmarks, showing greater sensitivity in the ROC plots in Fig 4. (In the SAM riboswitch benchmark, every method tested is able to fully separate homologous test and decoy sequences.) In the three alignment benchmarks (see table 2), the all-by-all HPM is more accurate than the no *e*_*kl*_ HPM. We conclude *e*_*kl*_’s generally add sensitivity to remote homology search and alignment.

**Table 2:**
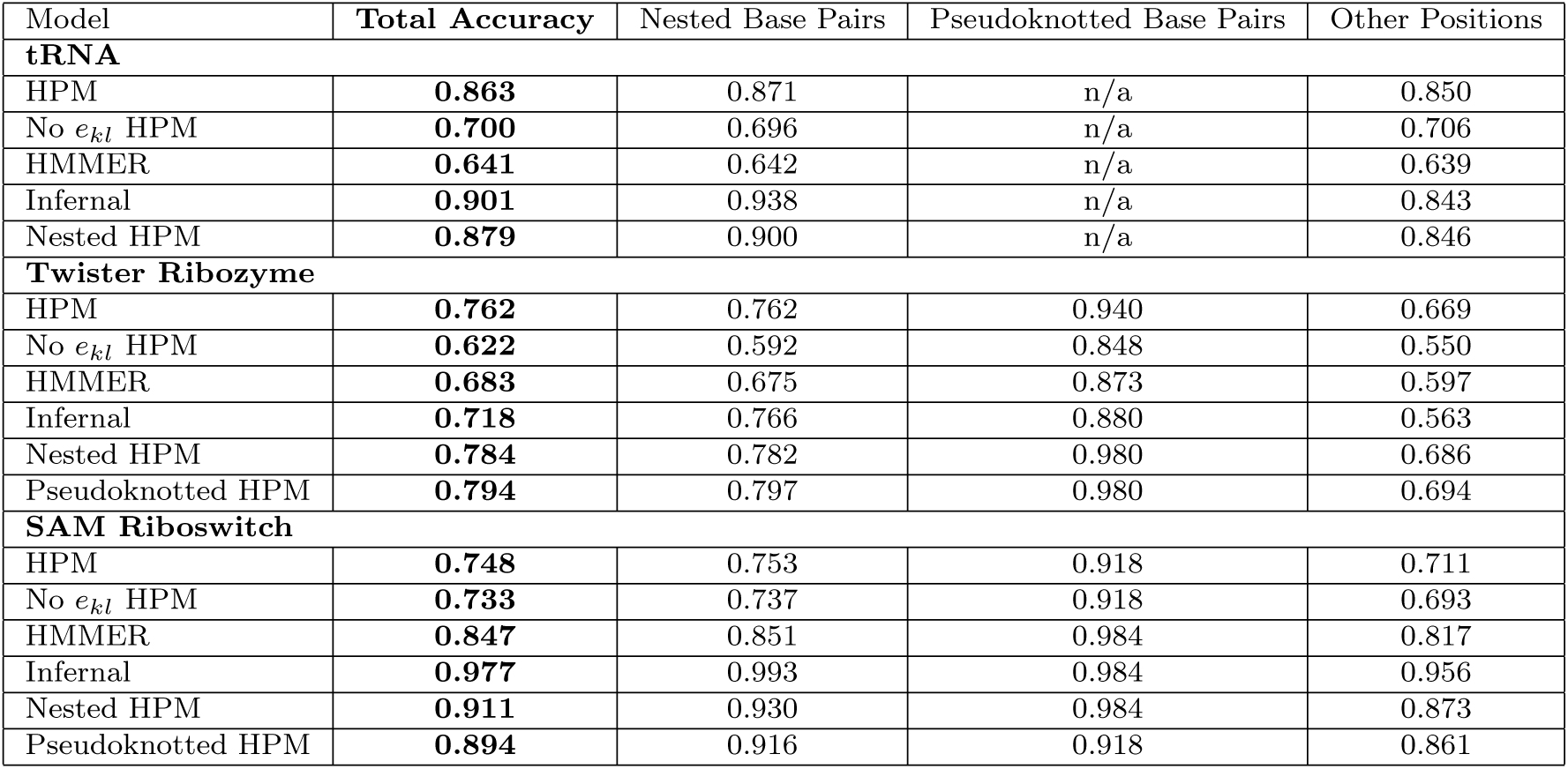
Remote homology alignment benchmark results. Total alignment accuracy is measured as the fraction of residues in consensus columns aligned correctly across all test sequences, relative to the reference alignment. Accuracy across consensus columns with different types of secondary structure annotation is also included.

**Figure 4:**
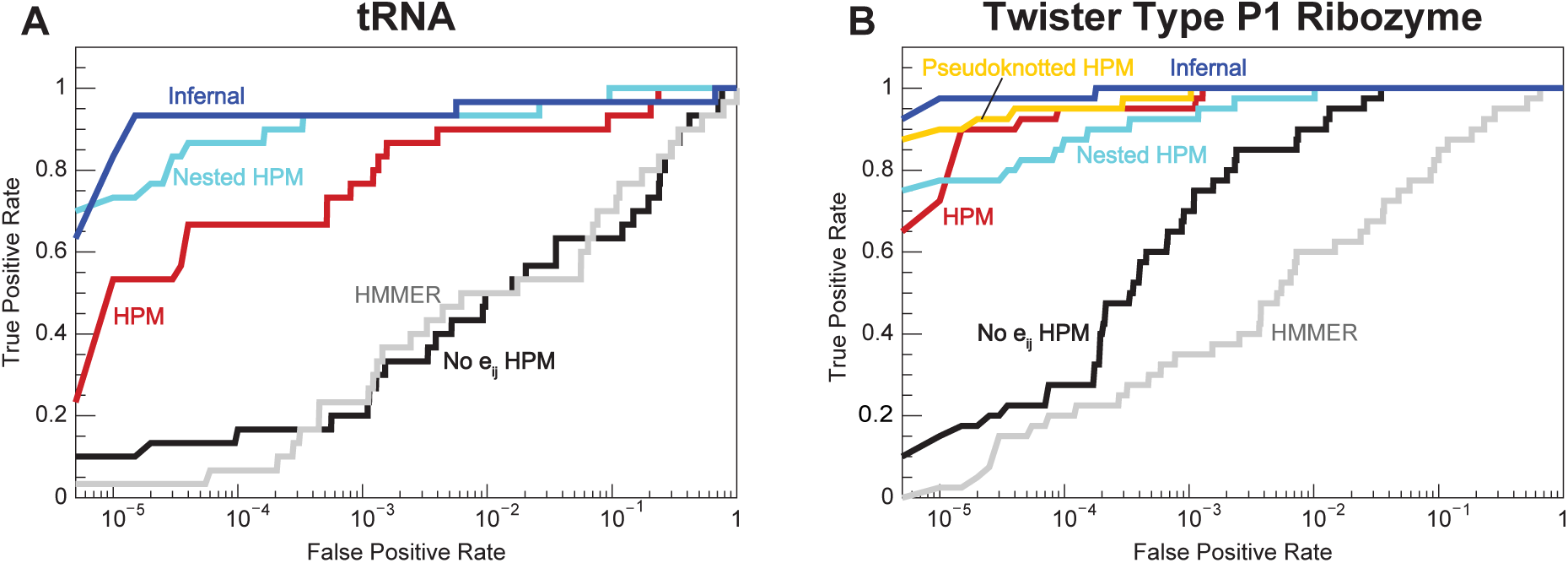
Remote homology scoring benchmark results. Receiver operating characteristic (ROC) plots for the tRNA (A) and Twister type P1 ribozyme (B) benchmarks. The X axis is the fraction of decoys that score higher than a certain threshold (false positive rate), and the Y axis is the fraction of homologous test sequences that score higher than the same threshold (true positive rate, or specificity). Not shown is the ROC plot for the SAM riboswitch benchmark, in which each model perfectly discriminates homologs from decoys.

But does our proof of principle HPM implementation outperform existing methods? We compare the performance of the HPM to HMMER and Infernal, pHMM and pSCFG homology search tools, respectively [46, 7] (gray and blue lines in Fig 4). The HPM generally outperforms HMMER, only falling short in the SAM riboswitch alignment benchmark. Nevertheless, Infernal is usually more sensitive than the HPM, with the only exception being the Twister ribozyme alignment benchmark.

Why is our HPM implementation not outperforming Infernal? One difference between the two methods is model parameterization. Maximum a posteriori pSCFG parameters are calculated analytically from weighted residue frequencies and priors, and Infernal’s choices for weighting and priors have been optimized for homology search and alignment over decades. In contrast, pseudolikelihood maximization of Potts model parameters is an approximation, and it has not previously been used to train homology search and alignment models. To gain more insight, we bypassed pseudolikelihood maximization by creating “masked” HPMs with non-zero *e*_*kl*_ terms restricted to annotated nested base pairs (“nested HPM”, cyan line in Fig 4) or to all annotated disjoint base pairs (“pseudoknotted” HPM, yellow line). For a model with a correlation structure consisting solely of disjoint base pairs, maximum a posteriori Potts model parameters can be estimated directly and analytically like pSCFG parameters by setting *e*_*kl*_ terms to negative log joint probabilities and *h*_*k*_, *h*_*l*_ terms to zero for *k, l* pairs. These “masked” HPMs were almost always more sensitive than the original HPM, with performance is closer to Infernal’s. We conclude that using the pseudolikelihood approximation to parameterize a Potts model is one of the obstacles to using HPMs for homology search.

The improved performance of the masked HPMs relative to the default HPM prompted us to look further into the HPM’s Potts model parameterization. We examine pairwise probabilities of nucleotides under the HPM at annotated base pairs, estimated using Markov chain Monte Carlo. These probabilities do not reflect the observed pairwise frequencies in the training MSA. For instance, Fig 5 shows disagreement between HPM probabilities and training MSA frequencies for an annotated base pair in the SAM riboswitch training MSA. We see many such examples in the SAM riboswitch benchmark.

**Figure 5:**
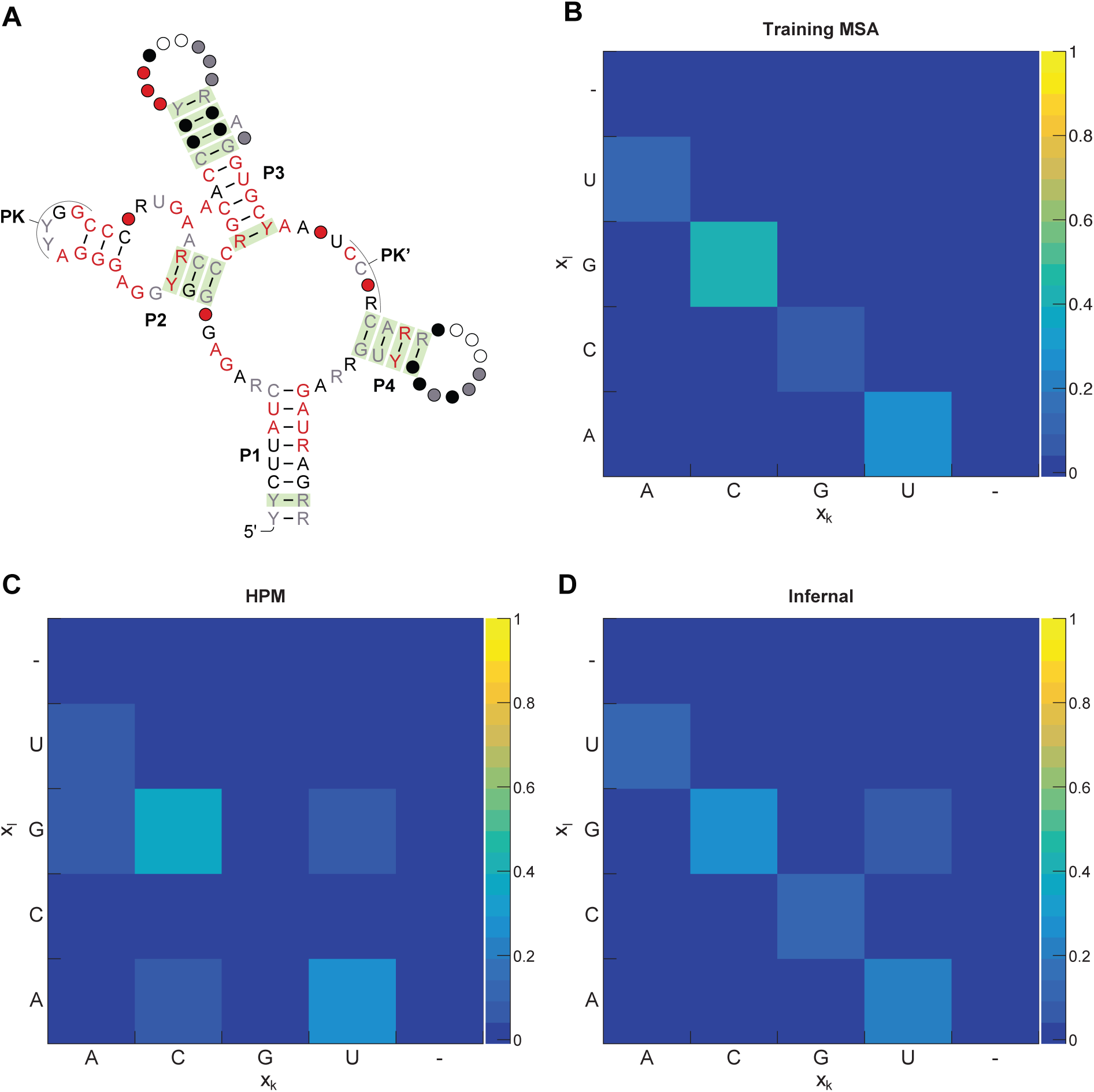
Hidden Potts model emission probabilities do not match training alignment statistics. (A) Rfam consensus structure for the class I SAM riboswitch. Base pairs supported by statistically significant covariation in analysis by R-scape in the RF00162 seed alignment are shaded green [40]. (B) Observed pairwise nucleotide frequencies for one base pair in the P3 stem (sites 52 and 62) in our RF00162 benchmark training alignment (192 sequences). (C) Pairwise nucleotide probabilities at sites 52 and 62 under the RF00162 training HPM. (D) Pairwise nucleotide probabilities at sites 52 and 62 under the RF00162 training Infernal pSCFG.

However, the Gremlin-trained Potts model is accurate in its intended purpose: predicting molecular structure from aligned sequence data. For all three benchmarks, Gremlin’s predicted contacts heavily overlap with annotated base pairs or other observed structural interactions (Fig 6). Future work is needed to optimize Potts model training for remote homology search rather than structure prediction alone.

**Figure 6:**
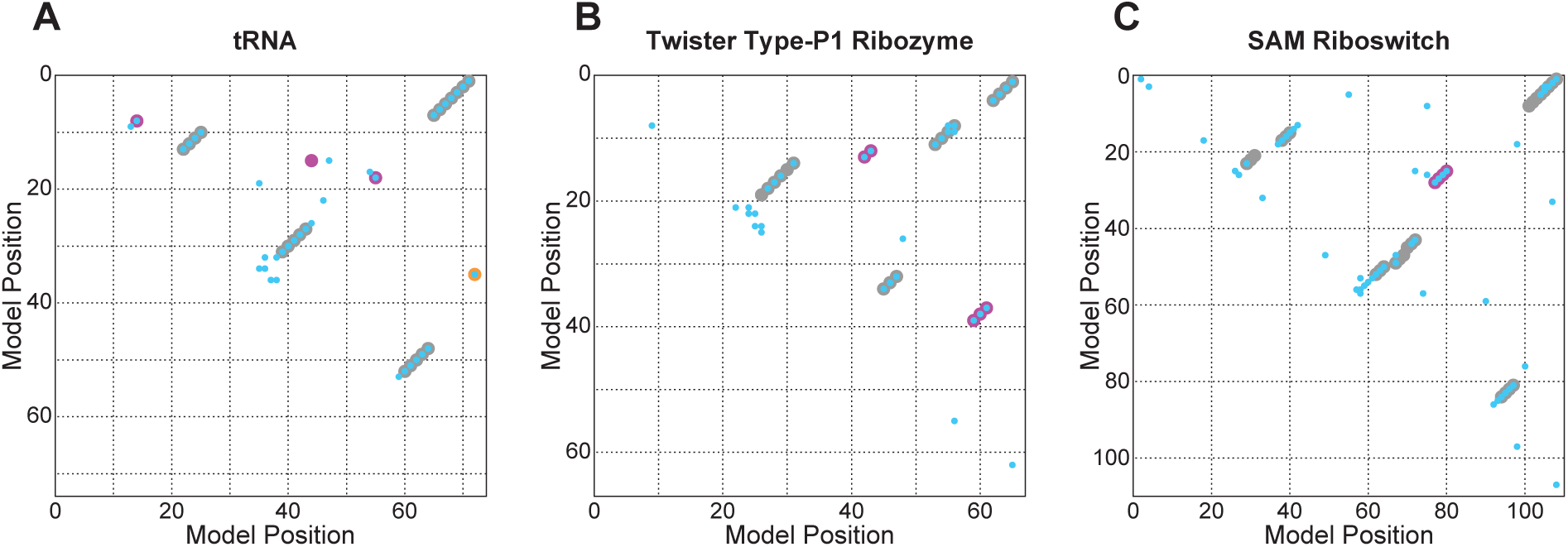
Gremlin-trained Potts models accurately predict 3D RNA contacts. Top 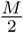 predicted 3D contacts from Gremlin-trained Potts models (cyan) against annotated nested base pairs (gray) and annotated tertiary contacts/pseudoknotted positions (pink) from the tRNA (A), Twister type P1 ribozyme (B), and SAM riboswitch (C) training alignments. For tRNA, annotated tertiary contacts are from a yeast tRNA-phe crystal structure [47]. A non-structural covariation between the tRNA discriminator base and the middle nucleotide in the anticodon (caused by aminoacyl-tRNA synthetase recognition preferences, not conserved base-pairing) is noted in orange [48].

## Discussion

We present hidden Potts models, generative probability models for sequence homology search and alignment. An HPM is a hybrid between a Potts model, successfully used in molecular structure prediction, and a profile HMM, used in protein and nucleic acid homology search and alignment. The hybrid model configuration uses advantages inherent to both models. By using a pHMM’s transition architecture, an HPM can model variable length sequences, a feature previously incompatible with Potts models. The all-by-all pairwise *e*_*kl*_ terms in an HPM’s emission process, as in a Potts model, allow for non-nested structural interactions like pseudoknots to be captured, going beyond the primary-sequence capability of a pHMM and the nested base pair modeling of a pSCFG. Finally, we have developed an efficient algorithm that uses importance sampling to align and score variable length sequences to an HPM.

In initial benchmarking of RNA remote homology search and alignment, we found that HPMs perform promisingly. In experiments where we compare HPMs to HPMs with pairwise *e*_*kl*_ terms forced to zero (thus a level comparison, within the same HPM implementation, of using versus not using correlation information), fully parameterized HPMs produce more accurate alignments and are generally more sensitive for homology searches. HPMs also generally outperformed HMMER, an existing pHMM method. This indicates that hidden Potts models capture useful conserved higher-order correlation structure information in an alignment-capable model. On the other hand, our pilot implementation of HPMs generally underperforms Infernal, an existing pSCFG implementation for capturing nested pairwise correlations in conserved RNA secondary structure. This suggests to us that the gains from being able to model pseudoknots and other non-nested RNA correlations are outweighed by deficiencies in our pilot implementation. Our additional experiments implicate model parameterization is a key issue. When we bypass pseudolikelihood maximization by using “masked” HPMs limited to nonzero maximum likelihood *e*_*kl*_’s only for a disjoint set of base pairs, HPM performance tends much closer to Infernal performance. Additionally, we are not yet using any informative prior distributions in the Potts parameterization, comparable to the use of informative mixture Dirichlet priors in pHMMs and pSCFGs.

Besides improved parameterization, there are other issues to address to make HPMs as the basis for useful homology search and alignment software tools. First, while the importance sampling alignment algorithm is efficient relative to directly enumerating over all possible alignments, it still takes seconds to minutes to align a single sequence. The method will need to be greatly accelerated. Second, while HPMs are able to handle insertions relative to a consensus with a standard affine gap-open, gap-extend probability, they do not explicitly model deletions. It would be desirable to find a cleaner model of deletions. Other work has attempted to combine Potts models with insertions and deletions using techniques based in statistical physics, though this method has not yet been applied to remote homology search and alignment [49]. Third, we only do “glocal” alignment, where a complete match to the HPM consensus model is identified in a possibly longer target sequence. We do not yet see how to do local sequence alignment to an HPM.

Another recent paper uses a different method to align protein and RNA sequences to Potts models [50]. DCAlign, developed by Muntoni et al., uses an algorithm based on message passing rather than importance sampling. They report promising results when comparing alignment accuracy to HMMER and Infernal. However, they do not attempt to score remotely homologous sequences.

Although we chose to do our initial testing with RNA sequence alignments, protein sequence homology search and alignment will also be of interest in future work. Unlike the case for pSCFGs for RNA, the state of the art for protein sequence homology search and alignment remains primary sequence methods such as HMMER and BLAST. Evolutionarily conserved protein structure creates a complex correlation structure unlike the simpler pairwise correlation patterns that dominate RNA analysis; pairwise correlations in protein alignments are difficult to treat by anything simpler than an all-by-all network in which *any* pair of sites is potentially correlated, making protein sequence analysis ripe for Potts models.

## Materials and methods

### HPM software implementation

Our prototype HPM software implementation builds hidden Potts models from multiple sequence alignments, aligns and scores sequences with an HPM using importance sampling, and generates sequences from an HPM using Markov chain Monte Carlo. The software uses functions from HMMER version 4 (in development, see https://github.com/EddyRivasLab/hmmer/tree/h4-develop) and the Easel sequence library (see https://github.com/EddyRivasLab/easel).

To train Potts models, we use a modified version of the Gremlin C++ software package [11], included with our HPM implementation. Code is available at https://github.com/gwwilburn/HPMHomology, with the version used to produce results for this paper archived at https://github.com/gwwilburn/HPMHomology/commits/master under tag 845ef8e. A tarball of our software is also included in S1 Code.

### HPM training

HPMs are trained by our program `hpmbuild`. *P*_*t*_ transition parameters and *P*_*i*_ insert emission parameters are taken from a pHMM trained by `hmmbuild` (HMMER v4). *P*_*m*_ Potts emission parameters are taken from a model trained by Gremlin C++ (version 1.0) modified to use Henikoff position-based sequence weights in the training procedure [51], the same relative weighting scheme used by HMMER.

### Masked HPM training

So-called masked HPMs, with only a disjoint subset of *e*_*kl*_ terms allowed to be non-zero, are trained by our program `hpmbuild_masked`. Annotated base pairs are determined by the secondary structure consensus line in an input Stockholm MSA file. Potts model parameters are negative log likelihoods estimated from weighted counts with a +1 Laplace pseudocount. Henikoff position-based sequence weights are used [51]. For sites belonging to annotated base pairs, *e*_*kl*_’s are trained using the weighted pairwise nucleotide frequencies in the input MSA; these sites’ *h*_*k*_ terms set to zero. For unpaired sites, *h*_*k*_ terms are trained using the weighted single-site nucleotide frequencies, and the corresponding *e*_*kl*_’s are set to zero.

### Aligning and scoring sequences with an HPM

Sequences are aligned to and scored by an HPM with our program `hpmscoreIS`. This program takes an HPM, a pHMM with the same number of match states, and a sequence file as input. It returns the unnormalized log odds scores for each sequence estimated using importance sampling with the pHMM as a proposal distribution. Additionally, aligned sequences are outputted to a Stockholm MSA file. For each input sequence, 1 million alignments are sampled from the pHMM.

### Estimation of single-site and pairwise probabilities under an HPM

Probabilities are estimated empirically from 10,000 synthetic sequences emitted from an HPM using our program `hpmemit`.

We use Markov chain Monte Carlo to emit match sequence 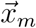 from the Potts model. Each sequence is generated independently using the Metropolis-Hastings algorithm with a burn-in period of 1000 steps.

To create the HPM path 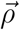, we perform a random walk through the HPM’s state transitions. Given 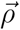, we emit insertion sequence 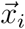 by drawing residues independently from 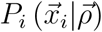.

### Software and database versions used

For building pHMMs, we use HMMER 4 (in development). The code is available at https://github.com/EddyRivasLab/hmmer, inside the branch `h4-develop`. The specific version of code we use is archived under tag 7994cc7 at https://github.com/EddyRivasLab/hmmer/commits/h4-develop. The software is also included in S1 Code.

We use HMMER 4 functions to create programs `hmmalign_uniglocal` and `hmmscore_uniglocal` that glocally score and align sequences with pHMMs, respectively. These programs are included in the HPM prototype software implementation.

For analysis with pSCFGs, we use Infernal version 1.1.3 [7]. The code is available at http://eddylab.org/infernal/ and included with S1 Code. Rfam version 14.1 is used for the SAM riboswitch anecdote [36].

For consensus RNA structure prediction by comparative sequence analysis, we used CaCoFold in version 1.4.0 of the R-scape software package [40, 41].

### Benchmark datasets

Training and test alignments used for each anecdote are included as supplementary material with S1 Code. Benchmark procedures follow methods in [44].

### Analysis of alignments

RNA multiple sequence alignments were visualized using the Ralee RNA alignment editor in Emacs [52].

## Supporting information

**S1 Appendix. Derivation of importance sampling alignment algorithm** (PDF).

**S1 Code. Contains the HPM software implementation, versions of HMMER and Infernal used in benchmarking, the Easel sequence library, and benchmark datasets**. (ZIP)

## Acknowledgments

We thank Elena Rivas, Tom Jones, Nick Carter, and Sergey Ovchinnikov for insightful discussions throughout the project. Additionally, we thank Eric Nawrocki and members of the Eddy and Rivas labs for comments on the manuscript. Some ideas in this work were conceived at workshops hosted at the Centro de Ciencias de Benasque Pedro Pascual (Benasque, Spain), and the Aspen Center for Physics (supported by National Science Foundation grant PHY-1066293). Computations were run on the Cannon cluster, supported by the Harvard FAS Division of Science’s Research Computing Group. Our work was funded by the Howard Hughes Medical Institute and by the National Human Genome Research Institute of the National Institutes of Health (award R01-HG009116). The content is solely the responsibility of the authors and does not necessarily represent the official views of the National Institutes of Health.

## References

1. Durbin R, Eddy SR, Krogh A, Mitchison GJ. Biological Sequence Analysis: Probabilistic Models of Proteins and Nucleic Acids. Cambridge UK: Cambridge University Press; 1998.

2. Weisman CM, Murray AW, Eddy SR. Many but Not All Lineage-Specific Genes Can Be Explained by Homology. Detection Failure. biorXiv 968420v2 [Preprint]. 2020 [Cited 11 June 2020]. Available from: https://www.biorxiv.org/content/10.1101/2020.02.27.968420v2 doi: 10.1101/2020.02.27.968420

3. Altschul SF, Gish W, Miller W, Myers EW, Lipman DJ. Basic Local Alignment Search Tool. J Mol Biol. 1990;215:403–410.

4. Haussler D, Krogh A, Mian IS, Sjolander K. Protein Modeling Using Hidden Markov Models: Analysis of Globins. In: Proceedings of the Twenty-Sixth Hawaii International Conference on System Sciences; 1993. p. 792–802.

5. Eddy SR. Profile Hidden Markov Models. Bioinformatics. 1998;14:755–763.

6. Eddy SR, Durbin R. RNA Sequence Analysis Using Covariance Models. Nucl Acids Res. 1994;22:2079–2088.

7. Nawrocki EP, Eddy SR. Infernal 1.1: 100-fold Faster RNA Homology Searches. Bioinformatics. 2013;29:2933–2935.

8. Lapedes AS, Giraud BG, Liu LC, Stormo GD. A Maximum Entropy Formalism for Disentangling Chains of Correlated Sequence Positions. Lecture Notes-Monograph Series, Statistics in Molecular Biology and Genetics. 1999;33:236–256.

9. Weigt M, White RA, Szurmant H, Hoch JA, Hwa T. Identification of Direct Residue Contacts in Protein–Protein Interaction by Message Passing. Proc Natl Acad Sci USA. 2009;106:67–72.

10. Morcos F, Pagnani A, Lunt B, Bertolino A, Marks DS, Sander C, et al. Direct-Coupling Analysis of Residue Coevolution Captures Native Contacts Across Many Protein Families. Proc Natl Acad Sci USA. 2011;108:E1293–E1301.

11. Kamisetty H, Ovchinnikov S, Baker D. Assessing the Utility of Coevolution-based Residue–Residue Contact Predictions in a Sequence-and Structure-Rich Era. Proc Natl Acad Sci USA. 2013;110(39):15674–15679.

12. Ekeberg M, Lövkvist C, Lan Y, Weigt M, Aurell E. Improved Contact Prediction in Proteins: Using Pseudolikelihoods to Infer Potts Models. Physical Review E. 2013;87(1):012707.

13. De Leonardis E, Lutz B, Ratz S, Cocco S, Monasson R, Schug A, et al. Direct-Coupling Analysis of Nucleotide Coevolution Facilitates RNA Secondary and Tertiary Structure Prediction. Nucl Acids Res. 2015;43:10444–10455.

14. Weinreb C, Riesselman AJ, Ingraham JB, Gross T, Sander C, Marks DS. 3D RNA and Functional Interactions from Evolutionary Couplings. Cell. 2016;165:963–975.

15. White JV, Muchnik I, Smith TF. Modeling Protein Cores with Markov Random Fields. Math Biosci. 1994;124:149–179.

16. Lathrop RH, Smith TF. Global Optimum Protein Threading with Gapped Alignment and Empirical Pair Score Functions. J Mol Biol. 1996;255:641–645.

17. Thomas J, Ramakrishnan N, Bailey-Kellogg C. Graphical Models of Residue Coupling in Protein Families. IEEE/ACM Trans Comp Biol Bioinf. 2008;5:183–197.

18. Liu Y, Carbonell J, Gopalakrishnan V, Weigele P. Conditional Graphical Models for Protein Structural Motif Recognition. J Comput Biol. 1996;255:641–645.

19. Menke M, Berger B, Cowen L. Markov Random Fields Reveal an N-Terminal Double Beta-Propeller Motif as Part of a Bacterial Hybrid Two-Component Sensor System. Proc Natl Acad Sci USA. 2010;107:4069–4074.

20. Peng J, Xu J. A Multiple-Template Approach to Protein Threading. Proteins. 2010;79:1930–1939.

21. Daniels NM, Hosur R, Berger B, Cowen L. SMURFLite: Combining Simplified Markov Random Fields with Simulated Evolution Improves Remote Homology Detection for Beta-Structural Proteins into the Twilight Zone. Bioinformatics. 2012;28:1216–1222.

22. Ovchinnikov S, Kamisetty H, Baker D. Robust and Accurate Prediction of Residue-Residue Interactions across Protein Interfaces Using Evolutionary Information. eLife. 2014;113:e02030.

23. Bitbol AF, Dwyer RS, Colwell LJ, Wintergreen NS. Inferring Interaction Partners from Protein Sequences. Proc Natl Acad Sci USA. 2016;106:67–72.

24. Gueudre T, Baldassi C, Zamparo M, Weigt M, Pagnani A. Simultaneous Identification of Specifically Interacting Paralogs and Interprotein Contacts by Direct Coupling Analysis. Proc Natl Acad Sci USA. 2016;113:12185–12191.

25. Cong Q, Anishchenko I, Ovchinnikov S, Baker D. Protein Interaction Networks Revealed by Proteome Coevolution. Science. 2019;365:185–189.

26. Cheng RR, Nordesjo O, Hayes RL, Levine H, Flores SC, Onuchic JN, et al. Connecting the Sequence-Space of Bacterial Signaling Proteins to Phenotypes Using Coevolutionary Landscapes. Mol Biol Evol. 2016;33:3054–3064.

27. Figliuzzi M, Jacquier H, Schug A, Tenaillon O, Weigt M. Coevolutionary Landscape Inference and the Context-Dependence of Mutations in Beta-Lactamase TEM-1. Mol Biol Evol. 2016;33:268–280.

28. Levy RM, Haldane A, Flynn WF. Potts Hamiltonian Models of Protein Co-variation, Free Energy Landscapes, and Evolutionary Fitness. Curr Opin Struct Biol. 2017;43:55–62.

29. Hopf TA, Ingraham JB, Poelwijk FJ, Sharfe CP, Springer M, Sander C, et al. Mutation Effects Predicted from Sequence Co-variation. Nature Biotechnology. 2017;35:128–135.

30. Salinas VH, Ranganathan R. Coevolution-Based Inference of Amino Acid Interactions Underlying Protein Function. eLife. 2018;7:e34300.

31. Graner F, Glazier JA. Simulation of Biological Cell Sorting Using a Two-Dimensional Extended Potts Model. Physical Review Letters. 1992;69:2013–2016.

32. Schneidmann E, II MJB, Segev R, Bialek W. Weak Pairwise Correlations Imply Strongly Correlated Network States in a Neural Population. Nature. 2007;440:1007–1012.

33. Chiu DKY, Kolodziejczak T. Inferring Consensus Structure from Nucleic Acid Sequences. Comput Applic Biosci. 1991;7:347–352.

34. Gutell RR, Power A, Hertz GZ, Putz EJ, Stormo GD. Identifying Constraints on the Higher-Order Structure of RNA: Continued Development and Application of Comparative Sequence Analysis Methods. Nucl Acids Res. 1992;20:5785–5795.

35. Finn RD, Bateman A, Clements J, Coggill P, Eberhardt RY, Eddy SR, et al. Pfam: The Protein Families Database. Nucl Acids Res. 2014;42:D222–D230.

36. Kalvari I, Argasinska J, Quinones-Olvera N, Nawrocki EP, Rivas E, Eddy SR, et al. Rfam 13.0: Shifting to a Genome-Centric Resource for Non-Coding RNA Families. Nucl Acids Res. 2018;46:D335–D342.

37. Besag J. Efficiency of Pseudolikelihood Estimation for Simple Gaussian Fields. Biometrika. 1977;64:616–618.

38. Eddy SR. Multiple Alignment Using Hidden Markov Models. In: Rawlings C, Clark D, Altman R, Hunter L, Lengauer T, Wodak S, editors. Proc. Third Int. Conf. Intelligent Systems for Molecular Biology. Menlo Park, CA: AAAI Press; 1995. p. 114–120.

39. Schneider TD, Stormo GD, Gold L, Ehrenfeucht A. Information Content of Binding Sites on Nucleotide Sequences. J Mol Biol. 1986;188:415–431.

40. Rivas E, Clements J, Eddy SR. A Statistical Test for Conserved RNA Structure Shows Lack of Evidence for Structure in lncRNAs. Nature Methods. 2017;14:45–48.

41. Rivas E. RNA Structure Prediction Using Positive and Negative Evolutionary Information. biorXiv 933952v2 [Preprint]. 2020 [Cited 11 June 2020]. Available from: https://www.biorxiv.org/content/10.1101/2020.02.04.933952v2 doi: 10.1101/2020.02.04.933952

42. Sprinzl M, Horn C, Brown M, Ioudovitch A, Steinberg S. Compilation of tRNA Sequences and Sequences of tRNA Genes. Nucl Acids Res. 1998;26:148–153.

43. Roth A, Weinberg Z, Chen AG, Kim PB, Ames TD, Breaker RR. A Widespread Self-Cleaving Ribozyme Class is Revealed by Bioinformatics. Nat Chem Biol. 2014;10:56–60.

44. Nawrocki EP, Eddy SR. Query-Dependent Banding (QDB) for Faster RNA Similarity Searches. PLOS Comput Biol. 2007;3:e56.

45. Montange R, Batey RT. Structure of the S-adenosylmethionine riboswitch regulatory mRNA element. Nature. 2006;441:1172–1175.

46. Eddy SR. Accelerated profile HMM searches. PLOS Comp Biol. 2011;7:e1002195.

47. Westhof E, Sundaralingam M. Restrained Refinement of the Monoclinic Form of Yeast Phenylalanine Transfer RNA. Temperature Factors and Dynamics, Coordinated Waters, and Base-Pair Propeller Twist Angles. Biochemistry. 1986;25:4868–4878.

48. Crothers DM, Seno T, Soll DG. Is There a Discriminator Site in tRNA? Proc Natl Acad Sci USA. 1972;69:3063–3067.

49. Kinjo AR. A Unified Statistical Model of Protein Multiple Sequence Alignment Integrating Direct Coupling and Insertions. Biophysics and Physicobiology. 2016;13:45–62.

50. Muntoni AP, Pagnani A, Weigt M, Zamponi F. Aligning Biological Sequences by Exploiting Residue Conservation and Coevolution; 2020.

51. Henikoff S, Henikoff JG. Protein Family Classification Based on Searching a Database of Blocks. Genomics. 1994;19:97–107.

52. Griffiths-Jones S. RALEE–RNA ALignment Editor in Emacs. Bioinformatics. 2005;21:257–259.

