## Supplemental Appendix S1 for "Remote homology search with hidden Potts models"

### Importance Sampling Alignment Algorithm

Technical appendix accompanying Wilburn & Eddy (2020),

### 1 Introduction

When aligning or scoring sequences with a fully parameterized hidden Potts model, we use an approximate method called importance sampling. Unlike with profile HMMs (pHMMs) and profile SCFGs (pSCFGs) [1, 2, 3], HPMs are incompatible with dynamic programming algorithms that complete in polynomial time. In importance sampling, we randomly sample possible alignments from a pHMM and "propose" them to an HPM, which rescores them. Below, we review background of the technique and our justification for its use in HPM homology search and alignment. The derivations below were inspired by [4] and [5].

### 2 Background and derivation

#### 2.1 Monte Carlo estimation

Consider a discrete variable $a$. We are given a normalized probability distribution $P(a)$ such that $\sum_a P(a) = 1$. Now consider a function $f(a)$. The expectation value of $f(a)$ under the distribution $P(a)$ is given by:

$$F \equiv \langle f(a) \rangle_{a \sim P} = \sum_a P(a) f(a) \tag{1}$$

If the sum in equation 1 is difficult to evaluate, we can approximate this expectation value using **Monte Carlo estimation**. If we can sample $R$ independent values $\left\{a^{(r)}\right\}_{r=1}^{R}$ from $P(a)$, the expectation value is approximated as:

$$F \approx \hat{f} \equiv \frac{1}{R} \sum_{r=1}^{R} f\left(a^{(r)}\right) \tag{2}$$

It can be shown:

- The expectation value $\langle \hat{f} \rangle$ is equal to $F$ for any number of samples $R$.
- If $f(x)$ has variance $\sigma_f^2$, the variance of $\hat{f}$ is $\frac{\sigma_f^2}{R}$.

For example, Monte Carlo estimation can be used to calculate marginal probability distributions. Consider a joint probability distribution $P(a, b)$. The marginal probability of $b$ is:

$$P(b) = \sum_a P(a, b) \tag{3}$$

Using Bayes' theorem, we can rewrite the argument of the sum as:

$$P(b) = \sum_a P(b|a) P(a) \tag{4}$$

In this case, $P(b|a)$ is analogous to $f(a)$ in our first example. We can approximate $P(b)$ via Monte Carlo estimation.

$$P(b) \approx \frac{1}{R} \sum_{r=1}^{R} P\left(b|a^{(r)}\right) \tag{5}$$

#### 2.2 Importance sampling

We may not want to sample from $P(a)$ for any number of reasons. For one, it may not be possible to generate samples from $P(a)$. A more subtle issue may be that the value of $f(a)$ is negligibly small for all but a small region of $a$ space, meaning most of the samples $\left\{a^{(r)}\right\}$ contribute almost nothing to the sum in equation 2.

Consider another distribution $Q(a)$ from which we are able to sample. We will call $Q(a)$ the *proposal distribution.* So long as $Q(a)$ is nonzero for every $a$ at which the product of $f(a)$ and $P(a)$ is nonzero, we can use it to estimate $\langle f(a)\rangle_{(a\sim P)}$.

First, we multiply the right hand side of equation 1 by one.

$$F = \sum_{a} P(a) f(a) \frac{Q(a)}{Q(a)} \tag{6}$$

Rearranging, we have an expectation value of the product of multiple terms under $Q(a)$.

$$F = \sum_{a} Q(a) \frac{P(a)}{Q(a)} f(a) = \langle \frac{P(a)}{Q(a)} f(a) \rangle_{a\sim Q} \tag{7}$$

By drawing samples from $Q(a)$, we can approximate the expectation value as:

$$F \approx \frac{1}{R} \sum_{r=1}^{R} \frac{P\left(a^{(r)}\right)}{Q\left(a^{(r)}\right)} f\left(a^{(r)}\right) \equiv \hat{f}_Q \tag{8}$$

Introducing the *importance weight* $w\left(a^{(r)}\right) \equiv \frac{P\left(a^{(r)}\right)}{Q\left(a^{(r)}\right)}$, our estimation becomes:

$$\hat{f}_Q = \sum_{r=1}^{R} w\left(a^{(r)}\right) f\left(a^{(r)}\right) \tag{9}$$

The importance weight corrects for the fact that we are sampling from $Q(a)$ while trying to calculate an expectation value under $P(a)$. For values of $a$ where $Q(a)$ is larger than $P(a)$, we will generate more samples using importance sampling than we would if sampling from $P(a)$, but these samples will be downweighted. For values of $a$ where $Q(a)$ is smaller than $P(a)$, we will generate fewer samples under importance sampling, but the samples will be upweighted.

In the case of marginalization, given a proposal distribution $Q(a)$ from which to sample, the marginal probability can be approximated as:

$$P(b) \approx \frac{1}{R} \sum_{r=1}^{R} P\left(b|a^{(r)}\right) \frac{P\left(a^{(r)}\right)}{Q\left(a^{(r)}\right)} \tag{10}$$

#### 2.3 Is importance sampling accurate?

In order for importance sampling to work, we need the value of our estimator $\hat{f}_Q$ to converge to the value of $F$. We can show $\hat{f}_Q$ is an unbiased estimator of $F$, as the expectation value of $\hat{f}_Q$ is $F$ for any $R$. Mackay Exercise 29.1 shows this for the continuous case [4], but we will show it for the discrete case here. The expectation value of $\hat{f}_Q$ is given by:

$$\langle \hat{f}_Q \rangle = \langle \frac{1}{R} \sum_{r=1}^{R} f\left(a^{(r)}\right) w\left(a^{(r)}\right) \rangle \tag{11}$$

As each sample $a^{(r)}$ is drawn independently from $Q(a)$, we can rewrite the expectation of the sum as a sum of expectation values.

$$\langle \hat{f}_Q \rangle = \frac{1}{R} \sum_{r=1}^{R} \langle f\left(a^{(r)}\right) w\left(a^{(r)}\right) \rangle = \langle f(a)\, w(a) \rangle \tag{12}$$

Under the distribution $Q(a)$, the right hand side can be written as a sum over all possible values of $a$.

$$\langle \hat{f}_Q \rangle = \sum_{a} f(a)\, w(a)\, Q(a) \tag{13}$$

Using the definition of the importance weight, we see that $\langle \hat{f}_Q \rangle$ is equal to $F$.

$$\langle \hat{f}_Q \rangle = \sum_{a} f(a) \frac{P(a)}{Q(a)} Q(a) = \sum_{a} f(a)\, P(a) = F \tag{14}$$

Note that the above does not necessarily hold true if $Q(a)$ is zero for certain values of $a$ for which the product of $P(a)$ and $f(a)$ is nonzero!

### 3 Applying importance sampling to HPM scoring and alignment

of

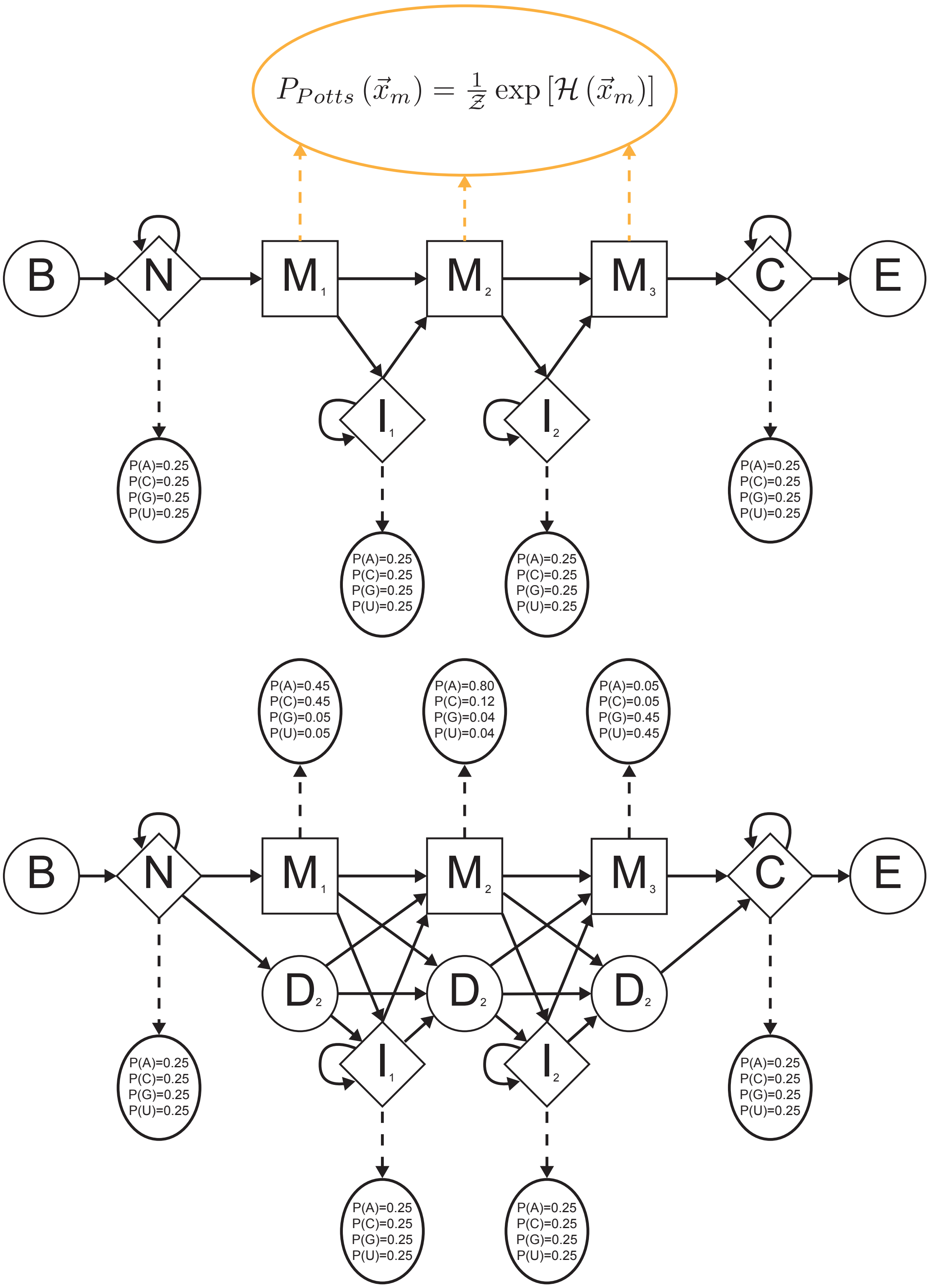


Figure 1: Model architecture of an 3-site HPM (top) and corresponding pHMM (bottom).

Given an unaligned sequence $\vec{x}$ and a fully parameterized hidden Potts model over all possible sequences and alignments (or "paths"), $P(\vec{x}, \text{ali})$, we wish to calculate the marginal distribution $P(\vec{x})$ when scoring sequences.

$$P(\vec{x}) = \sum_{\text{ali}} P(\vec{x}, \text{ali}) \tag{15}$$

We will use importance sampling. For our proposal distribution $Q(\text{ali})$, we require the following qualities:

1. We can generate samples from $Q$ in polynomial time.
2. The probability mass of $Q(\text{ali})$ is concentrated in a region of probability space in which $P(\vec{x}, \text{ali})$ is generally non-negligible.
3. $Q(\text{ali})$ is non-zero everywhere $P(\vec{x}, \text{ali})$ is non-zero.

As our proposal distribution, we will generate alignments from the probability distribution of alignments under a pHMM conditioned on sequence $\vec{x}$, as it satisfies the above criteria. First, samples can be generated efficiently using dynamic programming algorithms. Given a sequence $\vec{x}$ with $L$ characters, stochastic traceback generates a pHMM path in $\mathcal{O}(L)$ time [3]. At initiation, the stochastic traceback needs the dynamic programming matrix calculated by the forward algorithm, which takes $\mathcal{O}(LM)$ time to complete, where $M$ is the number of match states in the profile HMM. However, the forward algorithm only needs to be run once per sequence prior to generating paths.

Second, we have observed that pHMMs generally align noncoding RNAs *pretty well* despite only capturing primary sequence conservation (see Table 2 in main text). It is our experience that pHMMs align plausible homologs mostly right, even in the case of highly structured RNAs such as tRNA.

Third, as the transition structure of the HPM model architecture is derived from a pHMM, there are generally no paths with zero probability under a pHMM and non-zero probability under an HPM.

So our importance sampling estimate for $P\left(\vec{x}\right)$ becomes:

$$P\left(\vec{x}\right) \approx \frac{1}{R}\sum_{r=1}^{R}\frac{P\left(\vec{x},\text{ali}^{(r)}\right)}{P_{HMM}\left(\text{ali}^{(r)}|\vec{x}\right)} \tag{16}$$

However, there is a critical issue: a path under an HPM, $\vec{\rho}$, is different than a path under a pHMM, $\vec{\pi}$. pHMM paths handle match columns in an MSA with two types of states: match states, which emit characters; and delete states, which allow for the absence of a character in a match column. HPM paths only use match states to represent MSA match columns, with deletions represented as an extra character emitted by match states. As such, an HPM path $\vec{\rho}$ does not correspond to a single pHMM path $\vec{\pi}$, but rather $2^M$ possible pHMM paths. However, the converse is true: a given $\vec{\pi}$ corresponds to one and only one $\vec{\rho}$.

As the stochastic traceback algorithm samples pHMM paths from $P_{HMM}\left(\vec{\pi}|\vec{x}\right)$, we are sampling paths from one sample space (pHMM space) and proposing them to a probability distribution defined over a different sample space (HPM space). In order to use $P_{HMM}\left(\vec{\pi}|\vec{x}\right)$, we need to define HPMs and pHMMs using the same variables.

#### 3.1 Describing HPMs in terms of four variables

We usually think of generative sequence homology models as being distributions over two variables: the observed data, corresponding to a biological sequence in our case; and a path, representing columns in a multiple sequence alignment. Equivalently, an HPM can be cast as a probability distribution over four variables:

- $\vec{x}$: an unaligned, observed sequence.
- $\vec{x}_m$: characters emitted from match states (including deletion characters).
- $\vec{x}_i$: characters emitted from intermediate insert states ($I_k$, where $k$ denotes the index of the preceding match column) and flanking insert states ($N$, $C$).
- $\vec{\rho}$: An HPM-style path through the model.

The joint probability of these four variables under the HPM is the product of four terms.

$$P\left(\vec{x}, \vec{x}_m, \vec{x}_i, \vec{\rho}\right) = P\left(\vec{x} | \vec{x}_m, \vec{x}_i, \vec{\rho}\right) P\left(\vec{x}_m, \vec{x}_i, \vec{\rho}\right) \tag{17}$$

The first term on the right hand side of equation 17 is the *compatibility function.*

$$P\left(\vec{x} | \vec{x}_m, \vec{x}_i, \vec{\rho}\right) = \delta\left(\vec{x}, \vec{x}_m, \vec{x}_i, \vec{\rho}\right) \tag{18}$$

The compatibility function is 1 if $\vec{x}_m$, $\vec{x}_i$, and $\vec{\rho}$ are *compatible* with unaligned sequence $\vec{x}$, and it is 0 otherwise. Compatibility means that $\vec{x}_m$, $\vec{x}_i$, and $\vec{\rho}$ can be combined and *dealigned* to produce unaligned sequence $\vec{x}$.

For instance, given the 3-state HPM in figure 1 and sequence $\vec{x}$ = "ACU", $\vec{x}_m$ = "AU", $\vec{x}_i$ = 'C', and $\vec{\rho} = B \rightarrow N \rightarrow M_1 \rightarrow I_1 \rightarrow M_2 \rightarrow M_3 \rightarrow C \rightarrow E$ are compatible with $\vec{x}$. However, $\vec{x}_m$ = "AC-", $\vec{x}_i$ = 'U', and $\vec{\rho} = B \rightarrow N \rightarrow M_1 \rightarrow I_1 \rightarrow M_2 \rightarrow M_3 \rightarrow C \rightarrow E$ are not compatible with $\vec{x}$, as they form unaligned sequence "AUC" rather than"ACU".

For a given $\vec{x}$, many possible combinations of $\vec{x}_m$, $\vec{x}_i$, and $\vec{\rho}$ yield a compatibility value of 1. However, given $\vec{x}_m$, $\vec{x}_i$, and $\vec{\rho}$, only one $\vec{x}$ is compatible.

The second term on the right hand side of equation 17 factorizes into three terms.

$$P\left(\vec{x}_m, \vec{x}_i, \vec{\rho}\right) = P_m\left(\vec{x}_m\right) P_i\left(\vec{x}_i|\vec{\rho}\right) P_t\left(\vec{\rho}\right) \tag{19}$$

$P_m\left(\vec{x}_m\right)$ is the Potts distribution, $P_i\left(\vec{x}_i|\vec{\rho}\right)$ is the HPM's site-independent insert emission probability model, and $P_t\left(\vec{\rho}\right)$ is the HPM's transition probability model.

Here are the steps to generate an unaligned sequence $\vec{x}$ under an HPM using this four variable scheme.

1. Generate a path $\vec{\rho}$ from $P_t\left(\vec{\rho}\right)$.

    - This is a Markov process; we perform a random, memoryless walk between the states.

- Knowing $\vec{\rho}$ does not mean we know where deletion characters are.
- However, $\vec{\rho}$ does tell us which insert states (both flanking and intermediate) are visited and emit characters.

2. Generate $\vec{x}_i$ from $P_i\left(\vec{x}_i|\vec{\rho}\right)$.

   - This process is the independent emission of characters from the insert states visited in $\vec{\rho}$, just as in a pHMM.

3. Generate $\vec{x}_m$ from the Potts distribution, $P_m\left(\vec{x}_m\right)$.

   - Note that this distribution is not conditioned on $\vec{\rho}$. Every HPM path passes through every match state.
   - All match characters, including deletion characters, are generated in a single emission from the Potts model.

4. Choose $\vec{x}$ from $\delta\left(\vec{x}, \vec{x}_m, \vec{x}_i, \vec{\rho}\right)$.

   - Once we have $\vec{x}_m$, $\vec{x}_i$, and $\vec{\rho}$, only one $\vec{x}$ is possible.

To obtain the probability of $\vec{x}$ under an HPM, we need to marginalize over the other 3 variables.

$$P\left(\vec{x}\right) = \sum_{\vec{x}_m}\sum_{\vec{x}_i}\sum_{\vec{\rho}} P\left(\vec{x}, \vec{x}_m, \vec{x}_i, \vec{\rho}\right) \tag{20}$$

Writing this in terms of the compatibility function, we have:

$$P\left(\vec{x}\right) = \sum_{\vec{x}_m}\sum_{\vec{x}_i}\sum_{\vec{\rho}} \delta\left(\vec{x}, \vec{x}_m, \vec{x}_i, \vec{\rho}\right) P\left(\vec{x}_m, \vec{x}_i, \vec{\rho}\right) \tag{21}$$

We can now approximate $P(\vec{x})$ by sampling $\vec{x}_m$, $\vec{x}_i$, and $\vec{\rho}$ from a pHMM. However, we first need to formulate a pHMM under the four variable scheme.

#### 3.2 Redefining profile HMMs in terms of four variables

Just as with HPMs, profile HMMs can be recast using $\vec{x}$, $\vec{x}_m$, $\vec{x}_i$, and $\vec{\rho}$. $\vec{x}_m$ will still contain deletion characters in this formulation. For importance sampling, we will need to define $P_{HMM}(\vec{\pi}|\vec{x})$ in the four variable scheme.

##### 3.2.1 Relating pHMM paths to HPM paths

First, we must define the HPM path $\vec{\rho}$ in terms of the pHMM path $\vec{\pi}$. The HPM path is a modification of a pHMM path in which delete and match states corresponding to a given MSA column are combined into one state. Given $\vec{\rho}$, we know a path through match columns, but not whether a character or a deletion is present in a match column.

A specific $\vec{\rho}$ corresponds to $2^M$ possible pHMM paths $\vec{\pi}$, as each HPM match state corresponds to a pHMM match or delete state. However, given $\vec{\pi}$, we know $\vec{\rho}$: $P_{HMM}(\vec{\rho}|\vec{\pi})$ is 1 for only one $\vec{\rho}$, and 0 otherwise.

The probability of $\vec{\rho}$ under a pHMM is given by the sum of the probabilities of all $2^M$

$\vec{\pi}$'s that correspond to that $\vec{\rho}$.

$$P_{HMM}\left(\vec{\rho}\right) = \sum_{\vec{\pi}} P_{HMM}\left(\vec{\rho}|\vec{\pi}\right) P_{HMM}\left(\vec{\pi}\right) = \sum_{\vec{\pi}\in\vec{\rho}} P_{HMM}\left(\vec{\pi}\right) \tag{22}$$

Given a $\vec{\rho}$, the probability of a specific corresponding path $\vec{\pi}$ is the ratio of the probability of $\vec{\pi}$ over the probability of $\vec{\rho}$.

$$P_{HMM}\left(\vec{\pi}|\vec{\rho}\right) = \frac{P_{HMM}\left(\vec{\rho}|\vec{\pi}\right) P_{HMM}\left(\vec{\pi}\right)}{P_{HMM}\left(\vec{\rho}\right)} = \frac{P_{HMM}\left(\vec{\pi}\right)}{\sum_{\vec{\pi}'\in\vec{\rho}} P_{HMM}\left(\vec{\pi}'\right)} \tag{23}$$

##### 3.2.2 Relation of the pHMM joint probability distributions in the different schemes

Here is the procedure for producing an unaligned sequence $\vec{x}$ with a pHMM under the four variable scheme. This method for sequence generation is much more convoluted than the method under the two variable scheme, but we are including it for the sake of clarity.

1. Generate an HPM-style path $\vec{\rho}$ from $P_{HMM}\left(\vec{\rho}\right)$, as defined in equation 22.
2. Generate $\vec{x}_i$ from $P_{HMM}\left(\vec{x}_i|\vec{\rho}\right)$ by emitting from the insert states in $\vec{\rho}$.
3. Generate $\vec{x}_m$ from $P_{HMM}\left(\vec{x}_m|\vec{\rho}\right)$. This is done in two steps:
   (a) Given $\vec{\rho}$ from step 1, pick a pHMM-style path $\vec{\pi}$ from $P_{HMM}\left(\vec{\pi}|\vec{\rho}\right)$ using equation 23.
   (b) Generate $\vec{x}_m$ by emitting from the match and delete states in $\vec{\pi}$ using $P_{HMM}\left(\vec{x}_m|\vec{\pi}\right)$.
4. Choose $\vec{x}$ from $\delta\left(\vec{x}, \vec{x}_m, \vec{x}_i, \vec{\rho}\right)$

- This is the exact same compatibility function we used with HPMs.

The joint probability of all 4 variables under a pHMM factors into individual terms.

$$P_{HMM}\left(\vec{x}, \vec{x}_m, \vec{x}_i, \vec{\rho}\right) = P_{HMM}\left(\vec{\rho}\right) P_{HMM}\left(\vec{x}_i|\vec{\rho}\right) P_{HMM}\left(\vec{x}_m|\vec{\rho}\right) \delta\left(\vec{x}, \vec{x}_m, \vec{x}_i, \vec{\rho}\right) \tag{24}$$

However, we need to write this probability in terms of $\vec{\pi}$ rather than $\vec{\rho}$ if we are to have any hope of using a pHMM as a proposal distribution in importance sampling.

First, we examine the insert emission process. Both $\vec{\pi}$ and $\vec{\rho}$ specify insert states in the same manner. Therefore, the probability of a particular $\vec{x}_i$ is the same if either $\vec{\rho}$ or $\vec{\pi}$ is given.

$$P_{HMM}\left(\vec{x}_i|\vec{\rho}\right) = P_{HMM}\left(\vec{x}_i|\vec{\pi}\right) \tag{25}$$

Next, we turn to the emission of match characters. Step 3 suggests the conditional probability of a match sequence given $\vec{\rho}$ can be factored into two terms.

$$P_{HMM}\left(\vec{x}_m|\vec{\rho}\right) = P_{HMM}\left(\vec{\pi}|\vec{\rho}\right) P_{HMM}\left(\vec{x}_m|\vec{\pi}\right) \tag{26}$$

We can arrive at this result in a different way by introducing $\vec{\pi}$ as a nuisance variable.

$$P_{HMM}\left(\vec{x}_m|\vec{\rho}\right) = \sum_{\vec{\pi}} P_{HMM}\left(\vec{x}_m, \vec{\pi}|\vec{\rho}\right) = \sum_{\vec{\pi}} P_{HMM}\left(\vec{x}_m|\vec{\pi}, \vec{\rho}\right) P_{HMM}\left(\vec{\pi}|\vec{\rho}\right) \tag{27}$$

Only pHMM paths that correspond to the given $\vec{\rho}$ will yield non-zero $P_{HMM}\left(\vec{\pi}|\vec{\rho}\right)$, meaning we only sum over these pHMM paths.

$$P_{HMM}\left(\vec{x}_m|\vec{\rho}\right) = \sum_{\vec{\pi}\in\vec{\rho}} P_{HMM}\left(\vec{x}_m|\vec{\pi}, \vec{\rho}\right) P_{HMM}\left(\vec{\pi}|\vec{\rho}\right) \tag{28}$$

Of the $2^M$ $\vec{\pi}$'s that correspond to our given $\vec{\rho}$, only one will have delete states in the match columns in which $x_m$ has deletion characters, so our sum collapses to only one term.

$$P_{HMM}\left(\vec{x}_m|\vec{\rho}\right) = P_{HMM}\left(\vec{x}_m|\vec{\pi},\vec{\rho}\right) P_{HMM}\left(\vec{\pi}|\vec{\rho}\right) \tag{29}$$

This result looks like what we have in equation 26, except $\vec{x}_m$ is conditioned on $\vec{\rho}$. We argue that we can eliminate this condition, as we only need $\vec{\pi}$ to generate $\vec{x}_m$.

$$P_{HMM}\left(\vec{x}_m|\vec{\rho}\right) = P_{HMM}\left(\vec{x}_m|\vec{\pi}\right) P_{HMM}\left(\vec{\pi}|\vec{\rho}\right) \tag{30}$$

Aside: This is in analogy to a Markov chain. If we transition from state $A$ to state $B$, and then state $B$ to state $C$, the probability of being in state $C$ is given by:

$$P\left(C\right) = P\left(C|B\right) P\left(B|A\right) P\left(A\right) \tag{31}$$

That is, $C$ is not *directly* conditioned on $A$. (end aside)

Now, we can write the joint probability of the four variables under the pHMM in terms of $\vec{\pi}$.

$$P_{HMM}\left(\vec{x},\vec{x}_m,\vec{x}_i,\vec{\rho}\right) = P_{HMM}\left(\vec{\rho}\right) P_{HMM}\left(\vec{x}_i|\vec{\pi}\right) P_{HMM}\left(\vec{x}_m|\vec{\pi}\right) P_{HMM}\left(\vec{\pi}|\vec{\rho}\right) \delta\left(\vec{x},\vec{x}_m,\vec{x}_i,\vec{\rho}\right) \tag{32}$$

Using equation 23, we can replace $P_{HMM}\left(\vec{\pi}|\vec{\rho}\right)$.

$$P_{HMM}\left(\vec{x},\vec{x}_m,\vec{x}_i,\vec{\rho}\right) = P_{HMM}\left(\vec{\rho}\right) P_{HMM}\left(\vec{x}_i|\vec{\pi}\right) P_{HMM}\left(\vec{x}_m|\vec{\pi}\right) \frac{P_{HMM}\left(\vec{\pi}\right)}{P_{HMM}\left(\vec{\rho}\right)} \delta\left(\vec{x},\vec{x}_m,\vec{x}_i,\vec{\rho}\right) \tag{33}$$

The $P_{HMM}(\vec{\rho})$ terms cancel.

$$P_{HMM}(\vec{x}, \vec{x}_m, \vec{x}_i, \vec{\rho}) = P_{HMM}(\vec{x}_i|\vec{\pi}) P_{HMM}(\vec{x}_m|\vec{\pi}) P_{HMM}(\vec{\pi}) \delta(\vec{x}, \vec{x}_m, \vec{x}_i, \vec{\rho}) \tag{34}$$

Note that this looks like the joint probability of $\vec{x}$ and $\vec{\pi}$ under the two variable scheme, multiplied by the delta function.

$$P_{HMM}(\vec{x}, \vec{\pi}) = P_{HMM}(\vec{x}|\vec{\pi}) P_{HMM}(\vec{\pi}) \tag{35}$$

By breaking $\vec{x}$ into two noncontiguous subsequences $\vec{x}_m$ and $\vec{x}_i$, we have:

$$P_{HMM}(\vec{x}, \vec{\pi}) = P_{HMM}(\vec{x}_i|\vec{\pi}) P_{HMM}(\vec{x}_m|\vec{\pi}) P_{HMM}(\vec{\pi}) \tag{36}$$

We can eliminate the $\delta$-function in equation 34 by noting that if we know $\vec{x}$ and $\vec{\pi}$, we can uniquely determine $\vec{x}_m$, $\vec{x}_i$, and $\vec{\rho}$; that is to say, $\delta(\vec{x}, \vec{x}_m, \vec{x}_i, \vec{\rho}) = 1$.

So we have proven $P_{HMM}(\vec{x}, \vec{\pi}) = P_{HMM}(\vec{x}, \vec{x}_m, \vec{x}_i, \vec{\rho})$.

##### 3.2.3 Relation of the pHMM posterior probability distributions across the different schemes

Using Bayes' theorem, the probability of $\vec{x}_m$, $\vec{x}_i$, and $\vec{\rho}$ given $\vec{x}$ is the ratio of the joint probability of all four variables and the marginal probability of $\vec{x}$.

$$P_{HMM}(\vec{x}_m, \vec{x}_i, \vec{\rho}|\vec{x}) = \frac{P_{HMM}(\vec{x}, \vec{x}_m, \vec{x}_i, \vec{\rho})}{P_{HMM}(\vec{x})} \tag{37}$$

The numerator we have shown to be $P_{HMM}(\vec{x}, \vec{\pi})$, while the denominator is calculated via the forward algorithm [3].

$$P_{HMM}(\vec{x}_m, \vec{x}_i, \vec{\rho}|\vec{x}) = \frac{P_{HMM}(\vec{x}, \vec{\pi})}{P_{HMM}(\vec{x})} \tag{38}$$

Therefore, $P_{HMM}(\vec{x}, \vec{x}_m, \vec{x}_i, \vec{\rho}|\vec{x}) = P_{HMM}(\vec{\pi}|\vec{x})$. So by sampling pHMM paths from $P_{HMM}(\vec{\pi}|\vec{x})$, we are simultaneously sampling $\vec{x}_m$, $\vec{x}_i$, and $\vec{\rho}$.

#### 3.3 Redefining the importance estimator under the four variable scheme

We can estimate the marginal probability of $\vec{x}$ under an HPM by using importance sampling. Our importance estimator is defined as:

$$\Phi \equiv \frac{1}{R} \sum_{r=1}^{R} \delta\left(\vec{x}, \vec{x}_m^{(r)}, \vec{x}_i^{(r)}, \vec{\rho}^{(r)}\right) \frac{P\left(\vec{x}_m^{(r)}, \vec{x}_i^{(r)}, \vec{\rho}^{(r)}\right)}{P_{HMM}\left(\vec{x}_m^{(r)}, \vec{x}_i^{(r)}, \vec{\rho}^{(r)}|\vec{x}\right)} \tag{39}$$

In this case, our importance weight is the rightmost term in the sum.

$$w\left(\vec{x}_m^{(r)}, \vec{x}_i^{(r)}, \vec{\rho}^{(r)}\right) = \frac{P\left(\vec{x}_m^{(r)}, \vec{x}_i^{(r)}, \vec{\rho}^{(r)}\right)}{P_{HMM}\left(\vec{x}_m^{(r)}, \vec{x}_i^{(r)}, \vec{\rho}^{(r)}|\vec{x}\right)} \tag{40}$$

In order for $\Phi$ to be an accurate, unbiased estimator of $P(\vec{x})$, the expectation value of $\Phi$ under the proposal distribution must equal $P(\vec{x})$. We will prove this is so.

$$\langle\Phi\rangle = \langle\frac{1}{R} \sum_{r=1}^{R} \delta\left(\vec{x}, \vec{x}_m^{(r)}, \vec{x}_i^{(r)}, \vec{\rho}^{(r)}\right) w\left(\vec{x}_m^{(r)}, \vec{x}_i^{(r)}, \vec{\rho}^{(r)}\right)\rangle \tag{41}$$

As we sample each combination of $\vec{x}_m$, $\vec{x}_i$, and $\vec{\rho}$ independently, we can rewrite the expectation of the sum as the sum of $R$ independent expectation values.

$$\langle\Phi\rangle = \langle\delta\left(\vec{x}, \vec{x}_m, \vec{x}_i, \vec{\rho}\right) w\left(\vec{x}_m, \vec{x}_i, \vec{\rho}\right)\rangle \tag{42}$$

Now we can write the expectation as a weighted average over all $\vec{x}_m$, $\vec{x}_i$, and $\vec{\rho}$ with non-zero probability under $P_{HMM}\left(\vec{x}_m^{(r)}, \vec{x}_i^{(r)}, \vec{\rho}^{(r)}|\vec{x}\right)$.

$$\langle\Phi\rangle = \sum_{\vec{x}_m}\sum_{\vec{x}_i}\sum_{\vec{\rho}}\delta\left(\vec{x}, \vec{x}_m, \vec{x}_i, \vec{\rho}\right) w\left(\vec{x}_m, \vec{x}_i, \vec{\rho}\right) P_{HMM}\left(\vec{x}_m, \vec{x}_i, \vec{\rho}|\vec{x}\right) \tag{43}$$

The factor of $P_{HMM}\left(\vec{x}, \vec{x}_m, \vec{x}_i, \vec{\rho}|\vec{x}\right)$ in the denominator of $w$ cancels with the rightmost term in the sum, so we are left with a sum over two terms.

$$\langle\Phi\rangle = \sum_{\vec{x}_m}\sum_{\vec{x}_i}\sum_{\vec{\rho}}\delta\left(\vec{x}, \vec{x}_m, \vec{x}_i, \vec{\rho}\right) P\left(\vec{x}_m, \vec{x}_i, \vec{\rho}\right) \tag{44}$$

This is the marginal probability defined in equation 20. We have proven $\langle\Phi\rangle = P\left(\vec{x}\right)$

### 4 Testing the accuracy of importance sampling scores

While the importance sampling estimate of $P\left(\vec{x}\right)$ converges to the exact answer in theory, two important questions remain: does importance sampling accurately estimate $P\left(\vec{x}\right)$ in the case of real biological sequences; and if so, how many sampled paths are sufficient? Ideally, we would answer these questions by comparing importance sampling estimates of $P\left(\vec{x}\right)$ with exact values calculated by enumerating all possible paths under an HPM. However, this is not tractable with HPMs representing actual sequence families.

As a control, we instead score sequences with a covariance model (CM), a type of stochastic context free grammar specific to RNA homology search [2], using the software package Infernal [6]. A CM can be viewed as a special case of Potts model in which all $e_{kl}$ terms are zero except for a set of disjoint nested basepairing interactions. The marginal probability of a sequence under a CM, $P_{CM}\left(\vec{x}\right)$, can be calculated *exactly* using the dynamic programming inside algorithm [3]. Importance sampling a CM using a pHMM as a proposal distribution is also feasible.

$$P_{CM}\left(\vec{x}\right) \approx \frac{1}{R}\sum_{r=1}^{R}\frac{P_{CM}\left(\vec{x},\vec{\pi}^{(r)}\right)}{P_{HMM}\left(\vec{\pi}^{(r)}|\vec{x}\right)} \tag{45}$$

Therefore, we can compare our importance sampling estimates to exact, efficiently calculated probability values under a model similar to a Potts model.

We train a CM on our tRNA training MSA (modified from data in [7]) using Infernal (see 4.1 Control experiment methods). For a proposal distribution, we once again use a pHMM trained by HMMER 4 (in progress). We then score the train, test, and decoy sequences from our tRNA benchmark using the inside algorithm and importance sampling with 5 million sampled paths per sequence. Rather than marginal probability, Infernal reports a log odds score, comparing $P_{CM}\left(\vec{x}\right)$ to the probability that $\vec{x}$ is generated by a null model, $P_N\left(\vec{x}\right)$.

$$S_{CM}\left(\vec{x}\right) = \log\frac{P_{CM}\left(\vec{x}\right)}{P_N\left(\vec{x}\right)} \tag{46}$$

Figure 2 shows that for most tRNA homologs, importance sampling accurately estimates

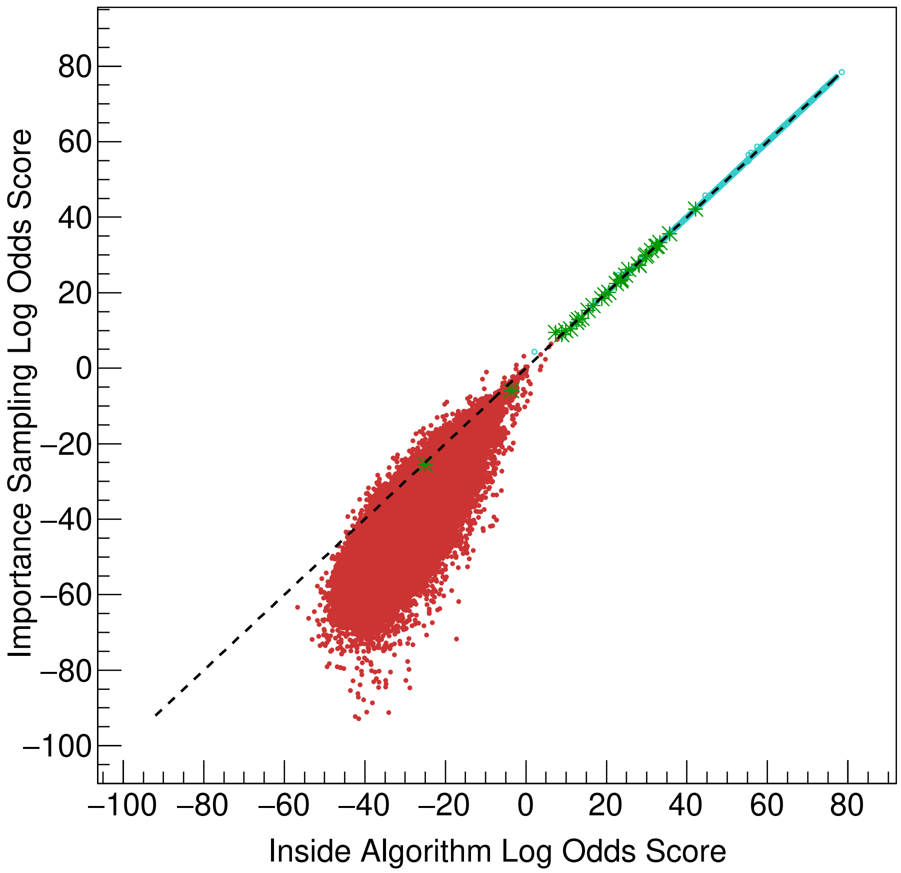


Figure 2: Scores for tRNA benchmark sequences under the training CM calculated with the inside algorithm (X axis) and estimated using importance sampling (Y axis). Each point represents an individual sequence from the tRNA remote homology benchmark, with training sequences in blue, homologous test sequences in green, and non-homologous decoy sequences in red. The dashed line represents $y = x$.

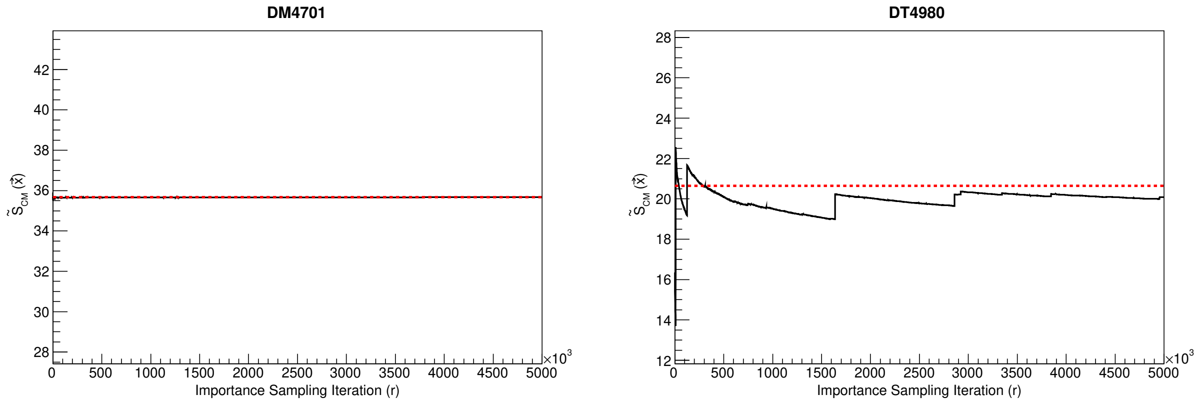


Figure 3: Importance sampling estimate of sequence log odds score for an individual sequence (Y axis) versus number of sampled paths (X axis). The red dashed line represents the exact log odds of the sequence under the CM, as calculated by the inside algorithm. Left: tRNA test sequence DM4701, a mitochondrial methionine tRNA from *Mytilus edulis*. Right: tRNA test sequence DT4980, a mitochondrial threonine tRNA from *Pisaster ochraceus* [7].

$S_{cm}(\vec{x})$. For decoys, the agreement is not as sharp, and importance sampling generally underestimates the exact value of $S_{cm}(\vec{x})$. We conclude importance sampling is an accurate method for estimating the marginal probability of a homologous sequence under a CM, does not inflate the score of non-homologous decoys on average, and is therefore likely accurate in estimating $P(\vec{x})$ under an HPM.

In general, the importance sampling estimate of $S_{CM}(\vec{x})$ quickly converges to the exact value. For instance, figure 3 shows the log odds score for tRNA test sequence DM4701 converges well before 1 million samples. A few sequences' scores do not converge, instead showing sudden "jumps" in score estimation, like tRNA sequence DT4980. Such jumps in importance sampling score are a known phenomenon [4]. They appear when the denominator in equation 45 is very small compared to the numerator, meaning the posterior probability of the path under the pHMM is much lower than the joint probability of the path and the sequence under the CM. Jumps in score are rare and appear to be confined to a minority of problematic homologous sequences. One million sampled paths per sequence appears to be sufficient for most sequences' importance sampling scores to converge, and future work will focus on increasing convergence of importance sampling scores and eliminating score jumps.

#### 4.1 Control experiment methods

CMs are built from the tRNA remote homology benchmark training MSA using Infernal 1.1.3 program `cmbuild`. As Infernal generally aligns and scores sequences locally, transitions to and from flanking insert states (analogous to the N and C states in figure 1) are not trained using observed frequencies in the MSA by default. As importance sampling works glocally, we build the CM such that the flanking insert transitions are learned using the `--iflank` option. In addition, we specify the match columns in the MSA using `--hand`.

```
cmbuild --hand --iflank <infernal_model.cm> <training_msa.sto>
```

The pHMM is built with HMMER 4, again using the `--hand` option to specify the consensus columns in the input MSA.

```
hmmer build --hand <training_msa.sto> <hmmer_model.hmm>
```

To exactly score the sequences with the training CM using the inside algorithm, we use Infernal 1.1.3 program `cmalign`. We use the `-g` and `--notrunc` options for glocal scoring. We additionally use the `--nonbanded` option to run the inside algorithm without any approximation schemes.

```
cmalign -g --notrunc --nonbanded --sfile <inside_scores.tbl>
↪  <infernal_model.cm> <seqfile.fa>
```

To score the sequences with the training CM using importance sampling, we use our own program, `cmscoreISh4`.

```
cmscoreISh4 <infernal_model.cm> <hmmer_model.hmm> <seqfile.fa>
↪  <IS_scorefile.csv>
```

Code used in this experiment is included with **S1 Code**.

### References


[1] Haussler D, Krogh A, Mian IS, Sjolander K. Protein Modeling Using Hidden Markov Models: Analysis of Globins. In: Proceedings of the Twenty-Sixth Hawaii International Conference on System Sciences; 1993. p. 792–802.

[2] Eddy SR, Durbin R. RNA Sequence Analysis Using Covariance Models. Nucl Acids Res. 1994;22:2079–2088.

[3] Durbin R, Eddy SR, Krogh A, Mitchison GJ. Biological Sequence Analysis: Probabilistic Models of Proteins and Nucleic Acids. Cambridge UK: Cambridge University Press; 1998.

[4] Mackay DJC. Information Theory, Inference, and Learning Algorithms. Cambridge UK: Cambridge University Press; 2003.

[5] Anderson EC. Monte Carlo Methods and Importance Sampling: Lecture notes for Stat 578C Statistical Genetics; 1999 [cited 11 June 20]. Available from: `http://ib.berkeley.edu/labs/slatkin/eriq/classes/guest_lect/mc_lecture_notes.pdf`.

[6] Nawrocki EP, Eddy SR. Infernal 1.1: 100-fold Faster RNA Homology Searches. Bioinformatics. 2013;29:2933–2935.

[7] Sprinzl M, Horn C, Brown M, Ioudovitch A, Steinberg S. Compilation of tRNA Sequences and Sequences of tRNA Genes. Nucl Acids Res. 1998;26:148–153.
