## Supplemental Code S1 for "Remote homology search with hidden Potts models": 00README

I Contents:

- ./infernal-1.1.3/ : Infernal version 1.1.3 (must be cloned from GitHub)
- ./hmmer4/ : HMMER version 4 (in development) (must be cloned from GitHub)
- ./HPMHomology/ : code to build HPMs, align and score sequences using HPMs
- ./CMIS/ : code to score and align sequences with Infernal CMs using importance sampling
- ./data/ : benchmark datasets for tRNA, Twister ribozyme, and SAM riboswitch benchmarks
- ./Tutorial.txt: instructions for how to recreate the Twister ribozyme benchmark results.
- ./compile.sh: bashh script for compiling

II Compiling instructions

A. Programs required to compile:
 - git
 - make
 - gcc
 - autoconf

B. Quick install

To clone Infernal and HMMER4, run
> ./clone.sh

Next, to compile, run:
> ./compile.sh

Or follow the step-by-step instructions below.

C. Step-by-step install

1. Instructions for cloning Infernal from Git
> git clone https://github.com/EddyRivasLab/infernal.git
>
> mv infernal infernal-1.1.3
>
> cd infernal-1.1.3
>
> git checkout a358a49
>
> git clone https://github.com/EddyRivasLab/hmmer.git
>
> git clone https://github.com/EddyRivasLab/easel.git
>
> cd hmmer
>
> git checkout bcd05b0
>
> cd ../easel
>
> git checkout fa17f53
>
> cd ../..


2. Instructions for cloning HMMER4 from git
> git clone https://github.com/EddyRivasLab/hmmer.git
>
> mv hmmer hmmer4
>
> cd hmmer4
>
> git checkout 7994cc7
>
> git clone https://github.com/EddyRivasLab/easel.git
>
> cd easel
>
> git checkout 66745ea
>
> cd ../..

3. Instructions to compile Infernal
> cd infernal-1.1.3/
>
> autoconf
>
> ./configure
>
> make
>
> cd ..

4. Instructions to compile HMMER4
> cd hmmer4/easel
>
> make
>
> cd ..
>
> ln -s ${PWD}/easel lib/easel
>
> autoconf
>
> ./configure
>
> make
>
> cd nwo/
>
> make
>
> cd ../..

5. Instructions to compile HPMHomology code
> cd HPMHomology/src/
>
> make
>
> g++ -O3 -std=c++0x -o gremlin_h2_henikoff gremlin_h2_henikoff.cpp
>
> cd ../..

6. Instructions to compile CMIS code
> cd CMIS/src/
>
> make
>
> cd ../..
