## Supplemental Code S1 for "Remote homology search with hidden Potts models": clone.sh

```
#!/bin/bash
# bash script to clone infernal and hmmer4 from github

# Grab infernal from git
echo "Cloning infernal and associated code from git"
git clone https://github.com/EddyRivasLab/infernal.git

mv infernal infernal-1.1.3

cd infernal-1.1.3

# checkout tag used for paper
git checkout a358a49

git clone https://github.com/EddyRivasLab/hmmer.git

git clone https://github.com/EddyRivasLab/easel.git

cd hmmer

git checkout bcd05b0

cd ../easel

git checkout fa17f53

cd ../..

# Grab hmmer4 from github
echo "Cloning hmmer4 and associated code from git"

git clone https://github.com/EddyRivasLab/hmmer.git

mv hmmer hmmer4

cd hmmer4

git checkout 7994cc7

git clone https://github.com/EddyRivasLab/easel.git

cd easel

git checkout 66745ea

cd ../..
```
