## Supplemental Code S1 for "Remote homology search with hidden Potts models": cmemit_posterior.c

```
#include <stdio.h>
#include <stdlib.h>
#include <string.h>

#include "easel.h"
#include "esl_alphabet.h"
#include "esl_getopts.h"
#include "esl_msa.h"
#include "esl_random.h"
#include "esl_sq.h"
#include "esl_sqio.h"
#include "esl_vectorops.h"

#include "h4_config.h"
#include "h4_hmmfile.h"
#include "h4_mode.h"
#include "h4_path.h"
#include "h4_profile.h"
#include "h4_refmx.h"
#include "logsum.h"
#include "reference_dp.h"

#include "infernal.h"
#include "config.h"

#include "h4_path_cmis.h"

/* declaration of internal functions */
int generate_msa(CM_t *cm, ESL_SQ **sq, ESL_RANDOMNESS *rng, int nseq, int R, float *cmsc_ld, float *insc_ld, ESL_MSA **ret_msa, char *errbuf);
int msa_score_h4(H4_PROFILE *hmm, ESL_MSA *msa, ESL_SQ **sq, int nseq, int R, float *hmmsc_ld, float *fwdsc_ld);
int write_comp_scorefile(ESL_SQ **sq, float *insc_ld, float *cmsc_ld, float *hmmsc_ld, float *fwdsc_ld, int nseq, int R, char *scorefile);

static ESL_OPTIONS options[] = {
        /* name       type        default    env range togs  reqs  incomp            help                                             docgroup */
        { "-h",       eslARG_NONE,    FALSE, NULL, NULL, NULL, NULL, NULL,            "help; show brief info on version and usage",   1 },

        /* Options forcing which alphabet we're working in (normally autodetected) */
        { "--amino",  eslARG_NONE,    FALSE, NULL, NULL, NULL, NULL, "--dna,--rna",   "We are dealing with protein sequences",        2 },
        { "--dna",    eslARG_NONE,    FALSE, NULL, NULL, NULL, NULL, "--amino,--rna", "We are dealing with dna sequences",            2 },
        { "--rna",    eslARG_NONE,    FALSE, NULL, NULL, NULL, NULL, "--amino,--dna", "We are dealing with rna sequences",            2 },

        /* options for controlling sampling */
        { "-R",       eslARG_INT,     "10",  NULL, NULL, NULL, NULL, NULL,            "Number of CM paths sampled per sequence",      3 },
        { "-s",       eslARG_INT,      "0",   NULL, NULL, NULL, NULL, NULL,           "Set random number seed to <n>",                3 },

        { 0,0,0,0,0,0,0,0,0,0 },
};

static char banner[] = "Emit alignments from P_cm (path | sequence)";
static char usage[]  = "[-options] <cmfile> <hmmfile> <seqfile> <msa_outfile> <score_outfile>";

static void
cmdline_failure(char *argv0, char *format, ...)
{
        va_list argp;
        printf("\nERROR: ");
        va_start(argp, format);
        vfprintf(stderr, format, argp);
        va_end(argp);
        esl_usage(stdout, argv0, usage);
        printf("\nTo see more help on available options, do %s -h\n\n", argv0);
        exit(1);
}

int main (int argc, char *argv[]) {
   ESL_GETOPTS      *go               = NULL;                 /* application configuration                    */
   ESL_ALPHABET     *abc              = NULL;                 /* biological alphabet                          */
   ESL_RANDOMNESS   *rng              = NULL;                 /* random number generator                      */
   char             *cmfile           = NULL;                 /* input cm filepath                            */
   char             *hmmfile          = NULL;                 /* input hmmm filepath                          */
   char             *seqfile          = NULL;                 /* input seq filepath                           */
   char             *msafile          = NULL;                 /* output msa filepath                          */
   char             *scorefile        = NULL;                 /* output score csv filepath                    */
   CM_FILE          *cmfp             = NULL;                 /* open input CM file stream                    */
   H4_HMMFILE       *hmmfp            = NULL;                 /* open input hmm file stream                   */
   ESL_SQFILE       *sqfp             = NULL;                 /* open seq file stream                         */
   CM_t             *cm               = NULL;                 /* cm                                           */
   H4_PROFILE       *hmm              = NULL;                 /* hmm                                          */
   ESL_SQ          **sq               = NULL;                 /* array of sequences                           */
   int               format           = eslSQFILE_UNKNOWN;    /* seq file format                              */
   FILE             *afp              = NULL;                 /* output alignment file (-A)                   */
   ESL_MSA          *msa              = NULL;                 /* alignment output object                      */
   float            *cmsc_ld          = NULL;                 /* array of S_cm (x,pi)                         */
   float            *hmmsc_ld         = NULL;                 /* array of S_hmm (x,pi)                        */
   float            *insc_ld          = NULL;                 /* array insde scores for each input seq        */
   float            *fwdsc_ld         = NULL;                 /* array of (raw) fwd scores for each input seq */
   int               outfmt           = eslMSAFILE_STOCKHOLM; /* alignment output format                      */
   int               n                = 0;                    /* sequence index                               */
   int               nseq             = 0;                    /* total number of seqs in seq file             */
   int               R;                                       /* number of paths sampled per sequence         */
   int               status;                                  /* easel return code                            */
   char              errbuf[eslERRBUFSIZE];


   /* parse command line */
   go = esl_getopts_Create(options);
   if (esl_opt_ProcessCmdline(go, argc, argv) != eslOK) cmdline_failure(argv[0], "Failed to parse command line: %s\n", go->errbuf);
   if (esl_opt_VerifyConfig(go)               != eslOK) cmdline_failure(argv[0], "Error in app configuration:   %s\n", go->errbuf);

   if (esl_opt_GetBoolean(go, "-h") ) {
      esl_banner(stdout, argv[0], banner);
      esl_usage (stdout, argv[0], usage);
      puts("\n where options are:");
      esl_opt_DisplayHelp(stdout, go, 1, 2, 80);
      puts("\n Alphabet options:");
      esl_opt_DisplayHelp(stdout, go, 2, 2, 80);
      puts("\n Sampling options:");
      esl_opt_DisplayHelp(stdout, go, 3, 2, 80);

      exit(0);
   }

   /* read arguments */
   if (esl_opt_ArgNumber(go) != 5)  cmdline_failure(argv[0], "Incorrect number of command line arguments.\n", go->errbuf);

   cmfile    =  esl_opt_GetArg(go, 1);
   hmmfile   =  esl_opt_GetArg(go, 2);
   seqfile   =  esl_opt_GetArg(go, 3);
   msafile   =  esl_opt_GetArg(go, 4);
   scorefile =  esl_opt_GetArg(go, 5);


   /* if user has defined an alphabet we define it here */
   if        (esl_opt_GetBoolean(go, "--amino"))       abc = esl_alphabet_Create(eslAMINO);
   else if   (esl_opt_GetBoolean(go, "--rna"))         abc = esl_alphabet_Create(eslRNA);
   else if   (esl_opt_GetBoolean(go, "--dna"))         abc = esl_alphabet_Create(eslDNA);

   /* get number of paths */
   R = esl_opt_GetInteger(go, "-R");

   /* create random number generator */
   rng = esl_randomness_Create(esl_opt_GetInteger(go, "-s"));

   /* open the .cm file */
   status = cm_file_Open(cmfile, NULL, FALSE, &cmfp, errbuf);
   if      (status == eslENOTFOUND) cm_Fail("File existence/permissions problem in trying to open CM file %s.\n%s\n", cmfile, errbuf);
   else if (status == eslEFORMAT)   cm_Fail("File format problem in trying to open CM file %s.\n%s\n",                cmfile, errbuf);
   else if (status != eslOK)        cm_Fail("Unexpected error %d in opening CM file %s.\n%s\n",               status, cmfile, errbuf);

   /* read first cm in file */
   status = cm_file_Read(cmfp, TRUE, &abc, &cm);
   if      (status == eslEFORMAT)   cm_Fail("Bad file format in CM file %s:\n%s\n",          cmfp->fname, cmfp->errbuf);
   else if (status == eslEINCOMPAT) cm_Fail("CM in %s is not in the expected %s alphabet\n", cmfp->fname, esl_abc_DecodeType(abc->type));
   else if (status == eslEOF)       cm_Fail("Empty CM file %s? No CM data found.\n",         cmfp->fname);
   else if (status != eslOK)        cm_Fail("Unexpected error in reading CMs from %s\n",     cmfp->fname);

   cm_file_Close(cmfp);

   /* configure CM */
   cm->config_opts |= CM_CONFIG_SCANMX;
   cm_Configure(cm, errbuf, -1);

   /* set search options */
   cm->search_opts |= CM_SEARCH_INSIDE;
   cm->search_opts |= CM_SEARCH_NONBANDED;

   /* initlalize logsum lookup table magic */
   init_ilogsum();
   FLogsumInit();

   /* print all configuration flags */
   DumpCMFlags(stdout, cm);

   /* open the h4 hmm file */
   if ( h4_hmmfile_Open(hmmfile, NULL, &hmmfp) != eslOK) esl_fatal("couldn't open profile file %s", hmmfile);
   /* read first hmm from hmm file */
   if ( h4_hmmfile_Read(hmmfp, &abc, &hmm)     != eslOK) esl_fatal("failed to read profile from file %s", hmmfile);
   h4_hmmfile_Close(hmmfp);

   /* initiate h4's logsum table lookup magic */
   h4_logsum_Init();

   /* check that cm, hmm have same # of match states */
   if (hmm->M != cm->clen) esl_fatal("The hmm and cm have different numbers of consensus columns!\n hmm->M: %d \n cm->M: %d\n", hmm->M, cm->clen);

   /* open the sequence file */
   status = esl_sqfile_OpenDigital(abc, seqfile, format, NULL, &sqfp);
   if      (status == eslENOTFOUND) esl_fatal("No such file.");
   else if (status == eslEFORMAT)   esl_fatal("Format unrecognized.");
   else if (status == eslEINVAL)    esl_fatal("Can't autodetect stdin or .gz.");
   else if (status != eslOK)        esl_fatal("Open failed, code %d.", status);

   /* read sequences into array */
   ESL_REALLOC(sq, sizeof(ESL_SQ *) * (n + 1));
   sq[n] = esl_sq_CreateDigital(abc);
   while ((status = esl_sqio_Read(sqfp, sq[n+nseq])) == eslOK) {
      nseq++;
      ESL_REALLOC(sq, sizeof(ESL_SQ *) * (n+nseq+1));
      sq[n+nseq] = esl_sq_CreateDigital(abc);
   }

   /* error handling and cleanup */
   if      (status == eslEFORMAT) esl_fatal("Parse failed (sequence file %s):\n%s\n",
                                            sqfp->filename, esl_sqfile_GetErrorBuf(sqfp));
   else if (status != eslEOF)     esl_fatal("Unexpected error %d reading sequence file %s",
                                            status, sqfp->filename);

   /* allocate space for score arrays */
   ESL_ALLOC(cmsc_ld,  sizeof(float *) * nseq * R);
   ESL_ALLOC(hmmsc_ld, sizeof(float *) * nseq * R);
   ESL_ALLOC(insc_ld,  sizeof(float *) * nseq);
   ESL_ALLOC(fwdsc_ld, sizeof(float *) * nseq);
   esl_vec_FSet(cmsc_ld,  nseq*R, 0.0);
   esl_vec_FSet(hmmsc_ld, nseq*R, 0.0);
   esl_vec_FSet(insc_ld,  nseq,   0.0);
   esl_vec_FSet(fwdsc_ld, nseq,   0.0);

   if ((status = generate_msa(cm, sq, rng, nseq, R, cmsc_ld, insc_ld, &msa, errbuf) != eslOK))  esl_fatal("issue running generate_msa(), returned code %d", status);

   if ((status = msa_score_h4(hmm, msa, sq, nseq, R, hmmsc_ld, fwdsc_ld) != eslOK))  esl_fatal("issue running msa_score_h4(), returned code %d", status);

   /* write MSA to output file */
   if ( (afp = fopen(msafile,"w"))== NULL)  esl_fatal("Failed to open alignment file %s for writing\n", msafile);
   esl_msafile_Write(afp, msa, outfmt);
   fclose(afp);

   /* write score comparison output csv */
   write_comp_scorefile(sq, insc_ld, cmsc_ld, hmmsc_ld, fwdsc_ld, nseq, R, scorefile);

   /* clean up and return */
   for (n = 0; n < nseq+1; n++)
   {
      esl_sq_Destroy(sq[n]);
   }

   esl_sqfile_Close(sqfp);
   esl_msa_Destroy(msa);
   free(sq);
   FreeCM(cm);
   free(cmsc_ld);
   free(hmmsc_ld);
   free(insc_ld);
   free(fwdsc_ld);
   h4_profile_Destroy(hmm);
   esl_randomness_Destroy(rng);
   esl_alphabet_Destroy(abc);
   esl_getopts_Destroy(go);
   return 0;

   ERROR:
      return status;
}

int generate_msa(CM_t *cm, ESL_SQ **sq, ESL_RANDOMNESS *rng, int nseq, int R, float *cmsc_ld, float *insc_ld, ESL_MSA **ret_msa, char *errbuf) {
   int           nrow = R * nseq;             /* total number of paths we will generate                   */
   int           n;                           /* sequence index [0,...,nseq-1]                            */
   int           r;                           /* samp;e index [0,...,R-1]                                 */
   int           idx;                         /* combined sequence-sample index [0,...,nrow-1]            */
   ESL_MSA      *msa = NULL;                  /* msa we're building                                       */
   Parsetree_t **pstr = NULL;                 /* array of parsetrees we are generating                    */
   ESL_SQ      **sq_dummy = NULL;             /* dummy sequence array for output MSA                      */
   CM_MX        *mx    = cm_mx_Create(cm->M); /* DP matrix for running inside algorithm                   */
   int           status;                      /* esl return code                                          */


   /* allocate space for parsetree array */
   ESL_ALLOC(pstr, nrow * sizeof(Parsetree_t * ));

   /* allocate space for sequence array */
   ESL_ALLOC(sq_dummy, sizeof(P7_TRACE *) * nrow);
   for (idx = 0; idx < nrow; idx++) sq_dummy[idx] = esl_sq_CreateDigital(cm->abc);

   for (n = 0; n < nseq; n++) {

      /* run inside algorithm */
      cm_InsideAlign(cm, errbuf, sq[n]->dsq, sq[n]->n, 512.0, mx, &insc_ld[n]);

      for (r = 0; r < R; r++) {
         idx = (n*R) + r;

         /* copy sequence into dummy array, make unique name */
         esl_sq_Copy(sq[n], sq_dummy[idx]);
         sprintf(sq_dummy[idx]->name, "%s_%d", sq[n]->name, r);

         /* sample parsetree */
         if((status = cm_StochasticParsetree(cm, errbuf, sq[n]->dsq, sq[n]->L, mx, rng, &pstr[idx], &cmsc_ld[idx])) != eslOK) return status;
         //ParsetreeDump(stdout, pstr[n*nseq + r], cm, sq[n]->dsq);
      }
   }

   /* generate MSA from sampled parsetrees */
   if ((status = Parsetrees2Alignment(cm, errbuf, cm->abc, sq_dummy, NULL, pstr, NULL, nrow, NULL, NULL, TRUE, FALSE, &msa))) return status;
   /* digitize msa (needed for h4 path creation */
   if ((status = esl_msa_Digitize(cm->abc, msa, errbuf))!= eslOK) return status;
   *ret_msa = msa;

   /* clean up and return */
   for (idx = 0; idx < nrow; idx++)
   {
      FreeParsetree(pstr[idx]);
      esl_sq_Destroy(sq_dummy[idx]);
   }

   free(pstr);
   cm_mx_Destroy(mx);
   free(sq_dummy);
   return eslOK;

   ERROR:
      return status;
}


int msa_score_h4(H4_PROFILE *hmm, ESL_MSA *msa, ESL_SQ **sq, int nseq, int R, float *hmmsc_ld, float *fwdsc_ld)
{
   int8_t       *matassign = NULL;                    /* flag for each msa column: match or not [0,...,msa->alen] */
   H4_PATH     **pi       = NULL;                     /* array of h4 paths                                        */
   H4_MODE      *mo     = h4_mode_Create();           /* h4 profile hmm mode                                      */
   H4_REFMX     *fwd     = h4_refmx_Create(100, 100); /* DP matrix for forward algorithm                          */
   int           optflags = 0;                        /* options for creating MSA (not yet used...)               */
   int           apos;                                /* msa column index                                         */
   int           idx;                                 /* msa row (sequence) index                                 */
   int           r;                                   /* counter over sampled paths per seq                       */
   int           n;                                   /* counter over seqs in input seq file                      */
   int           status;                              /* esl return code                                          */

   /* set HMM mode to uniglocal  */
   h4_mode_SetUniglocal(mo);

   //h4_profile_Dump(stdout, hmm);

   /* make matassign array */
   ESL_ALLOC(matassign, sizeof(int8_t) * (msa->alen + 1));

   /* we ignore column zero */
   matassign[0] = 0;

   /* read RF line to build matassign */
   for (apos=0; apos < msa->alen; apos++){
      if (msa->rf[apos] != '.') matassign[apos+1] = 1;
      else                      matassign[apos+1] = 0;
   }

   ESL_ALLOC(pi, sizeof(H4_PATH *) * msa->nseq);

   /* convert msa to h4_path array */
   h4_path_FromMSA(msa, matassign, optflags, pi);

   /* loop over paths and score them with hmm */
   for (n = 0; n < nseq; n++) {
      /* set mode length */
      h4_mode_SetLength(mo, sq[n]->n);

      /* run forward matrix for this sequence */
      h4_reference_Forward(sq[n]->dsq, sq[n]->n, hmm, mo, fwd, &fwdsc_ld[n]);

      for (r = 0; r < R; r++) {
         idx = (n*R) + r;
         h4_path_Score(pi[idx], sq[n]->dsq, hmm, mo, &hmmsc_ld[idx]);
         /* get path dump */
         //h4_path_Dump(stdout, pi[idx]);
         //h4_path_DumpAnnotated(stdout, pi[idx], hmm, mo, sq[n]->dsq);
      }
   }

   /* clean up and return */
   for (idx = 0; idx < msa->nseq; idx++)
   {
      h4_path_Destroy(pi[idx]);
   }
   h4_mode_Destroy(mo);
   h4_refmx_Destroy(fwd);
   free(pi);
   free(matassign);
   return eslOK;

   ERROR:
      return status;

}


int write_comp_scorefile(ESL_SQ **sq, float *insc_ld, float *cmsc_ld, float *hmmsc_ld, float *fwdsc_ld, int nseq, int R, char *scorefile) {
   int    n;
   int    r;
   int    idx;
   FILE  *sfp  = NULL;  /* output score csv file */


   if ((sfp = fopen(scorefile, "w")) == NULL) esl_fatal("Failed to open output score file %s for writing", scorefile);

   fprintf(sfp, "seq,path,cmsc,insc,lpcm(path|seq),hmmsc,fwdsc,lphmm(path|seq),is_sum_cont\n");
   for (n = 0; n < nseq; n++) {
      for (r = 0; r < R; r++){
         idx = (n*R) + r;
         fprintf(sfp,
                 "%s,%d,%.2f,%.2f,%.2f,%.2f,%.2f,%.2f,%.2f\n",
                 sq[n]->name,
                 r,
                 cmsc_ld[idx],
                 insc_ld[n],
                 cmsc_ld[idx] - insc_ld[n],
                 hmmsc_ld[idx],
                 fwdsc_ld[n],
                 hmmsc_ld[idx] - fwdsc_ld[n],
                 cmsc_ld[idx] - hmmsc_ld[idx]);
      }
   }

   fclose(sfp);

   return eslOK;
}
```
