## Supplemental Code S1 for "Remote homology search with hidden Potts models": cmis_scoreset.c

```
/* cmis_scoreset.c */

#include <stdio.h>

#include "easel.h"
#include "esl_vectorops.h"

#include "cmis_scoreset.h"

CMIS_SCORESET *
cmis_scoreset_Create(int nseq) {
   CMIS_SCORESET *cmis_ss = NULL;
   int status;


   ESL_ALLOC(cmis_ss, sizeof(CMIS_SCORESET));

   /* set number of sequences */
   cmis_ss->nseq = nseq;

   /* allocate memory for sequence name array */
   ESL_ALLOC(cmis_ss->sqname, sizeof(char *) * nseq);

   /* allocate memory for probabilities/scores */
   ESL_ALLOC(cmis_ss->R,           sizeof(int)     * nseq);
   ESL_ALLOC(cmis_ss->fsc,         sizeof(float)   * nseq);
   ESL_ALLOC(cmis_ss->ntsc,        sizeof(float)   * nseq);
   ESL_ALLOC(cmis_ss->insc,        sizeof(float)   * nseq);
   ESL_ALLOC(cmis_ss->scansc,      sizeof(float)   * nseq);
   ESL_ALLOC(cmis_ss->seq_from,    sizeof(int)     * nseq);
   ESL_ALLOC(cmis_ss->seq_to,      sizeof(int)     * nseq);
   ESL_ALLOC(cmis_ss->cmis_ld,     sizeof(float)   * nseq);


   /* initialize values */
   esl_vec_ISet(cmis_ss->R,         nseq,   0);
   esl_vec_FSet(cmis_ss->fsc,       nseq, 0.0);
   esl_vec_FSet(cmis_ss->ntsc,      nseq, 0.0);
   esl_vec_FSet(cmis_ss->insc,      nseq, 0.0);
   esl_vec_FSet(cmis_ss->scansc,    nseq, 0.0);
   esl_vec_ISet(cmis_ss->seq_from,  nseq,   0);
   esl_vec_ISet(cmis_ss->seq_to,    nseq,   0);
   esl_vec_FSet(cmis_ss->cmis_ld,   nseq, 0.0);

   return cmis_ss;

   ERROR:
      return NULL;
}


void
cmis_scoreset_Destroy( CMIS_SCORESET *cmis_ss) {

   /* ignore NULL pointer */
   if (cmis_ss == NULL) return;

   /* free arrays */
   if (cmis_ss->sqname)    free(cmis_ss->sqname);
   if (cmis_ss->R)         free(cmis_ss->R);
   if (cmis_ss->fsc)       free(cmis_ss->fsc);
   if (cmis_ss->ntsc)      free(cmis_ss->ntsc);
   if (cmis_ss->insc)      free(cmis_ss->insc);
   if (cmis_ss->scansc)    free(cmis_ss->scansc);
   if (cmis_ss->seq_from)  free(cmis_ss->seq_from);
   if (cmis_ss->seq_to)    free(cmis_ss->seq_to);
   if (cmis_ss->cmis_ld)   free(cmis_ss->cmis_ld);

   /* free the scoreset object */
   free(cmis_ss);

   return;
}

int
cmis_scoreset_Write(FILE *fp, CMIS_SCORESET *cmis_ss) {
   int n;

   /* write csv header line */
   fprintf(fp, "id,N_sample,raw_fwd,null_tran,InsideAlign_logodds,InsideScan_logodds,seq_from,seq_to,cmis_logodds\n");

   /* loop over sequences, print id and scores */
   for (n = 0; n < cmis_ss->nseq; n++) {
      fprintf(fp, "%s,%d,%.4f,%.4f,%.4f,%.4f,%d,%d,%.4f\n",
                  cmis_ss->sqname[n],
                  cmis_ss->R[n],
                  cmis_ss->fsc[n],
                  cmis_ss->ntsc[n],
                  cmis_ss->insc[n],
                  cmis_ss->scansc[n],
                  cmis_ss->seq_from[n],
                  cmis_ss->seq_to[n],
                  cmis_ss->cmis_ld[n]);
    }


   return eslOK;
}
```
