## Supplemental Code S1 for "Remote homology search with hidden Potts models": cmis_scoreset.h

```
#ifndef CMIS_SCORESET_INCLUDED
#define CMIS_SCORESET_INCLUDED

#include <stdio.h>

#include "easel.h"
#include "esl_vectorops.h"

typedef struct cmisscoreset_s{
   char   **sqname;    /* sequence names, [0...nseq-1][], \0-terminated                 */
   int      nseq;      /* number of sequences                                           */
   int     *R;         /* number of samples/sequence, [0...nseq-1]                      */
   float   *fsc;       /* forward partial log-odds scores, in bits [0...nseq-1]         */
   float   *ntsc;      /* hmmer null model transition scores, in bits [0...nseq-1]      */
   float   *insc;      /* inside align alg log odds scores, in bits, [0...nseq-1]       */
   float   *scansc;    /* inside scan alg log odds scores, in bits, [0...nseq-1]        */
   int     *seq_from;  /* start point of inside scan hit, in seq coords                 */
   int     *seq_to;    /* end point of inside scan hit, in seq coords                   */
   float   *cmis_ld;   /* CM importance sampling log-odds scores, in bits, [0...nseq-1] */

} CMIS_SCORESET;


/* cmis_scoreset.c */
extern CMIS_SCORESET       *cmis_scoreset_Create(int nseq);
extern void                 cmis_scoreset_Destroy(CMIS_SCORESET *cmis_ss);
extern int                  cmis_scoreset_Write(FILE *fp, CMIS_SCORESET *cmis_ss);


#endif
```
