## Supplemental Code S1 for "Remote homology search with hidden Potts models": cmis_trace.c

```
/*cmis_trace.c */

#include <string.h>

#include "hmmer.h"
#include "p7_config.h"

#include "easel.h"
#include "esl_vectorops.h"

#include "cmis_trace.h"

int
cmis_trace_Copy(const P7_TRACE *src, P7_TRACE *dst)
{
   /* copy non-pointer objects */
   dst->N         = src->N;         /* length of traceback                               */
   dst->nalloc    = src->nalloc;    /* allocated length of traceback                     */
   dst->M         = src->M;         /* model length M (maximum k)                        */
   dst->L         = dst->L;         /* sequence length L (maximum i)                     */
   dst->ndom      = src->ndom;      /* number of domains in trace (= # of B or E states) */
   dst->ndomalloc = src->ndomalloc; /* current allocated size of these stacks            */

   /* copy over array objects */
   /* src->tr is a character array, not a NULL-terminated  "string" */
   /* => use memcpy(), not strcpy() */
   memcpy(dst->st, src->st, src->N);  /* state type code [0..N-1] */

   if (src->k != NULL)       esl_vec_ICopy(src->k, src->N, dst->k);                /* node index; 1..M if M,D,I; else 0 [0..N-1]  */
   if (src->i != NULL)       esl_vec_ICopy(src->i, src->N, dst->i);                /* pos emitted in dsq, 1..L; else 0  [0..N-1]  */
   if (src->pp != NULL)      esl_vec_FCopy(src->pp, src->N, dst->pp);              /* pos emitted in dsq, 1..L; else 0  [0..N-1]  */
   if (src->tfrom != NULL)   esl_vec_ICopy(src->tfrom, src->ndom, dst->tfrom);     /* locations of B states in trace (0..tr->N-1) */
   if (src->tto != NULL)     esl_vec_ICopy(src->tto, src->ndom, dst->tto);         /* locations of E states in trace (0..tr->N-1) */
   if (src->sqfrom != NULL)  esl_vec_ICopy(src->sqfrom, src->ndom, dst->sqfrom);   /* first M-emitted residue on sequence (1..L)  */
   if (src->hmmfrom != NULL) esl_vec_ICopy(src->hmmfrom, src->ndom, dst->hmmfrom); /* first M/D state on model (1..M)             */
   if (src->hmmto != NULL)   esl_vec_ICopy(src->hmmto, src->ndom, dst->hmmto);     /* last M/D state on model (1..M)              */

   return eslOK;
}

P7_TRACE *
cmis_trace_Clone(const P7_TRACE *tr) {

   P7_TRACE *tr2     = p7_trace_Create();
   int       status;

   /* make sure tr2 has enough memory allocated */
   p7_trace_GrowTo(tr2, tr->N);
   p7_trace_GrowIndexTo(tr2, tr->ndom);
   if ((status = cmis_trace_Copy(tr, tr2)) != eslOK) goto ERROR;

   return tr2;

   ERROR:
      p7_trace_Destroy(tr2);
      return NULL;

}
```
