## Supplemental Code S1 for "Remote homology search with hidden Potts models": cmis_trace.h

```
/* cmis_trace.h */

#ifndef CMISTRACE_INCLUDED
#define CMISTRACE_INCLUDED

#include <string.h>

#include "hmmer.h"
#include "p7_config.h"

#include "easel.h"
#include "esl_vectorops.h"

/* hpm_trace.c */
extern int       cmis_trace_Copy(const P7_TRACE *src, P7_TRACE *dst);
extern P7_TRACE *cmis_trace_Clone(const P7_TRACE *tr);

#endif
```
