## Supplemental Code S1 for "Remote homology search with hidden Potts models": cmscore_path.c

```
#include <stdio.h>
#include <stdlib.h>
#include <string.h>

#include "easel.h"
#include "esl_alphabet.h"
#include "esl_getopts.h"
#include "esl_msa.h"
#include "esl_msafile.h"
#include "esl_sq.h"

#include "infernal.h"
#include "config.h"

static ESL_OPTIONS options[] = {
   /* name        type        default    env range togs  reqs  incomp            help                                            docgroup */
   { "-h",        eslARG_NONE,    FALSE, NULL, NULL, NULL, NULL, NULL,            "help; show brief info on version and usage",  1 },

   /* Options forcing which alphabet we're working in (normally autodetected) */
   { "--amino",   eslARG_NONE,    FALSE, NULL, NULL, NULL, NULL, "--dna,--rna",   "We are dealing with protein sequences",       2 },
   { "--dna",     eslARG_NONE,    FALSE, NULL, NULL, NULL, NULL, "--amino,--rna", "We are dealing with dna sequences",           2 },
   { "--rna",     eslARG_NONE,    FALSE, NULL, NULL, NULL, NULL, "--amino,--dna", "We are dealing with rna sequences",           2 },

   { 0,0,0,0,0,0,0,0,0,0 },
};

static char banner[] = "Score sequence(s) and path(s) with a cm";
static char usage[]  = "[-options] <cmfile> <msafile>";

static void
cmdline_failure(char *argv0, char *format, ...)
{
   va_list argp;
   printf("\nERROR: ");
   va_start(argp, format);
   vfprintf(stderr, format, argp);
   va_end(argp);
   esl_usage(stdout, argv0, usage);
   printf("\nTo see more help on available options, do %s -h\n\n", argv0);
   exit(1);

}

int main(int argc, char *argv[]){

   ESL_GETOPTS      *go        = NULL;
   ESL_ALPHABET     *abc       = NULL;                 /* alphabet for both cm and msa                   */
   char             *cmfile    = NULL;                 /* input cm filepath                              */
   char             *msafile   = NULL;                 /* input msa filepath                             */
   CM_FILE          *cmfp      = NULL;                 /* open input CM file stream                      */
   ESL_MSAFILE      *afp       = NULL;                 /* open input msa file stream                     */
   CM_t             *cm        = NULL;                 /* cm                                             */
   ESL_MSA          *msa       = NULL;                 /* msa                                            */
   ESL_SQ          **sq        = NULL;                 /* array of sequences                             */
   Parsetree_t     **pstr      = NULL;                 /* array of parsetrees for input MSA              */
   Parsetree_t      *mtr       = NULL;                 /* the guide tree; for constructing pstr from msa */
   CM_MX            *mx        = NULL;                 /* DP matrix for inside algorithm                 */
   float             cmsc;                             /* Joint cm score of seq and path                 */
   float             insc;                             /* inside score of a sequence                     */
   int               n;                                /* msa sequence (row) index                       */
   int               fmt       = eslMSAFILE_STOCKHOLM; /* input MSA format.                              */
   int               status;                           /* easel return code                              */
   char              errbuf[eslERRBUFSIZE];            /* for verbose error reporting                    */

   /* parse command line */
   go = esl_getopts_Create(options);
   if (esl_opt_ProcessCmdline(go, argc, argv) != eslOK) cmdline_failure(argv[0], "Failed to parse command line: %s\n", go->errbuf);
   if (esl_opt_VerifyConfig(go)               != eslOK) cmdline_failure(argv[0], "Error in app configuration:   %s\n", go->errbuf);

   /* output help if requested */
   if (esl_opt_GetBoolean(go, "-h") ) {
      esl_banner(stdout, argv[0], banner);
      esl_usage (stdout, argv[0], usage);
      puts("\n where options are:");
      esl_opt_DisplayHelp(stdout, go, 1, 2, 80);
      puts("\n Alphabet options:");
      esl_opt_DisplayHelp(stdout, go, 2, 2, 80);
      exit(0);
   }

   if       (esl_opt_GetBoolean(go, "--amino")) abc = esl_alphabet_Create(eslAMINO);
   else if  (esl_opt_GetBoolean(go, "--dna"))   abc = esl_alphabet_Create(eslDNA);
   else if  (esl_opt_GetBoolean(go, "--rna"))   abc = esl_alphabet_Create(eslRNA);


   /* read arguments */
   if (esl_opt_ArgNumber(go) != 2)  cmdline_failure(argv[0], "Incorrect number of command line arguments.\n", go->errbuf);

   cmfile = esl_opt_GetArg(go, 1);
   msafile = esl_opt_GetArg(go, 2);

   /* read the cm file */
   status = cm_file_Open(cmfile, NULL, FALSE, &cmfp, errbuf);
   if      (status == eslENOTFOUND) cm_Fail("File existence/permissions problem in trying to open CM file %s.\n%s\n", cmfile, errbuf);
   else if (status == eslEFORMAT)   cm_Fail("File format problem in trying to open CM file %s.\n%s\n",                cmfile, errbuf);
   else if (status != eslOK)        cm_Fail("Unexpected error %d in opening CM file %s.\n%s\n",               status, cmfile, errbuf);

   /* read first cm in file */
   status = cm_file_Read(cmfp, TRUE, &abc, &cm);
   if      (status == eslEFORMAT)   cm_Fail("Bad file format in CM file %s:\n%s\n",          cmfp->fname, cmfp->errbuf);
   else if (status == eslEINCOMPAT) cm_Fail("CM in %s is not in the expected %s alphabet\n", cmfp->fname, esl_abc_DecodeType(abc->type));
   else if (status == eslEOF)       cm_Fail("Empty CM file %s? No CM data found.\n",         cmfp->fname);
   else if (status != eslOK)        cm_Fail("Unexpected error in reading CMs from %s\n",     cmfp->fname);
   cm_file_Close(cmfp);

   /* configure CM */
   cm->config_opts |= CM_CONFIG_SCANMX;
   cm_Configure(cm, errbuf, -1);

   /* set search options */
   cm->search_opts |= CM_SEARCH_INSIDE;
   cm->search_opts |= CM_SEARCH_NONBANDED;

   /* initlalize logsum lookup table magic */
   init_ilogsum();
   FLogsumInit();

   /* print all configuration flags */
   DumpCMFlags(stdout, cm);

   /* allow '-' and '.' as gap */
   esl_alphabet_SetEquiv(abc, '.', '-');

   /* read the msa file */
   if ((status = esl_msafile_Open(&abc, msafile, NULL, fmt, NULL, &afp)) != eslOK) esl_msafile_OpenFailure(afp, status);
   if ((status = esl_msafile_Read(afp, &msa))                            != eslOK) esl_fatal ("Failed to read MSA");
   esl_msafile_Close(afp);

   /* extract sequences from MSA */
   ESL_ALLOC(sq, sizeof(ESL_SQ *) * msa->nseq);
   for (n = 0; n < msa->nseq; n++) {
      fprintf(stdout, "n: %d\n", n);
      esl_sq_FetchFromMSA(msa, n, &sq[n]);
   }

   if (! (msa->flags & eslMSA_DIGITAL)) esl_fatal("need a digital msa");
   if (msa->rf == NULL)                 esl_fatal("msa lacks an RF line");

   /* create guide tree from MSA; needed for making parse tree array */
   status = HandModelmaker(msa, errbuf,
                           TRUE,  /* use_rf */
                           FALSE, /* use_el, no */
                           FALSE, /* use_wts, irrelevant */
                           0.5,   /* gapthresh, irrelevant */
                           NULL,  /* returned CM, irrelevant */
                           &mtr); /* guide tree */

   if (status != eslOK)  ESL_XFAIL(status, errbuf, "Issue running HandModelmaker()\n");

   /* create parse tree alignment from MSA */
   Alignment2Parsetrees(msa, cm, mtr, errbuf, NULL, &pstr);

   /* initiate DP matrix */
   mx = cm_mx_Create(cm->M);

   /* score sequences and paths */
   for (n=0;  n < msa->nseq; n++) {

      /* run inside algroithm */
      cm_InsideAlign(cm, errbuf, sq[n]->dsq, sq[n]->n, 512.0, mx, &insc);

      /* score sequence and given path */
      ParsetreeScore(cm, NULL, errbuf, pstr[n], sq[n]->dsq, 0, &cmsc, NULL, NULL, NULL, NULL);

      ParsetreeDump(stdout, pstr[n], cm, sq[n]->dsq);

      fprintf(stdout, "seq: %s\n", sq[n]->name);
      fprintf(stdout, "Inside score: %.2f\n", insc);
      fprintf(stdout, "CM score of sequence, path: %.2f\n",  cmsc);
      fprintf(stdout, "Log posterior probability of path under CM: %.2f\n", cmsc - insc);
   }

   fprintf(stdout, "hello world\n");

   /* clean up and return */
   for (n = 0; n < msa->nseq; n++) {
      esl_sq_Destroy(sq[n]);
      FreeParsetree(pstr[n]);
   }
   free(pstr);
   free(sq);
   FreeParsetree(mtr);
   esl_msa_Destroy(msa);
   FreeCM(cm);
   cm_mx_Destroy(mx);
   esl_alphabet_Destroy(abc);
   esl_getopts_Destroy(go);
   return 0;

   ERROR:
      return status;

}
```
