## Supplemental Code S1 for "Remote homology search with hidden Potts models": cmscoreISh4.c

```
#include <stdio.h>
#include <stdlib.h>
#include <string.h>

#include "easel.h"
#include "esl_alphabet.h"
#include "esl_getopts.h"
#include "esl_msa.h"
#include "esl_random.h"
#include "esl_sq.h"
#include "esl_sqio.h"
#include "esl_vectorops.h"

/* h4 nwo includes */
#include "h4_hmmfile.h"
#include "h4_mode.h"
#include "h4_profile.h"
#include "h4_refmx.h"
#include "general.h"
#include "logsum.h"
#include "reference_dp.h"

/* infernal includes */
#include "infernal.h"
#include "config.h"

/* CMIS includes */
#include "cmis_scoreset.h"
#include "cmis_trace.h"
#include "h4_path_cmis.h"
#include "h4_pathalign.h"

/* declaration of internal functions */
int Calculate_IS_scores(CM_t *cm, H4_PROFILE *hmm, ESL_SQ **sq, ESL_RANDOMNESS *rng, CMIS_SCORESET *cmis_ss, ESL_MSA **msa, char *scoreprogfile, char *aliprogfile, int start, int end, int totseq, int user_R, int A, int verbose);
int map_ss_cons(CM_t *cm, ESL_MSA *msa, char *errbuf);
int find_jumps(double *pr_unsorted, int r, int R_batch, ESL_SQ *sq, H4_PATH **pi, H4_PROFILE *hmm, H4_MODE *mo, float fsc, CM_t *cm, Parsetree_t **pstr, char *errbuf);
char *strsafe(char *dest, const char *src);

static ESL_OPTIONS options[] = {
        /* name              type        default    env range togs  reqs  incomp            help                                                     docgroup */
        { "-h",              eslARG_NONE,    FALSE, NULL, NULL, NULL, NULL, NULL,            "help; show brief info on version and usage",               1 },

        /* Options forcing which alphabet we're working in (normally autodetected) */
        { "--amino",         eslARG_NONE,    FALSE, NULL, NULL, NULL, NULL, "--dna,--rna",   "We are dealing with protein sequences",                    2 },
        { "--dna",           eslARG_NONE,    FALSE, NULL, NULL, NULL, NULL, "--amino,--rna", "We are dealing with dna sequences",                        2 },
        { "--rna",           eslARG_NONE,    FALSE, NULL, NULL, NULL, NULL, "--amino,--dna", "We are dealing with rna sequences",                        2 },

        /* options for bounding sequences we score */
        { "--seqstart",      eslARG_INT,     "-1",  NULL, NULL, NULL, NULL, NULL,            "Start sequence index",                                     3 },
        { "--seqend",        eslARG_INT,     "-2",  NULL, NULL, NULL, NULL, NULL,            "End sequence index",                                       3 },

        /* options for controlling number of alignments sampled per sequence */
        { "-R",              eslARG_INT,     "-1",  NULL, NULL, NULL, NULL, NULL,            "Number of HMM paths sampled per sequence",                 4 },
        { "-s",              eslARG_INT,     "0",   NULL, NULL, NULL, NULL, NULL,            "Set random number seed to <n>",                            4 },

        /* control of output */
        { "-A",              eslARG_OUTFILE, NULL,  NULL, NULL, NULL, NULL, NULL,            "save multiple alignment of all seqs to file <s>",          5 },
        { "--scoreprog",     eslARG_OUTFILE, NULL,  NULL, NULL, NULL, NULL, NULL,            "save .csv with info on IS scoring progress to file <s>",   5 },
        { "--aliprog",       eslARG_OUTFILE, NULL,  NULL, NULL, NULL, "-A", NULL,            "save .csv with info on IS alignment prgoress to file <s>", 5 },

        /* debugging tools */
        { "-v",              eslARG_NONE,    FALSE, NULL, NULL, NULL, NULL, NULL,            "Verbose mode: print info on intermediate scoring steps",   6 },

        { 0,0,0,0,0,0,0,0,0,0 },
};

static char banner[] = "Align and score sequences with a CM by importance sampling a CM/SCFG";
static char usage[]  = "[-options] <cmfile> <hmmfile> <seqfile> <score_outfile>";

static void
cmdline_failure(char *argv0, char *format, ...)
{
        va_list argp;
        printf("\nERROR: ");
        va_start(argp, format);
        vfprintf(stderr, format, argp);
        va_end(argp);
        esl_usage(stdout, argv0, usage);
        printf("\nTo see more help on available options, do %s -h\n\n", argv0);
        exit(1);
}

int main (int argc, char *argv[]) {
   ESL_GETOPTS      *go               = NULL;                 /* application configuration                */
   ESL_ALPHABET     *abc              = NULL;                 /* biological alphabet                      */
   ESL_RANDOMNESS   *rng              = NULL;                 /* random number generator                  */
   char             *cmfile           = NULL;                 /* input cm filepath                        */
   char             *hmmfile          = NULL;                 /* input hmm filepath                       */
   char             *seqfile          = NULL;                 /* input seq filepath                       */
   char             *scorefile        = NULL;                 /* output score filepath                    */
   char             *scoreprogfile    = NULL;                 /* output IS scoring progress filepath      */
   char             *aliprogfile      = NULL;                 /* output IS alignment progress filepath    */
   CM_FILE          *cmfp             = NULL;                 /* open input CM file stream                */
   H4_HMMFILE       *hmmfp            = NULL;                 /* open input hmm file stream               */
   ESL_SQFILE       *sqfp             = NULL;                 /* open seq file stream                     */
   CM_t             *cm               = NULL;                 /* cm                                       */
   H4_PROFILE       *hmm              = NULL;                 /* hmm                                      */
   ESL_SQ          **sq               = NULL;                 /* array of sequences                       */
   CMIS_SCORESET    *cmis_ss          = NULL;                 /* for sequence scores                      */
   int               format           = eslSQFILE_UNKNOWN;    /* seq file format                          */
   FILE             *afp              = NULL;                 /* output alignment file (-A)               */
   ESL_MSA          *msa              = NULL;                 /* alignment output object (-A)             */
   int               outfmt           = eslMSAFILE_STOCKHOLM; /* alignment output format (-A)             */
   FILE             *cmis_ss_fp       = NULL;                 /* file for sequence scores                 */
   int               n                =  0;                   /* seq_index                                */
   int               nseq             =  0;                   /* number of seqs in seq file               */
   int               totseq           =  0;                   /* number of seqs we deal with              */
   int               start,end;                               /* start and end seq indices                */
   int               R                = -1;                   /* number of ali samples/seq                */
   int               A                = 0;                    /* Boolean for output alignment             */
   int               v                = 0;                    /* Boolean for verbose mode                 */
   int               status;                                  /* easel return code                        */
   char              errbuf[eslERRBUFSIZE];


   /* parse command line */
   go = esl_getopts_Create(options);
   if (esl_opt_ProcessCmdline(go, argc, argv) != eslOK) cmdline_failure(argv[0], "Failed to parse command line: %s\n", go->errbuf);
   if (esl_opt_VerifyConfig(go)               != eslOK) cmdline_failure(argv[0], "Error in app configuration:   %s\n", go->errbuf);

   if (esl_opt_GetBoolean(go, "-h") ) {
      esl_banner(stdout, argv[0], banner);
      esl_usage (stdout, argv[0], usage);
      puts("\n where options are:");
      esl_opt_DisplayHelp(stdout, go, 1, 2, 80);
      puts("\n Alphabet options:");
      esl_opt_DisplayHelp(stdout, go, 2, 2, 80);
      puts("\n Options for bounding sequences we score:");
      esl_opt_DisplayHelp(stdout, go, 3, 2, 80);
      puts("\n Sampling options:");
      esl_opt_DisplayHelp(stdout, go, 4, 2, 80);
      puts("\n MSA Output options:");
      esl_opt_DisplayHelp(stdout, go, 5, 2, 80);
      puts("\n Debug options:");
      esl_opt_DisplayHelp(stdout, go, 6, 2, 80);
      exit(0);
   }

   /* read arguments */
   if (esl_opt_ArgNumber(go) != 4)  cmdline_failure(argv[0], "Incorrect number of command line arguments.\n", go->errbuf);

   cmfile    =  esl_opt_GetArg(go, 1);
   hmmfile   =  esl_opt_GetArg(go, 2);
   seqfile   =  esl_opt_GetArg(go, 3);
   scorefile =  esl_opt_GetArg(go, 4);

   /* check for verbose mode */
   if (esl_opt_GetBoolean(go, "-v")) v=1;

   /* if output msa requested by user, try to open it */
   if (esl_opt_IsOn(go, "-A")) {
      if ((afp = fopen(esl_opt_GetString(go, "-A"), "w")) == NULL) esl_fatal("Failed to open alignment file %s for writing\n", esl_opt_GetString(go, "-A"));
      A = 1;
   }

   /* if IS intermediate scoring progress csv is requested by user, set variables*/
   if (esl_opt_IsOn(go, "--scoreprog")) {
      scoreprogfile = (esl_opt_GetString(go, "--scoreprog"));
   }

   /* if IS intermediate alignment progress csv is requested by user, set variables*/
   if (esl_opt_IsOn(go, "--aliprog")) {
      aliprogfile = (esl_opt_GetString(go, "--aliprog"));
   }


   /* if user has defined an alphabet we define it here */
   if        (esl_opt_GetBoolean(go, "--amino"))       abc = esl_alphabet_Create(eslAMINO);
   else if   (esl_opt_GetBoolean(go, "--rna"))         abc = esl_alphabet_Create(eslRNA);
   else if   (esl_opt_GetBoolean(go, "--dna"))         abc = esl_alphabet_Create(eslDNA);

   /* check if user has specified the number of samples/sequence */
   if (esl_opt_GetInteger(go, "-R") > 0) R = esl_opt_GetInteger(go, "-R");

   /* create random number generator */
   rng = esl_randomness_Create(esl_opt_GetInteger(go, "-s"));

   /* open the .hmm file */
   if ( h4_hmmfile_Open(hmmfile, NULL, &hmmfp) != eslOK) esl_fatal("couldn't open profile file %s", hmmfile);
   /* read first hmm  from hmm file */
   if ( h4_hmmfile_Read(hmmfp, &abc, &hmm)     != eslOK) esl_fatal("failed to read profile from file %s", hmmfile);
   h4_hmmfile_Close(hmmfp);

   /* open the .cm file */
   status = cm_file_Open(cmfile, NULL, FALSE, &cmfp, errbuf);
   if      (status == eslENOTFOUND) cm_Fail("File existence/permissions problem in trying to open CM file %s.\n%s\n", cmfile, errbuf);
   else if (status == eslEFORMAT)   cm_Fail("File format problem in trying to open CM file %s.\n%s\n",                cmfile, errbuf);
   else if (status != eslOK)        cm_Fail("Unexpected error %d in opening CM file %s.\n%s\n",               status, cmfile, errbuf);

   /* read first cm in file */
   status = cm_file_Read(cmfp, TRUE, &abc, &cm);
   if      (status == eslEFORMAT)   cm_Fail("Bad file format in CM file %s:\n%s\n",          cmfp->fname, cmfp->errbuf);
   else if (status == eslEINCOMPAT) cm_Fail("CM in %s is not in the expected %s alphabet\n", cmfp->fname, esl_abc_DecodeType(abc->type));
   else if (status == eslEOF)       cm_Fail("Empty CM file %s? No CM data found.\n",         cmfp->fname);
   else if (status != eslOK)        cm_Fail("Unexpected error in reading CMs from %s\n",     cmfp->fname);

   cm_file_Close(cmfp);

   /* check that cm, hmm have same # of match states */
   //if (hmm->M != cm->clen) esl_fatal("The hmm and cm have different numbers of consensus columns!\n hmm->M: %d \n cm->M: %d\n", hmm->M, cm->clen);

   /* configure CM */
   cm->config_opts |= CM_CONFIG_SCANMX;
   cm_Configure(cm, errbuf, -1);

   /* set search options */
   cm->search_opts |= CM_SEARCH_INSIDE;
   cm->search_opts |= CM_SEARCH_NONBANDED;

   /* initlalize logsum lookup table magic */
   init_ilogsum();
   FLogsumInit();
   h4_logsum_Init();

   /* print all configuration flags */
   DumpCMFlags(stdout, cm);

   /* open the sequence file */
   status = esl_sqfile_OpenDigital(abc, seqfile, format, NULL, &sqfp);
   if      (status == eslENOTFOUND) esl_fatal("No such file.");
   else if (status == eslEFORMAT)   esl_fatal("Format unrecognized.");
   else if (status ==eslEINVAL)     esl_fatal("Can't autodetect stdin or .gz.");
   else if (status != eslOK)        esl_fatal("Open failed, code %d.", status);

   /* read sequences into array */
   ESL_REALLOC(sq, sizeof(ESL_SQ *) * (n + 1));
   sq[n] = esl_sq_CreateDigital(abc);
   while ((status = esl_sqio_Read(sqfp, sq[n+nseq])) == eslOK) {
      nseq++;
      ESL_REALLOC(sq, sizeof(ESL_SQ *) * (n+nseq+1));
      sq[n+nseq] = esl_sq_CreateDigital(abc);
   }

   /* error handling and cleanup */
   if      (status == eslEFORMAT) esl_fatal("Parse failed (sequence file %s):\n%s\n",
                                          sqfp->filename, esl_sqfile_GetErrorBuf(sqfp));
   else if (status != eslEOF)     esl_fatal("Unexpected error %d reading sequence file %s",
                                            status, sqfp->filename);

   /* if user specified bounds, set them here */
   start = 0;
   end   = nseq;
   if (esl_opt_GetInteger(go, "--seqstart") > -1) start = esl_opt_GetInteger(go, "--seqstart");
   if (esl_opt_GetInteger(go, "--seqend")   > -1)     end = esl_opt_GetInteger(go, "--seqend");
   /* check if end is beyond number of sequences in input file */
   if (end > nseq) end = nseq;


   /* total number of sequences we will be scoring */
   totseq = end-start;
   fprintf(stdout, "totseq: %d\n", totseq);

   /* create scoreset object */
   cmis_ss = cmis_scoreset_Create(totseq);

   /* calculate importance sampling score for all seqs */
   if ((status =Calculate_IS_scores(cm, hmm, sq, rng, cmis_ss, &msa, scoreprogfile, aliprogfile, start, end, totseq, R, A, v)) != eslOK) {
      esl_fatal("Error running Calculate_IS_scores(), returned code %d\n", status);
   }

   /* write score file */
   if ((cmis_ss_fp = fopen(scorefile, "w")) == NULL) esl_fatal("Failed to open output CMIS scoreset file %s for writing", scorefile);
   if (cmis_scoreset_Write(cmis_ss_fp, cmis_ss) != eslOK) esl_fatal("Failed to write scores to CMIS scoreset file %s",    scorefile);
   fclose(cmis_ss_fp);

   if(cm->flags & CMH_LOCAL_BEGIN) fprintf(stdout, "Local begins on!\n");
   if(cm->flags & CMH_LOCAL_END) fprintf(stdout, "Local ends on!\n");

   /* if output msa requested, write it to output file */
   if (A) {
      esl_msafile_Write(afp, msa, outfmt);
      fclose(afp);
      esl_msa_Destroy(msa);
   }

   /* clean up and return */
   for (n = 0; n < nseq+1; n++)
   {
      esl_sq_Destroy(sq[n]);
   }

   esl_sqfile_Close(sqfp);
   free(sq);
   FreeCM(cm);
   cmis_scoreset_Destroy(cmis_ss);
   h4_profile_Destroy(hmm);
   esl_randomness_Destroy(rng);
   esl_alphabet_Destroy(abc);
   esl_getopts_Destroy(go);
   return 0;

   ERROR:
   return status;
}

int Calculate_IS_scores(CM_t *cm, H4_PROFILE *hmm, ESL_SQ **sq, ESL_RANDOMNESS *rng, CMIS_SCORESET *cmis_ss,
                        ESL_MSA **msa, char *scoreprogfile, char *aliprogfile, int start, int end, int totseq,
                        int user_R, int A, int verbose) {

   H4_MODE        *mo         = h4_mode_Create();           /* h4 profile hmm mode                           */
   H4_REFMX       *fwd        = h4_refmx_Create(100, 100);  /* forward DP matrix                             */
   ESL_SQ        **sq_dummy;                                /* dummy sequence array for sampled alignments   */
   ESL_SQ        **out_sq      = NULL;                      /* traces used to construct output msa (-A)      */
   H4_PATH       **pi_dummy;                                /* dummy path array for sampled alignments       */
   H4_PATH       **out_pi      = NULL;                      /* paths used to construct output MSA            */
   CM_MX          *mx           = cm_mx_Create(cm->M);      /* DP matrix for running inside algorithm        */
   FILE           *scoreprogfp = NULL;                      /* score progress output file (--scoreprog)      */
   FILE           *aliprogfp   = NULL;                      /* alignment progress output file (--aliprog)    */
   int             i,j;                                     /* sequence indices                              */
   int             b;                                       /* batch index                                   */
   int             r,s;                                     /* alignment sample indices                      */
   int             R           = 1000;                      /* number of sampled alignments per sequence     */
   //int             R_batch     = 10;                        /* sampled alignment batch size                  */
   int             R_batch     = 1000;                      /* sampled alignment batch size                  */
   int             N_batch;                                 /* total number of batches                       */
   int             nBetter;                                 /* number of better alignments we've found       */
   int             allcons     = 1;                         /* bool for keeping all consensus columns in MSA */
   float           fsc;                                     /* forward partial log-odds score, in bits       */
   float           hmmsc_ld;                                /* hmm log odds S(x,pi), in bits                 */
   float           cmsc_ld;                                 /* cm log odds S(x, pi), in bits                 */
   float           cmsc_max;                                /* best cm log odds S(x, pi) for all paths       */
   float           insc;                                    /* log-odds score for inside algorithm           */
   float           ldprev;                                  /* previous IS log-odds score                    */
   float           ld;                                      /* importance sampling log odds score S(x)       */
   float           ls;                                      /* log of importance sampling sum                */
   double         *pr, *pr_unsorted;                        /* terms in importance sampling sum              */
   int             status;                                  /* easel return code                             */
   char            errbuf[eslERRBUFSIZE];                   /* buffer for easel errors                       */

   /* if score progress file is requested, open it */
   if (scoreprogfile) {
      if ((scoreprogfp = fopen(scoreprogfile, "w")) == NULL) p7_Fail("Failed to open file %s for writing\n", scoreprogfile);
      fprintf(scoreprogfp, "id,iter,inside_logodds,cmis_logodds\n");
   }

   /* if alignment progress file is requested, open it */
   if (aliprogfile) {
      if ((aliprogfp = fopen(aliprogfile, "w")) == NULL) p7_Fail("Failed to open file %s for writing\n", aliprogfile);
      fprintf(aliprogfp, "id,iter,cmis_logodds\n");
   }

   if (A) {
      /* allocate memory for paths for output MSA */
      ESL_ALLOC(out_pi, sizeof(H4_PATH *) * totseq);
      ESL_ALLOC(out_sq, sizeof(ESL_SQ *) * (totseq));
   }

   /* set number of samples */
   if (user_R > 0) R = user_R;

   /* calculate number of batches */
   N_batch = (R / R_batch) + (R % R_batch != 0);
   fprintf(stdout, "N_batch: %d\n", N_batch);

   /* allocate space for dummy trace  and sequence arrays */
   ESL_ALLOC(sq_dummy, sizeof(P7_TRACE *) * R_batch);
   ESL_ALLOC(pi_dummy, sizeof(H4_PATH *) * R_batch);
   for (r = 0; r < R_batch; r++) {
      sq_dummy[r] = esl_sq_CreateDigital(hmm->abc);
      pi_dummy[r] = h4_path_Create();
   }

   /* allocate array to store terms in importance sampling sum */
   ESL_ALLOC(pr, R*sizeof(double));
   ESL_ALLOC(pr_unsorted, R*sizeof(double));
   esl_vec_DSet(pr, R, 0.0);
   esl_vec_DSet(pr_unsorted, R, 0.0);


   /* set HMM mode to uniglocal  */
   h4_mode_SetUniglocal(mo);

   /* set up cm for scoring */
   if((status = CMLogoddsify(cm)) != eslOK) ESL_FAIL(status, errbuf, "problem logodisfying CM");


    /* outer loop over sequences */
   for (i=start; i < end; i++) {

      nBetter = 0;
      ldprev = 0.0;


      /* index for score set: must start at 0 */
      j=i-start;

      /* for getting optimal path under hpm */
      if (A) {
         cmsc_max  = -eslINFINITY;
         out_sq[j] = esl_sq_CreateDigital(hmm->abc);
         esl_sq_Copy(sq[i], out_sq[j]);
      }

      /* set profile length */
      h4_mode_SetLength(mo, sq[i]->n);

      /* run forward algorithm */
      h4_reference_Forward(sq[i]->dsq, sq[i]->n, hmm, mo, fwd, &fsc);

      //fprintf(stdout, "\nseq: %s\n", sq[i]->name);
      //fprintf(stdout, "fsc: %.2f\n", fsc);
      //fprintf(stdout, "nullsc: %.2f\n", mo->nullsc);
      //fprintf(stdout, "forward score: %.2f bits\n", fsc - mo->nullsc);


      /* run inside algorithm */
      cm_InsideAlign(cm, errbuf, sq[i]->dsq, sq[i]->n, 512.0, mx, &insc);

      /* prep for sampling paths */
      esl_vec_DSet(pr, R, 0.0);
      esl_vec_DSet(pr_unsorted, R, 0.0);

      /* middle loop over batches of sampled alignments */
      for (b = 0; b < N_batch; b++) {
         ESL_MSA        *msa_dummy;                            /* MSA of different aligmnetss of seq i  */
         Parsetree_t    *mtr       = NULL;                     /* the guide tree                        */
         Parsetree_t   **pstr      = NULL;                     /* array of parse trees created from MSA */

         /* inner loop 1: sample alignments from HMM */
         for (s = 0;  s < R_batch; s++) {

            /* calculte overall sample number */
            r = b*R_batch + s;

            /* if we've sampled as much as requested, skip inner loop 1 */
            if (r >= R) break;

            /* copy sequence into dummy array */
            esl_sq_Copy(sq[i], sq_dummy[s]);

            /* get stochastic traceback */
            h4_reference_StochasticTrace(rng, NULL, hmm, mo, fwd, pi_dummy[s]);

            //h4_path_Dump(stdout, pi_dummy[s]);

            h4_path_Score(pi_dummy[s], sq[i]->dsq, hmm, mo, &hmmsc_ld);

            pr[r] -= hmmsc_ld;

         }

         /* in between inner loops: convert h4 paths to MSA to parse trees */

         /* step 1: convert paths to an MSA */

         status = h4_pathalign_Seqs(sq_dummy, pi_dummy, R_batch, hmm->M, allcons, hmm, &msa_dummy);
         //fprintf(stdout, "i: %d, h4_pathalign_Seqs() return code: %d\n", i, status);


         map_ss_cons(cm, msa_dummy, errbuf);
         //int outfmt = eslMSAFILE_STOCKHOLM;
         //FILE *afp = NULL;
         //afp = fopen("foo.sto", "w");
         //esl_msafile_Write(afp, msa_dummy, outfmt);
         //fclose(afp);

         /* create guide tree */
         status = HandModelmaker(msa_dummy, errbuf,
                                 TRUE,  /* use_rf */
                                 FALSE, /* use_el, no */
                                 FALSE, /* use_wts, irrelevant */
                                 0.5,   /* gapthresh, irrelevant */
                                 NULL,  /* returned CM, irrelevant */
                                 &mtr); /* guide tree */

         if (status != eslOK)  ESL_XFAIL(status, errbuf, "Issue running HandModelmaker()\n");

         /* create parse tree alignment from MSA */
         Alignment2Parsetrees(msa_dummy, cm, mtr, errbuf, NULL, &pstr);

         /* inner loop 2: score traces with CM */
         for (s = 0; s < R_batch; s++) {

            /* calculate overall sample number */
            r = b*R_batch + s;

            /* if we've sampled enough, skip scoring and go to clean up */
            if (r >= R) goto cleanup;


            /* score this sequence and alignment */
            ParsetreeScore(cm,
                           NULL,
                           errbuf,
                           pstr[s],
                           sq[i]->dsq,
                           0,
                           &cmsc_ld,
                           NULL,
                           NULL,
                           NULL,
                           NULL);

             pr[r] += cmsc_ld;

            /* for output alignment , see if we've got a better path */
            if (A && cmsc_ld > cmsc_max) {
               cmsc_max = cmsc_ld;

               /* if we're tracking alignment progress, print S(x,pi) of new best alignment */
               if (aliprogfile) fprintf(aliprogfp, "%s,%d,%.2f\n", sq[i]->name, r, cmsc_max);

               /* if we've already created out_pi[j], free it before recreating it */
               if (nBetter > 0) h4_path_Destroy(out_pi[j]);
               out_pi[j] = h4_path_Clone(pi_dummy[s]);
               nBetter++;
            }


         }

         /* if requested, track intermediate scoring progress for this batch */
         if (scoreprogfile) {
            esl_vec_DCopy(pr,r+1, pr_unsorted);
            esl_vec_DSortIncreasing(pr, r+1);
            float ls = esl_vec_DLog2Sum(pr, r+1);
            float ld = fsc - log2f(r+1) + ls;

            //fprintf(stdout, "test!\n");
            fprintf(stdout, "r: %d, intermediate score: %2f\n", r, ld);

            fprintf(scoreprogfp, "%s,%d,%.2f,%.2f\n", sq[i]->name, r, insc, ld);

            /* look for paths that make IS scores jump */
            if ( (r > 1e6) && (ld-ldprev > 0.25))  find_jumps(pr_unsorted, r, R_batch, sq[i], pi_dummy, hmm, mo, fsc, cm, pstr, errbuf);

             ldprev = ld;
         }


         /* clean up for next iteration of middle loop */
         cleanup: for (s = 0; s < R_batch; s++) {
            esl_sq_Reuse(sq_dummy[s]);
            h4_path_Reuse(pi_dummy[s]);
            FreeParsetree(pstr[s]);
         }

         esl_msa_Destroy(msa_dummy);
         FreeParsetree(mtr);
         free(pstr);
      }

      /* calculate IS approximation for this sequence */
      /* sort to increase accuracy if LogSum */
      esl_vec_DSortIncreasing(pr, R);

      /* run log sum */
      ls = esl_vec_DLog2Sum(pr, R);

      /* incorporate alignment-independent bits into score */
      ld = fsc - log2f(R) + ls;

      /* add scoring info to scoreset object */
      cmis_ss->sqname[j]   = sq[i]->name;
      cmis_ss->R[j]        = R;
      cmis_ss->fsc[j]      = fsc;
      cmis_ss->ntsc[j]     = mo->nullsc;
      cmis_ss->insc[j]     = insc;
      cmis_ss->scansc[j]   = 0.0;
      cmis_ss->seq_from[j] = -1;
      cmis_ss->seq_to[j]   = -2;
      cmis_ss->cmis_ld[j]  = ld;

   }

   /* if output msa requested, create it from traces */
   if (A) {
      h4_pathalign_Seqs(out_sq, out_pi, totseq, hmm->M, TRUE, hmm, msa);
   }

   /* clean up and return */
   if (A) {
      for (j=0; j<totseq; j++) {
         h4_path_Destroy(out_pi[j]);
         esl_sq_Destroy(out_sq[j]);
      }
      free(out_pi);
      free(out_sq);
   }
   if (scoreprogfp) fclose(scoreprogfp);
   if (aliprogfp) fclose(aliprogfp);
   for (s = 0; s < R_batch; s++) {
      esl_sq_Destroy(sq_dummy[s]);
      h4_path_Destroy(pi_dummy[s]);
   }
   free(sq_dummy);
   free(pi_dummy);
   free(pr);
   free(pr_unsorted);
   h4_refmx_Destroy(fwd);
   h4_mode_Destroy(mo);
   cm_mx_Destroy(mx);
   return eslOK;

   ERROR:
      return status;
}


/* Function: find_jumps()
 *
 * Purpose:  Poor importance sampling often shows sudden "jumps" in bitscore
 *           due to a single sample. Given a set of <R_batch> sampled paths
 *           of sequence <sq> through model <hmm>, find the path(s) that
 *           cause(s) the jump in score. Print S_HMM(x,pi) and S_CM(x,pi) to
 *           stdout. Also, output the alignment to an output msafile in the
 *           current working directly.
 *
 *           In case this isn't clear, this function is mostly for debugging.
 *           It's only called with the --scoreprog option set.
 *
 *
 *
 * args:    pr_unsorted   - vector containing ratios of bitscores. At least
 *                          the last <R_batch> elements are in the order in
 *                          which they were sampled.
 *          r             - sample index of the **last** path in the batch.
 *          sq            - sequence we are scoring
 *          pi            - <R_batch>-sized path array with batch of sampled
 *                          paths.
 *          hmm           - HMM used for generating paths.
 *          fsc           - forward score of <sq> under HMM, in bits.
 *          cm            - cm, only used here for mapping ss_cons to output
 *                          MSA('s)
 *          errbuf        - Error information. Feature more for the future tbh.
 *
 * Returns: eslOK on success, eslERROR if call to map_ss_cons() has issues.
 *
 * */

int find_jumps(double *pr_unsorted, int r, int R_batch, ESL_SQ *sq, H4_PATH **pi, H4_PROFILE *hmm, H4_MODE *mo, float fsc, CM_t *cm, Parsetree_t **pstr, char *errbuf){
   double    *pr_sorted;                                  /* vector of log-odds ratios, sorted from largest to smallest */
   float      ls;                                         /* log_sum of pr_sorted                                       */
   float      ld;                                         /* IS log odds score                                          */
   float      ldprev;                                     /* Previous iteration's IS log-odds score                     */
   float      hmmsc_ld;                                   /* log-odds score of a sequence and path given <hmm>          */
   int        s,t;                                        /* IS iteration indices                                       */
   H4_PATH  **out_pi             = NULL;                  /* 1-element path array for outputting path to an MSA         */
   ESL_SQ   **out_sq;                                     /* 1-element seq array for outputting path to an MSA          */
   ESL_MSA  *out_msa             = NULL;                  /* output MSA object                                          */
   FILE      *afp                = NULL;                  /* output alignment file                                      */
   char       out_msa_path[200];                          /* output MSA filepath                                        */
   char       sqname_safe[100];                           /* sequence name with '/' replace with '_'                    */
   int        outfmt             = eslMSAFILE_STOCKHOLM;  /* output MSA format. I am unwavering on this matter.         */
   int        njump              = 0;                     /* keep track of number of jumping paths in this batch        */
   int        status             = 0;                     /* esl return code                                            */

   strsafe(sqname_safe, sq->name);

   //fprintf(stdout, "in find jumps. r: %d\n", r);
   //fprintf(stdout, "sqname_safe: %s\n", sqname_safe);

   /* allocate memory */
   ESL_ALLOC(out_pi, sizeof(H4_PATH *));
   ESL_ALLOC(out_sq, sizeof(ESL_SQ *));
   ESL_ALLOC(pr_sorted, r*sizeof(double));

   /* create the output sequence */
   out_sq[0] = esl_sq_CreateDigital(hmm->abc);
   esl_sq_Copy(sq, out_sq[0]);

   /* figure out what the IS score was before this batch */
   s = r-R_batch;
   /* copy pr_unsorted to pr_unsorted */
   esl_vec_DCopy(pr_unsorted, r-R_batch+1, pr_sorted);
   /* sort pr_sorted for more accurate log summing */
   esl_vec_DSortIncreasing(pr_sorted, s+1);
   ls = esl_vec_DLog2Sum(pr_sorted,s+1);
   /* calcuate full log odds-score */
   ldprev = fsc - log2f(r-R_batch+1) + ls;

   /* now loop through paths and find jump(s) */
   for (s = r-R_batch+1; s<r; s++){

      /* index w/in batch: t = 0,...,R_batch-1 */
      t = s-r+R_batch-1;

      fprintf(stdout, "\tt: %d\n", t);

      /* calculate log-odds score at this iteration */
      esl_vec_DCopy(pr_unsorted,s+1, pr_sorted);
      esl_vec_DSortIncreasing(pr_sorted, s+1);
      ls = esl_vec_DLog2Sum(pr_sorted, s+1);
      ld = fsc - log2f(s+1) + ls;

      /* calculate HMM score of this seq and sampled path */
      h4_path_Score(pi[t], sq->dsq, hmm, mo, &hmmsc_ld);

      /* put path that causes jump in an output MSA file */
      if (fabsf(ld-ldprev) > 0.25) {
         if (njump > 0) {
            h4_path_Destroy(out_pi[0]);
            esl_msa_Destroy(out_msa);
         }

         out_pi[0] = h4_path_Clone(pi[t]);
         h4_pathalign_Seqs(out_sq, out_pi, 1, hmm->M, TRUE, hmm, &out_msa);

         /* map cm's SS_cons onto MSA */
         if (map_ss_cons(cm, out_msa, errbuf) != eslOK) ESL_FAIL(eslFAIL, errbuf, "Issue running map_ss_cons from find_jumps\n");

         /* create unique output MSA path */
         sprintf(out_msa_path,"%s_path_%d.sto", sqname_safe, s);

         /* write the msa file */
         afp = fopen(out_msa_path, "w");
         esl_msafile_Write(afp, out_msa, outfmt);
         fclose(afp);

         /* print info about jumping path */
         fprintf(stdout, "\nJumping IS score for sequence %s, path %d\n", sq->name, s);
         fprintf(stdout, "IS score at previous iteration:                  %.2f\n", ldprev);
         fprintf(stdout, "IS score at this iteration:                      %.2f\n", ld);
         fprintf(stdout, "CM log-odds score for this seq, path:            %.2f\n",  pr_unsorted[s] + hmmsc_ld);
         fprintf(stdout, "HMM log-odds score for this seq, path:           %.2f\n",  hmmsc_ld);
         fprintf(stdout, "This path's contribution to IS sum (in bits):    %.2f\n",  pr_unsorted[s]);
         fprintf(stdout, "Log posterior probability of path under HMM:     %.2f\n",  hmmsc_ld - fsc);

         ParsetreeDump(stdout, pstr[t], cm, sq->dsq);
         h4_path_DumpAnnotated(stdout, out_pi[0], hmm, mo, sq->dsq);
         /* note that we've seen a jump */
         njump++;
      }

      ldprev = ld;
   }


   /* clean up and return */
   if (njump > 0) {
      h4_path_Destroy(out_pi[0]);
      esl_msa_Destroy(out_msa);
   }

   esl_sq_Destroy(out_sq[0]);
   free(out_pi);
   free(out_sq);
   free(pr_sorted);

   ERROR:
      return status;


   return eslOK;
}

/* Function: map_ss_cons()
 *
 * Purpose:  Copy the secondary structure annotation from <cm>
 *           onto the ss_cons line of <msa>. Useful because
 *           msa often has insert columns => mapping non-trivial.
 *
 *
 *
 * args:    cm            - covariance model
 *          msa           - multiple sequence alignment
 *          errbuf        - optRETURN: space for an informative
 *                          error message if something fails.
 *                          Feature more for the future tbh.
 *
 * Returns: eslOK on success
 *
 * */

int map_ss_cons(CM_t *cm, ESL_MSA *msa, char *errbuf) {

   int cpos   = 0;    /* cm node index    */
   int k;             /* msa column index */
   int status;        /* esl return code  */

   ESL_ALLOC(msa->ss_cons, (sizeof(char) * (msa->alen+1)));

   /* loop over columns and add ss_cons annotation */
   for (k = 0; k <= msa->alen; k++) {
      /* if the RF line is an 'x' here, we have a match column! */
      /* 120 = 'x' in ASCII. If you don't know, now you know.   */
      if (msa->rf[k] == 120) {
         msa->ss_cons[k] = cm->cmcons->cstr[cpos];
         cpos++;
      }
      else {
         msa->ss_cons[k] = '.';
      }
   }

   /* terminate ss_cons w/ null byte */
   msa->ss_cons[msa->alen] = '\0';

   return eslOK;

   ERROR:
      return status;

}


char *strsafe(char *dest, const char *src)
{

   int i;
   for (i=0; src[i] != '\0'; i++) {
      if (src[i] == '/') dest[i] = '_';
      else               dest[i] = src[i];
   }

   dest[i] = '\0';

   return dest;

}
```
