## Supplemental Code S1 for "Remote homology search with hidden Potts models": h4_path_cmis.c

```
/* h4_path_cmis.c */
/* Grey's functions with h4 paths */

#include <stdlib.h>
#include <stdio.h>
#include <stdarg.h>

#include "easel.h"
#include "esl_alphabet.h"
#include "esl_msa.h"

#include "h4_config.h"
#include "h4_mode.h"
#include "h4_profile.h"
#include "h4_path.h"


int h4_path_FromMSA(ESL_MSA *msa, int8_t *matassign, int optflags, H4_PATH **pi){
   int  idx;                      /* counter over seqs in MSA */
   int  lpos, rpos;               /* leftmost, rightmost consensus column, 1..alen */
   int  status          = eslOK;

   /* make sure all traces are null before we procede */
   for (idx = 0; idx < msa->nseq; idx++) pi[idx] = NULL;

   /* Find leftmost and rightmost consensus columns */
   for (lpos = 1;         lpos <= msa->alen; lpos ++)  if (matassign[lpos]) break;
   for (rpos = msa->alen; rpos >= 1;         rpos--)   if (matassign[rpos]) break;

   fprintf(stdout, "lpos: %d, rpos: %d\n", lpos, rpos);

   for (idx = 0; idx < msa->nseq; idx++) {

      fprintf(stdout, "\tidx: %d\n", idx);

      /* allocate space for path */
      if ((pi[idx] = h4_path_Create()) == NULL) goto ERROR;

      /* infer path from correspoding row in **digital** msa */
      h4_path_InferGlocal(msa->abc, msa->ax[idx], msa->alen, matassign, lpos, rpos, pi[idx]);

      //h4_path_Dump(stdout, pi[idx]);
   }


   return eslOK;

   ERROR:
      return status;
}

int h4_path_DumpAnnotated(FILE *fp, const H4_PATH *pi, const H4_PROFILE *hmm, const H4_MODE *mo, const ESL_DSQ *dsq) {

   float sc = (pi->Z ? 0. : -eslINFINITY);      /* final score for path; -inf if trace is empty      */
   int   i  = 0;                                /* seq position index                                */
   int   k  = 0;                                /* hmm node index                                    */
   char  resi;                                  /* emitted residue character                         */
   int   z,y;                                   /* counters over path elements and their runlengths  */
   float tsc;                                   /* single transition score                           */
   float esc;                                   /* single emission score                             */

   /* print header */
   fprintf(fp, "#  z      y  st   k     i   x_i  transit  emission \n");
   fprintf(fp, "#----- ----- -- ----- ----- ---  -------- -------- \n");

   /* loop over state runs in trace */
   for (z = 0; z < pi->Z; z++) {


      /* handle each state differently */
      switch (pi->st[z]) {

         /* N-terminus (5') flanking insert states */
         case h4P_N:

            /* loop over all runs in state */
            for (y = 0; y < pi->rle[z]; y++)
            {

               /* handle last instance of N state: we don't assign a residue here */
               if (y == pi->rle[z] - 1) {
                  tsc = mo->xsc[h4_N][h4_MOVE];  /* N -> N */
                  resi = ' ';
                  /* silent states are assigned sequence position 0 */
                  /* don't worry, we'll fix it after the for loop */
                  i = 0;
               }
               /* handle all other N states; we say they emit */
               else {
                  i++;
                  tsc = mo->xsc[h4_N][h4_LOOP];  /* N -> G */
                  resi = hmm->abc->sym[dsq[i]];
               }

               /* print info to outfile */
               fprintf(fp, "%5d %5d %2s %5d %5d  %c  %8.4f %8.4f\n",
                       z,
                       y,
                       h4_path_DecodeStatetype(pi->st[z]),
                       k,
                       i,
                       resi,
                       tsc,
                       0.0);
               sc += tsc;
            }

            /* set i to be number of residues emitted by N state */
            i = pi->rle[z]-1;
            break;

         /* G global entry state (not the airport kind) */
         case h4P_G:
            /* figure out if next state is a match or delete */
            switch(pi->st[z+1]) {
               case h4P_MG: tsc = hmm->tsc[0][h4_MM]; break;
               case h4P_DG: tsc = hmm->tsc[0][h4_MD]; break;
            }

            /* print info to outfile */
            fprintf(fp, "%5d %5d %2s %5d %5d  %c  %8.4f %8.4f\n",
                    z,
                    0,
                    h4_path_DecodeStatetype(pi->st[z]),
                    k,
                    0,
                    ' ',
                    tsc,
                    0.0);

            /* update total score */
            sc += tsc;

            break;

         /* global match state */
         case h4P_MG:

            /* loop over states in this run */
            for (y = 0; y < pi->rle[z]; y++){
               /* increment node and sequence index */
               k++;
               i++;

               /* get transition score */
               /* look to next run if we're at the end of the run*/
               if (y == pi->rle[z] -1) {
                  switch(pi->st[z+1]) {
                     case h4P_MG: tsc = hmm->tsc[k][h4_MM]; break;
                     case h4P_DG: tsc = hmm->tsc[k][h4_MD]; break;
                     case h4P_IG: tsc = hmm->tsc[k][h4_MI]; break;
                     case h4P_C:  tsc = hmm->tsc[k][h4_MM]; break;

                  }
               }
               /* all intra-run transitions in run are M->M */
               else tsc = hmm->tsc[k][h4_MM];

               /* get emission score */
               esc = hmm->rsc[dsq[i]][k];

               /* print info to outfile */
               fprintf(fp, "%5d %5d %2s %5d %5d  %c  %8.4f %8.4f\n",
                       z,
                       y,
                       h4_path_DecodeStatetype(pi->st[z]),
                       k,
                       i,
                       hmm->abc->sym[dsq[i]],
                       tsc,
                       esc);

               /* update total score */
               sc += tsc + esc;


            }

            break;

         /* Global delete state */
         case h4P_DG:

            /* loop over all states in the run */
            for (y = 0; y < pi->rle[z]; y++){
               /* increment node index ONLY */
               k++;

               /* get transition scores */
               /* look a head if we're at the end of the run */
               if (y == pi->rle[z] -1) {
                   switch(pi->st[z+1]) {
                     case h4P_MG: tsc = hmm->tsc[k][h4_DM]; break;
                     case h4P_DG: tsc = hmm->tsc[k][h4_DD]; break;
                     case h4P_IG: tsc = hmm->tsc[k][h4_DI]; break;
                     case h4P_C:  tsc = hmm->tsc[k][h4_DM]; break;
                   }
               }

               /* all intra-run transitions are D->D */
               else tsc = hmm->tsc[k][h4_DD];

               /* print info to outfile */
               fprintf(fp, "%5d %5d %2s %5d %5d  %c  %8.4f %8.4f\n",
                       z,
                       y,
                       h4_path_DecodeStatetype(pi->st[z]),
                       k,
                       0,
                       '-',
                       tsc,
                       0.0);

               sc += tsc;
            }

            break;

         /* global insert states */
         case h4P_IG:
            /* loop over all states in the run */
            for (y = 0; y < pi->rle[z]; y++){

               /* increment the sequence position index ONLY */
               i++;

               /* get transition score */
               /* if we're at the end of the run, look ahead */
               if (y == pi->rle[z] -1) {
                   switch(pi->st[z+1]) {
                     case h4P_MG: tsc = hmm->tsc[k][h4_IM]; break;
                     case h4P_DG: tsc = hmm->tsc[k][h4_ID]; break;
                     case h4P_IG: tsc = hmm->tsc[k][h4_II]; break;
                   }
               }
               /* all intra-run transitions are I->I */
               else tsc = hmm->tsc[k][h4_II];

               /* print info to outfile */
               fprintf(fp, "%5d %5d %2s %5d %5d  %c  %8.4f %8.4f\n",
                       z,
                       y,
                       h4_path_DecodeStatetype(pi->st[z]),
                       k,
                       i,
                       hmm->abc->sym[dsq[i]],
                       tsc,
                       0.0);

               /* update total score */
               sc += tsc;
            }

            break;

         /* C-terminus (3') flanking insert states */
         case h4P_C:

            /* loop over all states in the run */
            for (y = 0; y < pi->rle[z]; y++)
            {

               /* get transition score */
               /* handle last instance of C state */
               if (y == pi->rle[z] - 1) {
                  tsc = mo->xsc[h4_C][h4_MOVE];   /* C->T */
                  resi = ' ';
                  i = 0;                          /* assign i = 0 to silent states */
               }
               /* all but last C state emit residues, transition C->C */
               else {
                  /* increment the sequence index ONLY */
                  i++;
                  tsc = mo->xsc[h4_C][h4_LOOP];   /* C->C */
                  resi = hmm->abc->sym[dsq[i]];
               }


               /* print info to outfile */
               fprintf(fp, "%5d %5d %2s %5d %5d  %c  %8.4f %8.4f\n",
                       z,
                       y,
                       h4_path_DecodeStatetype(pi->st[z]),
                       k,
                       i,
                       resi,
                       tsc,
                       0.0);
               /* update total score */
               sc += tsc;
            }

            break;

         /* for non-uniglocal states, tell the caller */
         default: return eslEINVAL;

      }
   }

   /* print footer */
   fprintf(fp, "#----- ----- -- ----- ----- ---  -------- -------- \n");
   fprintf(fp, "# path score: %4f\n", sc);

   /* clean up and return */
   return eslOK;
}
```
