## Supplemental Code S1 for "Remote homology search with hidden Potts models": h4_path_cmis.h

```
/* h4_path_cmis.h*/
/* Grey's functions with h4 paths */


#ifndef h4PATHCMIS_INCLUDED
#define h4PATH_INCLUDED


#include <stdlib.h>
#include <stdio.h>
#include <stdarg.h>

#include "easel.h"
#include "esl_alphabet.h"
#include "esl_msa.h"

#include "h4_config.h"
#include "h4_mode.h"
#include "h4_profile.h"
#include "h4_path.h"


extern int h4_path_FromMSA(ESL_MSA *msa, int8_t *matassign, int optflags, H4_PATH **pi);
extern int h4_path_DumpAnnotated(FILE *fp, const H4_PATH *pi, const H4_PROFILE *hmm, const H4_MODE *mo, const ESL_DSQ *dsq);
#endif
```
