## Supplemental Code S1 for "Remote homology search with hidden Potts models": h4_pathalign.c

```
#include "easel.h"
#include "esl_alphabet.h"
#include "esl_msa.h"
#include "esl_sq.h"
#include "esl_vectorops.h"

#include "h4_path.h"
#include "h4_profile.h"

/* declaration of internal functions */
static int map_new_msa(H4_PATH **pi, int nseq, int M, int allcons, int **ret_inscount, int **ret_matuse, int **ret_matmap, int *ret_alen);
static int make_digital_msa(ESL_SQ **sq, const ESL_MSA *premsa, H4_PATH **pi, int nseq, const int *matuse, const int *matmap, int M, int alen, int allcons, ESL_MSA **ret_msa);
static int annotate_rf(ESL_MSA *msa, int M, const int *matuse, const int *matmap);
static int rejustify_insertions_digital(ESL_MSA *msa, const int *inserts, const int *matmap, const int *matuse, int M);

int
h4_pathalign_Seqs(ESL_SQ **sq, H4_PATH **pi, int nseq, int M, int allcons, H4_PROFILE *hmm, ESL_MSA **ret_msa)
{
   ESL_MSA            *msa        = NULL;       /* new msa to return                                                   */
   int                *inscount   = NULL;       /* array of max gaps between aligned columns                           */
   int                *matmap     = NULL;       /* matmap[k] = apos of match k matmap[1..M] = [1..alen]                */
   int                *matuse     = NULL;       /* TRUE if an alignment column is associated with match state k [1..M] */
   int                 idx;                     /* sequence index                                                      */
   int                 alen;                    /* width of alignment, aka # of columns                                */
   //int                 apos;                    /* width of alignment, aka # of columns                                */
   int                 status;                  /* Easel return code                                                   */

   /* map match positions to MSA columns */
   if ((status = map_new_msa(pi, nseq, M, allcons, &inscount, &matuse, &matmap, &alen))        != eslOK) return status;
   /* make a new ESL_MSA */
   if ((status = make_digital_msa(sq, NULL, pi, nseq, matuse, matmap, M, alen, allcons, &msa)) != eslOK) goto ERROR;

   /* add RF annotation */
   if ((status = annotate_rf(msa, M, matuse, matmap))                                          != eslOK) goto ERROR;
   /* rejustify insertions */
   if ((status = rejustify_insertions_digital(msa, inscount, matmap, matuse, M))               != eslOK) goto ERROR;


   /* set sequence names, weights, etc */
   for (idx = 0; idx < nseq; idx++)
   {
      esl_msa_SetSeqName(msa, idx, sq[idx]->name, -1);
      if (sq[idx]->acc[0]  != '\0') esl_msa_SetSeqAccession  (msa, idx, sq[idx]->acc,  -1);
      if (sq[idx]->desc[0] != '\0') esl_msa_SetSeqDescription(msa, idx, sq[idx]->desc, -1);
      msa->wgt[idx] = 1.0;
      if (msa->sqlen != NULL) msa->sqlen[idx] = sq[idx]->n;

      //for(apos = 0; apos < alen; apos++){
      //   fprintf(stdout, "\tidx: %d, apos; %d, residue: %d\n", idx, apos, msa->ax[idx][apos]);
      //}
   }

   /* clean up and return */
   *ret_msa = msa;
   free(inscount);
   free(matmap);
   free(matuse);
   return eslOK;

   ERROR:
      if (msa      != NULL) esl_msa_Destroy(msa);
      if (inscount != NULL) free(inscount);
      if (matmap   != NULL) free(matmap);
      if (matuse   != NULL) free(matuse);
      *ret_msa = NULL;
      return status;
}

static int
map_new_msa(H4_PATH **pi, int nseq, int M, int allcons, int **ret_inscount,
            int **ret_matuse, int **ret_matmap, int *ret_alen)
{
   int *inscount = NULL;  /* inscount[k=0..M] == max # of inserts in node k                            */
   int *insnum   = NULL;  /* insct[k=0..M] == # of inserts in node k in current trace                  */
   int *matuse   = NULL;  /* matuse[k=1..M] == TRUE|FALSE: does node k map to an alignment column      */
   int *matmap   = NULL;  /* matmap[k=1..M]: if matuse[k] TRUE, what column 1..alen does node k map to */
   int  idx;              /* counter over sequences                                                    */
   int  z;                /* index into path positions                                                 */
   int  y;                /* index over path position run lengths                                      */
   int  alen;             /* length of alignment                                                       */
   int  k;                /* counter over nodes 1..M                                                   */
   int  status;

   /* allocate space for arrays */
   ESL_ALLOC(inscount, sizeof(int) * (M+1));
   ESL_ALLOC(insnum,   sizeof(int) * (M+1));
   ESL_ALLOC(matuse,   sizeof(int) * (M+1)); matuse[0] = 0;
   ESL_ALLOC(matmap,   sizeof(int) * (M+1)); matmap[0] = 0;

   /* initialize insert count array */
   esl_vec_ISet(inscount, M+1, 0);

   /* if we are commanded to keep all match coumns, set this array to true  (except node 0) */
   if    (allcons) esl_vec_ISet(matuse+1, M, TRUE);
   else  esl_vec_ISet(matuse+1, M, FALSE);

   /* Collect inscount[], matuse[] */
   for (idx = 0; idx < nseq; idx++) {
      /* reset single-trace insert counts */
      esl_vec_ISet(insnum, M+1, 0);

      //h4_path_Dump(stdout, pi[idx]);

      for (z = 0; z < pi[idx]->Z; z++) {            // z=0 is always an N.
         //fprintf(stdout, "z: %d\n", z);

         switch(pi[idx]->st[z]) {

            /* N-terminus (5') flanking insert states */
            case h4P_N:

               //fprintf(stdout, "number of residues in N state: %d\n", pi[idx]->rle[z]-1);

               /* record how many residues this path has in its N state */
               /* N state emits on loop transition */
               insnum[0] = pi[idx]->rle[z]-1;
               k = 0;
               break;

            /* global match states */
            case h4P_MG:
               for (y = 0; y < pi[idx]->rle[z]; y++){

                  /* increment the node counter */
                  k++;

                  /* note that this match state is occupied */
                  matuse[k] = TRUE;

               }
               break;

            /* global delete states */
            case h4P_DG:
               /* increase the node counter by the number of delete states*/
               k += pi[idx]->rle[z];
               break;

            /*global insert states */
            case h4P_IG:
               //fprintf(stdout, "number of residues in insert state %d: %d\n", k, pi[idx]->rle[z]);

               /* record how many residues this path has in insert state k */
               insnum[k] = pi[idx]->rle[z];
               break;

            /* C-terminus (3') flanking insert states */
            case h4P_C:
           /* global delete states */
               //fprintf(stdout, "number of residues in C state: %d\n", pi[idx]->rle[z]-1);

               /* record how many residues this path has in its N state */
               /* C state emits on loop transition */
               insnum[M] = pi[idx]->rle[z]-1;

            /* silent uniglogal states that don't show up in the MSA */
            case h4P_S:
            case h4P_B:
            case h4P_G:
            case h4P_E:
            case h4P_T:
            case h4P_NST:
               break;

            /* states we don't deal with in uniglocal mode */
            case h4P_L:
            case h4P_ML:
            case h4P_IL:
            case h4P_DL:
            case h4P_J:
               break;


         }

      }
      /* update inscount if this path  widens the alignment */
      for (k = 0; k <= M; k++) {
         inscount[k] = ESL_MAX(inscount[k], insnum[k]);
      }
   }

   /* Use inscount, matuse to set the matmap[] */
   alen      = inscount[0];
   for (k = 1; k <= M; k++) {
      //fprintf(stdout, "node %d\n", k);
      //fprintf(stdout, "\tmatuse: %d\n", matuse[k]);
      //fprintf(stdout, "\tinscount: %d\n", inscount[k] );

      if (matuse[k]) { matmap[k] = alen+1; alen += 1+inscount[k]; }
      else           { matmap[k] = alen;   alen +=   inscount[k]; }
      //fprintf(stdout, "\tmatmap[k]: %d\n", matmap[k]);
   }


   /* clean up and return */
   free(insnum);
   *ret_inscount = inscount;
   *ret_matuse   = matuse;
   *ret_matmap   = matmap;
   *ret_alen     = alen;
   return eslOK;

   ERROR:
      if (inscount) free(inscount);
      if (insnum)   free(insnum);
      if (matuse)   free(matuse);
      if (matmap)   free(matmap);
      *ret_inscount = NULL;
      *ret_matuse   = NULL;
      *ret_matmap   = NULL;
      *ret_alen     = 0;
      return status;
}

static int
make_digital_msa(ESL_SQ **sq, const ESL_MSA *premsa, H4_PATH **pi, int nseq, const int *matuse, const int *matmap, int M, int alen, int allcons, ESL_MSA **ret_msa) {
   const ESL_ALPHABET *abc;         /* esl alphabet              */
   ESL_MSA            *msa = NULL;  /* msa we are buiilding      */
   int                 z,y;         /* index over path positions */
   int                 i;           /* sequence position index   */
   int                 k;           /* match state/node index    */
   int                 idx;         /* index over sequences      */
   int                 apos;        /* index over MSA columns    */
   int                 status;      /* esl return code           */

   /* make sure we have an actual sequence array */
   if (sq == NULL) return eslEINVAL;

   /* set alphabet */
   abc =  sq[0]->abc;

   /* allocate memory for msa */
   if ((msa = esl_msa_CreateDigital(abc, nseq, alen)) == NULL) { status = eslEMEM; goto ERROR;  }

   for (idx = 0; idx < nseq; idx++){
      /* initialize MSA row to all gaps, flanked by eslDSQ_SENTINEL flag */
      msa->ax[idx][0]      = eslDSQ_SENTINEL;
      for (apos = 1; apos <= alen; apos++) msa->ax[idx][apos] = esl_abc_XGetGap(abc);
      msa->ax[idx][alen+1] = eslDSQ_SENTINEL;

      /* now fill in this MSA row by traversing the trace */
      apos = 1;
      i    = 0;
      k    = 0;

      //h4_path_Dump(stdout, pi[idx]);

      for (z = 0; z < pi[idx]->Z; z++) {
         switch(pi[idx]->st[z]) {

            /* N-terminus (5') or C-terminus (3') flanking insert states*/
            case h4P_N:
            case h4P_C:
               for (y = 1; y < pi[idx]->rle[z]; y++) {

                  /* increment the sequence position index ONLY */
                  i++;

                  /* set MSA element to appropriate character */
                  msa->ax[idx][apos] = sq[idx]->dsq[i];
                  //fprintf(stdout, "\tFlanking state! i = %d, a = %d, k = %d, residue = %d\n", i, apos, k, msa->ax[idx][apos]);
                  /* increment the sequence postion and MSA column indices */
                  apos++;
               }
               break;

            /* global match states */
            case h4P_MG:
               /* loop over all match states in ths run */
               for (y = 0; y < pi[idx]->rle[z]; y++){
                  /* increment both the node and sequence position indices */
                  k++;
                  i++;

                  /* note that we are using matmap to get apos here */
                  /* Allows us to skip insert columns for which this sequence has no residue */
                  apos = matmap[k];

                  /* set MSA element to appropriate character */
                  msa->ax[idx][apos] = sq[idx]->dsq[i];

                  //fprintf(stdout, "\tMatch state! i = %d, a = %d, k = %d, residue = %d\n", i, matmap[k], k, msa->ax[idx][apos]);

                  /* set apos to be the next MSA column */
                  apos = matmap[k] + 1;

               }
               break;

            /* global delete states */
            case h4P_DG:
               /* loop over all delete states in this run */
               for (y = 0; y < pi[idx]->rle[z]; y++){
                  /* increment the node index ONLY */
                  k++;

                  /* note that we are using matmap to get apos here */
                  apos = matmap[k];

                  /* set MSA element to appropriate gap char */
                  msa->ax[idx][apos] = esl_abc_XGetGap(abc);

                  //fprintf(stdout, "\tDelete state! i = %d, a = %d, k = %d, residue = %d\n", i, apos, k, msa->ax[idx][apos]);

                  /* set apos to be the next MSA column */
                  apos = matmap[k] + 1;

               }
               break;

            case h4P_IG:
               /* loop over all insert states in this run */
               for (y = 0; y < pi[idx]->rle[z]; y++){

                  /* increment the sequence position index ONLY */
                  i++;

                  /* set MSA element to appropriate character */
                  msa->ax[idx][apos] = sq[idx]->dsq[i];
                  //fprintf(stdout, "\tInsert state! i = %d, a = %d, k = %d, residue = %d\n", i, apos, k, msa->ax[idx][apos]);

                  /* move over one MSA column */
                  apos++;
               }
               break;

            default:
               break;
         }
      }

      //for (apos = 0; apos < alen; apos++) {
      //   fprintf(stdout, "\tidx: %d, apos; %d, residue: %d\n", idx, apos, msa->ax[idx][apos]);
      //}
   }

   /* set MSA attributes and return */
   msa->nseq = nseq;
   msa->alen = alen;
   *ret_msa = msa;
   return eslOK;

   ERROR:
      if (msa) esl_msa_Destroy(msa);
      *ret_msa = NULL;
      return status;


}

static int
annotate_rf(ESL_MSA *msa, int M, const int *matuse, const int *matmap)
{
   int apos;   /* alignment column index */
   int k;      /* match state index      */
   int status; /* esl return code        */

   /* allocate memory for RF line */
   ESL_ALLOC(msa->rf, sizeof(char) * (msa->alen+1));

   /* start by making every position a '.' */
   for (apos = 0; apos < msa->alen; apos++) {
      msa->rf[apos] = '.';
      msa->rf[msa->alen] = '\0';
   }

   /* mark match columns with an 'x' */
   for (k = 1; k <= M; k++)
      if (matuse[k]) msa->rf[matmap[k]-1] = 'x'; /* watch off by one: rf[0..alen-1]; matmap[] = 1..alen */

   return eslOK;

   ERROR:
      return status;

}

static int
rejustify_insertions_digital(ESL_MSA *msa, const int *inserts, const int *matmap, const int *matuse, int M)
{
   int idx;         /* sequence index */
   int k;           /* match state index */
   int apos;        /* MSA column index  */
   int nins;        /* index over residues in this insert state */
   int npos, opos;

   for (idx = 0; idx < msa->nseq; idx++)
   {
      for (k = 0; k < M; k++)
      {
         if (inserts[k] > 1)
         {
             for (nins = 0, apos = matmap[k]+1; apos <= matmap[k+1]-matuse[k+1]; apos++)
             {
               if (esl_abc_XIsResidue(msa->abc, msa->ax[idx][apos])) nins++;
             }

             if (k == 0) nins = 0;    /* N-terminus is right justified */
             else        nins /= 2;   /* split in half; nins now = # of residues left left-justified  */

             opos = npos = matmap[k+1]-matuse[k+1];
             while (opos >= matmap[k]+1+nins) {
               if (esl_abc_XIsGap(msa->abc, msa->ax[idx][opos])) opos--;
                else {
                   msa->ax[idx][npos] = msa->ax[idx][opos];
                   npos--;
                   opos--;
                }
             }

             while (npos >= matmap[k]+1+nins) {
                msa->ax[idx][npos] = esl_abc_XGetGap(msa->abc);
                if (msa->pp != NULL && msa->pp[idx] != NULL) msa->pp[idx][npos-1] = '.';
                npos--;
             }
         }
      }
   }


   return eslOK;
}
```
