## Supplemental Code S1 for "Remote homology search with hidden Potts models": h4_pathalign.h

```
#ifndef h4PATHALIGN_INCLUDED
#define h4PATHALIGN_INCLUDED

#include "easel.h"
#include "esl_alphabet.h"
#include "esl_msa.h"
#include "esl_sq.h"
#include "esl_vectorops.h"

#include "h4_path.h"
#include "h4_profile.h"

extern int  h4_pathalign_Seqs(ESL_SQ **sq, H4_PATH **pi, int nseq, int M, int allcons, H4_PROFILE *hmm, ESL_MSA **ret_msa);

#endif /* h4PATHALIGN_INCLUDED */
```
