## Supplemental Code S1 for "Remote homology search with hidden Potts models": h4score_path.c

```
#include <stdio.h>
#include <stdlib.h>
#include <string.h>

#include "easel.h"
#include "esl_alphabet.h"
#include "esl_getopts.h"
#include "esl_msa.h"
#include "esl_msafile.h"
#include "esl_sq.h"

#include "h4_config.h"
#include "h4_hmmfile.h"
#include "h4_mode.h"
#include "h4_profile.h"
#include "h4_path.h"
#include "h4_refmx.h"
#include "logsum.h"
#include "reference_dp.h"

#include "h4_path_cmis.h"

static ESL_OPTIONS options[] = {
   /* name        type        default    env range togs  reqs  incomp            help                                            docgroup */
   { "-h",        eslARG_NONE,    FALSE, NULL, NULL, NULL, NULL, NULL,            "help; show brief info on version and usage",  1 },

   /* Options forcing which alphabet we're working in (normally autodetected) */
   { "--amino",   eslARG_NONE,    FALSE, NULL, NULL, NULL, NULL, "--dna,--rna",   "We are dealing with protein sequences",       2 },
   { "--dna",     eslARG_NONE,    FALSE, NULL, NULL, NULL, NULL, "--amino,--rna", "We are dealing with dna sequences",           2 },
   { "--rna",     eslARG_NONE,    FALSE, NULL, NULL, NULL, NULL, "--amino,--dna", "We are dealing with rna sequences",           2 },

   { 0,0,0,0,0,0,0,0,0,0 },
};

static char banner[] = "Score sequence(s) and path(s) with an HMM";
static char usage[]  = "[-options] <hmmfile> <msafile>";

static void
cmdline_failure(char *argv0, char *format, ...)
{
   va_list argp;
   printf("\nERROR: ");
   va_start(argp, format);
   vfprintf(stderr, format, argp);
   va_end(argp);
   esl_usage(stdout, argv0, usage);
   printf("\nTo see more help on available options, do %s -h\n\n", argv0);
   exit(1);

}

int main(int argc, char *argv[]){

   ESL_GETOPTS      *go        = NULL;
   ESL_ALPHABET     *abc       = NULL;
   char             *hmmfile   = NULL;                       /* input hmm filepath                       */
   char             *msafile   = NULL;                       /* input msa filepath                       */
   H4_HMMFILE       *hmmfp     = NULL;                       /* open input hmm file stream               */
   ESL_MSAFILE      *afp       = NULL;                       /* oopen input msa file stream              */
   H4_PROFILE       *hmm       = NULL;                       /* input hmm                                */
   H4_MODE          *mo        = h4_mode_Create();           /* h4 profile hmm mode                      */
   ESL_MSA          *msa       = NULL;                       /* input msa                                */
   ESL_SQ          **sq        = NULL;                       /* array of sequences                       */
   H4_REFMX         *fwd        = h4_refmx_Create(100, 100); /* forward DP matrix                        */
   H4_PATH         **pi         = NULL;                      /* dummy path array for sampled alignments  */
   int8_t            *matassign = NULL;                       /* MAT state assignments if 1; 1..alen      */
   float             hmmsc;                                  /* Joint hmm score of seq and path, in nats */
   float             fsc;                                    /* forward score of a sequence, in nats     */
   int               i;                                      /* msa position (column) index              */
   int               n;                                      /* msa sequence (row) index                 */
   int               fmt       = eslMSAFILE_STOCKHOLM;       /* input msa format #sorrynotsorry          */
   int               status;                                 /* easel return code                        */

   /* parse command line */
   go = esl_getopts_Create(options);
   if (esl_opt_ProcessCmdline(go, argc, argv) != eslOK) cmdline_failure(argv[0], "Failed to parse command line: %s\n", go->errbuf);
   if (esl_opt_VerifyConfig(go)               != eslOK) cmdline_failure(argv[0], "Error in app configuration:   %s\n", go->errbuf);

   /* output help if requested */
   if (esl_opt_GetBoolean(go, "-h") ) {
      esl_banner(stdout, argv[0], banner);
      esl_usage (stdout, argv[0], usage);
      puts("\n where options are:");
      esl_opt_DisplayHelp(stdout, go, 1, 2, 80);
      puts("\n Alphabet options:");
      esl_opt_DisplayHelp(stdout, go, 2, 2, 80);
      exit(0);
   }

   if       (esl_opt_GetBoolean(go, "--amino")) abc = esl_alphabet_Create(eslAMINO);
   else if  (esl_opt_GetBoolean(go, "--dna"))   abc = esl_alphabet_Create(eslDNA);
   else if  (esl_opt_GetBoolean(go, "--rna"))   abc = esl_alphabet_Create(eslRNA);


   /* read arguments */
   if (esl_opt_ArgNumber(go) != 2)  cmdline_failure(argv[0], "Incorrect number of command line arguments.\n", go->errbuf);

   hmmfile = esl_opt_GetArg(go, 1);
   msafile = esl_opt_GetArg(go, 2);

   /* read the hmm file */
   if ( h4_hmmfile_Open(hmmfile, NULL, &hmmfp) != eslOK) esl_fatal("couldn't open profile file %s", hmmfile);
   /* read first hmm  from hmm file */
   if ( h4_hmmfile_Read(hmmfp, &abc, &hmm)     != eslOK) esl_fatal("failed to read profile from file %s", hmmfile);
   h4_hmmfile_Close(hmmfp);

   /* set HMM mode to uniglocal  */
   h4_mode_SetUniglocal(mo);

   /* allow '-' and '.' as gap */
   esl_alphabet_SetEquiv(abc, '.', '-');

   /* read the msa file */
   if ((status = esl_msafile_Open(&abc, msafile, NULL, fmt, NULL, &afp)) != eslOK) esl_msafile_OpenFailure(afp, status);
   if ((status = esl_msafile_Read(afp, &msa))                            != eslOK) esl_fatal ("Failed to read MSA");
   esl_msafile_Close(afp);

   /* extract sequences from MSA */
   ESL_ALLOC(sq, sizeof(ESL_SQ *) * msa->nseq);
   for (n = 0; n < msa->nseq; n++) {
      fprintf(stdout, "n: %d\n", n);
      esl_sq_FetchFromMSA(msa, n, &sq[n]);
   }

   if (! (msa->flags & eslMSA_DIGITAL)) esl_fatal("need a digital msa");
   if (msa->rf == NULL)                 esl_fatal("msa lacks an RF line");

   /* extract match states from alignment */
   ESL_ALLOC(matassign, sizeof(int8_t) * (msa->alen + 1));
   for (i=0; i < msa->alen; i++){
      if (msa->rf[i] != '.') matassign[i+1] = 1;
      else                   matassign[i+1] = 0;
   }

   /* generate trace array from msa */
   ESL_ALLOC(pi, sizeof(H4_PATH *) * msa->nseq);
   h4_path_FromMSA(msa, matassign, 0, pi);

   /* run h4 logsum magic */
   h4_logsum_Init();

   /* score sequences and paths */
   for (n=0;  n < msa->nseq; n++) {

      /* configure profile and background models using seq length */
      h4_mode_SetLength(mo, sq[n]->n);

      /* rescale the forward matrix */
      h4_refmx_GrowTo(fwd, hmm->M, sq[n]->n);

      /* run forward algorithm */
      h4_reference_Forward(sq[n]->dsq, sq[n]->n, hmm, mo, fwd, &fsc);

      /* score sequence and given path */
      h4_path_Score(pi[n], sq[n]->dsq, hmm, mo, &hmmsc);

      h4_path_DumpAnnotated(stdout, pi[n], hmm, mo, sq[n]->dsq);

      fprintf(stdout, "seq: %s\n", sq[n]->name);
      fprintf(stdout, "Raw forward score: %.2f\n", fsc);
      fprintf(stdout, "Forward score of sequence: %.2f\n", fsc - mo->nullsc );
      fprintf(stdout, "HMM score of sequence, path: %.2f\n", hmmsc);
      fprintf(stdout, "Log posterior probability of path under HMM:  %.2f\n", hmmsc - fsc );
   }

   fprintf(stdout, "hello world\n");

   /* clean up and return */
   free(matassign);
   for (n = 0; n < msa->nseq; n++) {
      esl_sq_Destroy(sq[n]);
      h4_path_Destroy(pi[n]);
   }
   free(sq);
   free(pi);
   esl_msa_Destroy(msa);
   h4_profile_Destroy(hmm);
   h4_mode_Destroy(mo);
   h4_refmx_Destroy(fwd);
   esl_alphabet_Destroy(abc);
   esl_getopts_Destroy(go);
   return 0;

   ERROR:
      return status;


}
```
