## Supplemental Code S1 for "Remote homology search with hidden Potts models": hmmscore_path.c

```
#include <stdio.h>
#include <stdlib.h>
#include <string.h>

#include "easel.h"
#include "esl_alphabet.h"
#include "esl_getopts.h"
#include "esl_msa.h"
#include "esl_msafile.h"
#include "esl_sq.h"

#include "p7_config.h"
#include "hmmer.h"


static ESL_OPTIONS options[] = {
   /* name        type        default    env range togs  reqs  incomp            help                                            docgroup */
   { "-h",        eslARG_NONE,    FALSE, NULL, NULL, NULL, NULL, NULL,            "help; show brief info on version and usage",  1 },

   /* Options forcing which alphabet we're working in (normally autodetected) */
   { "--amino",   eslARG_NONE,    FALSE, NULL, NULL, NULL, NULL, "--dna,--rna",   "We are dealing with protein sequences",       2 },
   { "--dna",     eslARG_NONE,    FALSE, NULL, NULL, NULL, NULL, "--amino,--rna", "We are dealing with dna sequences",           2 },
   { "--rna",     eslARG_NONE,    FALSE, NULL, NULL, NULL, NULL, "--amino,--dna", "We are dealing with rna sequences",           2 },

   { 0,0,0,0,0,0,0,0,0,0 },
};

static char banner[] = "Score sequence(s) and path(s) with an HMM";
static char usage[]  = "[-options] <hmmfile> <msafile>";

static void
cmdline_failure(char *argv0, char *format, ...)
{
   va_list argp;
   printf("\nERROR: ");
   va_start(argp, format);
   vfprintf(stderr, format, argp);
   va_end(argp);
   esl_usage(stdout, argv0, usage);
   printf("\nTo see more help on available options, do %s -h\n\n", argv0);
   exit(1);

}

int main(int argc, char *argv[]){

   ESL_GETOPTS      *go        = NULL;
   ESL_ALPHABET     *abc       = NULL;
   char             *hmmfile   = NULL;                 /* input hmm filepath                       */
   char             *msafile   = NULL;                 /* input msa filepath                       */
   P7_HMMFILE       *hmmfp     = NULL;                 /* open input hmm file stream               */
   ESL_MSAFILE      *afp       = NULL;
   P7_HMM           *hmm       = NULL;                 /* hmm                                      */
   ESL_MSA          *msa       = NULL;
   P7_PROFILE       *gm        = NULL;                 /* h3 profile model                            */
   P7_BG            *bg        = NULL;                 /* h3 background model                         */
   ESL_SQ          **sq        = NULL;                 /* array of sequences                       */
   P7_TRACE        **tr        = NULL;                 /* trace for alignment paths                */
   P7_GMX           *fwd       = p7_gmx_Create(100, 100);   /* h3 forward matrix                           */
   int              *matassign = NULL;                 /* MAT state assignments if 1; 1..alen      */
   float             hmmsc;                            /* Joint hmm score of seq and path, in nats */
   float             ntsc;                             /* Null transition score under hmmer's hmm  */
   float             fsc;                              /* forward score of a sequence, in nats     */
   int               i;                                /* msa position (column) index              */
   int               n;                                /* msa sequence (row) index                 */
   int               fmt       = eslMSAFILE_STOCKHOLM;
   int               status;                           /* easel return code                        */

   /* parse command line */
   go = esl_getopts_Create(options);
   if (esl_opt_ProcessCmdline(go, argc, argv) != eslOK) cmdline_failure(argv[0], "Failed to parse command line: %s\n", go->errbuf);
   if (esl_opt_VerifyConfig(go)               != eslOK) cmdline_failure(argv[0], "Error in app configuration:   %s\n", go->errbuf);

   /* output help if requested */
   if (esl_opt_GetBoolean(go, "-h") ) {
      esl_banner(stdout, argv[0], banner);
      esl_usage (stdout, argv[0], usage);
      puts("\n where options are:");
      esl_opt_DisplayHelp(stdout, go, 1, 2, 80);
      puts("\n Alphabet options:");
      esl_opt_DisplayHelp(stdout, go, 2, 2, 80);
      exit(0);
   }

   if       (esl_opt_GetBoolean(go, "--amino")) abc = esl_alphabet_Create(eslAMINO);
   else if  (esl_opt_GetBoolean(go, "--dna"))   abc = esl_alphabet_Create(eslDNA);
   else if  (esl_opt_GetBoolean(go, "--rna"))   abc = esl_alphabet_Create(eslRNA);


   /* read arguments */
   if (esl_opt_ArgNumber(go) != 2)  cmdline_failure(argv[0], "Incorrect number of command line arguments.\n", go->errbuf);

   hmmfile = esl_opt_GetArg(go, 1);
   msafile = esl_opt_GetArg(go, 2);

   /* read the hmm file */
   if (p7_hmmfile_OpenE(hmmfile, NULL, &hmmfp, NULL) != eslOK) p7_Fail("Failed to open HMM file %s", hmmfile);
   if (p7_hmmfile_Read(hmmfp, &abc, &hmm)              != eslOK) p7_Fail("Failed to read HMM");
   p7_hmmfile_Close(hmmfp);

   /* create a profile from the HMM */
   gm = p7_profile_Create(hmm->M, hmm->abc);

   /* create null model*/
   bg = p7_bg_Create(hmm->abc);

   /* configure profile with dummy length (we'll change it later) */
   p7_ProfileConfig(hmm, bg, gm, 100, p7_UNIGLOCAL);


   /* allow '-' and '.' as gap */
   esl_alphabet_SetEquiv(abc, '.', '-');

   /* read the msa file */
   if ((status = esl_msafile_Open(&abc, msafile, NULL, fmt, NULL, &afp)) != eslOK) esl_msafile_OpenFailure(afp, status);
   if ((status = esl_msafile_Read(afp, &msa))                            != eslOK) p7_Fail ("Failed to read MSA");
   esl_msafile_Close(afp);

   /* extract sequences from MSA */
   ESL_ALLOC(sq, sizeof(ESL_SQ *) * msa->nseq);
   for (n = 0; n < msa->nseq; n++) {
      fprintf(stdout, "n: %d\n", n);
      esl_sq_FetchFromMSA(msa, n, &sq[n]);
   }

   if (! (msa->flags & eslMSA_DIGITAL)) p7_Fail("need a digital msa");
   if (msa->rf == NULL)                 p7_Fail("msa lacks an RF line");

   /* extract match states from alignment */
   ESL_ALLOC(matassign, sizeof(int) * (msa->alen + 1));
   for (i=0; i < msa->alen; i++){
      if (msa->rf[i] != '.') matassign[i+1] = 1;
      else                   matassign[i+1] = 0;
   }

   /* generate trace array from msa */
   ESL_ALLOC(tr, sizeof(P7_TRACE *) * msa->nseq);
   p7_trace_FauxFromMSA(msa, matassign, p7_DEFAULT, tr);


   /* score sequences and paths */
   for (n=0;  n < msa->nseq; n++) {

      /* configure profile and background models using seq length */
      p7_ReconfigLength(gm, sq[n]->n);
      p7_bg_SetLength(bg, sq[n]->n);

      /* rescale the forward matrix */
      p7_gmx_GrowTo(fwd, gm->M, sq[n]->n);
      /* run forward algorithm */
      p7_GForward(sq[n]->dsq, sq[n]->n, gm, fwd, &fsc);

      /* get null transition score */
      p7_bg_NullOne(bg, sq[n]->dsq, sq[n]->n, &ntsc);

      /* score sequence and given path */
      p7_trace_Score(tr[n], sq[n]->dsq, gm, &hmmsc);

      /* add flanking tranition scores */
      /* THIS IS A HARDCODED HACK THAT I NEED TO FIX */
      /* p7_faux_from_trace doesn't include N and C states! */
      hmmsc += 2.0*(-3.8286);

      p7_trace_Dump(stdout, tr[n], gm, sq[n]->dsq);

      fprintf(stdout, "seq: %s\n", sq[n]->name);
      fprintf(stdout, "Raw forward score: %.2f\n", (fsc) / eslCONST_LOG2);
      fprintf(stdout, "Forward score of sequence: %.2f\n", (fsc - ntsc) / eslCONST_LOG2);
      fprintf(stdout, "HMM score of sequence, path: %.2f\n", (hmmsc) / eslCONST_LOG2);
      fprintf(stdout, "Log posterior probability of path under HMM:     %.2f\n", (hmmsc - fsc) / eslCONST_LOG2);
   }

   fprintf(stdout, "hello world\n");

   /* clean up and return */
   free(matassign);
   for (n = 0; n < msa->nseq; n++) esl_sq_Destroy(sq[n]);
   free(sq);
   p7_trace_DestroyArray(tr, msa->nseq);
   esl_msa_Destroy(msa);
   p7_profile_Destroy(gm);
   p7_bg_Destroy(bg);
   p7_hmm_Destroy(hmm);
   p7_gmx_Destroy(fwd);
   esl_alphabet_Destroy(abc);
   esl_getopts_Destroy(go);
   return 0;

   ERROR:
      return status;


}
```
