## Supplemental Code S1 for "Remote homology search with hidden Potts models": Makefile

```
#useful directories
ESLDIR = ../../infernal-1.1.3/easel
HMMERDIR = ../../infernal-1.1.3/hmmer/src
INFERNALDIR = ../../infernal-1.1.3/src
H4DIR = ../../hmmer4/nwo

#variables that I will run
INCLUDES = -I${INFERNALDIR} -I${HMMERDIR} -I${H4DIR} -I${ESLDIR}
IFLAGS   = -I${ESLDIR} -I${HMMERDIR} -I${INFERNALDIR} -I${H4DIR} -I../src
LFLAGS   = -L${ESLDIR} -L${HMMERDIR} -I${INFERNALDIR}  -I${H4DIR} -L../src
LIBS     = -lm
SLIBS    = ${INFERNALDIR}/libinfernal.a ${HMMERDIR}/libhmmer.a ${H4DIR}/libhmmer.a ${ESLDIR}/libeasel.a

#compiler information
CC = gcc
CFLAGS = -Wall -g

all: cmis_scoreset cmis_trace h4_pathalign h4_path_cmis cmscoreIS cmscoreISh4 cmscore_path cmemit_posterior h4score_path hmmscore_path

cmis_scoreset: cmis_scoreset.c
	${CC} ${CFLAGS} -c -o cmis_scoreset.o ${IFLAGS} cmis_scoreset.c

cmis_trace: cmis_trace.c
	${CC} ${CFLAGS} -c -o cmis_trace.o ${IFLAGS} cmis_trace.c

h4_pathalign: h4_pathalign.c
	${CC} ${CFLAGS} -c -o h4_pathalign.o ${IFLAGS} h4_pathalign.c

h4_path_cmis: h4_path_cmis.c
	${CC} ${CFLAGS} -c -o h4_path_cmis.o ${IFLAGS} h4_path_cmis.c

cmscoreIS: cmscoreIS.c
	${CC} ${CFLAGS} -o cmscoreIS ${IFLAGS} ${LFLAGS} cmscoreIS.c cmis_scoreset.o cmis_trace.o  ${SLIBS} ${LIBS}

cmscoreISh4: cmscoreISh4.c
	${CC} ${CFLAGS} -o cmscoreISh4 ${IFLAGS} ${LFLAGS} cmscoreISh4.c cmis_scoreset.o cmis_trace.o h4_path_cmis.o h4_pathalign.o  ${SLIBS} ${LIBS}

cmscore_path: cmscore_path.c
	${CC} ${CFLAGS} -o cmscore_path ${IFLAGS} ${LFLAGS} cmscore_path.c  ${SLIBS} ${LIBS}

cmemit_posterior: cmemit_posterior.c
	${CC} ${CFLAGS} -o cmemit_posterior ${IFLAGS} ${LFLAGS} cmemit_posterior.c h4_path_cmis.o  ${SLIBS} ${LIBS}

h4score_path: h4score_path.c
	${CC} ${CFLAGS} -o h4score_path ${IFLAGS} ${LFLAGS} h4score_path.c h4_path_cmis.o  ${SLIBS} ${LIBS}

hmmscore_path: hmmscore_path.c
	${CC} ${CFLAGS} -o hmmscore_path ${IFLAGS} ${LFLAGS} hmmscore_path.c  ${SLIBS} ${LIBS}
clean:
	rm -f cmscoreIS cmscoreISh4 cmemit_posterior cmscore_path h4score_path hmmscore_path cmis_scoreset.o cmis_trace.o h4_pathalign.o h4_path_cmis.o

.PHONY: all clean
```
