## Supplemental Code S1 for "Remote homology search with hidden Potts models": compile.sh

```
#!/bin/bash
# bash script to compile code

# compile Infernal
echo -e "Compiling Infernal\n\n\n"
cd infernal-1.1.3/

autoconf

./configure

make

cd ..

# compile HMMER4
echo -e "\n\n\nCompiling HMMER4\n\n\n"

# compile HMMER3-like parts of HMMER4/
cd hmmer4


# link easel (could be fixed...)
ln -s ${PWD}/easel lib/easel

autoconf

./configure

make

# compile HMMER4 nwo
cd nwo

make

cd ../..

# Install HPMHomology
echo -e "\n\n\nCompiling HPMHomology\n\n\n"

cd HPMHomology/src/

make

# compile Gremlin cpp code
g++ -O3 -std=c++0x -o gremlin_h2_henikoff gremlin_h2_henikoff.cpp

cd ../..

# compile CMIS
echo -e "\n\n\nCompiling CMIS\n\n\n"

cd CMIS/src/

make

cd ../..
```
