## Supplemental Code S1 for "Remote homology search with hidden Potts models": gremlin_h2_henikoff.cpp

```
#include <stdio.h>
#include <iostream>
#include <iomanip>
#include <vector>
#include <fstream>
#include <sstream>
#include <math.h>
#include <algorithm>
#include <string>

using namespace std;
typedef vector<bool> bool_1D;
typedef vector<bool_1D> bool_2D;
typedef vector<int> int_1D;
typedef vector<int_1D> int_2D;
typedef vector<double> double_1D;
typedef vector<double_1D> double_2D;
typedef vector<double_2D> double_3D;
typedef vector<string> string_1D;

class Msa{
public:
	// vars
	int_2D X;           // msa
	int_1D c2f;         // mapping of index from (c)ut to (f)ull msa
	int_1D f2c;
	double_1D eff;      // (eff)ctive weight for each sequence
	double neff;        // (n)umber of (eff)ective sequences
	double_1D H;        // Entropy
	double_1D f;        // frequency
	int nr;             // number of (r)ows in msa
	int nc;             // number of (c)olumns in msa
	int na;             // number of characters in (a)lphabet
	int na_gap;
	int pair_size;      // number of residue pairs
	int N1;             // number of 1bd vars
	int N2;             // number of 1bd+2bd vars
	double lam_v;       // lambda weight for 1bd penality
	double lam_w;       // lambda weight for 2bd penality

	// functions
	void load(string msa_i, string alphabet, double gap);
	void get_eff(double cutoff);
	void get_wgt_henikoff();
	void get_wgt_henikoff_neff();
	void get_H(bool only_v);

	int v_n(int i, int a){return(i*na + a);}
	int w_n(int w, int a, int b){return(N1 + w*na*na + a*na + b);}

	int_2D pair;
	void all_pair();
	void load_pair(string pair_i);
};

class Mrf{
public:
	double_1D x;
	double_1D g;

	void load(string file);
	void save(string file, Msa &msa, bool only_v);
	void saveRSC(string file, Msa &msa, bool only_v);
	void resize(int N){x.resize(N,0);g.resize(N,0);}
	size_t size(){return x.size();}
	void reset_g(){for(int n = 0; n < g.size(); n++){g[n] = 0;}}
};


class Opt{
public:
	string msa_i;
	string pair_i;
	string mrf_i;
	string mrf_o;
	string preds_out;
	string alphabet = "protein";
	string min_type = "lbfgs";
	int max_iter = 100;
	double gap_cutoff = 1.0;
	double eff_cutoff = 0.8;
	double lambda = 0.01;
	bool only_v = 0;
	void get(string_1D &arg);
};

// functions
int aa2int (char aa, string alphabet);

typedef double eval(Mrf &mrf, Msa &msa);
double eval_V(Mrf &mrf, Msa &msa);
double eval_VW(Mrf &mrf, Msa &msa);

void lbfgs(eval func, Mrf &mrf, Msa &msa, int max_iter);
void cg(eval func, Mrf &mrf, Msa &msa, int max_iter);

double_1D mrf2mtx(Mrf &mrf, Msa &msa);
void save_mtx(string preds_out, double_1D &mtx, Msa &msa);

void set_1D(double_1D &M, double val);
double vecdot(const double_1D &v1, const double_1D &v2);
double vec_sum(const double_1D &v1);
double vec_L2(const double_1D &v);
double vec_L2norm(const double_1D &v);

bool exists (const string &name){ifstream f(name);return f.good();}

int main(int argc, const char * argv[]) {
	// parse input arguments
	Opt opt; string_1D arg(argv+1,argv+argc); opt.get(arg);

	// load msa (multiple sequence alignment)
	Msa msa; msa.load(opt.msa_i,opt.alphabet,opt.gap_cutoff);

	// load list of pairs
	if(!opt.only_v){
		if(opt.pair_i.empty()){msa.all_pair();}
		else{msa.load_pair(opt.pair_i);}
	}else{msa.N1 = msa.nc * msa.na;}

	// get effective weight for each sequence
	msa.get_eff(opt.eff_cutoff);
	cout << "Neff: " << msa.neff << endl;
	// get henikoff weights
	//msa.get_wgt_henikoff();
	msa.get_wgt_henikoff_neff();

	// get entropy
	msa.get_H(opt.only_v);

	// lambda weight for V and W
	msa.lam_v = opt.lambda;
	msa.lam_w = opt.lambda * ((double)msa.nc - 1.0) * ((double)msa.na - 1.0);

	// mrf parameters
	Mrf mrf;

	// load mrf
	if(!opt.mrf_i.empty()){
		if(opt.only_v){mrf.resize(msa.N1);}
		else{mrf.resize(msa.N2);}
		mrf.load(opt.mrf_i);
	}
	// minimize using cg or lbfgs, which calls eval()
	cout << "# learning MRF ..." << endl;
	if(opt.min_type == "cg"){
		if(mrf.size() <= msa.N1){
			mrf.resize(msa.N1);
			cg(eval_V,mrf,msa,100);
		}
		if(!opt.only_v){
			mrf.resize(msa.N2);
			cg(eval_VW,mrf,msa,opt.max_iter);
		}
	}
	if(opt.min_type == "lbfgs"){
		if(mrf.size() <= msa.N1){
			mrf.resize(msa.N1);
			lbfgs(eval_V,mrf,msa,opt.max_iter);
		}
		if(!opt.only_v){
			mrf.resize(msa.N2);
			lbfgs(eval_VW,mrf,msa,opt.max_iter);
		}
	}
	if(!opt.only_v){
		// convert mrf to mtx by taking l2norm of each pair 20x20 matrix
		double_1D mtx = mrf2mtx(mrf,msa);
		// save mtx and apply entropy correction
		save_mtx(opt.preds_out,mtx,msa);
	}
	// save mrf
	if(!opt.mrf_o.empty()){
		//mrf.save(opt.mrf_o,msa,opt.only_v);
		mrf.saveRSC(opt.mrf_o,msa,opt.only_v);

	}
	return 0;
}
double eval_V(Mrf &mrf, Msa &msa){
	mrf.reset_g();

	// function we want to maximize
	double fx = 0;
	double reg = 0;
	double_2D PC(msa.nc,double_1D(msa.na,0));

	#pragma omp parallel for
	for(int i = 0; i < msa.nc; i++){
		int d = msa.v_n(i,0);
		for(int a = 0; a < msa.na; a++){
			PC[i][a] += mrf.x[d];
			d++;
		}
	}
	#pragma omp parallel for reduction(+:fx)
	for(int i = 0; i < msa.nc; i++){
		// compute Z
		double Z = 0;
		for(int a = 0; a < msa.na; a++){
			Z += exp(PC[i][a]);
		}
		Z = log(Z);
		// compute fx
		for(int a = 0; a < msa.na; a++){
			fx += msa.f[msa.v_n(i,a)] * (PC[i][a] - Z);
			PC[i][a] = exp(PC[i][a] - Z);
		}
	}
	// compute gradient and reg
	#pragma omp parallel for reduction(+:reg)
	for(int i = 0; i < msa.nc; i++){
		int d = msa.v_n(i, 0);
		for(int a = 0; a < msa.na; a++){
			mrf.g[d] += (PC[i][a]*msa.neff) - (msa.f[msa.v_n(i,a)]) + (msa.lam_v*2*mrf.x[d]);
			reg += msa.lam_v * pow(mrf.x[d],2);
			d++;
		}
	}
	// flip direction, since we are passing function to a minimizer
	return -1.0 * (fx-reg);
}
double eval_VW(Mrf &mrf, Msa &msa){

	mrf.reset_g();
	int N = mrf.size();
	// function we want to maximize
	double fx = 0;
	double reg = 0;

	double_3D PCN(msa.nr,double_2D(msa.nc,double_1D(msa.na,0)));
	// for each sequence
	#pragma omp parallel for reduction(+:fx)
	for(int n = 0; n < msa.nr; n++){
		// precompute sum(V+W) for each position "i" and amino acids "a"
		// assuming all other positions are fixed
		double_2D PC(msa.nc,double_1D(msa.na,0));

		// for each position i
		for(int i = 0; i < msa.nc; i++){
			// for each amino acid
			for(int a = 0; a < msa.na; a++){
				// 1bd
				PC[i][a] += mrf.x[msa.v_n(i,a)];
			}
		}
		if(N == msa.N2){
			for(int w = 0; w < msa.pair_size; w++){
				int i = msa.pair[w][0];
				int j = msa.pair[w][1];
				int xni = msa.X[n][i];
				int xnj = msa.X[n][j];
				for(int a = 0; a < msa.na; a++){
					PC[i][a] += mrf.x[msa.w_n(w,a,xnj)];
					PC[j][a] += mrf.x[msa.w_n(w,xni,a)];
				}
			}
		}
		for(int i = 0; i < msa.nc; i++){
			// compute local Z
			double Z = 0;
			for(int a = 0; a < msa.na; a++){Z += exp(PC[i][a]);}
			Z = log(Z);

			// compute fx
			int xni = msa.X[n][i];
			fx += (PC[i][xni] - Z) * msa.eff[n];

			// needed for (g)radient calculation
			for(int a = 0; a < msa.na; a++){
				PCN[n][i][a] = exp(PC[i][a] - Z) * msa.eff[n];
			}
		}
	}
	if(N >= msa.N1){
		// compute (g)radient for 1bd
		#pragma omp parallel for
		for(int i = 0; i < msa.nc; i++){
			for(int n = 0; n < msa.nr; n++){
				int xni = msa.X[n][i];
				mrf.g[msa.v_n(i,xni)] -= msa.eff[n];
				for(int a = 0; a < msa.na; a++){
					mrf.g[msa.v_n(i,a)] += PCN[n][i][a];
				}
			}
		}
	}
	if(N == msa.N2){
		// compute (g)radient for 2bd
		#pragma omp parallel for
		for(int w = 0; w < msa.pair_size; w++){
			int i = msa.pair[w][0];
			int j = msa.pair[w][1];
			for(int n = 0; n < msa.nr; n++){
				int xni = msa.X[n][i];
				int xnj = msa.X[n][j];
				mrf.g[msa.w_n(w,xni,xnj)] -= 2.0 * msa.eff[n];
				for(int a = 0; a < msa.na; a++){
					mrf.g[msa.w_n(w,a,xnj)] += PCN[n][i][a];
					mrf.g[msa.w_n(w,xni,a)] += PCN[n][j][a];
				}
			}
		}
	}
	if(N >= msa.N1){
		// compute (reg)ularization and (g)raident for 1bd
		#pragma omp parallel for reduction(+:reg)
		for(int d = 0; d < msa.N1; d++){
			reg  += msa.lam_v * pow(mrf.x[d],2);
			mrf.g[d] += msa.lam_v * 2.0 * mrf.x[d];
		}
	}
	if(N == msa.N2){
		// compute (reg)ularization and (g)raident for 2bd
		#pragma omp parallel for reduction(+:reg)
		for(int d = msa.N1; d < msa.N2; d++){
			reg  += msa.lam_w * pow(mrf.x[d],2);
			mrf.g[d] += msa.lam_w * 2.0 * mrf.x[d];
		}
	}
	// flip direction, since we are passing function to a minimizer
	return -1.0 * (fx-reg);
}
void lbfgs(eval func, Mrf &mrf, Msa &msa, int max_iter){

	/* ----------------------------------------------------------------------------
	 * Limited-memory Broyden–Fletcher–Goldfarb–Shanno (LBFGS)
	 * ---------------------------------------------------------------------------
	 * Adopted from: https://github.com/js850/lbfgs_cpp
	 * "THE BEER-WARE LICENSE" (Revision 42):
	 * This <> wrote this function. As long as you retain this notice
	 * you can do whatever you want with this stuff. If we meet some day, and you
	 * think this stuff is worth it, you can buy me a beer in return Jacob Stevenson
	 * ---------------------------------------------------------------------------
	 * modified to remove convergence criteria, will continue until max_iter
	 * modified to remove maxstep check
	 * ---------------------------------------------------------------------------*/

	double max_f_rise = 1e-4;
	double H0 = 0.1;
	//double maxstep = 0.2;

	size_t N = mrf.size();
	int M = 5;

	// allocate memory
	double_2D y(M,double_1D(N));
	double_2D s(M,double_1D(N));
	double_1D rho(M);
	double_1D step(N);

	Mrf mrf_new;
	mrf_new.resize(N);

	double f = func(mrf,msa);
	cout << "# lbfgs::iter S_S fx: " << f << " gnorm: "<< vec_L2norm(mrf.g) << endl;
	for(int iter = 0; iter < max_iter; iter++){
		/////////////////////////////////
		// compute_lbfgs_step
		/////////////////////////////////
		if(iter == 0){
			double gnorm = vec_L2norm(mrf.g);
			if(gnorm > 1.0){gnorm = 1.0/gnorm;}
			#pragma omp parallel for
			for(int n = 0; n < N; n++){step[n] = -gnorm * H0 * mrf.g[n];}
		}
		else{
			step = mrf.g;
			int jmin = iter-M;if(jmin < 0){jmin = 0;}
			int jmax = iter;

			int i;
			double beta;
			double_1D alpha(M);
			// loop backwards through the memory
			for(int j = jmax - 1; j >= jmin; j--){
				i = j % M;
				alpha[i] = rho[i] * vecdot(s[i],step);
				#pragma omp parallel for
				for(int n = 0; n < N; n++){step[n] -= alpha[i] * y[i][n];}
			}
			// scale the step size by H0
			#pragma omp parallel for
			for(int n = 0; n < N; n++){step[n] *= H0;}
			// loop forwards through the memory
			for(int j = jmin; j < jmax; j++){
				i = j % M;
				beta = rho[i] * vecdot(y[i],step);
				#pragma omp parallel for
				for(int n = 0; n < N; n++){step[n] += s[i][n] * (alpha[i] - beta);}
			}
			// invert the step to point downhill
			#pragma omp parallel for
			for(int n = 0; n < N; n++){step[n] *= -1;}
		}
		/////////////////////////////////
		// backtracking_linesearch
		/////////////////////////////////
		double fnew;
		// if the step is pointing uphill, invert it
		if(vecdot(step,mrf.g) > 0.0){
			#pragma omp parallel for
			for(int n = 0; n < N; n++){step[n] *= -1;}
		}
		int attempt = 0;
		double factor = 1.0;
		double stepsize = vec_L2norm(step);

		// make sure the step is no larger than maxstep
		/* if (factor * stepsize > maxstep){factor = maxstep/stepsize;} */

		for(int nred = 0; nred < 10; nred++){
			attempt++;
			#pragma omp parallel for
			for(int n = 0; n < N; n++){mrf_new.x[n] = mrf.x[n] + factor * step[n];}
			fnew = func(mrf_new,msa);
			double df = fnew - f;
			if(df < max_f_rise){break;}
			else{factor /= 10.0;}
		}
		stepsize = stepsize * factor;
		/////////////////////////////////
		// update_memory
		/////////////////////////////////
		int klocal = iter % M;
		#pragma omp parallel for
		for(int n = 0; n < N; n++){
			y[klocal][n] = mrf_new.g[n] - mrf.g[n];
			s[klocal][n] = mrf_new.x[n] - mrf.x[n];
		}
		double ys = vecdot(y[klocal],s[klocal]);
		if(ys == 0.0){ys = 1.0;}
		rho[klocal] = 1.0/ys;

		double yy = vecdot(y[klocal],y[klocal]);
		if(yy == 0.0){yy = 1.0;}
		H0 = ys/yy;

		mrf.x = mrf_new.x;
		mrf.g = mrf_new.g;
		f = fnew;
		cout << "# lbfgs::iter " << iter << "_" << attempt << " fx: " << f << " gnorm: " << vec_L2norm(mrf.g) << endl;
	}
}
void cg(eval func, Mrf &mrf, Msa &msa, int max_iter)
{
	/* ----------------------------------------------------------------------------
	 * Nonlinear Conjugate Gradient (CG)
	 * ---------------------------------------------------------------------------
	 * Adopted from: https://bitbucket.org/soedinglab/libconjugrad
	 * CCMpred is released under the GNU Affero General Public License v3 or later.
	 * ---------------------------------------------------------------------------
	 * modified to remove convergence criteria, will continue until max_iter
	 * ---------------------------------------------------------------------------*/
	////////////////////////////////////
	double epsilon = 1e-5;
	double ftol = 1e-4;
	double wolfe = 0.1;
	double alpha_mul = 0.5;
	int max_line = 10;
	////////////////////////////////////
	int N = mrf.size();

	double_1D s(N);

	double gnorm_old = 0;
	double alpha_old = 0;
	double dg_old = 0;

	double fx = func(mrf,msa);

	double gnorm = vec_L2(mrf.g);

	double dg = 0;
	double alpha = 1/sqrt(gnorm);

	cout << "# cg::iter S_S fx: " << fx << " gnorm: " << sqrt(gnorm) << endl;

	for(int iter = 0; iter < max_iter; iter++){
		if(iter == 0){
			#pragma omp parallel for
			for(int n = 0; n < N; n++){s[n] = -mrf.g[n];}
			dg = vecdot(s,mrf.g);
		}else{
			// fletcher-reeves
			double beta = gnorm / gnorm_old;
			#pragma omp parallel for
			for(int n = 0; n < N; n++){s[n] = s[n] * beta - mrf.g[n];}
			dg = vecdot(s,mrf.g);
			alpha = alpha_old * dg_old/dg;
		}
		/////////////////////////////////////////////////////
		// linesearch
		////////////////////////////////////////////////////
		int attempts = 0;
		double dg_ini = dg;
		double dg_test = dg_ini * ftol;
		double fx_ini = fx;
		double old_alpha = 0;
		for(int line = 0; line < max_line; line++){
			attempts++;
			double step = alpha-old_alpha;
			#pragma omp parallel for
			for(int n = 0; n < N; n++){mrf.x[n] += s[n] * step;}
			double fx_step = func(mrf,msa);
			if(fx_step <= fx_ini + alpha * dg_test){
				if(vecdot(s,mrf.g) < wolfe * dg_ini){
					fx = fx_step;
					break;
				}
			}
			old_alpha = alpha;
			alpha *= alpha_mul;
		}
		//////////////////////////////////////////////////////
		gnorm_old = gnorm;
		gnorm = vec_L2(mrf.g);
		alpha_old = alpha;
		dg_old = dg;
		cout << "# cg::iter " << iter << "_" << attempts << " fx: " << fx << " gnorm: " << sqrt(gnorm) << endl;
	}
}
///////////////////////////////////////////////////////////////////
void Msa::load(string msa_i, string alphabet, double gap){

	string format = "aln";
	string line;
	ifstream in(msa_i);

	ostringstream out;
	int_2D X_tmp;
	int line_n = 0;
	while(getline(in,line)){
		istringstream is(line);
		string seq;
		is >> seq;
		if(line_n == 0 && seq[0] == '>'){format = "fasta";}
		if(format == "aln" || (format == "fasta" && seq[0] == '>')){
			X_tmp.push_back(int_1D());
			int n = X_tmp.size() - 1;
			if(n > 0 && X_tmp[0].size() != X_tmp[n-1].size()){
				// check that all sequences of same length
				cout << "# ERROR: sequence #" << n-1 << " length " << X_tmp[n-1].size() << " != " << X_tmp[0].size() << endl;
				exit(1);
			}
		}
		if(format == "aln" || (format == "fasta" && seq[0] != '>')){
			int n = X_tmp.size() - 1;
			for(int i = 0; i < seq.size(); i++){
				X_tmp[n].push_back(aa2int(seq[i],alphabet));
			}
		}
		line_n++;
	}
	// remove positions with too many gaps
	int gap_int = aa2int('-',alphabet);
	int nrow = X_tmp.size();
	int ncol = X_tmp[0].size();
	int gap_cutoff = nrow * gap;

	X.resize(nrow,int_1D());

	int c = 0;
	for(int i = 0; i < ncol; i++){
		int gaps = 0;
		for(int n = 0; n < nrow; n++){
			if(X_tmp[n][i] == gap_int){gaps++;}
		}
		if(gap_cutoff > gaps){
			for(int n = 0; n < nrow; n++){
				X[n].push_back(X_tmp[n][i]);
			}
			c2f.push_back(i);
			f2c.push_back(c);
			c++;
		}else{
			f2c.push_back(-1);
		}
	}
	int pos_rm = ncol - X[0].size();
	if(pos_rm > 0){
		cout << "# removing " << pos_rm << " out of " << ncol << " positions with >= " << gap*100 << "% gaps!" << endl;
	}

	// compute size of various vectors
	nr = X.size();
	nc = X[0].size();
	na = 21;
	if(alphabet == "rna"){na = 5;}
	if(alphabet == "binary"){na = 3;}
	na_gap = na-1;

	in.close();
}

void Msa::get_eff(double cutoff){
	eff.resize(nr,1);
	int_1D N(nr,1);
	int chk = nc * cutoff;
	for(int n = 0; n < nr; n++){
		//cout << "\t\tn: " << n << endl;
		int w = N[n];
		#pragma omp parallel for reduction(+:w)
		for(int m=n+1; m < nr; m++){
			//cout << "\t\tn: " << n << " m: " << m << endl;

			int hm = 0;
			for(int i = 0; i < nc; i++){
				if(X[n][i] == X[m][i]){hm++;}
			}
			if(hm > chk){N[m]++;w++;}
		}
		eff[n] = 1.0/(double)w;
		/* added by grey */
		//cout << "\tn: " << n << " m: " << w << " wgt: " << eff[n] << endl;
	}
	neff = vec_sum(eff);
	//cout << "# nc: " << nc << " neff: " << neff << endl;
}

void Msa::get_wgt_henikoff(){

	/* declare variables */
	int     i;                          /* column index */
	int     n;                          /* row (seq) index */
	int     a;                          /* residue index  */
	int     gap      = na-1;            /* gap idx */
	double  wgt_sum  = na-1;            /* sum of unnormalized weights */
	int*    r        = new int[nc];     /* number of different canonical residues used in each consensus column. r[j=0..ncons-1] */
	int*    l        = new int[nr];     /* length of sequence (match cols occupied) */
	double* wgt_un   = new double[nr];  /* unnormalized weghts */
	double* wgt      = new double[nr];  /* normalized weights */
	int**   ct       = new int*[nc];    /* matrix of symbol counts in each column. ct[0,...nc-1][0,..na] */
	for (i=0; i < nc; i++){
		ct[i] = new int[na-1];
		for (a=0; a < gap; a++) ct[i][a] = 0;
	}

	/* step 1: fill count array */
	for (n = 0; n < nr; n++){
		for (i = 0; i < nc; i++) {
			//cout << "seq: " << n << " col: " << i <<  " X[n][i]: " << X[n][i] << endl;
			if (X[n][i] != gap) ct[i][X[n][i]]++;
		}
	}

	/* print out count matrix */
	//for (i=0; i < nc; i++) {
	//	for (a=0; a<gap; a++){
	//		cout << "col: " << i << " resi: " << a << " ct[i][a]: " << ct[i][a] << endl;
	//	}
	//}

	/* step 2: find number of unique residues in each column */
	for (i = 0; i < nc; i++) {
		//cout << "col: " << i << endl;
		r[i] = 0;
		for (a = 0; a < gap; a++) {
			//cout << "\ta: " << a << endl;

			if (ct[i][a] > 0) r[i]++;
		}
		//cout << "col: " << i << " r[i]: " << r[i] << endl;
	}

	/* step 3: set unnormalized sequence weights */
	wgt_sum = 0.0;
	for (n = 0; n < nr; n++){
		wgt_un[n] = 0.0;
		wgt[n] = 0.0;
		l[n] = 0;
		for (i = 0; i < nc; i++){
			a = X[n][i];
			if (a != gap) {
				l[n]++;
				wgt_un[n] += 1.0/ (double)(r[i] * ct[i][a]);
			}
		}
		//cout << "seq: " << n << " len: " << l[n] << " wgt_un_un " << wgt_un[n] << endl;
		if (l[n] > 0) wgt_un[n] /= (double) l[n];
		//cout << " \twgt_un_ " << wgt_un[n] << endl;
		wgt_sum += wgt_un[n];
	}

	/* step 4: normalize weights so they sum to N */
	for (n = 0; n < nr; n++) {
		wgt[n] = wgt_un[n] * ( (double) nr / wgt_sum);
		eff[n] = wgt_un[n] * ( (double) nr / wgt_sum);
		cout << "seq: " << n  << " wgt: " << wgt[n] << endl;
	}

}


void Msa::get_wgt_henikoff_neff(){
	cout << "\thello world!" << endl;
	cout << "\tneff: " << neff << endl;

	/* declare variables */
	int     i;                          /* column index */
	int     n;                          /* row (seq) index */
	int     a;                          /* residue index  */
	int     gap      = na-1;            /* gap idx */
	double  wgt_sum  = na-1;            /* sum of unnormalized weights */
	int*    r        = new int[nc];     /* number of different canonical residues used in each consensus column. r[j=0..ncons-1] */
	int*    l        = new int[nr];     /* length of sequence (match cols occupied) */
	double* wgt_un   = new double[nr];  /* unnormalized weghts */
	double* wgt      = new double[nr];  /* normalized weights */
	int**   ct       = new int*[nc];    /* matrix of symbol counts in each column. ct[0,...nc-1][0,..na] */
	for (i=0; i < nc; i++){
		ct[i] = new int[na-1];
		for (a=0; a < gap; a++) ct[i][a] = 0;
	}

	/* step 1: fill count array */
	for (n = 0; n < nr; n++){
		for (i = 0; i < nc; i++) {
			//cout << "seq: " << n << " col: " << i <<  " X[n][i]: " << X[n][i] << endl;
			if (X[n][i] != gap) ct[i][X[n][i]]++;
		}
	}


	/* step 2: find number of unique residues in each column */
	for (i = 0; i < nc; i++) {
		//cout << "col: " << i << endl;
		r[i] = 0;
		for (a = 0; a < gap; a++) {
			//cout << "\ta: " << a << endl;

			if (ct[i][a] > 0) r[i]++;
		}
		//cout << "col: " << i << " r[i]: " << r[i] << endl;
	}

	/* step 3: set unnormalized sequence weights */
	wgt_sum = 0.0;
	for (n = 0; n < nr; n++){
		wgt_un[n] = 0.0;
		wgt[n] = 0.0;
		l[n] = 0;
		for (i = 0; i < nc; i++){
			a = X[n][i];
			if (a != gap) {
				l[n]++;
				wgt_un[n] += 1.0/ (double)(r[i] * ct[i][a]);
			}
		}
		//cout << "seq: " << n << " len: " << l[n] << " wgt_un_un " << wgt_un[n] << endl;
		if (l[n] > 0) wgt_un[n] /= (double) l[n];
		//cout << " \twgt_un_ " << wgt_un[n] << endl;
		wgt_sum += wgt_un[n];
	}

	/* step 4: normalize weights so they sum to Neff */
	for (n = 0; n < nr; n++) {
		wgt[n] = wgt_un[n] * ( (double) neff / wgt_sum);
		eff[n] = wgt_un[n] * ( (double) neff / wgt_sum);
		cout << "seq: " << n  << " wgt: " << wgt[n] << endl;
	}

}


void Msa::get_H(bool only_v){
	if(only_v){
		f.resize(N1,0);
		H.resize(nc,0);
	}else{
		f.resize(N2,0);
		H.resize(nc+pair_size,0);
	}
	#pragma omp parallel for
	for(int i = 0; i < nc; i++){
		for(int n = 0; n < nr; n++){
			int d = v_n(i,X[n][i]);
			f[d] += eff[n];
		}
		for(int a = 0; a < na; a++){
			int d = v_n(i,a);
			if(f[d] > 0){
				double P = f[d]/neff;
				H[i] += fabs(P*log(P));
			}
		}
	}
	if(!only_v){
		#pragma omp parallel for
		for(int w = 0; w < pair_size; w++){
			int i = pair[w][0];
			int j = pair[w][1];
			for(int n = 0; n < nr; n++){
				int xni = X[n][i];
				int xnj = X[n][j];
				int d = w_n(w,xni,xnj);
				f[d] += eff[n];
			}
			for(int a = 0; a < na; a++){
				for(int b = 0; b < na; b++){
					int d = w_n(w,a,b);
					if(f[d] > 0){
						double P = f[d]/neff;
						H[nc+w] += fabs(P*log(P));
					}
				}
			}
		}
	}
}
///////////////////////////////////////////////////////////////////
void Mrf::load(string file){
	cout << "# loading MRF: " << file << endl;
	string line;
	ifstream in(file);
	if(in.is_open()){
		int n = 0;
		while(getline(in,line)){
			istringstream is(line);
			string tag; is >> tag;
			while(n < x.size() and is >> x[n]){
				n++;
				//if(n > x.size()){cout << "# ERROR: input MRF size > "<< x.size() << endl; exit(1);}
			}
		}
		in.close();
	}
}
void Mrf::save(string file, Msa &msa, bool only_v){
	cout << "# saving MRF: " << file << endl;

	ofstream out(file);
	if(out.is_open()){
		int n = 0;
		for(int i = 0; i < msa.nc; i++){
			out << "V[" << msa.c2f[i] << "]";
			for(int a = 0; a < msa.na; a++){
				out << " " << x[n];
				n++;
			}
			out << endl;
		}
		if(!only_v){
			for(int w = 0; w < msa.pair_size; w++){
				int i = msa.pair[w][0];
				int j = msa.pair[w][1];
				out << "W[" << msa.c2f[i] << "][" << msa.c2f[j] << "]";
				for(int a = 0; a < msa.na; a++){
					for(int b = 0; b < msa.na; b++){
						out << " " << x[n];
						n++;
					}
				}
				out << endl;
			}
		}
		out.close();
	}
}

void Mrf::saveRSC(string file, Msa &msa, bool only_v){
	cout << "# saving MRF: " << file << endl;

	ofstream out(file);
	/* only print first 4 digits after decimal for floats */
	/* equivalent to %.4f in C */
	out.setf(std::ios_base::fixed, std::ios_base::floatfield);
	out.precision(4);

	if(out.is_open()){
		int n = 0;
		for(int i = 0; i < msa.nc; i++){
			out << msa.c2f[i];
			for(int a = 0; a < msa.na; a++){
				out << " " <<  x[n];
				n++;
			}
			out << endl;
		}
		/* print coupling terms (if they exist) */
		if(!only_v){
			for(int w = 0; w < msa.pair_size; w++){
				int i = msa.pair[w][0];
				int j = msa.pair[w][1];
				out <<  msa.c2f[i] << " " << msa.c2f[j] ;
				out << endl;
				for(int a = 0; a < msa.na; a++){
					for(int b = 0; b < msa.na; b++){
						out << x[n] <<" ";
						n++;
					}
					out << endl;

				}
				out << endl;
			}
		}
		else {
			for(int i = 0; i < msa.nc; i++){
				for(int j = i+1; j < msa.nc; j++){
					out <<  msa.c2f[i] << " " << msa.c2f[j] << endl;
					for(int a = 0; a < msa.na; a++){
						for(int b = 0; b < msa.na; b++){
							out << "0.0000" << " ";
						}
						out << endl;
					}
					out << endl;
				}
			}
		}
		/* print zeros in place of coupling terms if they dont exist */
		out.close();
	}
}


void Msa::load_pair(string file){
	int len = f2c.size();
	string line;
	ifstream in(file);
	if(in.is_open()){
		int i;
		int j;
		while(getline(in,line)){
			istringstream is(line);
			is >> i; is >> j;
			if(i > j){int tmp_i = i; i = j; j = tmp_i;}
			if(i == j){
				cout << "# pair invalid: " << i << " " << j << endl;
			}
			else if(i < len && j < len){
				if(f2c[i] != -1 && f2c[j] != -1){
					pair.push_back({f2c[i],f2c[j]});
				}
				else{cout << "# pair within gap region: " << i << " " << j << endl;}
			}
			else{cout << "# pair out of range: " << i << " " << j << endl;}
		}
		in.close();
	}
	pair_size = pair.size();
	N1 = nc * na;
	N2 = N1 + pair_size * na * na;
}
void Msa::all_pair(){
	for(int i = 0; i < nc; i++){
		for(int j = i+1; j < nc; j++){
			pair.push_back({i,j});
		}
	}
	pair_size = (nc*(nc-1))/2;
	N1 = nc * na;
	N2 = N1 + pair_size * na * na;
}
double_1D mrf2mtx(Mrf &mrf, Msa &msa){
	double_1D mtx(msa.pair_size,0);
	for(int w = 0; w < msa.pair_size; w++){
		double l2 = 0;
		for(int a = 0; a < msa.na_gap; a++){
			for(int b = 0; b < msa.na_gap; b++){
				l2 +=  pow(mrf.x[msa.w_n(w,a,b)], 2);
			}
		}
		mtx[w] = sqrt(l2);
	}
	return(mtx);
}
void save_mtx(string preds_out, double_1D &mtx, Msa &msa){
	ofstream out(preds_out);
	if(out.is_open()){

		double_1D rtot(msa.nc,0);
		double tot = 0;

		for(int w = 0; w < msa.pair_size; w++){
			int i = msa.pair[w][0];
			int j = msa.pair[w][1];

			rtot[i] += mtx[w];
			rtot[j] += mtx[w];
			tot += 2 * mtx[w];

		}
		out << "#W i j raw apc jen" << endl;
		for(int w = 0; w < msa.pair_size; w++){
			int i = msa.pair[w][0];
			int j = msa.pair[w][1];

			double raw = mtx[w];
			double apc = mtx[w] - (rtot[i] * rtot[j])/tot;
			double jen = mtx[w] / (1.0 + msa.H[msa.nc+w]);

			out << "W " << msa.c2f[i] << " " << msa.c2f[j] << " " << raw << " " << apc << " " << jen << endl;
		}
		out.close();
	}
}

int aa2int(char aa, string alphabet){
	if(alphabet == "protein"){
		if(aa == 'A'){return 0;}
		else if(aa == 'C'){return 1;}
		else if(aa == 'D'){return 2;}
		else if(aa == 'E'){return 3;}
		else if(aa == 'F'){return 4;}
		else if(aa == 'G'){return 5;}
		else if(aa == 'H'){return 6;}
		else if(aa == 'I'){return 7;}
		else if(aa == 'K'){return 8;}
		else if(aa == 'L'){return 9;}
		else if(aa == 'M'){return 10;}
		else if(aa == 'N'){return 11;}
		else if(aa == 'P'){return 12;}
		else if(aa == 'Q'){return 13;}
		else if(aa == 'R'){return 14;}
		else if(aa == 'S'){return 15;}
		else if(aa == 'T'){return 16;}
		else if(aa == 'V'){return 17;}
		else if(aa == 'W'){return 18;}
		else if(aa == 'Y'){return 19;}
		else              {return 20;}
	}
	else if(alphabet == "protein_sergey"){
		if(aa == 'A'){return 0;}
		else if(aa == 'R'){return 1;}
		else if(aa == 'N'){return 2;}
		else if(aa == 'D'){return 3;}
		else if(aa == 'C'){return 4;}
		else if(aa == 'Q'){return 5;}
		else if(aa == 'E'){return 6;}
		else if(aa == 'G'){return 7;}
		else if(aa == 'H'){return 8;}
		else if(aa == 'I'){return 9;}
		else if(aa == 'L'){return 10;}
		else if(aa == 'K'){return 11;}
		else if(aa == 'M'){return 12;}
		else if(aa == 'F'){return 13;}
		else if(aa == 'P'){return 14;}
		else if(aa == 'S'){return 15;}
		else if(aa == 'T'){return 16;}
		else if(aa == 'W'){return 17;}
		else if(aa == 'Y'){return 18;}
		else if(aa == 'V'){return 19;}
		else              {return 20;}
	}

	else if(alphabet == "rna" || alphabet == "dna"){
		if(aa == 'A'){return 0;}
		else if(aa == 'C'){return 1;}
		else if(aa == 'G'){return 2;}
		else if(aa == 'U'){return 3;}
		else if(aa == 'T'){return 3;}
		else              {return 4;}
	}
	else if(alphabet == "rna_sergey" || alphabet == "dna_sergey"){
		if(aa == 'A'){return 0;}
		else if(aa == 'U'){return 1;}
		else if(aa == 'T'){return 1;}
		else if(aa == 'C'){return 2;}
		else if(aa == 'G'){return 3;}
		else              {return 4;}
	}


	else if(alphabet == "binary"){
		if(aa == '0'){return 0;}
		if(aa == '1'){return 1;}
		else{return 2;}
	}
	else{return 0;}
}

// math functions
void set_1D(double_1D &M,double val){
#pragma omp parallel for
	for(int i = 0; i < M.size(); i++){M[i] = val;}
}
double vecdot(const double_1D &v1, const double_1D &v2){
	double dot = 0;
#pragma omp parallel for reduction(+:dot)
	for (int i=0; i < v1.size(); i++){dot += v1[i] * v2[i];}
	return dot;
}
double vec_sum(const double_1D &v1){
	double sum = 0;
#pragma omp parallel for reduction(+:sum)
	for (int i=0; i < v1.size(); i++){sum += v1[i];}
	return sum;
}
double vec_L2(const double_1D &v){return vecdot(v,v);}
double vec_L2norm(const double_1D &v){return sqrt(vecdot(v,v));}

void Opt::get(string_1D &arg)
{
	for (int a = 0; a < arg.size(); a++)
	{
		string val = arg[a];
		if (val[0] == '-')
		{
			if(val == "-i"              ){msa_i      = arg[a+1]; a++;}
			else if(val == "-o"         ){preds_out  = arg[a+1]; a++;}
			else if(val == "-pair_i"    ){pair_i     = arg[a+1]; a++;}
			else if(val == "-mrf_i"     ){mrf_i      = arg[a+1]; a++;}
			else if(val == "-mrf_o"     ){mrf_o      = arg[a+1]; a++;}
			else if(val == "-alphabet"  ){alphabet   = arg[a+1]; a++;}
			else if(val == "-min_type"  ){min_type   = arg[a+1]; a++;}
			else if(val == "-max_iter"  ){max_iter   = stoi(arg[a+1]); a++;}
			else if(val == "-gap_cutoff"){gap_cutoff = stof(arg[a+1]); a++;}
			else if(val == "-eff_cutoff"){eff_cutoff = stof(arg[a+1]); a++;}
			else if(val == "-lambda"    ){lambda     = stof(arg[a+1]); a++;}
			else if(val == "-only_v"    ){
				if(a+1 < arg.size() and arg[a+1][0] != '-'){
					only_v = stoi(arg[a+1]); a++;
				}else{
					only_v = 1;
				}
			}
		}
	}
	if(gap_cutoff > 1.0){gap_cutoff /= 100.0;}
	if(eff_cutoff > 1.0){eff_cutoff /= 100.0;}

	bool error = 0;

	ostringstream eout;
	if(msa_i.empty() || !exists(msa_i)){
		eout << "# ERROR: -i " << msa_i << endl;
		error = 1;
	}
	if(preds_out.empty()){
		eout << "# ERROR: -o " << preds_out << endl;
		error = 1;
	}
	if(!mrf_i.empty() && !exists(mrf_i)){
		eout << "# ERROR: -mrf_i " << mrf_i << endl;
		error = 1;
	}
	if(alphabet != "protein" && alphabet != "rna" && alphabet != "binary" && alphabet!= "protein_sergey" && alphabet != "rna_sergey"){
		eout << "# ERROR: -alphabet '" << alphabet << "' not valid" << endl;
		error = 1;
	}
	if(min_type != "lbfgs" && min_type != "cg" && min_type != "none"){
		eout << "# ERROR: -min_type '" << min_type << "' not valid" << endl;
		error = 1;
	}
	if(min_type == "none" && mrf_i.empty()){
		eout << "# ERROR: -min_type 'none' requires -mrf_i " << mrf_i << endl;
		error = 1;
	}
	if(gap_cutoff < 0.0 || gap_cutoff > 1.0){
		eout << "# ERROR: -gap_cutoff '" << gap_cutoff << "' should be between 0 and 1" << endl;
		error = 1;
	}
	if(eff_cutoff < 0.0 || eff_cutoff > 1.0){
		eout << "# ERROR: -eff_cutoff '" << eff_cutoff << "' should be between 0 and 1" << endl;
		error = 1;
	}
	if(lambda < 0.0){
		eout << "# ERROR: -lambda '" << lambda << "' should be > 0" << endl;
		error = 1;
	}

	if(error){
		cout << "# ---------------------------------------------------------------------------------------------" << endl;
		cout << "#                                GREMLIN_H (with entropy correction)                           " << endl;
		cout << "# ---------------------------------------------------------------------------------------------" << endl;
		cout << "#   -i            input alignment (either one sequence per line or in fasta format)"             << endl;
		cout << "#   -o            save output to"                                                                << endl;
		cout << "# ---------------------------------------------------------------------------------------------" << endl;
		cout << "#  Optional settings                                                                           " << endl;
		cout << "# ---------------------------------------------------------------------------------------------" << endl;
		cout << "#   -only_v       only compute v (1body-term)                    [Default=" << only_v     << "]" << endl;
		cout << "#   -gap_cutoff   remove positions with > X fraction gaps        [Default=" << gap_cutoff << "]" << endl;
		cout << "#   -alphabet     select: [protein|rna|binary]                   [Default=" << alphabet   << "]" << endl;
		cout << "#   -eff_cutoff   seq id cutoff for downweighting similar seqs   [Default=" << eff_cutoff << "]" << endl;
		cout << "#   -lambda       L2 regularization weight                       [Default=" << lambda     << "]" << endl;
		cout << "#   -mrf_i        load MRF"                                                                      << endl;
		cout << "#   -mrf_o        save MRF"                                                                      << endl;
		cout << "#   -pair_i       load list of residue pairs (one pair per line, index 0)"                       << endl;
		cout << "# ---------------------------------------------------------------------------------------------" << endl;
		cout << "#  Minimizer settings                                                                          " << endl;
		cout << "# ---------------------------------------------------------------------------------------------" << endl;
		cout << "#   -min_type     select: [lbgfs|cg|none]                        [Default=" << min_type   << "]" << endl;
		cout << "#   -max_iter     number of iterations                           [Default=" << max_iter   << "]" << endl;
		cout << "# ---------------------------------------------------------------------------------------------" << endl;
		if(arg.size() > 0){cout << eout.str();}
		exit(1);
	}
	else{
		cout << "# ---------------------------------------------------------------------------------------------" << endl;
		cout << "#                                GREMLIN_H (with entropy correction)                           " << endl;
		cout << "# ---------------------------------------------------------------------------------------------" << endl;
		cout << "#   -i           " << msa_i            << endl;
		cout << "#   -o           " << preds_out        << endl;
		cout << "# ---------------------------------------------------------------------------------------------" << endl;
		cout << "#   -only_v      " << only_v           << endl;
		cout << "#   -gap_cutoff  " << gap_cutoff       << endl;
		cout << "#   -alphabet    " << alphabet         << endl;
		cout << "#   -eff_cutoff  " << eff_cutoff       << endl;
		cout << "#   -lambda      " << lambda           << endl;
		if(!mrf_i.empty()){
			cout << "#   -mrf_i       " << mrf_i            << endl;}
		if(!mrf_o.empty()){
			cout << "#   -mrf_o       " << mrf_o            << endl;}
		if(!pair_i.empty()){
			cout << "#   -pair_i      " << pair_i           << endl;}
		cout << "# ---------------------------------------------------------------------------------------------" << endl;
		cout << "#   -min_type    " << min_type         << endl;
		cout << "#   -max_iter    " << max_iter         << endl;
		cout << "# ---------------------------------------------------------------------------------------------" << endl;
	}
}
```
