## Supplemental Code S1 for "Remote homology search with hidden Potts models": hmm_entropy.c

```
/* hmm_entropy.c */
#include "hmm_entropy.h"
#include "p7_config.h"
#include "dp_reference/p7_refmx.h"
#include "base/p7_profile.h"
#include "hmmer.h"
#include "easel.h"
#include "esl_alphabet.h"
#include "esl_arr2.h"
#include "esl_composition.h"
#include "esl_vectorops.h"
#include "potts.h"


double **G_Create(int NS);

double **Gx_Create(int NSx);

void G_Initialize(double **G, int NS);

void Gx_Initialize(double **Gx, int NSx);

void DLogNorm_NoNan(double *vec, int n);

int dp_IDX_standard(int k, int state);

int dp_IDX_special(int M, int state);


/* Function: hmm_entropy_Calculate()
 *
 * Synopsis: Calculate the posterior entropy of a path given a sequence
 *           and model, H ( \vec{\pi} | \vec{x} ).
 *
 * Purpose:  For a configured uniglocal profile HMM <gm> and precalculated
 *           forward matrix <fwd> which has been calculated using <gm> and an
 *           unaligned sequence of length L, return the posterior entropy of
 *           the path given the sequence and model, <ret_H>, in bits. 2^H
 *           yields the "perplexity", which is an estimate of the magnitude of
 *           the number of random q_samples needed to effectively characterize a
 *           probability distribution (see MacKay).
 *
 *           This function is based on the algorithm described in
 *           Hernando et al, 2003. The format of the function is  roughly
 *           based on p7_ReferenceForward().
 *
 *           I have yet to Tex up my handwritten notes on the math that
 *           underlies this function.
 *
 *
 * Args:     gm      : query profile w/ M match states and length L
 *           fwd     : precalculated P7_REFMX forward dp object (created
 *                    using p7_ReferenceForward() <gm> and a sequence
 *                    of length L, perhaps).
 *           ret_H   : RESULT: posterior entropy, in bits
 *           verbose : option to print intermediate variables to the terminal.
 *
 *
 */

int
hmm_entropy_Calculate(P7_PROFILE *gm, P7_REFMX *fwd, float *ret_H, int verbose)
{
	int          status;
	int          M            = gm->M;                   /* number of match states                                    */
	int          L            = gm->L;                   /* target sequence length                                    */
	int          i,k,s;                                  /* residue, node, and state indices                          */
	double     **G            = NULL;                    /* log P( \pi_{i-1} | \pi_i, \vec{x}_i ) for M, I, D states  */
	double     **g         	  = NULL;                    /* P( \pi_{i-1} | \pi_i, \vec{x}_i ) for M, I, D states      */
	double     **Gx           = NULL;						  /* log P( \pi_{i-1} | \pi_i, \vec{x}_i ) for special states  */
	double     **gx           = NULL;                    /* P( \pi_{i-1} | \pi_i, \vec{x}_i ) for special states      */
	P7_REFMX    *ent          = p7_refmx_Create(M, L);   /* dynamic programming object for posterior entropy          */
	float       *dpc, *dpp;                              /* pointers for stepping through ent->dp                     */
	float       *fwc, *fwp;                              /* pointers for stepping throw fwd->dp                       */
	int          idx_M        = 0;                       /* match state index for G, g, Gx, gx                        */
	int          idx_I        = 1;                       /* insert state index for G, g                               */
	int          idx_D        = 2;                       /* Delete state index for G, g, Gx, gx                       */
	int          idx_G        = 3;                       /* G state index for G, g                                    */
	int          idx_E        = 0;                       /* E state index for Gx, gx                                  */
	int          idx_C        = 1;                       /* C state index for Gx, gx                                  */
	int          NSG          = 3;                       /* number of standard states per node (MG, IG, DG)           */
	int          NSx          = 2;                       /* number of special states with non-zero entropy (E, C)     */
	float        mgv;                                    /* MG value for current cell row                             */
	float        dgv;                                    /* Pushed-ahead DG cell k+1 values                           */


	/* set dimensions for dp matrix */
	if ( (status = p7_refmx_GrowTo(ent, M, L)) != eslOK) goto ERROR;
	ent->M    = M;
	ent->L    = L;
	ent->type = p7R_UNSET;

	/* initialize dp matrix with all zero values                 */
	/* all i=0 entropies are 0.0, anyways, (initialization step) */
	p7_refmx_SetValues(ent, 0.0);

	/* create G array */
	G = G_Create(NSG);
	g = G_Create(NSG);

	/* create Gx array for special states G,C*/
	Gx = Gx_Create(NSx);
	gx = Gx_Create(NSx);

	/* Main DP recursion */
	for (i = 1; i <= L; i++) {
		if (verbose) fprintf(stdout, "i = %d\n", i);

		/* initialization for a new row */
		dpp = ent->dp[i-1];               /* previous row dpp is already set, and at k=0              */
		dpc = ent->dp[i];                 /* current DP row, skip k=0, start at k=1.                  */
		fwp = fwd->dp[i-1];               /* previous row of forward matrix, starting at k=0          */
		fwc = fwd->dp[i];                 /* current row of forward matrix, starting at k=0           */

		/* set entropies for all k=0 standard states to zero */
		for (s = 0; s < p7R_NSCELLS; s++){
			*dpc++ = 0.0;
		}
		/* now dpc points at k=1 */

		/* handle k = 1 specially                            */
      /* MG, DG will have zero entropy here => already set */

		/* initialize G matrix */
		G_Initialize(G, NSG);

		/* Calculate G values */
		/* only G_MI and G_MI matter */
		G[idx_I][idx_M] = P7P_TSC(gm, 1, p7P_MI) + fwp[dp_IDX_standard(1,p7R_MG)];
		G[idx_I][idx_I] = P7P_TSC(gm, 1, p7P_II) + fwp[dp_IDX_standard(1,p7R_IG)];

		/* normalize, keep in log space */
		DLogNorm_NoNan(G[idx_I], NSG+1);

		if (verbose) {
			fprintf(stdout, "\tk = 1\n");

			fprintf(stdout, "\tG_M1I1 (%d) = %f\n", i, G[idx_I][idx_M] );
			fprintf(stdout, "\tG_I1I1 (%d) = %f\n", i, G[idx_I][idx_I] );
			fprintf(stdout, "\n");
		}

		/* calculate g from G, move to linear space */
		esl_vec_DCopy(G[idx_I], NSG+1, g[idx_I]);
		esl_vec_DExp(g[idx_I], NSG+1);

		/* set entropies with pointer */
		/* Local match state ML1 */
		*dpc++ = 0.0;

		/* Global state MG1 */
		*dpc++ = 0.0;

		dpp += p7R_NSCELLS;	/* dpp advances to cells for node 1 */

		/* Local insert state IL1 */
		*dpc++ = 0.0;

		/* global insert state IG1                                */
		/*esl_vec_DEntropy() returns bits, we want nats (for now) */
		*dpc++ = (esl_vec_DEntropy(g[idx_I], NSG+1))/(1.44269504)
			      + (*(dpp+p7R_MG) * g[idx_I][idx_M])
					+ (*(dpp+p7R_IG) * g[idx_I][idx_I]);

		/* local delete state DL1 */
		*dpc++ = 0.0;

		/* global delete state DG1 */
		*dpc++ = 0.0;

		dgv = 0.0;
		mgv = 0.0;

		/* loop through intermediate nodes */
		for (k = 2; k < M; k++) {

			/* initialize G matrix */
			G_Initialize(G, NSG);

			/* calculate G_XM's */
			G[idx_M][idx_M] =  P7P_TSC(gm, k-1, p7P_MM) + fwp[dp_IDX_standard(k-1, p7R_MG)];
			G[idx_M][idx_I] =  P7P_TSC(gm, k-1, p7P_IM) + fwp[dp_IDX_standard(k-1, p7R_IG)];
			G[idx_M][idx_D] =  P7P_TSC(gm, k-1, p7P_DM) + fwp[dp_IDX_standard(k-1, p7R_DG)];
			G[idx_M][idx_G] =  P7P_TSC(gm, k-1, p7P_GM) + fwp[dp_IDX_special(M, p7R_G)];

			DLogNorm_NoNan(G[idx_M], NSG+1);

			/* calculate G_XI's */
			G[idx_I][idx_M] = P7P_TSC(gm, k, p7P_MI) + fwp[dp_IDX_standard(k, p7R_MG)];
			G[idx_I][idx_I] = P7P_TSC(gm, k, p7P_II) + fwp[dp_IDX_standard(k, p7R_IG)];

			DLogNorm_NoNan(G[idx_I], NSG+1);

			/* calculate G_XD's */
			G[idx_D][idx_M] = P7P_TSC(gm, k-1, p7P_DM) + fwc[dp_IDX_standard(k-1, p7R_MG)];
			G[idx_D][idx_D] = P7P_TSC(gm, k-1, p7P_DD) + fwc[dp_IDX_standard(k-1, p7R_DG)];

			DLogNorm_NoNan(G[idx_D], NSG+1);

			if (verbose) {
				fprintf(stdout, "\tk = %d\n", k);

				fprintf(stdout, "\tG_M%dM%d (%d) = %f\n", k-1, k, i, G[idx_M][idx_M] );
				fprintf(stdout, "\tG_I%dM%d (%d) = %f\n", k-1, k, i, G[idx_M][idx_I] );
				fprintf(stdout, "\tG_D%dM%d (%d) = %f\n", k-1, k, i, G[idx_M][idx_D] );
				fprintf(stdout, "\tG_GM%d (%d) = %f\n", k, i, G[idx_M][idx_G] );
				fprintf(stdout, "\n");

				fprintf(stdout, "\tG_M%dI%d (%d) = %f\n", k, k, i, G[idx_I][idx_M] );
				fprintf(stdout, "\tG_I%dI%d (%d) = %f\n", k, k, i, G[idx_I][idx_I] );
				fprintf(stdout, "\n");

				fprintf(stdout, "\tG_M%dD%d (%d) = %f\n", k-1, k, i, G[idx_D][idx_M] );
				fprintf(stdout, "\tG_D%dD%d (%d) = %f\n", k-1, k, i, G[idx_D][idx_D] );
				fprintf(stdout, "\n");
			}


			/* calculate g from G */
			esl_vec_DCopy(G[idx_M], NSG+1, g[idx_M]);
			esl_vec_DExp(g[idx_M], NSG+1);

			esl_vec_DCopy(G[idx_I], NSG+1, g[idx_I]);
			esl_vec_DExp(g[idx_I], NSG+1);

			esl_vec_DCopy(G[idx_D], NSG+1, g[idx_D]);
			esl_vec_DExp(g[idx_D], NSG+1);


			/* set entropy values */

			/* pre-set d-state entropy using H_{k-1}(i)'s, "deferred storage trick" */
			dgv =  esl_vec_DEntropy(g[idx_D], NSG+1)/(1.44269504)
				 +  (mgv * g[idx_D][idx_M])
				 +  (dgv * g[idx_D][idx_D]);

			/* state ML */
			*dpc++ = 0.0;

			/* state MG */
			mgv = *dpc++ = (esl_vec_DEntropy(g[idx_M], NSG+1))/(1.44269504)
				          + (*(dpp+p7R_MG) * g[idx_M][idx_M])
				          + (*(dpp+p7R_IG) * g[idx_M][idx_I])
				          + (*(dpp+p7R_DG) * g[idx_M][idx_D]);

			dpp += p7R_NSCELLS;	/* dpp advances to cells for states k   */

			/* state IL */
			*dpc++ = 0.0;

			/* state IG */
			*dpc++ = (esl_vec_DEntropy(g[idx_I], NSG+1))/(1.44269504)
				 + (*(dpp+p7R_MG) * g[idx_I][idx_M])
				 + (*(dpp+p7R_IG) * g[idx_I][idx_D]);


			/* state DL */
			*dpc++ = 0.0;


			/* Delete state, deferred storage trick */
			/* state DG */
			*dpc++ = dgv;

		}

		/* handle k = M specially */

		/* initialize G matrix */
		G_Initialize(G, NSG);

		/* calculate G_XM's */
		G[idx_M][idx_M] = P7P_TSC(gm, M-1, p7P_MM) + fwp[dp_IDX_standard(M-1, p7R_MG)];
		G[idx_M][idx_I] = P7P_TSC(gm, M-1, p7P_IM) + fwp[dp_IDX_standard(M-1, p7R_IG)];
		G[idx_M][idx_D] = P7P_TSC(gm, M-1, p7P_DM) + fwp[dp_IDX_standard(M-1, p7R_DG)];
		G[idx_M][idx_G] = P7P_TSC(gm, M-1, p7P_GM) + fwp[dp_IDX_special(M, p7R_G)];

		DLogNorm_NoNan(G[idx_M], NSG+1);

		/* state IM doesn't exist => ignore it's G's*/

		/* calculate G_XD's */
		G[idx_D][idx_M] = P7P_TSC(gm, M-1, p7P_MD) + fwc[dp_IDX_standard(M-1, p7R_MG)];
		G[idx_D][idx_D] = P7P_TSC(gm, M-1, p7P_DD) + fwc[dp_IDX_standard(M-1, p7R_DG)];

		DLogNorm_NoNan(G[idx_D], NSG+1);

		if (verbose) {
			fprintf(stdout, "\tk = M = %d\n", M);

			fprintf(stdout, "\tG_M%dM%d (%d) = %f\n", M-1, M, i, G[idx_M][idx_M] );
			fprintf(stdout, "\tG_I%dM%d (%d) = %f\n", M-1, M, i, G[idx_M][idx_I] );
			fprintf(stdout, "\tG_D%dM%d (%d) = %f\n", M-1, M, i, G[idx_M][idx_D] );
			fprintf(stdout, "\tG_GM%d (%d) = %f\n", M, i, G[idx_M][idx_G] );

			fprintf(stdout, "\n");

			fprintf(stdout, "\tG_M%dD%d (%d) = %f\n", M-1, M, i, G[idx_D][idx_M] );
			fprintf(stdout, "\tG_D%dD%d (%d) = %f\n", M-1, M, i, G[idx_D][idx_D] );
			fprintf(stdout, "\n");
		}

		/* calculate g from G */
		esl_vec_DCopy(G[idx_M], NSG+1, g[idx_M]);
		esl_vec_DExp(g[idx_M], NSG+1);

		esl_vec_DCopy(G[idx_D], NSG+1, g[idx_D]);
		esl_vec_DExp(g[idx_D], NSG+1);

		/* pre-set d-state entropy using H_k-1(i)'s , current node g array */
		dgv =  esl_vec_DEntropy(g[idx_D], NSG+1)/(1.44269504)
			 +  (mgv * g[idx_D][idx_M])
			 +  (dgv * g[idx_D][idx_D]);


		/* state MLM */
		*dpc++ = 0.0;

		/* state MGM */
		mgv = *dpc++ = (esl_vec_DEntropy(g[idx_M], NSG+1))/(1.44269504)
				       + (*(dpp+p7R_MG) * g[idx_M][idx_M])
				       + (*(dpp+p7R_IG) * g[idx_M][idx_I])
				       + (*(dpp+p7R_DG) * g[idx_M][idx_D]);


		/* precalculate DG entropy before advancing dpp */
		dpp  += p7R_NSCELLS;

		/* state ILM */
		*dpc++ = 0.0;

		/* state IGM */
		*dpc++ = 0.0;

		/* state DLM */
		*dpc++ = 0.0;

		/* state DGM */
		*dpc++ = dgv;

		/* handle "special" states GCC and GEC */

		Gx_Initialize(Gx, NSx);

		/* calculate G_XE */
		Gx[idx_E][idx_M] = fwc[dp_IDX_special(M, p7R_MG)];
		Gx[idx_E][idx_D-1] = fwc[dp_IDX_special(M, p7R_DG)];


		DLogNorm_NoNan(Gx[idx_E], NSx);

		Gx[idx_C][idx_E] = fwc[dp_IDX_special(M, p7R_E)];
		Gx[idx_C][idx_C] = fwp[dp_IDX_special(M, p7R_C)] + gm->xsc[p7P_C][p7P_LOOP];

		DLogNorm_NoNan(Gx[idx_C], NSx);

		if (verbose) {
			fprintf(stdout, "\tSpecial States:\n");
			fprintf(stdout, "\tG_M%dE(%d) = %f\n", M, i, Gx[idx_E][idx_M] );
			fprintf(stdout, "\tG_D%dE(%d) = %f\n", M, i, Gx[idx_E][idx_D-1] );
			fprintf(stdout, "\n");

			fprintf(stdout, "\tG_CE(%d) = %f\n", i, Gx[idx_C][idx_E] );
			fprintf(stdout, "\tG_CC(%d) = %f\n", i, Gx[idx_C][idx_C] );
			fprintf(stdout, "\n");
		}

		/* move to linear space */
		esl_vec_DCopy(Gx[idx_E], NSx, gx[idx_E]);
		esl_vec_DExp(gx[idx_E], NSx);

		esl_vec_DCopy(Gx[idx_C], NSx, gx[idx_C]);
		esl_vec_DExp(gx[idx_C], NSx);

		dpp  += p7R_NSCELLS;
		/* row i is finished, dpc[] is positioned on first special state E */

		/* Set special state entropies */
		dpc[p7R_E]  =  (esl_vec_DEntropy(gx[idx_E], NSx))/(1.44269504)
			         +  (gx[idx_E][idx_M] * mgv)
						+  (gx[idx_E][idx_M] * mgv);
		dpc[p7R_N]  =  0.0;
		dpc[p7R_J]  =  0.0;
		dpc[p7R_B]  =  0.0;
		dpc[p7R_C]  =  (esl_vec_DEntropy(gx[idx_C], NSx))/(1.44269504)
			         +  (gx[idx_C][idx_E] * dpc[p7R_E])
						+  (gx[idx_C][idx_C] * dpp[p7R_C]);
		dpc[p7R_L]  = 	0.0;
		dpc[p7R_G]  =  0.0;
		dpc[p7R_JJ] =  0.0;
		dpc[p7R_CC] =  0.0;

	}

	/* H_T (L+1) = H_C (L) = H ( \vec{pi} | \vec{x} ) */
	/* convert entropy fron nats to bits */
	*ret_H = 1.44269504 * dpc[p7R_C];

	if (verbose) {

		/* dump entropy dp matrix to terminal */
		p7_refmx_DumpWindow(stdout, ent, 0, L, 0, M);
	}

	if (verbose) fprintf(stdout, "Total entropy: %f\n", 1.44269504*dpc[p7R_C]);

	p7_refmx_Destroy(ent);
        esl_arr2_Destroy((void **) G, NSG);
        esl_arr2_Destroy((void **) g, NSG);
        esl_arr2_Destroy((void **) Gx, NSx);
        esl_arr2_Destroy((void **) gx, NSx);

	return eslOK;

	ERROR:
		return status;
}

int
hmm_entropy_CalculateHernando(P7_PROFILE *gm, P7_REFMX *fwd, float *ret_H, int opt_L, int verbose)
{
	int          status;
	int          M            = gm->M;                   /* number of match states                                    */
	int          L            = gm->L;                   /* target sequence length                                    */
	int          i,k,s;                                  /* residue, node, and state indices                          */
	double     **G            = NULL;                    /* log P( \pi_{i-1} | \pi_i, \vec{x}_i ) for M, I, D states  */
	double     **g         	  = NULL;                    /* P( \pi_{i-1} | \pi_i, \vec{x}_i ) for M, I, D states      */
	double     **Gx           = NULL;						  /* log P( \pi_{i-1} | \pi_i, \vec{x}_i ) for special states  */
	double     **gx           = NULL;                    /* P( \pi_{i-1} | \pi_i, \vec{x}_i ) for special states      */
	P7_REFMX    *ent          = p7_refmx_Create(M, L);   /* dynamic programming object for posterior entropy          */
	float       *dpc, *dpp;                              /* pointers for stepping through ent->dp                     */
	float       *fwc, *fwp;                              /* pointers for stepping throw fwd->dp                       */
	int          idx_M        = 0;                       /* match state index for G, g, Gx, gx                        */
	int          idx_I        = 1;                       /* insert state index for G, g                               */
	int          idx_D        = 2;                       /* Delete state index for G, g, Gx, gx                       */
	int          idx_G        = 3;                       /* G state index for G, g                                    */
	int          idx_E        = 0;                       /* E state index for Gx, gx                                  */
	int          idx_C        = 1;                       /* C state index for Gx, gx                                  */
	int          NSG          = 3;                       /* number of standard states per node (MG, IG, DG)           */
	int          NSx          = 2;                       /* number of special states with non-zero entropy (E, C)     */
	float        mgv;                                    /* MG value for current cell row                             */
	float        dgv;                                    /* Pushed-ahead DG cell k+1 values                           */

	if (opt_L > 0) L = opt_L;

	/* set dimensions for dp matrix */
	if ( (status = p7_refmx_GrowTo(ent, M, L)) != eslOK) goto ERROR;
	ent->M    = M;
	ent->L    = L;
	ent->type = p7R_UNSET;

	/* initialize dp matrix with all zero values                 */
	/* all i=0 entropies are 0.0, anyways, (initialization step) */
	p7_refmx_SetValues(ent, 0.0);

	/* create G array */
	G = G_Create(NSG);
	g = G_Create(NSG);

	/* create Gx array for special states G,C*/
	Gx = Gx_Create(NSx);
	gx = Gx_Create(NSx);

	/* Main DP recursion */
	for (i = 1; i <= L; i++) {
		if (verbose) fprintf(stdout, "i = %d\n", i);

		/* initialization for a new row */
		dpp = ent->dp[i-1];               /* previous row dpp is already set, and at k=0              */
		dpc = ent->dp[i];                 /* current DP row, skip k=0, start at k=1.                  */
		fwp = fwd->dp[i-1];               /* previous row of forward matrix, starting at k=0          */
		fwc = fwd->dp[i];                 /* current row of forward matrix, starting at k=0           */

		/* set entropies for all k=0 standard states to zero */
		for (s = 0; s < p7R_NSCELLS; s++){
			*dpc++ = 0.0;
		}
		/* now dpc points at k=1 */

		/* handle k = 1 specially                            */
      /* MG, DG will have zero entropy here => already set */

		/* initialize G matrix */
		G_Initialize(G, NSG);

		/* Calculate G values */
		/* only G_MI and G_MI matter */
		G[idx_I][idx_M] = P7P_TSC(gm, 1, p7P_MI) + fwp[dp_IDX_standard(1,p7R_MG)];
		G[idx_I][idx_I] = P7P_TSC(gm, 1, p7P_II) + fwp[dp_IDX_standard(1,p7R_IG)];


		/* normalize, keep in log space */
		DLogNorm_NoNan(G[idx_I], NSG+1);

		if (verbose) {
			fprintf(stdout, "\tk = 1\n");

			fprintf(stdout, "\tG_M1I1 (%d) = %f\n", i, G[idx_I][idx_M] );
			fprintf(stdout, "\tG_I1I1 (%d) = %f\n", i, G[idx_I][idx_I] );
			fprintf(stdout, "\n");
		}

		/* calculate g from G, move to linear space */
		esl_vec_DCopy(G[idx_I], NSG+1, g[idx_I]);
		esl_vec_DExp(g[idx_I], NSG+1);

		/* set entropies with pointer */
		/* Local match state ML1 */
		*dpc++ = 0.0;

		/* Global state MG1 */
		*dpc++ = 0.0;

		dpp += p7R_NSCELLS;	/* dpp advances to cells for node 1 */

		/* Local insert state IL1 */
		*dpc++ = 0.0;

		/* global insert state IG1                                */
		/*esl_vec_DEntropy() returns bits, we want nats (for now) */
		*dpc++ = (esl_vec_DEntropy(g[idx_I], NSG+1))/(1.44269504)
			      + (*(dpp+p7R_MG) * g[idx_I][idx_M])
					+ (*(dpp+p7R_IG) * g[idx_I][idx_I]);

		/* local delete state DL1 */
		*dpc++ = 0.0;

		/* global delete state DG1 */
		*dpc++ = 0.0;

		dgv = 0.0;
		mgv = 0.0;

		/* loop through intermediate nodes */
		for (k = 2; k < M; k++) {

			/* initialize G matrix */
			G_Initialize(G, NSG);

			/* calculate G_XM's */
			G[idx_M][idx_M] =  P7P_TSC(gm, k-1, p7P_MM) + fwp[dp_IDX_standard(k-1, p7R_MG)];
			G[idx_M][idx_I] =  P7P_TSC(gm, k-1, p7P_IM) + fwp[dp_IDX_standard(k-1, p7R_IG)];
			G[idx_M][idx_D] =  P7P_TSC(gm, k-1, p7P_DM) + fwp[dp_IDX_standard(k-1, p7R_DG)];
			G[idx_M][idx_G] =  P7P_TSC(gm, k-1, p7P_GM) + fwp[dp_IDX_special(M, p7R_G)];

			DLogNorm_NoNan(G[idx_M], NSG+1);

			/* calculate G_XI's */
			G[idx_I][idx_M] = P7P_TSC(gm, k, p7P_MI) + fwp[dp_IDX_standard(k, p7R_MG)];
			G[idx_I][idx_I] = P7P_TSC(gm, k, p7P_II) + fwp[dp_IDX_standard(k, p7R_IG)];

			DLogNorm_NoNan(G[idx_I], NSG+1);

			/* calculate G_XD's */
			//G[idx_D][idx_M] = P7P_TSC(gm, k-1, p7P_DM) + fwc[dp_IDX_standard(k-1, p7R_MG)];
			G[idx_D][idx_M] = P7P_TSC(gm, k-1, p7P_MD) + fwc[dp_IDX_standard(k-1, p7R_MG)];
			G[idx_D][idx_D] = P7P_TSC(gm, k-1, p7P_DD) + fwc[dp_IDX_standard(k-1, p7R_DG)];


			DLogNorm_NoNan(G[idx_D], NSG+1);

			if (verbose) {
				fprintf(stdout, "\tk = %d\n", k);

				fprintf(stdout, "\tG_M%dM%d (%d) = %f\n", k-1, k, i, G[idx_M][idx_M] );
				fprintf(stdout, "\tG_I%dM%d (%d) = %f\n", k-1, k, i, G[idx_M][idx_I] );
				fprintf(stdout, "\tG_D%dM%d (%d) = %f\n", k-1, k, i, G[idx_M][idx_D] );
				fprintf(stdout, "\tG_GM%d (%d) = %f\n", k, i, G[idx_M][idx_G] );
				fprintf(stdout, "\n");

				fprintf(stdout, "\tG_M%dI%d (%d) = %f\n", k, k, i, G[idx_I][idx_M] );
				fprintf(stdout, "\tG_I%dI%d (%d) = %f\n", k, k, i, G[idx_I][idx_I] );
				fprintf(stdout, "\n");

				fprintf(stdout, "\tG_M%dD%d (%d) = %f\n", k-1, k, i, G[idx_D][idx_M] );
				fprintf(stdout, "\tG_D%dD%d (%d) = %f\n", k-1, k, i, G[idx_D][idx_D] );
				fprintf(stdout, "\n");
			}


			/* calculate g from G */
			esl_vec_DCopy(G[idx_M], NSG+1, g[idx_M]);
			esl_vec_DExp(g[idx_M], NSG+1);

			esl_vec_DCopy(G[idx_I], NSG+1, g[idx_I]);
			esl_vec_DExp(g[idx_I], NSG+1);

			esl_vec_DCopy(G[idx_D], NSG+1, g[idx_D]);
			esl_vec_DExp(g[idx_D], NSG+1);


			/* set entropy values */

			/* pre-set d-state entropy using H_{k-1}(i)'s, "deferred storage trick" */
			dgv =  esl_vec_DEntropy(g[idx_D], NSG+1)/(1.44269504)
				 +  (mgv * g[idx_D][idx_M])
				 +  (dgv * g[idx_D][idx_D]);

			/* state ML */
			*dpc++ = 0.0;

			/* state MG */
			mgv = *dpc++ = (esl_vec_DEntropy(g[idx_M], NSG+1))/(1.44269504)
				          + (*(dpp+p7R_MG) * g[idx_M][idx_M])
				          + (*(dpp+p7R_IG) * g[idx_M][idx_I])
				          + (*(dpp+p7R_DG) * g[idx_M][idx_D]);

			dpp += p7R_NSCELLS;	/* dpp advances to cells for states k   */

			/* state IL */
			*dpc++ = 0.0;

			/* state IG */
			*dpc++ = (esl_vec_DEntropy(g[idx_I], NSG+1))/(1.44269504)
				 + (*(dpp+p7R_MG) * g[idx_I][idx_M])
				 //+ (*(dpp+p7R_IG) * g[idx_I][idx_D]);
				 + (*(dpp+p7R_IG) * g[idx_I][idx_I]);


			/* state DL */
			*dpc++ = 0.0;


			/* Delete state, deferred storage trick */
			/* state DG */
			*dpc++ = dgv;

		}

		/* handle k = M specially */

		/* initialize G matrix */
		G_Initialize(G, NSG);

		/* calculate G_XM's */
		G[idx_M][idx_M] = P7P_TSC(gm, M-1, p7P_MM) + fwp[dp_IDX_standard(M-1, p7R_MG)];
		G[idx_M][idx_I] = P7P_TSC(gm, M-1, p7P_IM) + fwp[dp_IDX_standard(M-1, p7R_IG)];
		G[idx_M][idx_D] = P7P_TSC(gm, M-1, p7P_DM) + fwp[dp_IDX_standard(M-1, p7R_DG)];
		G[idx_M][idx_G] = P7P_TSC(gm, M-1, p7P_GM) + fwp[dp_IDX_special(M, p7R_G)];

		DLogNorm_NoNan(G[idx_M], NSG+1);

		/* state IM doesn't exist => ignore it's G's*/

		/* calculate G_XD's */
		G[idx_D][idx_M] = P7P_TSC(gm, M-1, p7P_MD) + fwc[dp_IDX_standard(M-1, p7R_MG)];
		G[idx_D][idx_D] = P7P_TSC(gm, M-1, p7P_DD) + fwc[dp_IDX_standard(M-1, p7R_DG)];

		DLogNorm_NoNan(G[idx_D], NSG+1);

		if (verbose) {
			fprintf(stdout, "\tk = M = %d\n", M);

			fprintf(stdout, "\tG_M%dM%d (%d) = %f\n", M-1, M, i, G[idx_M][idx_M] );
			fprintf(stdout, "\tG_I%dM%d (%d) = %f\n", M-1, M, i, G[idx_M][idx_I] );
			fprintf(stdout, "\tG_D%dM%d (%d) = %f\n", M-1, M, i, G[idx_M][idx_D] );
			fprintf(stdout, "\tG_GM%d (%d) = %f\n", M, i, G[idx_M][idx_G] );

			fprintf(stdout, "\n");

			fprintf(stdout, "\tG_M%dD%d (%d) = %f\n", M-1, M, i, G[idx_D][idx_M] );
			fprintf(stdout, "\tG_D%dD%d (%d) = %f\n", M-1, M, i, G[idx_D][idx_D] );
			fprintf(stdout, "\n");
		}

		/* calculate g from G */
		esl_vec_DCopy(G[idx_M], NSG+1, g[idx_M]);
		esl_vec_DExp(g[idx_M], NSG+1);

		esl_vec_DCopy(G[idx_D], NSG+1, g[idx_D]);
		esl_vec_DExp(g[idx_D], NSG+1);

		/* pre-set d-state entropy using H_k-1(i)'s , current node g array */
		dgv =  esl_vec_DEntropy(g[idx_D], NSG+1)/(1.44269504)
			 +  (mgv * g[idx_D][idx_M])
			 +  (dgv * g[idx_D][idx_D]);


		/* state MLM */
		*dpc++ = 0.0;

		/* state MGM */
		mgv = *dpc++ = (esl_vec_DEntropy(g[idx_M], NSG+1))/(1.44269504)
				       + (*(dpp+p7R_MG) * g[idx_M][idx_M])
				       + (*(dpp+p7R_IG) * g[idx_M][idx_I])
				       + (*(dpp+p7R_DG) * g[idx_M][idx_D]);


		/* precalculate DG entropy before advancing dpp */
		dpp  += p7R_NSCELLS;

		/* state ILM */
		*dpc++ = 0.0;

		/* state IGM */
		*dpc++ = 0.0;

		/* state DLM */
		*dpc++ = 0.0;

		/* state DGM */
		*dpc++ = dgv;

		/* handle "special" states GCC and GEC */

		Gx_Initialize(Gx, NSx);

		/* calculate G_XE */
		//Gx[idx_E][idx_M] = fwc[dp_IDX_special(M, p7R_MG)];
		//Gx[idx_E][idx_D-1] = fwc[dp_IDX_special(M, p7R_DG)];

		Gx[idx_E][idx_M] = fwc[dp_IDX_standard(M, p7R_MG)];
		Gx[idx_E][idx_D-1] = fwc[dp_IDX_standard(M, p7R_DG)];


		DLogNorm_NoNan(Gx[idx_E], NSx);

		Gx[idx_C][idx_E] = fwc[dp_IDX_special(M, p7R_E)];
		Gx[idx_C][idx_C] = fwp[dp_IDX_special(M, p7R_C)] + gm->xsc[p7P_C][p7P_LOOP];

		DLogNorm_NoNan(Gx[idx_C], NSx);

		if (verbose) {
			fprintf(stdout, "\tSpecial States:\n");
			fprintf(stdout, "\tG_M%dE(%d) = %f\n", M, i, Gx[idx_E][idx_M] );
			fprintf(stdout, "\tG_D%dE(%d) = %f\n", M, i, Gx[idx_E][idx_D-1] );
			fprintf(stdout, "\n");

			fprintf(stdout, "\tG_CE(%d) = %f\n", i, Gx[idx_C][idx_E] );
			fprintf(stdout, "\tG_CC(%d) = %f\n", i, Gx[idx_C][idx_C] );
			fprintf(stdout, "\n");
		}

		/* move to linear space */
		esl_vec_DCopy(Gx[idx_E], NSx, gx[idx_E]);
		esl_vec_DExp(gx[idx_E], NSx);

		esl_vec_DCopy(Gx[idx_C], NSx, gx[idx_C]);
		esl_vec_DExp(gx[idx_C], NSx);

		dpp  += p7R_NSCELLS;
		/* row i is finished, dpc[] is positioned on first special state E */

		/* Set special state entropies */
		dpc[p7R_E]  =  (esl_vec_DEntropy(gx[idx_E], NSx))/(1.44269504)
			         +  (gx[idx_E][idx_M] * mgv)
						+  (gx[idx_E][idx_D-1] * dgv);
		dpc[p7R_N]  =  0.0;
		dpc[p7R_J]  =  0.0;
		dpc[p7R_B]  =  0.0;
		dpc[p7R_C]  =  (esl_vec_DEntropy(gx[idx_C], NSx))/(1.44269504)
			         +  (gx[idx_C][idx_E] * dpc[p7R_E])
						+  (gx[idx_C][idx_C] * dpp[p7R_C]);
		dpc[p7R_L]  = 	0.0;
		dpc[p7R_G]  =  0.0;
		dpc[p7R_JJ] =  0.0;
		dpc[p7R_CC] =  0.0;

	}

	/* H_T (L+1) = H_C (L) = H ( \vec{pi} | \vec{x} ) */
	/* convert entropy fron nats to bits */
	*ret_H = 1.44269504 * dpc[p7R_C];

	if (verbose) {

		/* dump entropy dp matrix to terminal */
		p7_refmx_DumpWindow(stdout, ent, 0, L, 0, M);
	}

	if (verbose) fprintf(stdout, "Total entropy: %f\n", 1.44269504*dpc[p7R_C]);

        /* clean up and return */
	p7_refmx_Destroy(ent);
        esl_arr2_Destroy((void **) G, NSG);
        esl_arr2_Destroy((void **) g, NSG);
        esl_arr2_Destroy((void **) Gx, NSx);
        esl_arr2_Destroy((void **) gx, NSx);


	return eslOK;

	ERROR:
		return status;
}


/* Function: hmm_entropy_CalculateSampling()
 */
int
hmm_entropy_CalculateSampling(P7_PROFILE *gm, P7_REFMX *fwd, ESL_RANDOMNESS *rng, int nsamp, float *ret_H, int verbose)
{
	ESL_KEYHASH      *h          = NULL;
	P7_TRACE         *tr         = p7_trace_Create();    /* for stochastic traceback     */
	float            *wrk        = NULL;
	double           *q_samp;
	int               nalloc;
	int               n,m;
	int               idx;
	int               status;


	/* create keyhash object */
	h = esl_keyhash_Create();
	if (verbose) fprintf(stdout, "h->kalloc: %d\n", h->kalloc);
	/* allocate memory for sample array */
	nalloc = h->kalloc;
	ESL_ALLOC(q_samp, sizeof(double) * nalloc);
	esl_vec_DSet(q_samp, nalloc, 0.0);


	for (n=0; n < nsamp; n++)
	{

		/* generate path */
		p7_reference_trace_Stochastic(rng, &wrk, gm, fwd, tr);
		/* store state string in keyhash (if it isn't there already) */
		esl_keyhash_Store(h, tr->st, tr->N, &idx);


		/* check if we need to reallocate memory in sample array */
		if (idx == nalloc) {
			ESL_REALLOC(q_samp, sizeof(double) * 2 * nalloc);

			/* initialize new array elements */
			for (m=nalloc; m < nalloc * 2; m++) {
				q_samp[m] = 0.0;
			}

			nalloc *= 2;

		}

		/* note which path we sampled */
		q_samp[idx] += 1.0;

		/* prepare for next iteration */
		p7_trace_Reuse(tr);
	}

	/* print out q_samp */
	//for (n = 0; n < nalloc; n++){
	//	fprintf(stdout, "key: %d, q_samp[key]: %.1f\n", n, q_samp[n]);
	//}
	/* normalize probablities and calculate entropy */
	esl_vec_DNorm(q_samp, nalloc);

	/* print out q_samp */
	//for (n = 0; n < nalloc; n++){
	//	fprintf(stdout, "key: %d, q_samp[key]: %.4f\n", n, q_samp[n]);
	//}


	*ret_H = esl_vec_DEntropy(q_samp, nalloc);
	if (verbose) {
		fprintf(stdout, "size of q_samp: %lu\n", sizeof(q_samp));
		fprintf(stdout, "n_elements of q_samp: %lu\n", sizeof(q_samp) / sizeof(int));
		fprintf(stdout, "h->kalloc: %d\n", h->kalloc);
		fprintf(stdout, "nalloc: %d\n", nalloc);
		fprintf(stdout, "h->nkeys: %d\n", h->nkeys);
	}

	free(q_samp);
	free(wrk);
	p7_trace_Destroy(tr);
	esl_keyhash_Destroy(h);

	return eslOK;

	ERROR:
		return status;

}


/* Function: G_Create()
 * Synopsis: Allocate space for a (NSG X NSG+1) element
 * 			 log probability matrix, G
 *
 * Purpose:  Create matrix for G_L (i) for one sequence
 *           position i, node k. Elements correspond to
 *           log P (\pi_{i=1} | \pi_i, \vec{x}_i )
 *
 *    		 Gx is used in calculating intermediate posterior
 *           entropies of a profile HMM path distribution
 *           H ( \vec{\pi} | \vec{x} ) involving "special"
 *           states with non-zero intermediate entropy
 *           (E and C for a glocal profile hmm).
 *           See Hernando et al, 2003
 *
 *           rows: current state (\pi_i)
 *           cols: previous state (\pi_i-1)
 *
 *				 Last index corresnponds to "G" HMM state, (note
 *				 second use of letter G, sorry) handle sglocal
 *				 handles glocal wing-fold entry to intermediate
 *				 match states.
 *
 */

double
**G_Create(int NS) {
	double **ret_G   = NULL;
	int      i;
	int      status;

	/* first level allocation */
	ESL_ALLOC(ret_G,   NS * sizeof(double *));

	/* second level allocation */
	for (i=0; i<NS; i++) {
		ESL_ALLOC(ret_G[i],   (NS+1) * sizeof(double));
	}

	G_Initialize(ret_G, NS);

	return ret_G;

	ERROR:
		fprintf(stdout, "ERROR in G_Create!\n");
		return NULL;
}

/* Function: Gx_Create()
 * Synopsis: Allocate space for a (NSx X NSx) element
 * 			 log probability matrix, Gx,
 *
 * Purpose:  Create matrix for G_L (i) for one sequence
 *           position i, node k. Elements correspond to
 *           log P (\pi_{i=1} | \pi_i, \vec{x}_i )
 *
 *    		 Gx is used in calculating intermediate posterior
 *           entropies of a profile HMM path distribution
 *           H ( \vec{\pi} | \vec{x} ) involving "special"
 *           states with non-zero intermediate entropy
 *           (E and C for a glocal profile hmm).
 *           See Hernando et al, 2003
 *
 *           rows: current state (\pi_i)
 *           cols: previous state (\pi_i-1)
 *
 *			    NSx: number of special states w/ non-zero
 *			    intermediate entropies (2, for glocal profile
 *			    hmm's).
 *
 */


double
**Gx_Create(int NSx) {
	double **ret_Gx   = NULL;
	int      i;
	int      status;

	/* first level allocation */
	ESL_ALLOC(ret_Gx,   NSx * sizeof(double *));

	/* second level allocation */
	for (i=0; i<NSx; i++) {
		ESL_ALLOC(ret_Gx[i],   (NSx) * sizeof(double));
	}

	Gx_Initialize(ret_Gx, NSx);

	return ret_Gx;

	ERROR:
		fprintf(stdout, "ERROR in Gx_Create!\n");
		return NULL;
}


/* Function: G_Initialize()
 * Synopsis: Set all elements of <G> to -eslINFINITY
 *
 *
 * Purpose:  For reusing or creating a G matrix,
 *           it's often convenient to set all the elements
 *           to log(0) = -infinity
 *
 *
 *			    NS: number of "standard" state with non-zero
 *			    intermediate entropies (2, for glocal profile
 *			    hmm's).
 *
 */

void
G_Initialize(double **G, int NS) {
	int i;
	for (i=0; i<NS; i++) {
		esl_vec_DSet(G[i], NS+1, -eslINFINITY);
	}
}


/* Function: Gx_Initialize()
 * Synopsis: Set all elements of G to -eslINFINITY
 *
 *
 * Purpose:  For reusing or creating a Gx matrix,
 *           it's often convenient to set all the elements
 *           to log(0) = -infinity
 *
 *
 *			    NSx: number of "special' states with non-zero
 *			    intermediate entropies (2, for glocal profile
 *			    hmm's).
 *
 */


void
Gx_Initialize(double **Gx, int NSx) {
	int i;
	for (i=0; i<NSx; i++) {
		esl_vec_DSet(Gx[i], NSx, -eslINFINITY);
	}
}

/* Function: DLogNorm_NoNan()
 *
 * Synopsis: Normalize a log-p vector, keep in log space.
 *
 * Purpose:  Normalize a log probability vector <vec> of length
 *           n such that the exponential of all of the elements
 *           sum to 1.
 *
 *           Unlike esl_vec_DLogNorm(), this function DOES NOT
 *           convert the log-p vector to a p-vector
 *
 *           Note: If all elements of <vec> are -eslINFINITY,
 *           <vec> will be unmodified.
 *
 * Returns: (void); <vec> is changed in place.
 *
 */

void
DLogNorm_NoNan(double *vec, int n) {
	double denom;

	if (esl_vec_DMax(vec, n) ==  -eslINFINITY) {
		esl_vec_DSet(vec, n, -eslINFINITY);
	}
	else {
		denom = esl_vec_DLogSum(vec, n);
		esl_vec_DIncrement(vec, n, -1.*denom);
	}
}

/* Function: dp_IDX_standard()
 *
 * Synopsis: Get an index corresponding to a certain "standard" state
 *           type and node for a P7_REFMX dp array.
 *
 * Purpose:  For a specific dp[i] of the P7_REFMX dynamic programming
 *           matrix, states corresponding to different nodes are lumped
 *           into a single 1-D array. This function maps node numbers
 *           (<k> =0,..,M) and state types (<state> = p7R_ML, p7R_MG,
 *           p7R_IL, p7R_IG, p7R_DL, or p7R_MG) to the corresponding
 *           1-D array index.
 *
 *           See p7_refmx.h for more info on the layout of the dp matrix.
 *
 */

int dp_IDX_standard(int k, int state) {
	return (k*p7R_NSCELLS) + state;
}

/* Function: dp_IDX_special()
 *
 * Synopsis: Get an index corresponding to a certain "special" state
 *           type for a P7_REFMX dp array
 *
 * Purpose:  For a specific row dp[i] of the P7_REFMX dynamic programming
 *           matrix, special states are lumped into a single 1-D array
 *           along with all of the standard states corresponding to the
 *           M nodes.
 *           Possible values for <state>: p7R_E, p7R_N, p7R_J, p7R_B,
 *           p7R_L, p7R_G, p7R_C, p7R_JJ, p7R_CC
 *
 *
 *           See p7_refmx.h for more info on the layout of the dp matrix.
 *
 */


int dp_IDX_special(int M, int state) {
	return ((M+1)*p7R_NSCELLS) + state;
}


/* Unit tests go here */
#ifdef HMMENTROPY_TESTDRIVE

/* single path unit test */
static void
utest_singlepath_entropy(ESL_RANDOMNESS *rng, int alphatype, int M, int verbose)
{
   char          msg[] = "hmm entropy singlepath unit test failed";
	ESL_ALPHABET *abc   = esl_alphabet_Create(alphatype);
	P7_BG        *bg    = p7_bg_Create(abc);
	P7_HMM       *hmm   = NULL;
	P7_PROFILE   *gm    = p7_profile_Create(M, abc);
	ESL_SQ       *sq    = esl_sq_CreateDigital(abc);
	P7_TRACE     *tr    = p7_trace_Create();
	P7_REFMX     *fwd   = NULL;
	float         tol   = 0.0001;
	float         fsc;
	float         H;

	if (verbose) fprintf(stdout, "\thello world! In single path test.\n");

	/* Create a profile that has only a single possible path (including
	 * emissions) thru it; requires configuring in uniglocal mode w/ L=0
	 */
	if ( p7_modelsample_SinglePathed(rng, M, abc, &hmm) != eslOK) esl_fatal(msg);
	if ( p7_profile_ConfigUniglocal(gm, hmm, bg, 0)     != eslOK) esl_fatal(msg);

	//if (verbose) p7_profile_Dump(stdout, gm);

	/* Sample that sequence and path */
	if ( p7_ProfileEmit(rng, hmm, gm, bg, sq, tr)    != eslOK) esl_fatal(msg);
	if (verbose) p7_trace_Dump(stdout, tr);

	/* run the forward algorithm */
	if ( (fwd = p7_refmx_Create(M, sq->n))            == NULL)  esl_fatal(msg);
	if (p7_ReferenceForward (sq->dsq, sq->n, gm, fwd,      &fsc)  != eslOK) esl_fatal(msg);
	//if (verbose) p7_refmx_Dump(stdout, fwd);

	/* calculate entropy with Hernando algorithm */
	/* hack: Since we set gm->L to 0, we have to tell
	 * the hernando algorithm the sequence length */
	//hmm_entropy_CalculateHernando(gm, fwd, &H, sq->n, verbose);
	hmm_entropy_CalculateHernando(gm, fwd, &H, sq->n, 0);

	if (verbose) fprintf(stdout, "single path entropy: %.4f\n", H);


	if (esl_FCompare(H, 0, tol) != eslOK) esl_fatal(msg);

	p7_refmx_Destroy(fwd);
	p7_trace_Destroy(tr);
	esl_sq_Destroy(sq);
	p7_profile_Destroy(gm);
	p7_hmm_Destroy(hmm);
	p7_bg_Destroy(bg);
	esl_alphabet_Destroy(abc);

}

/* sampling unit test */
static void
utest_sampling_entropy(ESL_RANDOMNESS *rng, int alphatype, int M, int L, int verbose)
{
	char          msg[] = "random sampling entropy unit test failed";
	ESL_ALPHABET *abc        = esl_alphabet_Create(alphatype);
	P7_HMM       *hmm        = NULL;
	P7_PROFILE   *gm         = p7_profile_Create(M, abc);
	P7_BG        *bg         = p7_bg_Create(abc);
	ESL_SQ       *sq         = esl_sq_CreateDigital(abc);
	P7_TRACE     *tr         = p7_trace_Create();
	P7_REFMX     *fwd        = p7_refmx_Create(M, L);
	float         tol        = 0.01;
	int           Nsamp      = 100000000;
	float         fsc, H_hernando, H_samp;

	if ( p7_modelsample(rng, M, abc, &hmm) != eslOK) esl_fatal(msg);

	if (verbose) fprintf(stdout, "hello world: in sampling test.\n");

	if ( p7_modelsample(rng, M, abc, &hmm)             != eslOK) esl_fatal(msg);


	if ( p7_profile_ConfigUniglocal(gm, hmm, bg, L)    != eslOK) esl_fatal(msg);


	/* emit a sequence from the hmm */
	if (p7_ProfileEmit(rng, hmm, gm, bg, sq, tr)       != eslOK) esl_fatal(msg);


	/* set the length of the model to the actual length sq->n */
	if (p7_profile_SetLength(gm, sq->n)                != eslOK) esl_fatal(msg);


	if (verbose) p7_trace_Dump(stdout, tr);
	if (verbose) fprintf(stdout, "gm->L: %d\n", gm->L);
	if (verbose) fprintf(stdout, "sq->n: %d\n", sq->n);

	/* run the forward algorithm */
	p7_ReferenceForward(sq->dsq, sq->n, gm, fwd, &fsc);

	/* run the Hernando algorithm */
	hmm_entropy_CalculateHernando(gm, fwd, &H_hernando, sq->n, verbose);
	if (verbose) fprintf(stdout, "hernando algorithm entropy: %.4f\n", H_hernando);

	/* estimate entropy empirically by sampling paths */
	hmm_entropy_CalculateSampling(gm, fwd, rng, Nsamp, &H_samp, verbose);
	if (verbose) fprintf(stdout, "path sampling entropy: %.4f\n", H_samp);

	/* compare two H values, fail if they are not within 1% */
	if (esl_FCompare(H_hernando, H_samp, tol) != eslOK) esl_fatal(msg);

	p7_refmx_Destroy(fwd);
	p7_trace_Destroy(tr);
	esl_sq_Destroy(sq);
	p7_bg_Destroy(bg);
	p7_profile_Destroy(gm);
	p7_hmm_Destroy(hmm);
	esl_alphabet_Destroy(abc);
}

#endif

/* unit test driver goes here */
#ifdef HMMENTROPY_TESTDRIVE

#include <stdio.h>
#include <string.h>

#include "easel.h"
#include "esl_getopts.h"
#include "esl_random.h"

#include "hmmer.h"

static ESL_OPTIONS options[] = {
	/* name           type      default  env  range toggles reqs incomp  help                                             docgroup*/
	{ "-h",      eslARG_NONE,     FALSE, NULL, NULL,  NULL,  NULL, NULL, "show brief help on version and usage",                0 },
	{ "-s",      eslARG_INT,        "0", NULL, NULL,  NULL,  NULL, NULL, "set random number seed to <n>",                       0 },
	{ "-v",      eslARG_NONE,     FALSE, NULL, NULL,  NULL,  NULL, NULL, "be verbose",                                          0 },
	{ "-L",      eslARG_INT,       "10", NULL, NULL,  NULL,  NULL, NULL, "size of random sequences to sample",                  0 },
	{ "-M",      eslARG_INT,       "10", NULL, NULL,  NULL,  NULL, NULL, "size of random models to sample",                     0 },
	{ "-N",      eslARG_INT,      "100", NULL, NULL,  NULL,  NULL, NULL, "number of random sequences to sample",                0 },
	{  0, 0, 0, 0, 0, 0, 0, 0, 0, 0 },
};

static char usage[]  = "[-options]";
static char banner[] = "unit test driver for the Hernando posterior path entropy algorithm implementation";

int main(int argc, char **argv)
{
	ESL_GETOPTS    *go   = p7_CreateDefaultApp(options, 0, argc, argv, banner, usage);
	ESL_RANDOMNESS *r    = esl_randomness_CreateFast(esl_opt_GetInteger(go, "-s"));
	int             L    = esl_opt_GetInteger(go, "-L");
	int             M    = esl_opt_GetInteger(go, "-M");
	int             N    = esl_opt_GetInteger(go, "-N");
	int             v    = 0;

	if (esl_opt_GetBoolean(go, "-v")) v = 1;


	fprintf(stderr, "## %s\n", argv[0]);
	utest_singlepath_entropy(r,eslAMINO,M,v);
	utest_singlepath_entropy(r,eslRNA,M,v);

	utest_sampling_entropy(r,eslAMINO,M,L,v);
	fprintf(stderr, "#  status = ok\n");

	/* clean up and return */
	esl_randomness_Destroy(r);
	esl_getopts_Destroy(go);
	return 0;
}

#endif


/* 3MER example */

#ifdef HPM_3MER
#include "p7_config.h"
#include "esl_alphabet.h"
#include "easel.h"
#include "esl_sq.h"
#include "esl_sqio.h"

static ESL_OPTIONS options[] = {
	/* name           type      default  env  range  toggles reqs incomp  help                                    docgroup*/
	{ "-h",        eslARG_NONE,   FALSE, NULL, NULL, NULL, NULL, NULL, "show brief help on version and usage",                 0 },
	{ "-v",        eslARG_NONE,   FALSE, NULL, NULL, NULL, NULL, NULL, "Verbose mode: print G arrays",                         0 },

	{  0, 0, 0, 0, 0, 0, 0, 0, 0, 0 },
};

static char usage[]  = "[-options] <hmmfile> <seqfile>";
static char banner[] = "example of calculating posterior path entropy for a given hmm and sequence.";

int main(int argc, char *argv[])
{
	ESL_GETOPTS   *go      = p7_CreateDefaultApp(options, 2, argc, argv, banner, usage);
	char          *hmmfile = esl_opt_GetArg(go, 1);
	char          *seqfile = esl_opt_GetArg(go, 2);
	ESL_ALPHABET  *abc     = NULL;
	P7_HMMFILE    *hfp     = NULL;
	P7_HMM        *hmm     = NULL;
	ESL_SQ       **sq      = NULL;                      /* array of sequences */
	ESL_SQFILE    *sqfp    = NULL;
	P7_BG         *bg      = NULL;
	P7_PROFILE    *gm      = NULL;
	P7_REFMX      *fwd     = p7_refmx_Create(100, 100);
	int            format  = eslSQFILE_UNKNOWN;
	int            totseq  = 0;
	int            nseq;
	int            status;
	int            i;
	int            v       = 0;
	float          fsc;
	float          nullsc;
	float          H       = 0;

	if (esl_opt_GetBoolean(go, "-v")) v=1;

	/* Read in one HMM */
	if (p7_hmmfile_OpenE(hmmfile, NULL, &hfp, NULL) != eslOK) p7_Fail("Failed to open HMM file %s", hmmfile);
	if (p7_hmmfile_Read(hfp, &abc, &hmm)            != eslOK) p7_Fail("Failed to read HMM");
	p7_hmmfile_Close(hfp);


	/* open sequence file */
	status = esl_sqfile_OpenDigital(abc, seqfile, format, NULL, &sqfp);
	if      (status == eslENOTFOUND) p7_Fail("No such file.");
	else if (status == eslEFORMAT)   p7_Fail("Format unrecognized.");
	else if (status == eslEINVAL)    p7_Fail("Can't autodetect stdin or .gz.");
	else if (status != eslOK)        p7_Fail("Open failed, code %d.", status);

	/* read sequences into array */
	ESL_REALLOC(sq, sizeof(ESL_SQ *) * (totseq + 1));
	sq[totseq] = esl_sq_CreateDigital(abc);
	nseq = 0;
	while ((status = esl_sqio_Read(sqfp, sq[totseq+nseq])) == eslOK) {
		nseq++;
		ESL_REALLOC(sq, sizeof(ESL_SQ *) * (totseq+nseq+1));
		sq[totseq+nseq] = esl_sq_CreateDigital(abc);
	}
	/* error handling and cleanup */
	if      (status == eslEFORMAT) esl_fatal("Parse failed (sequence file %s):\n%s\n",
			                                   sqfp->filename, esl_sqfile_GetErrorBuf(sqfp));
	else if (status != eslEOF)     esl_fatal("Unexpected error %d reading sequence file %s", status, sqfp->filename);
	totseq = nseq;


	/* configure background model */
	bg = p7_bg_Create(abc);

	/* Configure a profile from the HMM in uniglocal mode*/
	gm = p7_profile_Create(hmm->M, abc);
	if (p7_profile_ConfigUniglocal(gm, hmm, bg, 400) != eslOK) esl_fatal("failed to configure profile");

	/* calculate posterior entropy for each sequence */
	for (i=0; i < totseq; i++) {
		if (i % 1000 == 0) fprintf(stdout, "%d\n", i);
		/* Set the profile and null model's target length models */
		p7_bg_SetLength     (bg, sq[i]->n);
		p7_profile_SetLength(gm, sq[i]->n);

		/* get fwd matrix */
		p7_ReferenceForward (sq[i]->dsq, sq[i]->n, gm, fwd, &fsc);
		p7_bg_NullOne(bg, sq[i]->dsq, sq[i]->n, &nullsc);

		/* if verbose mode, dump forward matrix */
		if (v) {
			fprintf(stdout, "profile configured for seq %d\n", i);
			p7_profile_Dump(stdout, gm);
			fprintf(stdout, "Forward matrix for seq %d\n", i);
			p7_refmx_Dump(stdout, fwd);
			fprintf(stdout, "Forward score (in bits) for seq %d: %.4f\n\n\n",
					  i, (fsc-nullsc) /  eslCONST_LOG2);
		}

		/* calculate H(pi | x) */
		hmm_entropy_CalculateHernando(gm, fwd, &H, -1, v);

		/* print entropy to stdout */
		fprintf(stdout, "posterior entropy for seq %d: %.4f\n", i, H);

		/* reuse fwd matrix for next iteration */
		p7_refmx_Reuse(fwd);
	}

	fprintf(stdout, "\nhello world, I am entropy!\n");

	/* clean up and return */
	esl_getopts_Destroy(go);
	esl_alphabet_Destroy(abc);
	esl_sqfile_Close(sqfp);
	p7_hmm_Destroy(hmm);
	free(sq);
	p7_profile_Destroy(gm);
	p7_bg_Destroy(bg);

	return 0;

	ERROR:
		return status;

}

#endif

#ifdef ENTROPY_MSA
#include "p7_config.h"
#include "esl_alphabet.h"
#include "easel.h"
#include "esl_sq.h"
#include "esl_msa.h"
#include "esl_msafile.h"
#include "esl_getopts.h"
#include "esl_keyhash.h"
#include "p7_config.h"
#include "hmmer.h"
#include <math.h>
#include <stdio.h>
#include <string.h>

static ESL_OPTIONS options[] = {
	/* name           type      default  env  range  toggles reqs incomp       help                                  docgroup*/
	{ "-h",        eslARG_NONE, FALSE, NULL, NULL, NULL, NULL, NULL,           "show brief help on version and usage",    0 },
	{ "--v",       eslARG_NONE, FALSE, NULL, NULL, NULL, NULL, NULL,           "Verbose mode: print G arrays",            0 },

	/* Options forcing which alphabet we're working in (normally autodetected) */
	{ "--amino",  eslARG_NONE,  FALSE, NULL, NULL, NULL,NULL,"--dna,--rna",    "<seqfile> contains protein sequences",    1 },
	{ "--dna",    eslARG_NONE,  FALSE, NULL, NULL, NULL,NULL,"--amino,--rna",  "<seqfile> contains DNA sequences",        1 },
	{ "--rna",    eslARG_NONE,  FALSE, NULL, NULL, NULL,NULL,"--amino,--dna",  "<seqfile> contains RNA sequences",        1 },

	{  0, 0, 0, 0, 0, 0, 0, 0, 0, 0 },
};

static char usage[]  = "[-options] <hmmfile> <msafile>";
static char banner[] = "compare brute force entropy calculations using all possible paths (in input msa) to hernando algorithm";

static void
cmdline_failure(char *argv0, char *format, ...)
{
	va_list argp;
	va_start(argp, format);
   vfprintf(stderr, format, argp);
	va_end(argp);
	esl_usage(stdout, argv0, usage);
	printf("\nTo see more help on available options, do %s -h\n\n", argv0);
	exit(1);

}

int main(int argc, char *argv[])
{
	ESL_GETOPTS      *go         = p7_CreateDefaultApp(options, 2, argc, argv, banner, usage);
	char             *hmmfile    = esl_opt_GetArg(go, 1);
	char             *alifile    = esl_opt_GetArg(go, 2);
	ESL_ALPHABET     *abc        = NULL;
	P7_HMMFILE       *hfp        = NULL;
	P7_HMM           *hmm        = NULL;
	ESL_MSAFILE      *afp        = NULL;
   ESL_MSA		     *msa        = NULL;
	ESL_SQ           *sq         = NULL;
	P7_TRACE        **tr         = NULL;                 /* trace for alignment paths     */
	P7_TRACE         *sctr       = p7_trace_Create();    /* for stochastic traceback     */
	P7_BG            *bg         = NULL;
	P7_PROFILE       *gm         = NULL;
	P7_REFMX         *fwd        = p7_refmx_Create(100, 100);
	ESL_KEYHASH      *h          = NULL;
	ESL_RANDOMNESS   *rng        = esl_randomness_Create(0);
	float            *q_fwd;
	float            *q_samp;
	float            *lq_fwd;
	float            *lq_samp;
	float             H_fwd;
	float             H_samp;
	float             H_cs;
	float             H_hernando;
	float            *wrk        = NULL;
	int              *matassign;
   float             fsc,nullsc;
	float             sc_ld;
	int               status;
	int               i;
	int               idx;                                /* key hash index */
	int               nq_sample    = 10000000;            /* number of stochastic traces */


	if (esl_opt_VerifyConfig(go) 					 != eslOK) cmdline_failure(argv[0], "Error in app configuration:   %s\n", go->errbuf);
	if (esl_opt_ArgNumber(go)						 != 2) 	  cmdline_failure(argv[0], "Incorrect number of command line arguments.\n", go->errbuf);

	hmmfile = esl_opt_GetArg(go, 1);
	alifile = esl_opt_GetArg(go, 2);

	/* Determine alphabet */
	if 		(esl_opt_GetBoolean(go, "--amino"))		abc = esl_alphabet_Create(eslAMINO);
	else if  (esl_opt_GetBoolean(go, "--dna"))		abc = esl_alphabet_Create(eslDNA);
	else if  (esl_opt_GetBoolean(go, "--rna"))		abc = esl_alphabet_Create(eslRNA);

	/* open the msa file */
	/* alphabet should be set in function if not user-specified */
	status = esl_msafile_Open(&abc, alifile, NULL, eslMSAFILE_UNKNOWN, NULL, &afp);
	if (status != eslOK) esl_msafile_OpenFailure(afp, status);

	/* now that we have the alphabet, set '.' as gap, too */
	//esl_alphabet_SetEquiv(abc, '.', '-');

	/* Read in the MSA */
	status = esl_msafile_Read(afp, &msa);
	if (status != eslOK) esl_msafile_ReadFailure(afp, status);

	/* set up traces from msa */
	ESL_ALLOC(tr, sizeof(P7_TRACE *) * msa->nseq);
	ESL_ALLOC(matassign, sizeof(int) * (msa->alen + 1));

	/* extract match states from alignment */
	for (i=0; i < msa->alen; i++){
		if   (msa->rf[i] != '.') matassign[i+1] = 1;
		else matassign[i+1] = 0;
	}

	/* put msa in trace array */
	p7_trace_FauxFromMSA(msa, matassign, 0, tr);

	/* allocate space for probability arrays */
	ESL_ALLOC(q_fwd, sizeof(float) * msa->nseq);
	ESL_ALLOC(q_samp, sizeof(float) * msa->nseq);
	ESL_ALLOC(lq_samp, sizeof(float) * msa->nseq);
	ESL_ALLOC(lq_fwd, sizeof(float) * msa->nseq);
	esl_vec_FSet(q_fwd,msa->nseq,0.0);
	esl_vec_FSet(q_samp,msa->nseq,0.0);
	esl_vec_FSet(lq_fwd,msa->nseq,-eslINFINITY);
	esl_vec_FSet(lq_samp,msa->nseq,-eslINFINITY);

	/* read in HMM */
	if (p7_hmmfile_OpenE(hmmfile, NULL, &hfp, NULL) != eslOK) p7_Fail("Failed to open HMM file %s", hmmfile);
	if (p7_hmmfile_Read(hfp, &abc, &hmm)            != eslOK) p7_Fail("Failed to read HMM");
	p7_hmmfile_Close(hfp);

	/* Configure a profile from the HMM */
	bg = p7_bg_Create(abc);
	gm = p7_profile_Create(hmm->M, abc);


	/* method 1: compute path probabilites using profile */

	/* get forward score for sequence */
	sq = esl_sq_CreateDigital(abc);
	esl_sq_GetFromMSA(msa,0,sq);

	/* configure profile and background models */
	p7_profile_ConfigUniglocal(gm, hmm, bg, sq->n);
	p7_bg_SetLength(bg, sq->n);
	p7_profile_SetLength(gm, sq->n);

	/* dump profile */
	p7_profile_Dump(stdout, gm);

	/* run the forward algorithm, get partial log-odds forward score, in nats */
	p7_ReferenceForward (sq->dsq, sq->n, gm, fwd, &fsc);

	/* dump forward matrix */

	p7_refmx_Dump(stdout, fwd);

	/* get null score, in nats */
	p7_bg_NullOne(bg, sq->dsq, sq->n, &nullsc);

	fprintf(stdout, "forward score: %.4f\n", (fsc - nullsc) /  eslCONST_LOG2);
	fprintf(stdout, "i,log Q(pi | x)\n");

	/* loop through msa, score individual paths */
	for(i=0; i < msa->nseq; i++)
	{
		//fprintf(stdout, "i: %d\n", i);
		sc_ld = 0.0;
		p7_trace_Dump(stdout, tr[i]);
		//fprintf(stdout, "traceback length: %d\n", tr[i]->N);
		/* Calculate Q(x,\pi) under hmm */
		p7_trace_Score(tr[i], sq->dsq, gm, &sc_ld);
		lq_fwd[i] = sc_ld - fsc;

		fprintf(stdout, "%d,%.4f,%.4f,\n", i, sc_ld, sc_ld - fsc);

	}
	/* calculate probabilities in linear space */
	esl_vec_FCopy(lq_fwd, msa->nseq, q_fwd);
	//esl_vec_FLogNorm(q_fwd, msa->nseq);
	esl_vec_FExp(q_fwd, msa->nseq);
	/* Method 2: Sample paths stochastically, empirically calculate frequencies */


	/* hash paths */
	h = esl_keyhash_Create();
	for(i=0; i < msa->nseq; i++){
		/* make state string null-terminated?? */
		esl_keyhash_Store(h, tr[i]->st, tr[i]->N, &idx);
		//fprintf(stdout, "i = %d, idx = %d\n", i, idx);
	}

	/* loop through stochastic traces */
	for (i=0; i < nq_sample; i++){
		if (i % 1000000 == 0) fprintf(stdout, "i: %d\n", i);
		p7_reference_trace_Stochastic(rng, &wrk, gm, fwd, sctr);

		/* look up array index for this trace */
		esl_keyhash_Lookup(h, sctr->st, sctr->N, &idx);
		/* if trace is not accounted for in input msa, dump it */
		if (idx < 0 || idx > msa->nseq) {
			fprintf(stdout, "Unaccounted for trace!\n");
			p7_trace_Dump(stdout, tr[i]);
		}

		/* mark this trace in the q_sample array */
      else q_samp[idx] += 1.0;

		p7_trace_Reuse(sctr);

	}

	/* normalize probability vector */
	esl_vec_FNorm(q_samp, msa->nseq);
	/* convert to log space */
	esl_vec_FCopy(q_samp, msa->nseq, lq_samp);
	esl_vec_FLog(lq_samp, msa->nseq);


	/* print out elements of q_sample array */
	for (i=0; i < msa->nseq; i++){

		fprintf(stdout, "trace index: %d, q_fwd: %.8f, lq_fwd: %.8f, q_samp: %.8f, lq_samp: %.8f\n",
				  i, q_fwd[i], lq_fwd[i], q_samp[i], lq_samp[i]);
	}

	/* calculate entropy using easel vector functions */
	H_fwd  = esl_vec_FEntropy(q_fwd, msa->nseq);
	H_samp = esl_vec_FEntropy(q_samp, msa->nseq);


	/* method 2.5 estimate entropy using hmm_entropy_CalculateSampling */
	hmm_entropy_CalculateSampling(gm, fwd, rng, nq_sample, &H_cs, 0);

	/* method 3: calculate entropy from fernando algorithm */
	hmm_entropy_CalculateHernando(gm, fwd, &H_hernando, -1, 1);

	fprintf(stdout, "Method 1 entropy (calculate q(pi | x) from hmm): %.4f\n", H_fwd);
	fprintf(stdout, "Method 2 entropy (estimate q(pi | x) via stochastic traceback: %.4f\n", H_samp);
	fprintf(stdout, "Method 2.5 entropy (estimate q(pi | x) via v2 stochastic traceback: %.4f\n", H_cs);
	fprintf(stdout, "Method 3 entropy (hernando algorithm): %.4f\n", H_hernando);
	//p7_profile_Dump(stdout, gm);

	/* clean up */
	esl_getopts_Destroy(go);
	esl_sq_Destroy(sq);
	p7_trace_DestroyArray(tr,msa->nseq);
	p7_trace_Destroy(sctr);
	esl_msa_Destroy(msa);
	p7_hmm_Destroy(hmm);
	esl_alphabet_Destroy(abc);
	esl_randomness_Destroy(rng);
	esl_keyhash_Destroy(h);
	free(wrk);
	free(matassign);
	free(q_fwd);
	free(lq_fwd);
	free(lq_samp);
	p7_refmx_Destroy(fwd);
	p7_profile_Destroy(gm);
	p7_bg_Destroy(bg);
	esl_msafile_Close(afp);

	return 0;

	ERROR:
		return status;

}

#endif
```
