## Supplemental Code S1 for "Remote homology search with hidden Potts models": hmm_entropy.h

```
/* hmm_entropy.h */

#ifndef HMM_ENTROPY_INCLUDED
#define HMM_ENTROPY_INCLUDED

#include <stdio.h>


#include "easel.h"
#include "esl_alphabet.h"
#include "esl_composition.h"
#include "esl_vectorops.h"

#include "hmmer.h"
#include "p7_config.h"
#include "dp_reference/p7_refmx.h"
#include "base/p7_profile.h"

/* hmm_entropy.c */
extern int       hmm_entropy_Calculate(P7_PROFILE *gm, P7_REFMX *fwd, float *ret_H, int verbose);
extern int       hmm_entropy_CalculateHernando(P7_PROFILE *gm, P7_REFMX *fwd, float *ret_H, int opt_L, int verbose);
extern int       hmm_entropy_CalculateSampling(P7_PROFILE *gm, P7_REFMX *fwd, ESL_RANDOMNESS *rng, int nsamp, float *ret_H, int verbose);
#endif
```
