## Supplemental Code S1 for "Remote homology search with hidden Potts models": hmm_scoreset.c

```
#include "hmm_scoreset.h"

#include "easel.h"
#include "esl_alphabet.h"
#include "esl_composition.h"

/* Function hmm_scoreset_Create()
 *
 * Synopsis: Allocate score set object for hmm scoring
 *
 *
 * Args:     nseq: number of sequences to be scored
 *
 * Returns:  pointer to the new hmm score set object
 *
 * Throws:   <NULL> if any allocation or initialization fails
 */

HMM_SCORESET *
hmm_scoreset_Create(int nseq)
{
   HMM_SCORESET *hmm_ss = NULL; /* scoreset object to return */
   int           n;            /* sequence index            */
   int           status;        /* esl return code           */

   ESL_ALLOC(hmm_ss, sizeof(HMM_SCORESET));

   /* set number of sequences */
   hmm_ss->nseq = nseq;

   /* allocate memory for sequence name array */
   ESL_ALLOC(hmm_ss->sqname, sizeof(char *) * nseq);

   /* allocate memory for probabilities/scores */
   ESL_ALLOC(hmm_ss->vsc,       sizeof(float) * nseq);
   ESL_ALLOC(hmm_ss->fsc,       sizeof(float) * nseq);
   ESL_ALLOC(hmm_ss->bsc,       sizeof(float) * nseq);
   ESL_ALLOC(hmm_ss->nullsc,    sizeof(float) * nseq);

   for (n = 0; n < nseq; n++) {
      hmm_ss->vsc[n]     = 0.0;
      hmm_ss->fsc[n]     = 0.0;
      hmm_ss->bsc[n]     = 0.0;
      hmm_ss->nullsc[n]  = 0.0;
   }

   return hmm_ss;

   ERROR:
      return NULL;
}

/* Function:  hmm_scoreset_Destroy()
 * Synopsis:  Free an <HMM_SCORESET>
 *
 * Purpose:   Frees the body of an <HMM_SCORESET>
 *
 * Note:      Based on p7_hmm_Destroy()
 *
 * Returns:   (void).
 */

void hmm_scoreset_Destroy(HMM_SCORESET *hmm_ss)
{
   if (hmm_ss == NULL) return;

   if (hmm_ss->vsc)    free(hmm_ss->vsc);
   if (hmm_ss->fsc)    free(hmm_ss->fsc);
   if (hmm_ss->bsc)    free(hmm_ss->bsc);
   if (hmm_ss->nullsc) free(hmm_ss->nullsc);
   if (hmm_ss->sqname) free(hmm_ss->sqname);

   free(hmm_ss);

}

/* Function hmm_scoreset_Write()
 *
 * Synopsis: Write hmm scores to an output .csv file
 *
 * Args:     fp:     output file
 *           hmm_ss: hmm scoreset object with scores
 *
 * Returns:  <eslOK> on success
 */

int
hmm_scoreset_Write(FILE *fp, HMM_SCORESET *hmm_ss)
{
   int n; /* sequence index */

   /* write csv header line */
   fprintf(fp,"id,raw_viterbi,raw_forward,raw_backward,logodds_viterbi,logodds_forward,logodds_backward\n");

   /* loop over sequences, print id and scores */
   for (n = 0; n < hmm_ss->nseq; n++) {
      fprintf(fp, "%s,%.4f,%.4f,%.4f,%.4f,%.4f,%.4f\n",
              hmm_ss->sqname[n],
              /* raw viterbi score, in nats */
              hmm_ss->vsc[n],
              /* raw forward score, in nats */
              hmm_ss->fsc[n],
              /* raw backward score, in nats */
              hmm_ss->bsc[n],
              /* full viterbi score, in bits */
              (hmm_ss->vsc[n] - hmm_ss->nullsc[n]) / eslCONST_LOG2,
              /* full forward score, in bits */
              (hmm_ss->fsc[n] - hmm_ss->nullsc[n]) / eslCONST_LOG2,
              /* full backward score, in bits */
              (hmm_ss->bsc[n] - hmm_ss->nullsc[n]) / eslCONST_LOG2);
   }
   return eslOK;
}
```
