## Supplemental Code S1 for "Remote homology search with hidden Potts models": hmm_scoreset.h

```
/* hmm_scoreset.h */

#ifndef HMMSCORESET_INCLUDED
#define HMMSCORESET_INCLUDED

#include <stdio.h>

typedef struct hmm_scoreset_s{
   char   **sqname;   /* sequence names [0..nseq-1][], \0-terminated */
   int      nseq;     /* number of sequences                         */
   float   *vsc;      /* viterbi scores [0..nseq-1]                  */
   float   *fsc;      /* forward scores [0..nseq-1]                  */
   float   *bsc;      /* backward scores [0..nseq-1]                 */
   float   *nullsc;   /* null model scores [0..nseq-1]               */

} HMM_SCORESET;

/* hmm_scoreset.c */
extern HMM_SCORESET *hmm_scoreset_Create(int nseq);
extern void          hmm_scoreset_Destroy(HMM_SCORESET *hmm_ss);
extern int           hmm_scoreset_Write(FILE *fp, HMM_SCORESET *hpm_SS);
#endif
```
