## Supplemental Code S1 for "Remote homology search with hidden Potts models": hmmalign_uniglocal.c

```
/* hmmalign_uniglocal.c */
/* based on example section of h4/src's dp_reference/reference_viterbi.c */

#include "p7_config.h"
#include "hmmer.h"

#include "easel.h"
#include "esl_alphabet.h"
#include "esl_getopts.h"
#include "esl_sq.h"
#include "esl_sqio.h"
#include "esl_msa.h"


static ESL_OPTIONS options[] = {
   /* name      type        default  env  range toggles reqs  incomp           help                                   docgroup*/
   { "-h",      eslARG_NONE,  FALSE, NULL, NULL, NULL,  NULL, NULL,            "show brief help on version and usage", 1 },

   /* Options forcing which alphabet we're working in (normally autodetected) */
   { "--amino", eslARG_NONE,  FALSE, NULL, NULL, NULL,  NULL, "--dna,--rna",   "<seqfile> contains protein sequences", 2 },
   { "--rna",   eslARG_NONE,  FALSE, NULL, NULL, NULL,  NULL, "--dna,--amino", "<seqfile> contains RNA sequences",     2 },
   { "--dna",   eslARG_NONE,  FALSE, NULL, NULL, NULL,  NULL, "--rna,--amino", "<seqfile> contains DNA sequences",     2 },

   {  0, 0, 0, 0, 0, 0, 0, 0, 0, 0 },
};

static char usage[]  = "[-options] <hmmfile> <seqfile> <msafile>";
static char banner[] = "simple program to produce a uniglocal alignment to an hmm";

static void
cmdline_failure(char *argv0, char *format, ...)
{
   va_list argp;
   printf("\nERROR: ");
   va_start(argp, format);
   vfprintf(stderr, format, argp);
   va_end(argp);
   esl_usage(stdout, argv0, usage);
   printf("\nTo see more help on available options, do %s -h\n\n", argv0);
   exit(1);
}

int
main(int argc, char **argv)
{
   ESL_GETOPTS    *go      = NULL;                      /* application configuration             */
   char           *hmmfile = NULL;                      /* input HMM filepath                    */
   char           *seqfile = NULL;                      /* input seq filepath                    */
   char           *msafile = NULL;                      /* output msa filepath                   */
   ESL_ALPHABET   *abc     = NULL;                      /* biological alphabet                   */
   P7_HMMFILE     *hfp     = NULL;                      /* open input hmm file stream            */
   P7_HMM         *hmm     = NULL;                      /* input hmm                             */
   ESL_SQ        **sq      = NULL;                      /* array of input seqs                   */
   ESL_SQFILE     *sqfp    = NULL;                      /* open input seq filestream             */
   P7_BG          *bg      = NULL;                      /* H3 background model                   */
   P7_PROFILE     *gm      = NULL;                      /* H3 profile model                      */
   P7_REFMX       *vit     = p7_refmx_Create(200, 400); /* Viterbi DP matrix                     */
   P7_TRACE      **tr      = NULL;                      /* array of traces for output MSA        */
   ESL_MSA        *msa     = NULL;                      /* resulting multiple alignment          */
   FILE           *afp     = NULL;                      /* output alignment file                 */
   int             outfmt  = eslMSAFILE_STOCKHOLM;      /* optut file format                     */
   float           vsc;                                 /* raw viterbi score                     */
   int             totseq  = 0;                         /* total number of seqs                  */
   int             msaopts = 0;                         /* option flags for p7_tracealign_Seqs() */
   int             nseq;                                /* seq index                             */
   int             i;                                   /* another seq index                     */
   int             format  = eslSQFILE_UNKNOWN;         /* input seq file format                 */
   int             status;                              /* esl return code                       */

   /* parse command line */
   go = esl_getopts_Create(options);
   if (esl_opt_ProcessCmdline(go, argc, argv) != eslOK) cmdline_failure(argv[0], "Failed to parse command line: %s\n", go->errbuf);
   if (esl_opt_VerifyConfig(go)               != eslOK) cmdline_failure(argv[0], "Error in app configuration:   %s\n", go->errbuf);

   if (esl_opt_GetBoolean(go, "-h") ) {
      esl_banner(stdout, argv[0], banner);
      esl_usage (stdout, argv[0], usage);
      puts("\n where options are:");
      esl_opt_DisplayHelp(stdout, go, 1, 2, 80);
      puts("\n Alphabet options:");
      esl_opt_DisplayHelp(stdout, go, 2, 2, 80);
      exit(0);
   }

   if (esl_opt_ArgNumber(go) != 3) cmdline_failure(argv[0], "Incorrect number of command line arguments.\n", go->errbuf);

   hmmfile = esl_opt_GetArg(go, 1);
   seqfile = esl_opt_GetArg(go, 2);
   msafile = esl_opt_GetArg(go, 3);

   /* if user has specified an alphabet we define it here */
   if      (esl_opt_GetBoolean(go, "--amino")) abc = esl_alphabet_Create(eslAMINO);
   else if (esl_opt_GetBoolean(go, "--rna"))   abc = esl_alphabet_Create(eslRNA);
   else if (esl_opt_GetBoolean(go, "--dna"))   abc = esl_alphabet_Create(eslDNA);

   /* Read in one HMM */
   if (p7_hmmfile_OpenE(hmmfile, NULL, &hfp, NULL) != eslOK) esl_fatal("Failed to open HMM file %s", hmmfile);
   if (p7_hmmfile_Read(hfp, &abc, &hmm)            != eslOK) esl_fatal("Failed to read HMM");
   p7_hmmfile_Close(hfp);

   /* Open sequence file */
   status = esl_sqfile_OpenDigital(abc, seqfile, format, NULL, &sqfp);
   if      (status == eslENOTFOUND) esl_fatal("No such file.");
   else if (status == eslEFORMAT)   esl_fatal("Format unrecognized.");
   else if (status == eslEINVAL)    esl_fatal("Can't autodetect stdin or .gz.");
   else if (status != eslOK)        esl_fatal("Open failed, code %d.", status);

   /* read in sequences into array */
   ESL_REALLOC(sq, sizeof(ESL_SQ *) * (totseq + 1));
   sq[totseq] = esl_sq_CreateDigital(abc);
   nseq = 0;
   while ((status = esl_sqio_Read(sqfp, sq[totseq+nseq])) == eslOK) {
      nseq++;
      ESL_REALLOC(sq, sizeof(ESL_SQ *) * (totseq+nseq+1));
      sq[totseq+nseq] = esl_sq_CreateDigital(abc);
   }
   /* error handling and cleanup */
   if      (status == eslEFORMAT) esl_fatal("Parse failed (sequence file %s):\n%s\n",
                                            sqfp->filename, esl_sqfile_GetErrorBuf(sqfp));
   else if (status != eslEOF)     esl_fatal("Unexpected error %d reading sequence file %s", status, sqfp->filename);
   esl_sqfile_Close(sqfp);
   totseq = nseq;

   /* allocate memory for traces */
   ESL_REALLOC(tr, sizeof(P7_TRACE *) * totseq);
   for (i = 0; i < totseq; i++) {
      tr[i] = p7_trace_Create();
   }

   /* Configure a profile from the HMM */
   bg = p7_bg_Create(abc);
   gm = p7_profile_Create(hmm->M, abc);

   /* Configure in uniglocal mode */
   p7_profile_ConfigUniglocal(gm, hmm, bg, 400);

   /* loop through sequence array and align seqs */
   for (i = 0; i < totseq; i++) {
      /* Set the profile and null model's target length models */
      p7_bg_SetLength     (bg, sq[i]->n);
      p7_profile_SetLength(gm, sq[i]->n);

      /* Run Viterbi - get raw score and optimal trace */
      if ( (status = p7_ReferenceViterbi(sq[i]->dsq, sq[i]->n, gm, vit, tr[i], &vsc)) != eslOK) esl_fatal("p7_ReferenceViterbi() failed, returned code %s\n", status);;

      /* prepare dp matrix for next iteration */
      p7_refmx_Reuse(vit);
   }

   /* create MSA from traces and seqs */
   msaopts |= p7_ALL_CONSENSUS_COLS; /* include all consensus columns in alignment */
   p7_tracealign_Seqs(sq, tr, totseq, hmm->M, msaopts, hmm, &msa);

   /* write MSA to file*/
   if ((afp = fopen(msafile, "w")) == NULL) esl_fatal("Failed to open output msafile %s for writing", msafile);
   esl_msafile_Write(afp, msa, outfmt);
   fclose(afp);

   /* clean up and return */
   for (i = 0; i < totseq; i++) {
      esl_sq_Destroy(sq[i]);
      p7_trace_Destroy(tr[i]);
   }
   esl_sq_Destroy(sq[totseq]);
   free(sq);
   free(tr);
   p7_hmm_Destroy(hmm);
   esl_alphabet_Destroy(abc);
   esl_msa_Destroy(msa);
   esl_getopts_Destroy(go);
   p7_profile_Destroy(gm);
   p7_refmx_Destroy(vit);
   p7_bg_Destroy(bg);
   return 0;

   ERROR:
      return status;
}
```
