## Supplemental Code S1 for "Remote homology search with hidden Potts models": hmmscore_uniglocal.c

```
/* hmmscore_uniglocal.c */
/* inspired by example in HMMER's generic_fwdback.c */

#include "hmmer.h"
#include "p7_config.h"

#include "easel.h"
#include "esl_alphabet.h"
#include "esl_getopts.h"
#include "esl_sq.h"
#include "esl_sqio.h"

#include "hmm_scoreset.h"


static ESL_OPTIONS options[] = {
   /* name      type        default env   range toggles reqs  incomp           help                                   docgroup*/
   { "-h",      eslARG_NONE, FALSE, NULL, NULL, NULL,   NULL,  NULL,           "show brief help on version and usage",  1 },

   /* Options forcing which alphabet we're working in (normally autodetected) */
   { "--amino", eslARG_NONE, FALSE, NULL, NULL, NULL,   NULL, "--dna,--rna",   "We are dealing with protein sequences", 2 },
   { "--dna",   eslARG_NONE, FALSE, NULL, NULL, NULL,   NULL, "--amino,--rna", "We are dealing with dna sequences",     2 },
   { "--rna",   eslARG_NONE, FALSE, NULL, NULL, NULL,   NULL, "--amino,--dna", "We are dealing with rna sequences",     2 },

   { 0, 0, 0, 0, 0, 0, 0, 0, 0, 0 },
};

static char usage[]  = "[-options] <hmmfile> <seqfile> <scorefile>";
static char banner[] = "calculate forward and backward scores with a glocal alignment method";

static void
cmdline_failure(char *argv0, char *format, ...)
{
   va_list argp;
   printf("\nERROR: ");
   va_start(argp, format);
   vfprintf(stderr, format, argp);
   va_end(argp);
   esl_usage(stdout, argv0, usage);
   printf("\nTo see more help on available options, do %s -h\n\n", argv0);
   exit(1);
}

int main(int argc, char **argv) {
   ESL_GETOPTS    *go        = NULL;                      /* application configuration      */
   char           *hmmfile   = NULL;                      /* input HMM filepath             */
   char           *seqfile   = NULL;                      /* input seq filepath             */
   char           *scorefile = NULL;                      /* output score csv filepath      */
   ESL_ALPHABET   *abc       = NULL;                      /* biological alphabet            */
   P7_HMMFILE     *hfp       = NULL;                      /* open input hmm file stream     */
   P7_HMM         *hmm       = NULL;                      /* input hmm                      */
   ESL_SQ        **sq        = NULL;                      /* array for input seqs           */
   ESL_SQFILE     *sqfp      = NULL;                      /* open input seq filestream      */
   P7_BG          *bg        = NULL;                      /* H3 background model            */
   P7_PROFILE     *gm        = NULL;                      /* H3 profile model               */
   P7_REFMX       *fwd       = p7_refmx_Create(100, 100); /* forward DP matrix              */
   P7_REFMX       *bck       = p7_refmx_Create(100, 100); /* backward DP matrix             */
   P7_REFMX       *vit       = p7_refmx_Create(200, 400); /* Viterbi DP matrix              */
   P7_TRACE       *tr        = p7_trace_Create();         /* trace for scoring              */
   HMM_SCORESET   *hmm_ss    = NULL;                      /* scoreset object                */
   FILE           *hmm_ss_fp = NULL;                      /* score output file stream       */
   float           fsc;                                   /* raw forward score, in nats     */
   float           bsc;                                   /* raw backward score, in nats    */
   float           vsc;                                   /* raw viterbi score, in nats     */
   float           nullsc;                                /* null transition score, in nats */
   int             totseq    = 0;                         /* total number of seqs           */
   int             nseq;                                  /* seq index                      */
   int             i;                                     /* seq index                      */
   int             format    = eslSQFILE_UNKNOWN;         /* input seq file format          */
   int             status;                                /* esl return code                */

   /* parse command line */
   go = esl_getopts_Create(options);
   if (esl_opt_ProcessCmdline(go, argc, argv) != eslOK) cmdline_failure(argv[0], "Failed to parse command line: %s\n", go->errbuf);
   if (esl_opt_VerifyConfig(go)               != eslOK) cmdline_failure(argv[0], "Error in app configuration:   %s\n", go->errbuf);

   if (esl_opt_GetBoolean(go, "-h") ) {
      esl_banner(stdout, argv[0], banner);
      esl_usage (stdout, argv[0], usage);
      puts("\n where options are:");
      esl_opt_DisplayHelp(stdout, go, 1, 2, 80);
      puts("\n Alphabet options:");
      esl_opt_DisplayHelp(stdout, go, 2, 2, 80);
      exit(0);
   }

   if (esl_opt_ArgNumber(go) != 3) cmdline_failure(argv[0], "Incorrect number of command line arguments.\n", go->errbuf);

   hmmfile   = esl_opt_GetArg(go, 1);
   seqfile   = esl_opt_GetArg(go, 2);
   scorefile = esl_opt_GetArg(go, 3);

   /* if user has specified an alphabet we define it here */
   if      (esl_opt_GetBoolean(go, "--amino")) abc = esl_alphabet_Create(eslAMINO);
   else if (esl_opt_GetBoolean(go, "--rna"))   abc = esl_alphabet_Create(eslRNA);
   else if (esl_opt_GetBoolean(go, "--dna"))   abc = esl_alphabet_Create(eslDNA);

   /* Read in one HMM */
   if (p7_hmmfile_OpenE(hmmfile, NULL, &hfp, NULL) != eslOK) esl_fatal("Failed to open HMM file %s", hmmfile);
   if (p7_hmmfile_Read(hfp, &abc, &hmm)            != eslOK) esl_fatal("Failed to read HMM");
   p7_hmmfile_Close(hfp);

   /* open sequence file */
   status = esl_sqfile_OpenDigital(abc, seqfile, format, NULL, &sqfp);
   if      (status == eslENOTFOUND) esl_fatal("No such file.");
   else if (status == eslEFORMAT)   esl_fatal("Format unrecognized.");
   else if (status == eslEINVAL)    esl_fatal("Can't autodetect stdin or .gz.");
   else if (status != eslOK)        esl_fatal("Open failed, code %d.", status);

   /* read sequences into array */
   ESL_ALLOC(sq, sizeof(ESL_SQ *) * (totseq + 1));
   sq[totseq] = esl_sq_CreateDigital(abc);
   nseq = 0;
   while ( (status = esl_sqio_Read(sqfp, sq[totseq+nseq])) == eslOK) {
      nseq++;
      ESL_REALLOC(sq, sizeof(ESL_SQ *) * (totseq+nseq+1));
      sq[totseq+nseq] = esl_sq_CreateDigital(abc);
   }

   /* error handling and cleanup */
   if      (status == eslEFORMAT) esl_fatal("Parse failed (sequence file %s):\n%s\n",
                                            sqfp->filename, esl_sqfile_GetErrorBuf(sqfp));
   else if (status != eslEOF)     esl_fatal("Unexpected error %d reading sequence file %s",
                                            status, sqfp->filename);
   totseq = nseq;

   /* Configure a profile from the HMM */
   bg = p7_bg_Create(abc);
   gm = p7_profile_Create(hmm->M, abc);

   /* Configure model in unihit glocal mode */
   //p7_ProfileConfig(hmm, bg, gm, sq->n, p7_UNIGLOCAL);  \\ h3 function
   if (p7_profile_ConfigUniglocal(gm, hmm, bg, 400) != eslOK) esl_fatal("failed to configure profile\n");

   /* set up hmm scoreset object */
   if ( (hmm_ss = hmm_scoreset_Create(nseq)) == NULL) esl_fatal("Failed so create HMM scoreset object\n");

   /* main loop over input sequences */
   for (i = 0; i < totseq; i++) {

      if (i % 1000 == 0) fprintf(stdout, "%d\n", i);

      /* Set the profile and null model's target length models */
      p7_bg_SetLength(bg, sq[i]->n);
      p7_profile_SetLength(gm, sq[i]->n);

      /* get fwd and bck scores */
      if ( (status = p7_ReferenceForward(sq[i]->dsq, sq[i]->n, gm, fwd, &fsc))  != eslOK) esl_fatal("p7_ReferenceForward() failed, returned code %s\n", status);
      if ( (status = p7_ReferenceBackward(sq[i]->dsq, sq[i]->n, gm, bck, &bsc)) != eslOK) esl_fatal("p7_ReferenceBackward() failed, returned code %s\n", status);

      /* get viterbi score */
      if ( (status = p7_ReferenceViterbi(sq[i]->dsq, sq[i]->n, gm, vit, tr, &vsc)) != eslOK) esl_fatal("p7_ReferenceViterbi() failed, returned code %s\n", status);

      /* get null score */
      p7_bg_NullOne(bg, sq[i]->dsq, sq[i]->n, &nullsc);

      /* add scores to score file */
      hmm_ss->sqname[i] = sq[i]->name;
      hmm_ss->fsc[i]    = fsc;
      hmm_ss->bsc[i]    = bsc;
      hmm_ss->vsc[i]    = vsc;
      hmm_ss->nullsc[i] = nullsc;

      /* prepare objects for next iteration */
      p7_refmx_Reuse(fwd);
      p7_refmx_Reuse(bck);
      p7_refmx_Reuse(vit);
      p7_trace_Reuse(tr);
   }

   /* write scorefile */
   if ((hmm_ss_fp = fopen(scorefile, "w")) == NULL) esl_fatal("Failed to open output hmm score setfile %s for writing", scorefile);
   hmm_scoreset_Write(hmm_ss_fp, hmm_ss);
   fclose(hmm_ss_fp);

   /* Clean up and return */
   for (i = 0; i <= totseq; i++) {
      esl_sq_Destroy(sq[i]);
   }
   esl_sqfile_Close(sqfp);
   free(sq);
   p7_hmm_Destroy(hmm);
   esl_alphabet_Destroy(abc);
   p7_refmx_Destroy(fwd);
   p7_refmx_Destroy(bck);
   p7_refmx_Destroy(vit);
   p7_profile_Destroy(gm);
   p7_trace_Destroy(tr);
   hmm_scoreset_Destroy(hmm_ss);
   p7_bg_Destroy(bg);
   esl_getopts_Destroy(go);
   return 0;

   ERROR:
      return status;
}
```
