## Supplemental Code S1 for "Remote homology search with hidden Potts models": hpm.c

```
/* hpm.c */

#include "hpm.h"
#include "p7_config.h"
#include "hmmer.h"
#include "easel.h"
#include "esl_alphabet.h"
#include "esl_composition.h"
#include "potts.h"


/* Function:  hpm_Create()
 * Synopsis:  Allocate a new <HPM>
 *
 * Purpose:   Allocate a <HPM> of <M> nodes, for symbol
 *            alphabet <abc>, and return a pointer to it.
 *
 * Note:      Based on p7_hmm_Create()
 *
 * Throws:    NULL on allocation failure.
 */

HPM *
hpm_Create(int M, ESL_ALPHABET *abc)
{
   int   Kg          = abc->K+1;  /* alphabet size w/ gaps                    */
   int   Kg2 	     = Kg*Kg;     /* (alphabet size w/ gaps) ** 2             */
   int   nTransition = 4;         /* number of transition parameters per node */
   HPM  *hpm         = NULL;      /* hpm to return                            */
   int   status;                  /* esl return code                          */
   int   i;                       /* Potts site index                         */
   int   j;                       /* Potts site index                         */
   int   k;                       /* hpm node index                           */
   int   l;                       /* character index                          */

   ESL_ALLOC(hpm, sizeof(HPM));


   hpm->M   = M;
   hpm->abc = abc;
   hpm->nTransition = nTransition;


   /* allocate memory for transition and insert emission params */

   /* level 1 */
   ESL_ALLOC(hpm->t,   (M+1) * sizeof(double *));
   ESL_ALLOC(hpm->ins, (M+1) * sizeof(double *));
   hpm->t[0]   = NULL;
   hpm->ins[0] =  NULL;

   /* level 2 */
   ESL_ALLOC(hpm->t[0],   (nTransition*(M+1)) * sizeof(double));
   ESL_ALLOC(hpm->ins[0], ((abc->K)*(M+1))    * sizeof(double));

   for (k = 0; k < M+1; k++) {
      hpm->t[k]   = hpm->t[0]   + (k * nTransition);
      hpm->ins[k] = hpm->ins[0] + (k * hpm->abc->K);
      if (abc->type == eslAMINO) {
         esl_composition_SW50(hpm->ins[k]);
      }
      else {
         for (l=0; l < abc->K; l++){
            hpm->ins[k][l] = 0.25;
         }
      }
   }

   /* allocate memory for potts params */
   ESL_ALLOC(hpm->h,  sizeof(double  *) * (M+1));
   ESL_ALLOC(hpm->e,  sizeof(double **) * (M+1));

   for (i=0; i<M+1; i++){
      ESL_ALLOC(hpm->h[i], sizeof(double ) * Kg);
      ESL_ALLOC(hpm->e[i], sizeof(double  *) * (M+1));
      for (j=0; j<M+1; j++) {
         ESL_ALLOC(hpm->e[i][j], sizeof(double  ) * Kg2);
      }
   }

   return hpm;

   ERROR:
      return NULL;
}

/* Function:  hpm_Destroy()
 * Synopsis:  Free an <HPM>
 *
 * Purpose:   Frees the body of an <HPM>
 *
 * Note:      Based on p7_hmm_Destroy()
 *
 * Returns:   (void).
 */

void
hpm_Destroy(HPM *hpm)
{
   int i,j;

   if (hpm == NULL) return;

   if (hpm->ins) {  if (hpm->ins[0]) free(hpm->ins[0]); free(hpm->ins); }
   if (hpm->t)   {  if (hpm->t[0])   free(hpm->t[0]);   free(hpm->t);   }

   /* free hi array */
   if (hpm->h && hpm->M) {
      for (i=0; i < hpm->M+1; i++) {
         if (hpm->h[i]) free(hpm->h[i]);
      }
      free(hpm->h);
   }

   /* free eij array */
   if (hpm->e && hpm->M) {
      for (i=0; i < hpm->M+1; i++) {
         for (j=0; j < hpm->M+1; j++) {
            if (hpm->e[i][j]) free(hpm->e[i][j]);
         }
         if (hpm->e[i]) free(hpm->e[i]);
      }
      free(hpm->e);
   }

   free(hpm);
   return;
}

/* Function:  hpm_Create_hmm_potts()
 * Synopsis:  Create an HPM from an HMM and a Potts model
 *
 * Purpose:   Allocate an <HPM> of <M> nodes, for symbol
 *            alphabet <abc>. Assign the <HPM> the transition
 *            and insert emission parameters from the <P7_HMM>
 *            and the Potts model parameters from the <POTTS>
 *            for match state emiissions.
 *
 * Note:      hmm and potts must have same number of match states
 *            and the same alphabet.
 *
 * Throws:    NULL on allocation failure.
 */


HPM *
hpm_Create_hmm_potts(P7_HMM *hmm, POTTS *potts, ESL_ALPHABET *abc) {
   HPM   *hpm      = NULL;   /* hpm object to return   */
   int    status;            /* esl return code        */
   int    i;                 /* position index         */
   int    j;                 /* position index         */
   int    a;                 /* character index        */
   int    b;                 /* character index        */
   int    idx;               /* character combo-index  */
   int    idx_rev;           /* character combo-index  */
   float *mocc     = NULL;   /* hmm match state probs  */

   /* check to make sure number of match states are equal */
   if (potts->L != hmm->M)     p7_Fail("Length of potts model is %d. HMM has %d match states. These must match!\n", potts->L, hmm->M);

   /* check to make sure alphabets are equal */
   if (potts->abc != hmm->abc) p7_Fail("Potts alphabet and hmm alphabet do not match!\n", potts->L, hmm->M);

   ESL_ALLOC(mocc, sizeof(float)*(hmm->M+1));

   /* create hpm with appropriate  number of match states */
   hpm = hpm_Create(hmm->M, abc);

   /* copy insert emission from hmm as-is */
   /* this throws an error for now */
   /* but the swissprot freqs are assigned in hpm_create, will fix later */
   //hpm->ins = hmm->ins;

   /* copy over potts parameters */
   for (i=0; i<hpm->M; i++) {
      for (a = 0; a < (hpm->abc->K+1); a++){
         /* assign h_i's */
         hpm->h[i+1][a] = potts->h[i][a];

         for (j = i+1; j<hpm->M; j++) {
            for (b = 0; b < (hpm->abc->K+1); b++) {

               /* assign e_ij's */
               idx = IDX(a,b,hpm->abc->K+1);
               idx_rev = IDX(b,a,hpm->abc->K+1);
               hpm->e[i+1][j+1][idx]     = potts->e[i][j][idx];
               hpm->e[j+1][i+1][idx_rev] = potts->e[j][i][idx_rev];
            }
         }
      }
   }


   /* calculate hpm transition probahilities */
   /* indices for hmm transitions */
   int hmm_MM = 0;
   int hmm_MI = 1;
   int hmm_MD = 2;
   int hmm_IM = 3;
   int hmm_II = 4;

   /* obtain match state probabilities from hmm */
   p7_hmm_CalculateOccupancy(hmm, mocc, NULL);

   /* transitions from begin state */
   hpm->t[0][HPM_MM] = hmm->t[0][hmm_MM] + hmm->t[0][hmm_MD];
   hpm->t[0][HPM_MI] = hmm->t[0][hmm_MI];
   hpm->t[0][HPM_IM] = hmm->t[0][hmm_IM];
   hpm->t[0][HPM_II] = hmm->t[0][hmm_II];

   /* intermediate transitions */
   for (i = 1; i < hpm->M; i++) {
      hpm->t[i][HPM_MM] = ((hmm->t[i][hmm_MM] + hmm->t[i][hmm_MD]) * mocc[i]) + (1.0 - mocc[i]);
      hpm->t[i][HPM_MI] = hmm->t[i][hmm_MI] * mocc[i];
      hpm->t[i][HPM_IM] = hmm->t[i][hmm_IM];
      hpm->t[i][HPM_II] = hmm->t[i][hmm_II];
   }

   /* transitions into end state */
   hpm->t[hpm->M][HPM_MM] = (hmm->t[hpm->M][hmm_MM] * mocc[i]) + (1.0 - mocc[i]);
   hpm->t[hpm->M][HPM_MI] = mocc[i]*hmm->t[0][hmm_MI];
   hpm->t[hpm->M][HPM_IM] = hmm->t[0][hmm_IM];
   hpm->t[hpm->M][HPM_II] = hmm->t[0][hmm_II];


   /* clean up and return */
   free(mocc);
   return hpm;
   ERROR:
      return NULL;


}

/* Function:  hpm_Create_3mer()
 * Synopsis:  Create a 3-state HPM for testing
 *
 * Purpose:   Creates a 3-state <HPM> for simple tests.
 *            The parameters are predetermined and worked
 *            out in my "3MER" notes.
 *
 * Note:      e_{i,j} = 5.0 if (a+b) = alphabet size
 *            AND |i-j| = 1, else 0.
 *
 * Throws:    NULL on allocation failure.
 */

HPM *
hpm_Create_3mer(ESL_ALPHABET *abc) {
   HPM  *hpm     = NULL;      /* hpm object to return      */
   int   M       = 3;         /* number of nodes           */
   int   i;                   /* site index                */
   int   j;                   /* site index                */
   int   a;                   /* alphabet index            */
   int   b;                   /* alphabet index            */
   int   idx;                 /* alphabet index for eij's  */
   int   idx2;                /* alphabet index for eij's  */
   int   Kg      = abc->K+1;  /* alphabet size w/ gap      */

   /* create hpm with 3 match states */
   hpm = hpm_Create(M, abc);

   /* set potts parameters */
   for (i = 1; i < (M+1); i++) {
      for(a = 0; a < Kg; a++) {
         /* set all h_i's to 0 */
         hpm->h[i][a] = 0;

         /* set specific set of e_12's and e_23's to be 1, else 0 */
         for (j=i+1; j < (M+1); j++) {
            for (b=0; b < Kg; b++) {
               idx  = IDX(a,b,Kg);
               idx2 = IDX(b,a,Kg);
               if ( j == (i+1) && (a+b) == abc->K) {
                  hpm->e[i][j][idx]  = 5.0;
                  hpm->e[j][i][idx2] = 5.0;
               }
               else {
                  hpm->e[i][j][idx]  = 0.0;
                  hpm->e[j][i][idx2] = 0.0;
               }
            }
         }
      }
   }

   /* set transition parameters by hand */

   /* node 0 */
   hpm->t[0][HPM_MM] = 1.0;
   hpm->t[0][HPM_MI] = 0.0;
   hpm->t[0][HPM_IM] = 1.0;
   hpm->t[0][HPM_II] = 0.0;

   /* node 1*/
   hpm->t[1][HPM_MM] = 0.9;
   hpm->t[1][HPM_MI] = 0.1;
   hpm->t[1][HPM_IM] = 0.7;
   hpm->t[1][HPM_II] = 0.3;

   /* node 2 */
   hpm->t[2][HPM_MM] = 0.8;
   hpm->t[2][HPM_MI] = 0.2;
   hpm->t[2][HPM_IM] = 0.5;
   hpm->t[2][HPM_II] = 0.5;

   /* node 3 */
   hpm->t[3][HPM_MM] = 1.0;
   hpm->t[3][HPM_MI] = 0.0;
   hpm->t[3][HPM_IM] = 1.0;
   hpm->t[3][HPM_II] = 0.0;

   return hpm;
}


/* Function:  hmm_Create_3mer()
 * Synopsis:  Create a 3-state HPM for testing
 *
 * Purpose:   Creates a 3-state <P7_HMM> for simple tests.
 *            Parameters are those of an HMM trained on
 *            infinite ensemble of sequences emitted from
 *            the 3mer HPM created in hpm_Create_3mer().
 *            .
 * Throws:    NULL on allocation failure.
 */


P7_HMM *
hmm_Create_3mer(ESL_ALPHABET *abc)
{
   P7_HMM  *hmm   = NULL;           /* hmm object to return   */
   P7_BG   *bg    = NULL;           /* background model       */
   int      M     = 3;              /* number of match states */
   int      i;                      /* node index             */
   int      a;                      /* residue index          */
   char     errbuf[eslERRBUFSIZE];  /* mem for error message  */
   int      status;                 /* esl return code        */

   /* initiate hmm object */
   hmm = p7_hmm_Create(M, abc);

   /* allocate memory for things that aren't allocated by default */
   ESL_ALLOC(hmm->consensus,  (M+2) * sizeof(char));
   ESL_ALLOC(hmm->rf,         (M+2) * sizeof(char));
   ESL_ALLOC(hmm->cs,         (M+2) * sizeof(char));
   ESL_ALLOC(hmm->ca,         (M+2) * sizeof(char));
   ESL_ALLOC(hmm->mm,         (M+2) * sizeof(char));

   hmm->rf[0] = ' ';
   hmm->cs[0] = ' ';
   hmm->ca[0] = ' ';
   hmm->mm[0] = ' ';

   hmm->rf[M+1] = '\0';
   hmm->cs[M+1] = '\0';
   hmm->ca[M+1] = '\0';
   hmm->mm[M+1] = '\0';

   /* set corresponding bit flags */
   hmm->flags |= p7H_CONS;
   hmm->flags |= p7H_COMPO;
   hmm->flags |= p7H_RF;
   hmm->flags |= p7H_CS;
   hmm->flags |= p7H_CA;
   hmm->flags |= p7H_MMASK;

   p7_hmm_SetName(hmm, "3mer");

   /* set up background model, for insertion probabilities */
   bg = p7_bg_Create(abc);

   /* set transition probs by hand */

   /* node 0 */
   hmm->t[0][p7H_MM] = ((float) abc->K) / ( (float) abc->K +1.0);
   hmm->t[0][p7H_MD] = 1./ ( (float) abc->K + 1.0);
   hmm->t[0][p7H_MI] = 0.0;
   hmm->t[0][p7H_IM] = 1.0;
   hmm->t[0][p7H_II] = 0.0;
   hmm->t[0][p7H_DM] = 1.0;
   hmm->t[0][p7H_DD] = 0.0;

   /* node 1 */
   hmm->t[1][p7H_MM] = 0.950 - 0.105;
   hmm->t[1][p7H_MD] = 0.050;
   hmm->t[1][p7H_MI] = 0.105;
   hmm->t[1][p7H_IM] = 0.7;
   hmm->t[1][p7H_II] = 0.3;
   hmm->t[1][p7H_DM] = 0.994;
   hmm->t[1][p7H_DD] = 0.006;

   /* node 2 */
   hmm->t[2][p7H_MM] = 0.950 - 0.210;
   hmm->t[2][p7H_MD] = 0.050;
   hmm->t[2][p7H_MI] = 0.210;
   hmm->t[2][p7H_IM] = 0.5;
   hmm->t[2][p7H_II] = 0.5;
   hmm->t[2][p7H_DM] = 0.994;
   hmm->t[2][p7H_DD] = 0.006;

   /* node 3 */
   hmm->t[3][p7H_MM] = 1.0;
   hmm->t[3][p7H_MD] = 0.0;
   hmm->t[3][p7H_MI] = 0.0;
   hmm->t[3][p7H_IM] = 1.0;
   hmm->t[3][p7H_II] = 0.0;
   hmm->t[3][p7H_DM] = 1.0;
   hmm->t[3][p7H_DD] = 0.0;

   /* set node 0 transitions */
   for (a=0;a<abc->K;a++){
      hmm->ins[0][a] = bg->f[a];
   }

   /* set insertion and transitionprobabilities */
   for (i=1;i<M+1;i++){

      hmm->rf[i] = '-';
      hmm->cs[i] = '-';
      hmm->ca[i] = '-';
      hmm->mm[i] = '-';

      for (a=0;a<abc->K;a++) {
         hmm->mat[i][a] = 1./ ((float)abc->K);
         hmm->ins[i][a] = bg->f[a];
      }
   }

   /* set compo value */
   p7_hmm_SetComposition(hmm);

   p7_hmm_SetConsensus(hmm, NULL);

   /* check hmm before returning */
   if (p7_hmm_Validate (hmm, errbuf, 0.0001) != eslOK) p7_Fail("3mer HMM is bad!!\n%s\n", errbuf);

   /* clean up and return */
   p7_bg_Destroy(bg);
   return hmm;

   ERROR:
   return NULL;
}

/* Function:  IDX
 * Synopsis:  Return index for accessing e_ij array.
 *
 * Purpose:   Rather than having a 4-d array for e_ij(a,b),
 *            the two K-element alphabet dimensions are compressed
 *            into one K**2 element 1-D array. This is based off
 *            Elena's Potts model code w/in RScape.
 *
 * Args:      a:   character 1
 *            b:   character 2
 *            K:   alphabet size
 */


int IDX(int a, int b, int K) {
   return (a*K) + b;
}

/* example main for testing: */
/* creates 3_mer hmm and hpm */
/* compile with make 3-mer   */

#ifdef HPM_3MER
#include "p7_config.h"
#include "esl_alphabet.h"
#include "easel.h"
#include "esl_getopts.h"
#include "esl_vectorops.h"
#include "hpmfile.h"

static ESL_OPTIONS options[] = {
   /* name           type      default  env  range  toggles reqs incomp  help                                       docgroup*/
   { "-h",        eslARG_NONE,   FALSE, NULL, NULL,   NULL,  NULL, NULL, "show brief help on version and usage",             0 },
   {  0, 0, 0, 0, 0, 0, 0, 0, 0, 0 },
};

static char usage[]  = "[-options] <hmmoutfile> <hpmoutfile>";
static char banner[] = "example of producing and writing 3mer hmm and hpm";

int main(int argc, char *argv[]) {
   ESL_GETOPTS  *go      = p7_CreateDefaultApp(options, 2, argc, argv, banner, usage);
   HPM          *hpm     = NULL;                                                        /* 3mer hmm              */
   P7_HMM       *hmm     = NULL;                                                        /* 3mer hpm              */
   ESL_ALPHABET *abc     = NULL;                                                        /* alphabet              */
   FILE         *hpm_fp  = NULL;                                                        /* open hpm file stream  */
   FILE         *hmm_fp  = NULL;                                                        /* open hmm file  stream */
   char         *hmmfile = esl_opt_GetArg(go, 1);                                       /* hmm file path         */
   char         *hpmfile = esl_opt_GetArg(go, 2);                                       /* hpm file path         */

   abc = esl_alphabet_Create(eslAMINO);

   /* create hpm */
   hpm = hpm_Create_3mer(abc);

   /* write hpm */
   if ((hpm_fp = fopen(hpmfile, "w")) == NULL) esl_fatal("Failed to open output hpm file %s for writing", hpmfile);
   hpmfile_Write(hpm_fp, hpm);
   fclose(hpm_fp);

   /* create hmm */
   hmm = hmm_Create_3mer(abc);

   /* write hmm */
   if ((hmm_fp = fopen(hmmfile, "w")) == NULL) esl_fatal("Failed to open output hmm file %s for writing", hmmfile);
   p7_hmmfile_WriteASCII(hmm_fp, -1, hmm);
   fclose(hmm_fp);

   /* clean up and return */
   esl_alphabet_Destroy(abc);
   esl_getopts_Destroy(go);
   p7_hmm_Destroy(hmm);
   hpm_Destroy(hpm);
   return 0;
}

#endif
```
