## Supplemental Code S1 for "Remote homology search with hidden Potts models": hpm.h

```
/* hpm.h */

#ifndef HPM_INCLUDED
#define HPM_INCLUDED

#include <stdio.h>

#include "hmmer.h"
#include "p7_config.h"
#include "easel.h"
#include "esl_alphabet.h"
#include "potts.h"

enum hpm_transitions {
   HPM_MM = 0,
   HPM_MI = 1,
   HPM_IM = 2,
   HPM_II = 3,
};

#define HPM_NTRANSITIONS 4
#define HPM_TMAT(hpm, k) ((hpm)->t[k])
#define HPM_TINS(hpm, k) ((hpm)->t[k]+2)
#define HPM_NTMAT 2
#define HPM_NTINS 2


typedef struct hpm_s{
   int              M;            /* number of nodes in model            */
   double         **t;            /* transition probs                    */
   double         **ins;          /* insert emission probs               */
   float          **h;            /* site-specific h_i params            */
   float         ***e;            /* coupling e_ij params                */
   int              nTransition;  /* number of transitions, should be 4  */
   ESL_ALPHABET    *abc;          /* alphabet                            */
} HPM;


/* hpm.c */
extern HPM       *hpm_Create(int M, ESL_ALPHABET *abc);
extern void       hpm_Destroy(HPM *hpm);
extern HPM       *hpm_Create_hmm_potts(P7_HMM *hmm, POTTS *potts, ESL_ALPHABET *abc);
extern HPM       *hpm_Create_3mer(ESL_ALPHABET *abc);
extern P7_HMM    *hmm_Create_3mer(ESL_ALPHABET *abc);
extern int        IDX(int a, int b, int K);

#endif
```
