## Supplemental Code S1 for "Remote homology search with hidden Potts models": hpm_scoreops.c

```
/* hpm_scoreops.c */

#include "easel.h"
#include "hmmer.h"
#include "p7_config.h"
#include "esl_sq.h"
#include "hpm.h"
#include "hpm_scoreops.h"

/* h4 includes */
#include "h4_path.h"
#include "h4_mode.h"


/*****************************************************************
 * h3-compatible scoring functions (w/ p7_traces)
*****************************************************************/

int hpm_scoreops_CalculateHamiltonian(HPM *hpm, P7_TRACE *tr, ESL_DSQ *dsq, float *ret_hsc, float *ret_esc) {
   int   z;         /* index for trace elements                   */
   int   y;         /* index for trace elements                   */
   int   i;         /* match state index                          */
   int   j;         /* match state index                          */
   int   a = 0;     /* residue index                              */
   int   b = 0;     /* residue index                              */
   int   idx;       /* index for potts parameters                 */
   float hsc = 0;   /* Hamiltonian contribution from h_i's        */
   float esc = 0;   /* Hamiltonian contribution from e_ij's       */


   int K = hpm->abc->K;

   for (z = 0; z < tr->N; z++) {

      //fprintf(stdout, "%d\n", z);
      /* check if we are in a match or delete state */
      if (tr->st[z] == 2 || tr->st[z] == 6) {
         i = tr->k[z];

         /* we have a match position */
         if (tr->st[z] == 2) {
            a = dsq[tr->i[z]];
            if (a > K) a = K;

            //fprintf(stdout, "%d\n", a);
         }

         /* we have a delete position */
         else if (tr->st[z] == 6) {
            a = K;
         }
         hsc += hpm->h[i][a];

         /* now add e_ij terms to pseudo-energy */
         for (y = z+1; y < tr->N; y++) {

            /* check if we are in a match position */
            if (tr->st[y] == 2 || tr->st[y] == 6) {
               j = tr->k[y];

               /* we have a match state */
               if (tr->st[y] == 2) {
                  b = dsq[tr->i[y]];
                  if (b > K) b = K;
               }

               /* we have a delete state */
               else if (tr->st[y] == 6) {
                  b = K;
               }

               idx = IDX(a,b,K+1);
               //fprintf(stdout, "i=%d, j=%d, a=%d, b=%d, idx=%d\n", i, j, a, b, idx);
               esc += hpm->e[i][j][idx];
            }
         }
      }
   }

   *ret_hsc = hsc;
   *ret_esc = esc;
   return eslOK;
}


int hpm_scoreops_CalculateHamiltonianSingleSite(HPM *hpm, P7_TRACE *tr, ESL_DSQ *dsq, int i, float *ret_Ei) {
   int   y;           /* index for trace elements                   */
   int   z;           /* index for trace elements                   */
   int   j;           /* match state index                          */
   int   a = 0;       /* residue index                              */
   int   b = 0;       /* residue index                              */
   int   idx;         /* index for potts parameters                 */
   float Ei = 0.0;    /* Hamiltonian contribution for site i        */
   int K = hpm->abc->K;

	//fprintf(stdout, "\t in hpm_scoreops_CalculateHamiltonianSingleSite()\n");

	/* figure out which node and residue we are dealing with */
	for (y = 0; y < tr->N; y++) {

		//fprintf(stdout, "\t\ty: %d\n", y);
		if (tr->st[y] == p7T_MG || tr->st[y] == p7T_MG) {
			if (tr->k[y] == i) {
				z = y;
				a = dsq[tr->i[z]];
				//fprintf(stdout, "\t\t\tWe've found our node! z = %d k = %d a =%d\n",
				//		  z, tr->k[y], a);
				break;
			}
		}
	}

	/* get h_i term */
	Ei += hpm->h[i][a];
	for (y = 0; y < tr->N; y++) {
		if (z == y) continue;
		//fprintf(stdout, "\t\ty: %d\n", y);

		if (tr->st[y] == p7T_MG || tr->st[y] == p7T_DG) {

			j = tr->k[y];

			/* we have a match state */
			if (tr->st[y] == p7T_MG) {
				b = dsq[tr->i[y]];
				if (b > K) b = K;
			}

			/* we have a delete state */
			else if (tr->st[y] == p7T_DG) {
				b = K;
			}

			idx = IDX(a,b,K+1);
			//fprintf(stdout, "\t\t\ti=%d, j=%d, a=%d, b=%d, idx=%d, eij[a][b]=%.4f\n",
			//		  i, j, a, b, idx,hpm->e[i][j][idx]);
         Ei += hpm->e[i][j][idx];
		}
	}

	/* loop over trace to get e_ij terms */

   *ret_Ei = Ei;

   return eslOK;
}


int hpm_scoreops_ScoreInsertEmissions(HPM *hpm, P7_TRACE *tr, ESL_DSQ *dsq, float *ret_isc)
{
	int     z;              /* index for trace elements   */
	int     i;              /* match state index          */
	int     a;              /* residue index            	*/
	float   isc = 0.0;

	for (z = 0; z < tr->N; z++) {

		/* we have an insert position */
		/* IG, N, or C states */
		if (tr->st[z] == p7T_IG || (tr->st[z] == p7T_N && tr->i[z] > 0 ) || ((tr->st[z] == p7T_C && tr->i[z] > 0 )) ) {

			/* get node index */
			i = tr->k[z];
			/* get emitted residue */
			a = dsq[tr->i[z]];
			/* update insert emission log prob */
			isc += log(hpm->ins[i][a]);
		}
	}

	*ret_isc = isc;
	return eslOK;
}

int hpm_scoreops_ScoreNullEmissions(HPM *hpm, ESL_SQ *sq, float *ret_nesc) {
   int    i;                 /* match state index             */
   int    a;                 /* residue index                 */
   float  nesc = 0.0;        /* log prob of insert emissions  */
   int    K = hpm->abc->K;   /* alphabet size                 */

   for (i = 1; i < sq->n+1; i++) {
      a = sq->dsq[i];
      /* treat all degenerate residues as first letter in abc  */
      /* insert emissions cancel in log odds score anyways     */
      /* this is possibly a half baked idea, 11/13/2018        */
      if (a > K) a = 0;
         nesc += log(hpm->ins[0][a]);

   }

   *ret_nesc = nesc;
   return eslOK;
}


int hpm_scoreops_ScoreNullMatchEmissions(HPM *hpm, P7_TRACE *tr, ESL_DSQ *dsq, float *ret_nmesc) {
   int     z;            /* index for trace elements       */
   int     i;            /* match state index                 */
   int     a;            /* residue index                  */
   float   nmesc = 0.0;  /* log prob of insert emissions     */

   int K = hpm->abc->K;

   for (z = 0; z < tr->N; z++) {
      /* we have a match state */
      if (tr->st[z] == 2) {
         i = tr->k[z];
         a = dsq[tr->i[z]];
         /* treat all degenerate residues as first letter in abc  */
         /* insert emissions cancel in log odds score anyways     */
         /* this is possibly a half baked idea, 11/13/2018        */
         if (a > K) a = 0;
         nmesc += log(hpm->ins[i][a]);
      }
   }

   *ret_nmesc = nmesc;

   return eslOK;
}


int hpm_scoreops_ScoreTransitions(HPM *hpm, P7_TRACE *tr, ESL_DSQ *dsq, int L, float *ret_tsc) {
   int   z;              /* index for trace elements               */
   int   st;
   int   stprev  = -1;   /* state id index                         */
   int   i;              /* match state index                      */
   int   iprev;          /* match state index                      */
   float tsc     = 0.0;  /* log of transition probs                */
   float tsc_NN  = 0.0;  /* log of N->N and C->C transition prob   */
   float tsc_NB  = 0.0;  /* log of N->B and C->T transition prob   */

   /* calculate N->N and C->C transition probability, in log space */
   tsc_NN = log(L) - log(L+2);
   /* calculate N->B and C->T transition probability, in log space */
   tsc_NB = log(2) - log(L+2);

   /* loop over trace positions for this seq */
   for (z = 0; z < tr->N; z++) {
      /* state type */
      st = tr->st[z];
      /* node in model */
      i = tr->k[z];

      /* handle transitions into N state */
      if (st == 8) {
         /*ignore S->N transitions, they have prob 1 */

         /* handle N->N transitions */
         if (stprev == 8)                     tsc = tsc + tsc_NN;
      }

      /* handle transitions into B state */
      else if (st == 9) {
         /* handle N->B transitons */
         if (stprev == 8)                     tsc += tsc_NB;
         /* no other transitions into B state possible */
      }

      /* handle transitions into hybrid match-del states */
      else if (st == 2 || st == 6) {
         /* handle M->M transitions */
         if      (stprev == 2 || stprev == 6) tsc += log(hpm->t[iprev][HPM_MM]);
         /* handle I->M transitions */
         else if (stprev == 4)                tsc += log(hpm->t[iprev][HPM_IM]);
         /* ignore B->M transitions, they have prob 1 */
      }
      /* handle transitions into I states */
      else if (st == 4) {
         /* handle M->I transitions */
         if      (stprev == 2 || stprev == 6) tsc += log(hpm->t[iprev][HPM_MI]);
         /* handle I->I transitions */
         else if (stprev == 4)                tsc += log(hpm->t[iprev][HPM_II]);
      }

      /* handle transitions into C state */
      else if (st == 13) {
         if (stprev == 13)                    tsc += tsc_NN;
         /* ignore E->C transitions, they have prob 1 */
      }

      /*handle transitions into T state */
      else if (st == 15) {
         if (stprev == 13)                    tsc += tsc_NB;
         /* no other possible transitions to T state */
      }

      stprev = st;
      iprev = i;
   }

   *ret_tsc = tsc;

   return eslOK;
}

int hpm_scoreops_ScoreNullTransitions(P7_BG  *bg, ESL_SQ *sq, float *ret_ntsc) {
   float ntsc = 0.0;  /* log of null transition probs */

   p7_bg_SetLength     (bg, sq->n);
   p7_bg_NullOne(bg, sq->dsq, sq->n, &ntsc);

   *ret_ntsc = ntsc;

   return eslOK;

}

/*****************************************************************
 * h4-compatible scoring functions
*****************************************************************/

/* Function:  hpm_scoreops_PathScore()
 * Synopsis:  Calculate score of a path under an HPM.
 *
 * Purpose:   Calculate the score of path <pi> and digital
 *            sequence <dsq> under hpm <hpm>. Return the
 *            unnormalized log odds score in nats in <*ret_sc>.
 *
 *            Optionally, return the individual components of the
 *            HPM score: the sum of the h_i terms (<*opt_hsc>),
 *            the sum of the e_{ij} terms (<*opt_esc>), the
 *            normalized transition score (<*opt_tsc>), and the
 *            null match emission scores (<*opt_nmsc>)
 *
 * Args:      pi:      - H4 path score
 *            dsq      - digital sequence to score
 *            hpm      - query HPM
 *            mo       - H4 mode, used to calculate flanking
 *                       transition scores
 *            ret_sc   - RETURN: Full score of <pi> and <dsq>
 *                       under <hpm>
 *            opt_hsc  - optRETURN: sum of h_i terms
 *            opt_esc  - optRETURN: sum of e_{ij} terms
 *            opt_tsc  - optRETURN: transition score
 *            opt_nmsc - optRETURN: null match emission scores
 *
 * Returns:   eslOK on success. <*ret_sc> contains score.
 *
 **/

int hpm_scoreops_ScorePath(const H4_PATH *pi, const ESL_DSQ *dsq, const HPM *hpm, const H4_MODE *mo, float *ret_sc, float *opt_hsc, float *opt_esc, float *opt_tsc, float *opt_nmsc) {
   float hsc;             /* h_i sum of Potts score                               */
   float esc;             /* e_ij sum of Potts score                              */
   float tsc;             /* transition score                                     */
   float nmsc;            /* null match emission score                            */
   int  *matseq = NULL;   /* matseq[k] = apos of match k matseq[1..M] = [1..alen] */
   int   status;          /* esl return code                                      */

   /* score transitions */
   if ((status = hpm_scoreops_ScorePathTransitions(pi, dsq, hpm, mo, &tsc, &matseq) != eslOK))
      esl_fatal("hpm_scoreops_ScorePathTransitions failed, threw status %s", status);

   /* score match emissions */
   hpm_scoreops_ScoreMatchSeq(matseq, hpm, &hsc, &esc, &nmsc);

   /* clean up and return */
   *ret_sc = hsc + esc + tsc - nmsc;
   if (opt_hsc)  *opt_hsc  = hsc;
   if (opt_esc)  *opt_esc  = esc;
   if (opt_tsc)  *opt_tsc  = tsc;
   if (opt_nmsc) *opt_nmsc = nmsc;
   free(matseq);
   return eslOK;
}

/* Function:  hpm_scoreops_ScorePathTransitions()
 * Synopsis:  Calculate transition score of a path under an HPM.
 *
 * Purpose:   Calculate the transition score of path <pi> under
 *            hpm <hpm>. Return the score in <*ret_tsc>. <pi>
 *            MUST BE GLOCAL
 *
 *            Optionally, determine the match state residues of
 *            digital sequence <dsq> and return them in
 *            <*opt_matseq>. This array can then be passed to
 *            hpm_scoreops_ScoreMatchSeq to get the hpm match
 *            emission and null match emission scores. Doing
 *            this here allows us to only unfold <pi> once
 *            when getting the entire hpm score.
 *
 *
 * Args:      pi:        - H4 path to score
 *            dsq        - digital sequence, for determining
 *                         match residues
 *            hpm        - query HPM
 *            mo         - H4 mode, used to calculate flanking
 *                         transition scores
 *            ret_tsc    - RETURN: transition score of <pi>
 *                         under <hpm>
 *            opt_matseq - optRETURN: match sequence residues,
 *                         in digital mode
 *
 * Returns:   eslOK on success. <*ret_tsc> contains score.
 *
 * Throws     eslEINVAL if non-glocal states encountered.
 *
 **/


int  hpm_scoreops_ScorePathTransitions(const H4_PATH *pi, const ESL_DSQ *dsq, const HPM *hpm, const H4_MODE *mo, float *ret_tsc, int **opt_matseq) {
   float tsc    = 0.;   /* transition score for path                                                  */
   int   i      = 0;     /* seq position index                                                        */
   int   k      = 0;     /* hmm node index                                                            */
   int   z,y;            /* counters over path elements and their runlengths                          */
   int  *matseq = NULL;  /* matseq[k=1..M]: if matuse[k] TRUE, what column 1..alen does node k map to */
   int   status;         /* esl return code                                                           */

   /* allocate space for arrays */
   ESL_ALLOC(matseq,   sizeof(int) * (hpm->M+1)); matseq[0] = 0;

   /* loop over state runs in trace */
   for (z = 0; z < pi->Z; z++) {

      /* handle each state differently */
      switch (pi->st[z]) {

         /* N-terminus (5') flanking insert states */
         case h4P_N:

            /* loop over all runs in state */
            for (y = 0; y < pi->rle[z]; y++) {
               /* handle last instance of N state: we don't assign a residue here */
               if (y == pi->rle[z] - 1) {
                  tsc += mo->xsc[h4_N][h4_MOVE] / eslCONST_LOG2R;  /* N -> G */
               }

               /* handle all other N states; we say they emit */
               else {
                  i++;
                  tsc += mo->xsc[h4_N][h4_LOOP] / eslCONST_LOG2R;  /* N -> N */
               }
            }
            break;

         /* global match state */
         case h4P_MG:

            /* loop over states in this run */
            for (y = 0; y < pi->rle[z]; y++){
               /* increment node and sequence index */
               k++;
               i++;

               /* note the residue in this match state */
               matseq[k] = dsq[i];
               /* treat degenerate chars as gaps */
               if (dsq[i] > hpm->abc->K) matseq[k] = hpm->abc->K;

               /* get transition score */
               /* look to next run if we're at the end of the run */
               /* if we are in the last match state, we transit to C w/ prob 1 */
               if (y == pi->rle[z] -1) {
                  switch(pi->st[z+1]) {
                     case h4P_MG: tsc += log(hpm->t[k][HPM_MM]); break;
                     case h4P_DG: tsc += log(hpm->t[k][HPM_MM]); break;
                     case h4P_IG: tsc += log(hpm->t[k][HPM_MI]); break;
                     case h4_C:   tsc += 0.0;                    break;
                  }
               }
               /* all intra-run transitions in run are M->M */
               else {
                  tsc += log(hpm->t[k][HPM_MM]);
               }
            }

            break;

         /* global delete state */
         case h4P_DG:

            /* loop over all states in the run */
            for (y = 0; y < pi->rle[z]; y++){

               /* increment node index ONLY */
               k++;
               matseq[k] = hpm->abc->K;

               /* get transition scores */
               /* look ahead if we're at the end of the run */
               /* if we are in the last delete state, we transit to C w/ prob 1 */
               if (y == pi->rle[z] -1) {
                  switch(pi->st[z+1]) {
                     case h4P_MG: tsc += log(hpm->t[k][HPM_MM]); break;
                     case h4P_DG: tsc += log(hpm->t[k][HPM_MM]); break;
                     case h4P_IG: tsc += log(hpm->t[k][HPM_MI]); break;
                     case h4_C:   tsc += 0.0;               break;

                  }
               }
               /* all intra-run transitions are D->D */
               else tsc += log(hpm->t[k][HPM_MM]);
            }
            break;

         /* global insert states */
         case h4P_IG:

            /* loop over all states in the run */
            for (y = 0; y < pi->rle[z]; y++){

               /* increment the sequence position index ONLY */
               i++;

               /* get transition score */
               /* if we're at the end of the run, look ahead */
               /* I THINK WE CAN GET RID OF THE SWITCH STATEMENT HERE */
               if (y == pi->rle[z] -1) {
                  tsc += log(hpm->t[k][HPM_IM]);
                  //switch(pi->st[z+1]) {
                  //   case h4P_MG: tsc += hpm->t[k][HPM_IM]; break;
                  //   case h4P_DG: tsc += hpm->t[k][HPM_IM]; break;

                  //}
               }

               /* all intra-run transitions are I->I */
               else tsc +=log(hpm->t[k][HPM_II]);
            }

            break;

         /* C-terminus (3') flanking insert states */
         case h4P_C:

            /* loop over all states in the run */
            for (y = 0; y < pi->rle[z]; y++) {

               /* get transition score */
               /* handle last instance of C state */
               if (y == pi->rle[z] - 1) {
                  tsc += mo->xsc[h4_C][h4_MOVE] / eslCONST_LOG2R;   /* C->T */
               }

               /* all but last C state emit residues, transition C->C */
               else {
                  i++;
                  tsc += mo->xsc[h4_C][h4_LOOP] / eslCONST_LOG2R;   /* C->C */
               }
            }
            break;

         /* glocal states that we skip */
         case h4P_G: break;

         /* for non-uniglocal states, tell the caller */
         default: return eslEINVAL;

      }
   }

   /* clean up and return */
   *ret_tsc = tsc;
   if (opt_matseq) *opt_matseq = matseq;
   return eslOK;

   ERROR:
      if (matseq)   free(matseq);
      *opt_matseq   = NULL;
      return status;
}

/* Function:  hpm_scoreops_ScoreMatchSeq()
 * Synopsis:  Calculate Potts model emission score and
 *            null emission score for a sequence. .
 *
 * Purpose:   Calculate match emission score and null match
 *            emission score of match sequece <mat_seq> under
 *            <HPM>.
 *
 * Args:      matseq:  - match sequence to score
 *            hpm      - query HPM
 *            ret_hsc  - RETURN: sum of h_i terms
 *            ret_esc  - RETURN: sum of e_{ij} terms
 *            ret_nmsc - null match emission scores
 *
 * Returns:   eslOK on success. <*ret_hsc>, <*ret_esc>, and
 *            <*ret_nmsc> contains scores.
 *
 **/


int hpm_scoreops_ScoreMatchSeq(int *matseq, const HPM *hpm, float *ret_hsc, float *ret_esc, float *ret_nmsc) {
   float hsc  = 0.;
   float esc  = 0.;
   float nmsc = 0.;
   int   idx;
   int   k,l;       /* match state index */

   for (k = 1; k < hpm->M+1; k++) {
      hsc +=  hpm->h[k][matseq[k]];
      if (matseq[k] < hpm->abc->K) nmsc += log(hpm->ins[k][matseq[k]]);

      for (l = k+1; l < hpm->M+1; l++) {
         idx = IDX(matseq[k], matseq[l], hpm->abc->K+1);
         esc += hpm->e[k][l][idx];
      }

   }

   *ret_hsc  = hsc;
   *ret_esc  = esc;
   *ret_nmsc = nmsc;
   return eslOK;
}
```
