## Supplemental Code S1 for "Remote homology search with hidden Potts models": hpm_scoreops.h

```
/* hpm_scoreops.h */

#ifndef HPM_SCOREOPS_INCLUDED
#define HPM_SCOREOPS_INCLUDED

#include "easel.h"
#include "hmmer.h"
#include "p7_config.h"
#include "esl_sq.h"
#include "hpm.h"

#include "h4_path.h"
#include "h4_mode.h"

/* hpm_scoreops.c */
extern int hpm_scoreops_CalculateHamiltonian(HPM *hpm, P7_TRACE *tr, ESL_DSQ *dsq, float *ret_hsc, float *ret_esc);
extern int hpm_scoreops_CalculateHamiltonianSingleSite(HPM *hpm, P7_TRACE *tr, ESL_DSQ *dsq, int i, float *ret_Ei);
extern int hpm_scoreops_ScoreInsertEmissions(HPM *hpm, P7_TRACE *tr, ESL_DSQ *dsq, float *ret_isc);
extern int hpm_scoreops_ScoreNullEmissions(HPM *hpm, ESL_SQ *sq, float *ret_nesc);
extern int hpm_scoreops_ScoreNullMatchEmissions(HPM *hpm, P7_TRACE *tr, ESL_DSQ *dsq, float *ret_nmesc);
extern int hpm_scoreops_ScoreTransitions(HPM *hpm, P7_TRACE *tr, ESL_DSQ *dsq, int L, float *ret_tsc);
extern int hpm_scoreops_ScoreNullTransitions(P7_BG  *bg, ESL_SQ *sq, float *ret_ntsc);
extern int hpm_scoreops_ScorePath(const H4_PATH *pi, const ESL_DSQ *dsq, const HPM *hpm, const H4_MODE *mo, float *ret_sc, float *opt_hsc, float *opt_esc, float *opt_tsc, float *opt_nmsc);
extern int hpm_scoreops_ScorePathTransitions(const H4_PATH *pi, const ESL_DSQ *dsq, const HPM *hpm, const H4_MODE *mo, float *ret_tsc, int **opt_matseq);
extern int hpm_scoreops_ScoreMatchSeq(int *matseq, const HPM *hpm, float *ret_hsc, float *ret_esc, float *ret_nmsc);
#endif
```
