## Supplemental Code S1 for "Remote homology search with hidden Potts models": hpm_scoreset.c

```
#include "hpm_scoreset.h"

#include "p7_config.h"
#include "hmmer.h"
#include "easel.h"
#include "esl_alphabet.h"
#include "esl_arr2.h"
#include "esl_composition.h"

#include <string.h>

/* Function  hpm_scoreset_Create()
 *
 * Synopsis: Allocate score set object for original (viterbi path) hpm scoring method
 *
 * Args:     nseq: number of sequences to be scored
 *
 * Returns:  pointer to the new hpm scoreset object
 *
 * Throws:   <NULL> if any allocation or initialization fails
 */

HPM_SCORESET *
hpm_scoreset_Create(int nseq)
{
   HPM_SCORESET *hpm_ss = NULL;
   int           n;
   int           status;

   ESL_ALLOC(hpm_ss, sizeof(HPM_SCORESET));

   /* set number of sequences */
   hpm_ss->nseq = nseq;

   /* allocate memory for sequence name array */
   ESL_ALLOC(hpm_ss->sqname, sizeof(char *) * nseq);

   /* allocate memory for probabilities/scores */
   ESL_ALLOC(hpm_ss->E_hi,         sizeof(float) * nseq);
   ESL_ALLOC(hpm_ss->E_eij,        sizeof(float) * nseq);
   ESL_ALLOC(hpm_ss->lp_ins,       sizeof(float) * nseq);
   ESL_ALLOC(hpm_ss->lp_trans,     sizeof(float) * nseq);
   ESL_ALLOC(hpm_ss->lpnull_match, sizeof(float) * nseq);
   ESL_ALLOC(hpm_ss->lpnull_trans, sizeof(float) * nseq);

   for (n = 0; n < nseq; n++) {
      hpm_ss->E_hi[n]         = 0.0;
      hpm_ss->E_eij[n]        = 0.0;
      hpm_ss->lp_ins[n]       = 0.0;
      hpm_ss->lp_trans[n]     = 0.0;
      hpm_ss->lpnull_match[n] = 0.0;
      hpm_ss->lpnull_trans[n] = 0.0;
   }

   return hpm_ss;

   ERROR:
      return NULL;
}


/* Function hpm_is_scoreset_Create()
 *
 * Synopsis: Allocate score set object for v2 (importance sampling, or "is")
 *           hpm scoring method
 *
 * Args:     nseq: number of sequences to be scored
 *
 * Returns:  pointer to the new hpm_is scoreset object
 *
 * Throws:   <NULL> if any allocation or initialization fails
 */

HPM_IS_SCORESET *
hpm_is_scoreset_Create(int nseq)
{
   HPM_IS_SCORESET *hpm_is_ss = NULL;
   int              n;
   int              status;

   ESL_ALLOC(hpm_is_ss, sizeof(HPM_IS_SCORESET));

   /* set number of sequences */
   hpm_is_ss->nseq = nseq;

   /* allocate memory for sequence name array */
   ESL_ALLOC(hpm_is_ss->sqname, sizeof(char *) * nseq);

   /* allocate memory for probabilities/scores */
   ESL_ALLOC(hpm_is_ss->R,           sizeof(int)   * nseq);
   ESL_ALLOC(hpm_is_ss->H,           sizeof(float) * nseq);
   ESL_ALLOC(hpm_is_ss->fwd,         sizeof(float) * nseq);
   ESL_ALLOC(hpm_is_ss->is_ld,       sizeof(double) * nseq);

   for (n = 0; n < nseq; n++) {
      hpm_is_ss->R[n]         = 0;
      hpm_is_ss->H[n]         = 0.0;
      hpm_is_ss->fwd[n]       = 0.0;
      hpm_is_ss->is_ld[n]     = 0.0;
   }

   return hpm_is_ss;

   ERROR:
      return NULL;
}

/* Function:  hpm_scoreset_Destroy()
 * Synopsis:  Free an <HPM_SCORESET>
 *
 * Purpose:   Frees the body of an <HPM_SCORESET>
 *
 * Note:      Based on p7_hmm_Destroy()
 *
 * Returns:   (void).
 */

void
hpm_scoreset_Destroy(HPM_SCORESET *hpm_ss) {

   if (hpm_ss == NULL) return;

   free(hpm_ss);

   return;
}

/* Function:  hpm_is_scoreset_Destroy()
 * Synopsis:  Free an <HPM_IS_SCORESET>
 *
 * Purpose:   Frees the body of an <HPM_IS_SCORESET>
 *
 * Note:      Based on p7_hmm_Destroy()
 *
 * Returns:   (void).
 */

void
hpm_is_scoreset_Destroy(HPM_IS_SCORESET *hpm_is_ss) {

   if (hpm_is_ss == NULL) return;

   if (hpm_is_ss->sqname)  free(hpm_is_ss->sqname);
   if (hpm_is_ss->R)       free(hpm_is_ss->R);
   if (hpm_is_ss->H)       free(hpm_is_ss->H);
   if (hpm_is_ss->fwd)     free(hpm_is_ss->fwd);
   if (hpm_is_ss->is_ld)   free(hpm_is_ss->is_ld);

   free(hpm_is_ss);
   return;
}

/* Function hpm_scoreset_Write()
 *
 * Synopsis: Write original (Viterbi-path) hpm scores to an output .csv file
 *
 * Args:     fp:     output file
 *           hpm_ss: original hpm scoreset object with scores
 *
 * Returns:  <eslOK> on success
 */

int
hpm_scoreset_Write(FILE *fp, HPM_SCORESET *hpm_ss){
   int n; /* sequence index */

   /* write csv header line */
   fprintf(fp,"id,E_hi,E_eij,hamiltonian,lp_ins,lp_trans,lpnull_match,lpnull_trans,logodds_unnormalized\n");

   /* loop over sequences, print id and scores */
   for (n = 0; n < hpm_ss->nseq; n++) {
      fprintf(fp, "%s,%.4f,%.4f,%.4f,%.4f,%.4f,%.4f,%.4f,%.4f\n",
              hpm_ss->sqname[n],
              hpm_ss->E_hi[n],
              hpm_ss->E_eij[n],
              hpm_ss->E_hi[n] + hpm_ss->E_eij[n],
              hpm_ss->lp_ins[n],
              hpm_ss->lp_trans[n],
              hpm_ss->lpnull_match[n],
              hpm_ss->lpnull_trans[n],
              hpm_ss->E_hi[n] + hpm_ss->E_eij[n] + hpm_ss->lp_trans[n] - hpm_ss->lpnull_match[n] - hpm_ss->lpnull_trans[n]);
   }

   return eslOK;
}

/* Function hpm_is_scoreset_Write()
 *
 * Synopsis: Write v2 (importance sampling) hpm scores to an output .csv file
 *
 * Args:     fp:        output file
 *           hpm_is_ss: v2 hpm_is scoreset object with scores
 *
 * Returns:  <eslOK> on success
 */

int
hpm_is_scoreset_Write(FILE *fp, HPM_IS_SCORESET *hpm_is_ss){
   int n; /* sequence index */


   /* write csv header line */
   fprintf(fp,"id,N_sample,path_ent,fwd_logodds,HPM_IS_logodds\n");

   /* loop over sequences, print id and scores */
   for (n = 0; n < hpm_is_ss->nseq; n++) {
      fprintf(fp, "%s,%d,%.4f,%.4f,%4f\n",
              hpm_is_ss->sqname[n],
              hpm_is_ss->R[n],
              hpm_is_ss->H[n],
              hpm_is_ss->fwd[n],
              hpm_is_ss->is_ld[n]);
   }

   return eslOK;
}
```
