## Supplemental Code S1 for "Remote homology search with hidden Potts models": hpm_scoreset.h

```
/* hpm_scoreset.h */

#ifndef HPM_SCORESET_INCLUDED
#define HPM_SCORESET_INCLUDED

#include <stdio.h>

#include "hmmer.h"
#include "p7_config.h"
#include "easel.h"
#include "esl_alphabet.h"
#include "potts.h"
#include "esl_arr2.h"

typedef struct hpmscoreset_s{
   char   **sqname;        /* sequence names [0..nseq-1][], \0-terminated            */
   int      nseq;          /* number of sequences                                    */
   float   *E_hi;          /* Potts h_i pseudo-energies [0...nseq-1]                 */
   float   *E_eij;         /* Potts e_ij pseudo-energies [0...nseq-1]                */
   float   *lp_ins;        /* product of insertion probabilities [0...nseq-1]        */
   float   *lp_trans;      /* product of transition probabilities [0...nseq-1]       */
   float   *lpnull_match;  /* null model probabilities of match states [0...nseq-1]  */
   float   *lpnull_trans;  /* null model probabilities of transitions  [0...nseq-1]  */
} HPM_SCORESET;


typedef struct hpmisscoreset_s{
   char   **sqname;        /* sequence names [0..nseq-1][], \0-terminated            */
   int      nseq;          /* number of sequences                                    */
   int     *R;             /* number of stochastic traces                            */
   float   *H;             /* Posterior path entropies H(pi | x)                     */
   float   *fwd;           /* forward log-odds scores                                */
   double  *is_ld;         /* unnormalized importance sampling log-odds scores       */

} HPM_IS_SCORESET;


/* hpm_scoreset.c */
extern HPM_SCORESET     *hpm_scoreset_Create(int nseq);
extern HPM_IS_SCORESET  *hpm_is_scoreset_Create(int nseq);
extern void              hpm_scoreset_Destroy(HPM_SCORESET *hpm_ss);
extern void              hpm_is_scoreset_Destroy(HPM_IS_SCORESET *hpm_is_ss);
extern int               hpm_scoreset_Write(FILE *fp, HPM_SCORESET *hpm_ss);
extern int               hpm_is_scoreset_Write(FILE *fp, HPM_IS_SCORESET *hpm_is_ss);


#endif
```
