## Supplemental Code S1 for "Remote homology search with hidden Potts models": hpm_trace.h

```
/* hpm_trace.h */
#ifndef HPMTRACE_INCLUDED
#define HPMTRACE_INCLUDED

#include "hmmer.h"
#include "p7_config.h"

#include "easel.h"
#include "esl_vectorops.h"


#include "hpm_trace.h"

/* hpm_trace.c */
extern int         hpm_trace_Copy(const P7_TRACE *src, P7_TRACE *dst);
extern P7_TRACE   *hpm_trace_Clone(const P7_TRACE *tr);

#endif
```
