## Supplemental Code S1 for "Remote homology search with hidden Potts models": hpmalign.c

```
/* hpmalign.c */
/* align sequences to an hpm via importance sampling w/ an hmm */
#include <math.h>

#include "hmmer.h"
#include "easel.h"

#include "p7_config.h"

#include "esl_getopts.h"
#include "esl_alphabet.h"
#include "esl_sq.h"
#include "esl_sqio.h"
#include "esl_vectorops.h"

#include "hpm.h"
#include "hpmfile.h"
#include "hmm_entropy.h"
#include "hpm_scoreops.h"
#include "hpm_trace.h"

/* declaration of internal functions */
int IS_align(HPM *hpm, P7_HMM *hmm, ESL_SQ **sq, ESL_MSA **msa, ESL_RANDOMNESS *rng, int totseq, int verbose);


static ESL_OPTIONS options[] = {
	/* name         type        default env   range togs  reqs  incomp      help                                                   docgroup */
	{ "-h",         eslARG_NONE,  FALSE, NULL, NULL, NULL, NULL, NULL,           "help; show brief info on version and usage",                 0 },
	{ "-s",         eslARG_INT,     "0", NULL, NULL, NULL, NULL, NULL,           "set random number seed to <n>",                              0 },

	/* Options forcing which alphabet we're working in (normally autodetected) */
	{ "--amino",    eslARG_NONE,  FALSE, NULL, NULL, NULL,NULL,"--dna,--rna",    "<seqfile> contains protein sequences",                       1 },
	{ "--rna",      eslARG_NONE,  FALSE, NULL, NULL, NULL,NULL,"--dna,--amino",  "<seqfile> contains RNA sequences",                           1 },
	{ "--dna",      eslARG_NONE,  FALSE, NULL, NULL, NULL,NULL,"--rna,--amino",  "<seqfile> contains DNA sequences",                           1 },

	/* debugging tools */
	{ "--v",        eslARG_NONE,  FALSE, NULL, NULL, NULL,NULL,NULL,              "Verbose mode: print info on intermediate alignment steps",   2 },
	{  0, 0, 0, 0, 0, 0, 0, 0, 0, 0 },
};

static char usage[]  = "[-options] <hpmfile> <hmmfile> <seqfile> <msa_outfile>";
static char banner[] = "align sequences to a profile HPM";

int main(int argc, char **argv){

	ESL_GETOPTS      *go            =  p7_CreateDefaultApp(options, 4, argc, argv, banner, usage);
	ESL_RANDOMNESS   *rng           = esl_randomness_Create(esl_opt_GetInteger(go, "-s"));
	int               v             = 0;
	char             *hpmfile       = esl_opt_GetArg(go, 1);
	char             *hmmfile       = esl_opt_GetArg(go, 2);
	char             *seqfile       = esl_opt_GetArg(go, 3);
	char             *msafile       = esl_opt_GetArg(go, 4);
	ESL_ALPHABET     *abc           = NULL;
	HPM              *hpm           = NULL;
	P7_HMMFILE       *hmmfp         = NULL;
	P7_HMM           *hmm           = NULL;
	ESL_SQ          **sq            = NULL;                      /* array of sequences */
	ESL_SQFILE       *sqfp          = NULL;
	FILE             *afp           = NULL;                      /* output alignment file */
	ESL_MSA          *msa           = NULL;                      /* resulting msa */
	int               outfmt        = eslMSAFILE_STOCKHOLM;
	int               format        = eslSQFILE_UNKNOWN;
	int               totseq        = 0;
	int               nseq          = 0;
	int status;
	char              errbuf[eslERRBUFSIZE];

	/* check for verbose mode */
	if (esl_opt_GetBoolean(go, "--v")) v=1;

	/* if user has defined an alphabet we define it here */
	if        (esl_opt_GetBoolean(go, "--amino"))       abc = esl_alphabet_Create(eslAMINO);
	else if   (esl_opt_GetBoolean(go, "--rna"))         abc = esl_alphabet_Create(eslRNA);
	else if   (esl_opt_GetBoolean(go, "--dna"))         abc = esl_alphabet_Create(eslDNA);

	/* read in hmm, autodetect alphabet if not manually set by user  */
	if (p7_hmmfile_OpenE(hmmfile, NULL, &hmmfp, NULL) != eslOK) p7_Fail("Failed to open HMM file %s", hmmfile);
	if (p7_hmmfile_Read(hmmfp, &abc, &hmm)        != eslOK) p7_Fail("Failed to read HMM");
	p7_hmmfile_Close(hmmfp);

	/* open hpm */
	hpm = hpmfile_Read(hpmfile, abc, errbuf);

	/* open sequence file */
	status = esl_sqfile_OpenDigital(abc, seqfile, format, NULL, &sqfp);
	if      (status == eslENOTFOUND) p7_Fail("No such file.");
	else if (status == eslEFORMAT)   p7_Fail("Format unrecognized.");
	else if (status == eslEINVAL)    p7_Fail("Can't autodetect stdin or .gz.");
	else if (status != eslOK)        p7_Fail("Open failed, code %d.", status);

	ESL_REALLOC(sq, sizeof(ESL_SQ *) * (totseq + 1));
	sq[totseq] = esl_sq_CreateDigital(abc);
	while ((status = esl_sqio_Read(sqfp, sq[totseq+nseq])) == eslOK) {
		nseq++;
		ESL_REALLOC(sq, sizeof(ESL_SQ *) * (totseq+nseq+1));
		sq[totseq+nseq] = esl_sq_CreateDigital(abc);
	}
	totseq = nseq;

	/* align sequences */
	IS_align(hpm, hmm, sq, &msa, rng, totseq, v);


	/* write MSA to file*/
	if ((afp = fopen(msafile, "w")) == NULL) esl_fatal("Failed to open output msafile %s for writing", msafile);
	esl_msafile_Write(afp, msa, outfmt);
	fclose(afp);

	/* clean up and return */
	esl_sqfile_Close(sqfp);
	free(sq);
	p7_hmm_Destroy(hmm);
	free(hpm);
	esl_alphabet_Destroy(abc);
	esl_getopts_Destroy(go);
	esl_msa_Destroy(msa);

	return 0;

	ERROR:
		return status;
}


/* Function: IS_align()
 * Synopisis: align sequences to an hmm by importance sampling paths
 * (traces) from an hmm
 *
 * args: hpm     -  hpm object
 *       hpm     -  corresponding hmm object
 *       seq     -  array of sequences
 *       msa     -  msa to be created
 *       rng     -  <ESL_RANDOMNESS *> random number generator
 *       totseq  -  total number of sequences in seq
 *       verbose -  boolean integer specifying verbose mode
 *
 *       Returns: <eslOK> on success
 */


int IS_align(HPM *hpm, P7_HMM *hmm, ESL_SQ **sq, ESL_MSA **msa, ESL_RANDOMNESS *rng, int totseq, int verbose)
{
	P7_BG          *bg        = NULL;
	P7_PROFILE     *gm        = NULL;
	P7_REFMX       *fwd       = p7_refmx_Create(100, 100);
	P7_TRACE       *tr        = p7_trace_Create();
	P7_TRACE      **out_tr    = NULL;                     /* traces used to construct msa */
	float           fsc;
	float           hsc;
	float           esc;
	float           isc;
	float           tsc;
	float           hpmsc;
	float           hpmsc_max;
	float          *wrk       = NULL;
	int             i;                                    /* sequence index */
	int             r;                                    /* sample index */
	int             R         = 10;                        /* total number of samples/sequence */
	float           H;                                    /* posterior path entropy */
	int             msaopts   = 0;
	int             status;

	fprintf(stdout, "in funtion IS_align()\n");

	/* allocate memory for traces  and initialize */
	ESL_ALLOC(out_tr, sizeof(P7_TRACE *) * totseq);

	/* create a profile from the HMM */
	gm = p7_profile_Create(hmm->M, hmm->abc);
	/* create null model*/
	bg = p7_bg_Create(hmm->abc);
	if (p7_profile_ConfigUniglocal(gm, hmm, bg, 400) != eslOK) esl_fatal("failed to configure profile");

	/* outer loop over all sequences in seqfile */
	for (i = 0; i < totseq; i++) {
		hpmsc_max = -eslINFINITY;

		if (i == 0) fprintf(stdout, "%d\n", i);

		/* Set the profile and null model's target length models */
		p7_profile_SetLength(gm, sq[i]->n);

		/* calculate forward dp matrix, forward score */
		p7_ReferenceForward(sq[i]->dsq, sq[i]->n, gm, fwd, &fsc);

		/* calculate posterior entropy H(pi | x) */
		hmm_entropy_CalculateHernando(gm, fwd, &H, -1, 0);

		R = 2*pow(2,ceil(H));
		if (H > 17.0)
		{
			R = 130000;  /* approx 2^17 */
		}
		else if (R < 1000)
		{
			R = 1000;
		}
		//if (R < 1000) R = 1000;
		fprintf(stdout, "seq: %s, H: %.4f, R: %d\n", sq[i]->name, H, R);

		/* inner loop over samples */
		for (r = 0; r < R; r++) {
			if (verbose) fprintf(stdout, "\nsequence %s, stochastic trace %d\n", sq[i]->name, r);

			/* perform stochastic traceback */
			p7_reference_trace_Stochastic(rng, &wrk, gm, fwd, tr);

			/* calculate Potts Hamiltonian */
			hpm_scoreops_CalculateHamiltonian(hpm, tr, sq[i]->dsq, &hsc, &esc);

			/* score insert state  emissions */
			hpm_scoreops_ScoreInsertEmissions(hpm, tr, sq[i]->dsq, &isc);

			/* score transitions */
			hpm_scoreops_ScoreTransitions(hpm, tr, sq[i]->dsq, sq[i]->L, &tsc);

			hpmsc = hsc + esc + isc + tsc;

			if (hpmsc > hpmsc_max) {
				hpmsc_max = hpmsc;
				out_tr[i] = hpm_trace_Clone(tr);
			}

			if (verbose) {
				p7_trace_DumpAnnotated(stdout, tr, gm, sq[i]->dsq);
				fprintf(stdout, "\n");
				fprintf(stdout, "# length of traceback: %d\n", tr->N);
				fprintf(stdout, "# local field hamiltonian sum: %.4f\n", hsc);
				fprintf(stdout, "# coupling hamiltonian sum: %.4f\n", esc);
				fprintf(stdout, "# HPM insert emissions log prob: %.4f\n", isc);
				fprintf(stdout, "# HPM transitions log prob: %.4f\n", tsc);
				fprintf(stdout, "# Total HPM log prob: %.4f\n", hpmsc);
				fprintf(stdout, "# Max HPM log prob: %.4f\n", hpmsc_max);
				fprintf(stdout, "\n");
			}
			p7_trace_Reuse(tr);
		}
	}

	/* create MSA from trace */
	msaopts |= p7_ALL_CONSENSUS_COLS; /* include all consensus columns in alignment */
	p7_tracealign_Seqs(sq, out_tr, totseq, hmm->M, msaopts, hmm, msa);

	/* clean up and return */
	p7_profile_Destroy(gm);
	p7_bg_Destroy(bg);
	p7_refmx_Destroy(fwd);
	p7_trace_Destroy(tr);
	free(out_tr);
	return eslOK;

	ERROR:
		return status;

}
```
