## Supplemental Code S1 for "Remote homology search with hidden Potts models": hpmbuild.c

```
#include <stdio.h>
#include <stdlib.h>
#include <string.h>

#include "easel.h"
#include "esl_alphabet.h"
#include "esl_getopts.h"

#include "p7_config.h"
#include "hmmer.h"

#include "hpm.h"
#include "hpmfile.h"
#include "potts.h"
#include "pottsfile.h"

static ESL_OPTIONS options[] = {
   /* name       type         default env   range togs  reqs  incomp           help                                                  docgroup */
   { "-h",       eslARG_NONE,  FALSE, NULL, NULL, NULL, NULL, NULL,            "help; show brief info on version and usage",           1 },

   /* Options forcing which alphabet we're working in (normally autodetected) */
   { "--amino",  eslARG_NONE,  FALSE, NULL, NULL, NULL, NULL, "--dna,--rna",   "<pottsfile> and <hmmfile> are for protein sequences",  2 },
   { "--dna",    eslARG_NONE,  FALSE, NULL, NULL, NULL, NULL, "--amino,--rna", "<pottsfile> and <hmmfile> are for dna sequences",      2 },
   { "--rna",    eslARG_NONE,  FALSE, NULL, NULL, NULL, NULL, "--amino,--dna", "<pottsfile> and <hmmfile> are for rna sequences",      2 },

   /* Option to create 3-mer dummy protein hpm */
   { "--3mer",   eslARG_NONE,  FALSE, NULL, NULL, NULL, NULL, NULL,            "create dummy 3-mer hpm, usage = [-options] <hpmfile>", 3 },
   { 0,0,0,0,0,0,0,0,0,0 },
};

static char banner[] = "Build an HPM from an HMM and a Potts model (w/ same number of match states)";
static char usage[]  = "[-options] <hpmfile> <hmmfile> <pottsfile>";

static void
cmdline_failure(char *argv0, char *format, ...)
{
   va_list argp;
   va_start(argp, format);
   vfprintf(stderr, format, argp);
   va_end(argp);
   esl_usage(stdout, argv0, usage);
   printf("\nTo see more help on available options, do %s -h\n\n", argv0);

   exit(1);
}

int main(int argc, char *argv[]) {
   ESL_GETOPTS   *go;                             /* cmd line configuration    */
   ESL_ALPHABET  *abc                    = NULL;  /* esl_alphabet              */
   P7_HMMFILE    *hfp;                            /* input hmm file            */
   P7_HMM        *hmm;                            /* input hmm                 */
   POTTS         *potts;                          /* input potts model         */
   HPM           *hpm;                            /* hpm to be created         */
   char          *hpm_outfile            = NULL;  /* hpm output file path      */
   char          *hmmfile                = NULL;  /* hmm file path             */
   char          *pottsfile              = NULL;  /* hmm file path             */
   FILE          *hpm_outfp              = NULL;  /* hpm output file object    */
   int            status;                         /* easel return code         */
   char           errbuf[eslERRBUFSIZE];          /* for error messages        */

   /* parse command line */
   go = esl_getopts_Create(options);

   if (esl_opt_ProcessCmdline(go, argc, argv) != eslOK) cmdline_failure(argv[0], "Failed to parse command line: %s\n", go->errbuf);
   if (esl_opt_VerifyConfig(go)               != eslOK) cmdline_failure(argv[0], "Error in app configuration:   %s\n", go->errbuf);

   /* output help if requested */
   if (esl_opt_GetBoolean(go, "-h") ) {
      esl_banner(stdout, argv[0], banner);
      esl_usage (stdout, argv[0], usage);
      puts("\n where options are:");
      esl_opt_DisplayHelp(stdout, go, 1, 2, 80);
      puts("\n Alphabet options:");
      esl_opt_DisplayHelp(stdout, go, 2, 2, 80);
      puts("\n 3mer options:");
      esl_opt_DisplayHelp(stdout, go, 3, 2, 80);
      exit(0);
   }

   /* check and see if we are making a dummy 3-mer model */

   if (esl_opt_GetBoolean(go, "--3mer")) {

      /* we only want 1 argument */
      if (esl_opt_ArgNumber(go) != 1) cmdline_failure(argv[0], "Incorrect number of command line arguments w/ dummy 3-mer hpm.\n", go->errbuf);
      hpm_outfile = esl_opt_GetArg(go, 1);

      /* set alphabet to amino, forget what user says */
      abc = esl_alphabet_Create(eslAMINO);

      /* make 3-mer hpm*/
      hpm = hpm_Create_3mer(abc);

      /* write to output file */
      /* open hpm outfile for writing, write hpm to it */
      if ((hpm_outfp = fopen(hpm_outfile, "w")) == NULL) esl_fatal("Failed to open output hpm file %s for writing", hpm_outfile);
      hpmfile_Write(hpm_outfp, hpm);
      fclose(hpm_outfp);

      /* clean up and return */
      esl_alphabet_Destroy(abc);
      hpm_Destroy(hpm);
      esl_getopts_Destroy(go);

      return 0;
   }

   /* normal, non-3mer mode of operation */
   if (esl_opt_ArgNumber(go) != 3) cmdline_failure(argv[0], "Incorrect number of command line arguments.\n", go->errbuf);

   hpm_outfile = esl_opt_GetArg(go, 1);
   hmmfile     = esl_opt_GetArg(go, 2);
   pottsfile   = esl_opt_GetArg(go, 3);

   /* Set up the configuration structure shared amongst functions here */
   /* Determine alphabet: do what user says, else autodetermine w/ hmm */
   if      (esl_opt_GetBoolean(go, "--amino"))    abc = esl_alphabet_Create(eslAMINO);
   else if (esl_opt_GetBoolean(go, "--dna"))      abc = esl_alphabet_Create(eslDNA);
   else if (esl_opt_GetBoolean(go, "--rna"))      abc = esl_alphabet_Create(eslRNA);

   /* Open the .hmm file */
   status = p7_hmmfile_OpenE(hmmfile, NULL, &hfp, errbuf);
   /* error handling below copied from hmmsearch */
   if      (status == eslENOTFOUND) esl_fatal("File existence/permissions problem in trying to open HMM file %s.\n%s\n", hmmfile, errbuf);
   else if (status == eslEFORMAT)   esl_fatal("File format problem in trying to open HMM file %s.\n%s\n",                hmmfile, errbuf);
   else if (status != eslOK)        esl_fatal("Unexpected error %d in opening HMM file %s.\n%s\n",               status, hmmfile, errbuf);

   /* read the .hmm file */
   status = p7_hmmfile_Read(hfp, &abc, &hmm);
   if (status != eslOK) esl_fatal("ERROR reading hmm, returned code %d\n", status);
   p7_hmmfile_Close(hfp);

   /* read the potts file */
   potts = pottsfile_Read(pottsfile, abc, errbuf);
   if (potts == NULL) esl_fatal("Error reading potts file %s\n", pottsfile);

   /* combine hmm and potts object to create hpm object */
   hpm = hpm_Create_hmm_potts(hmm, potts, abc);

   /* open hpm outfile for writing, write hpm to it */
   if ((hpm_outfp = fopen(hpm_outfile, "w")) == NULL) esl_fatal("Failed to open output hpm file %s for writing", hpm_outfile);
   hpmfile_Write(hpm_outfp, hpm);

   /* close hpm outfile */
   esl_alphabet_Destroy(abc);
   hpm_Destroy(hpm);
   p7_hmm_Destroy(hmm);
   potts_Destroy(potts);
   esl_getopts_Destroy(go);
   fclose(hpm_outfp);
   return 0;

}
```
