## Supplemental Code S1 for "Remote homology search with hidden Potts models": hpmbuild_masked.c

```
#include <stdio.h>
#include <stdlib.h>
#include <string.h>

#include "easel.h"
#include "esl_alphabet.h"
#include "esl_arr2.h"
#include "esl_arr3.h"
#include "esl_getopts.h"
#include "esl_msa.h"
#include "esl_msafile.h"
#include "esl_msaweight.h"
#include "esl_vectorops.h"
#include "esl_wuss.h"

#include "p7_config.h"
#include "hmmer.h"

#include "hpm.h"
#include "hpmfile.h"
#include "potts.h"

/* declaration of internal functions */
static int trim_msa_rf(ESL_MSA *msa, ESL_ALPHABET *abc, char *errbuf);
static int find_basepairs(ESL_MSA *msa, int pk, int **ret_ct, char *errbuf);
static int count_msa(ESL_MSA *msa, int *ct, int use_weights, double ***ret_abc_ct, double ****ret_bp_ct, char *errbuf);
static int counts_to_potts(ESL_MSA *msa,  double **abc_ct, double ***bp_ct, int *ct, POTTS **potts);

static ESL_OPTIONS options[] = {
   /* name         type        default env   range togs  reqs  incomp            help                                                 docgroup */
   { "-h",         eslARG_NONE, FALSE, NULL, NULL, NULL, NULL, NULL,            "help; show brief info on version and usage",          1 },

   /* Options forcing which alphabet we're working in (normally autodetected) */
   { "--amino",   eslARG_NONE,  FALSE, NULL, NULL, NULL, NULL, "--dna,--rna",   "<pottsfile> and <hmmfile> are for protein sequences", 2 },
   { "--dna",     eslARG_NONE,  FALSE, NULL, NULL, NULL, NULL, "--amino,--rna", "<pottsfile> and <hmmfile> are for dna sequences",     2 },
   { "--rna",     eslARG_NONE,  FALSE, NULL, NULL, NULL, NULL, "--amino,--dna", "<pottsfile> and <hmmfile> are for rna sequences",     2 },

   /* training options */
   { "--pknot",   eslARG_NONE,  FALSE, NULL, NULL, NULL, NULL, NULL,            "train coupling terms for annotated pseudoknots",      3 },
   { "--weight",  eslARG_NONE,  FALSE, NULL, NULL, NULL, NULL, NULL,            "weight MSA counts with Henikiff PB weights",          3 },

   { 0,0,0,0,0,0,0,0,0,0 },
};

static char banner[] = "Train a hidden potts model using maximum likelihood and secondary sequence annotation";
static char usage[]  = "[-options] <hpmfile> <hmmfile> <msafile>";

static void
cmdline_failure(char *argv0, char *format, ...)
{
   va_list argp;
   printf("\nERROR: ");
   va_start(argp, format);
   vfprintf(stderr, format, argp);
   va_end(argp);
   esl_usage(stdout, argv0, usage);
   printf("\nTo see more help on available options, do %s -h\n\n", argv0);
   exit(1);
}

int main(int argc, char *argv[])  {
   ESL_GETOPTS  *go           = NULL;                /* application configuration                                    */
   ESL_ALPHABET *abc          = NULL;                /* biological alphabet                                          */
   char         *alifile      = NULL;                /* alignment file name                                          */
   ESL_MSAFILE  *afp          = NULL;                /* open msa file                                                */
   ESL_MSA      *msa          = NULL;                /* one multiple sequence alignment                              */
   int           fmt          = eslMSAFILE_UNKNOWN;  /* format code for alifile                                      */
   char         *hmmfile      = NULL;                /* hmm input file path                                          */
   P7_HMMFILE   *hfp;                                /* input hmm file                                               */
   P7_HMM       *hmm;                                /* input hmm                                                    */
   char         *hpm_outfile;                        /* hpm output file path                                         */
   FILE         *hpm_outfp    = NULL;                /* hpm output file object                                       */
   HPM          *hpm;                                /* hidden potts model                                           */
   POTTS        *potts        = NULL;                /* potts model                                                  */
   int          *ct           = NULL;                /* [0..alen] base pair partners array for SS_cons               */
   double      **abc_ct       = NULL;                /* [0..msa->alen-1][0..abc->K] number of each residue at each   *
                                                      * position (abc->K is gap)                                     */
   double    ***bp_ct         = NULL;                /* [0..msa->alen-1][0..msa->alen-1][0..abc->K][0..abc->K]       *
                                                      * number of each combination of residues at all pairs of sites */
   int           use_weights  = 0;                   /* use PB weights when calculating MSA frequencies              */
   int           pk           = 0;                   /* leep track of pseudoknotted basepairs                        */
   int           status;                             /* easel return code                                            */
   char          errbuf[eslERRBUFSIZE];              /* for error messages                                           */

   /* parse command line */
   go = esl_getopts_Create(options);
   if (esl_opt_ProcessCmdline(go, argc, argv) != eslOK) cmdline_failure(argv[0], "Failed to parse command line: %s\n", go->errbuf);
   if (esl_opt_VerifyConfig(go)               != eslOK) cmdline_failure(argv[0], "Error in app configuration:   %s\n", go->errbuf);

   if (esl_opt_GetBoolean(go, "-h") ) {
      esl_banner(stdout, argv[0], banner);
      esl_usage (stdout, argv[0], usage);
      puts("\n where options are:");
      esl_opt_DisplayHelp(stdout, go, 1, 2, 80);
      puts("\n Alphabet options:");
      esl_opt_DisplayHelp(stdout, go, 2, 2, 80);
      puts("\n Training options:");
      esl_opt_DisplayHelp(stdout, go, 3, 2, 80);
      exit(0);
   }

   if (esl_opt_ArgNumber(go) != 3) cmdline_failure(argv[0], "Incorrect number of command line arguments.\n", go->errbuf);

   /* read arguments */
   hpm_outfile = esl_opt_GetArg(go, 1);
   hmmfile     = esl_opt_GetArg(go, 2);
   alifile     = esl_opt_GetArg(go, 3);

   /* if user has defined an alphabet we define it here */
   if      (esl_opt_GetBoolean(go, "--amino")) abc = esl_alphabet_Create(eslAMINO);
   else if (esl_opt_GetBoolean(go, "--rna"))   abc = esl_alphabet_Create(eslRNA);
   else if (esl_opt_GetBoolean(go, "--dna"))   abc = esl_alphabet_Create(eslDNA);

   /* check options for training hpm */
   if (esl_opt_GetBoolean(go, "--pknot"))  pk          = 1;
   if (esl_opt_GetBoolean(go, "--weight")) use_weights = 1;

   /* Open the .hmm file */
   status = p7_hmmfile_OpenE(hmmfile, NULL, &hfp, errbuf);
   if      (status == eslENOTFOUND) esl_fatal("File existence/permissions problem in trying to open HMM file %s.\n%s\n", hmmfile, errbuf);
   else if (status == eslEFORMAT)   esl_fatal("File format problem in trying to open HMM file %s.\n%s\n",                hmmfile, errbuf);
   else if (status != eslOK)        esl_fatal("Unexpected error %d in opening HMM file %s.\n%s\n",               status, hmmfile, errbuf);

   /* read first hmm  from hmm file */
   if (p7_hmmfile_Read(hfp, &abc, &hmm) != eslOK) esl_fatal("Failed to read HMM");

   /* open the alignment file */
   if ( (status = esl_msafile_Open(&abc, alifile, NULL, fmt, NULL, &afp)) != eslOK) esl_msafile_OpenFailure(afp, status);

   /* read first msa from alignment file */
   if ( (status = esl_msafile_Read(afp, &msa)) != eslOK) esl_msafile_ReadFailure(afp, status);

   /* calculate Henikoff PB weights from MSA */
   if ( (status = esl_msaweight_PB(msa)) != eslOK) esl_fatal("esl_msaweight_PB() failed, returned status %d\n", status);

   /* trim msa to consensus */
   if ( (status = trim_msa_rf(msa, abc, errbuf)) != eslOK) esl_fatal("trim_msa_rf() failed, returned status %d\n%s\n", status,errbuf);

   /* find annotated basepairs */
   if ( (status = find_basepairs(msa, pk, &ct, errbuf)) != eslOK) esl_fatal("find_basepairs() failed, returned status %d\n%s\n", status, errbuf);

   /* calculate alignment frequencies */
   if ( (status = count_msa(msa, ct, use_weights, &abc_ct, &bp_ct, errbuf)) != eslOK) esl_fatal("count_msa() failed, returned status %d\n%s\n", status, errbuf);

   /* Convert frequencies to Potts model */
   if ( (counts_to_potts(msa, abc_ct, bp_ct, ct, &potts)) != eslOK) esl_fatal("counts_to_potts() failed, returned status %d\n", status);

   /* combine potts and HMM to produce HPM */
   if ( (hpm = hpm_Create_hmm_potts(hmm, potts,abc)) == NULL) esl_fatal("hpm_Create_hmm_potts() failed\n");

   /* open hpm outfile for writing, write hpm to it */
   if ( (hpm_outfp = fopen(hpm_outfile, "w")) == NULL) esl_fatal("Failed to open output hpm file %s for writing\n", hpm_outfile);
   if ( (status = hpmfile_Write(hpm_outfp, hpm)) != eslOK) esl_fatal("hpmfile_Write() failed, returned status %d\n", status);

   /* close hpm outfile */
   fclose(hpm_outfp);

   /* clean up and return */
   free(ct);
   esl_arr2_Destroy((void **)  abc_ct, msa->alen);
   esl_arr3_Destroy((void ***) bp_ct,  msa->alen, abc->K+1);
   esl_msafile_Close(afp);
   esl_msa_Destroy(msa);
   p7_hmmfile_Close(hfp);
   hpm_Destroy(hpm);
   p7_hmm_Destroy(hmm);
   potts_Destroy(potts);
   esl_alphabet_Destroy(abc);
   esl_getopts_Destroy(go);

   return 0;
}

/* Function: trim_msa_rf()
 * Synopsis: Trim <msa> to only its consensus columns
 *
 * args:     msa    - alignment to be trimmed
 *           abc    - alphabet
 *           errbuf - for error messages
 *
 * Returns:  eslOK on success, <msa> is trimmed
 */

static int trim_msa_rf(ESL_MSA *msa, ESL_ALPHABET *abc, char *errbuf) {
   int  status;       /* esl return code                    */
   int  rflen = 0;    /* number of consensus columnn        */
   int  apos  = 0;    /* alignment position index           */
   int *useme = NULL; /* mask for trimming MSA to consensus */

   ESL_ALLOC(useme, sizeof(int) * msa->alen);

   /* contract check */
   if(msa->rf == NULL) ESL_FAIL(eslEINVAL, errbuf, "Error, trying to map RF positions to alignment positions, but msa->rf is NULL.");

   /* loop over alignment columns, see which are annotated match columns */
   for (apos = 0; apos < msa->alen; apos++) {
      if ( (! esl_abc_CIsGap(abc, msa->rf[apos]))   &&
         (! esl_abc_CIsMissing(abc, msa->rf[apos])) &&
         (! esl_abc_CIsNonresidue(abc, msa->rf[apos]))) {

         rflen++;
         useme[apos] = TRUE;
      }
      else {
         useme[apos] = FALSE;
      }
   }

   /* apply rf mask to MSA */
   if ((status = esl_msa_ColumnSubset (msa, errbuf, useme)) != eslOK) esl_fatal(errbuf);

   /* clean up and return */
   free(useme);
   return eslOK;

	ERROR:
		return status;
}

/* Function:  find basepairs()
 * Synopsis:  locate the annotated basepairs in <msa>
 *
 * args:     msa    - alignment w/ SS_cons line
 *           pk     - boolean to consider annotated pseudoknots
 *           ret_pk - [1...msa-alen] array, to be returned
 *                    ret_ct[i] = annotated basepair of column i,
 *                    else 0 (CT format)
 *           errbuf - for error messages
 *
 * Returns:  eslOK on success, basepairs in <ret_pk>
 */

static int find_basepairs(ESL_MSA *msa, int pk, int **ret_ct, char *errbuf)
{
   int  *ct          = NULL;  /* [0..alen] base pair partners array for SS_cons */
   char *ss_nopseudo = NULL;  /* no-pseudoknot version of structure             */
   int   status;              /* esl return code                                */

   /* find annotated basepairs */
   ESL_ALLOC(ct, sizeof(int) * (msa->alen+1));

   /* if we keep track of pseudoknots, read ss_cons as is */
   if (pk) {
      if ((status = esl_wuss2ct(msa->ss_cons, msa->alen, ct)) != eslOK) ESL_XFAIL(status, errbuf, "Consensus structure string is inconsistent.");
   }

   /* if no pseudoknots, then remove them from ss_cons */
   else {
      ESL_ALLOC(ss_nopseudo, sizeof(char) * (msa->alen+1));
      esl_wuss_nopseudo(msa->ss_cons, ss_nopseudo);
      if ((status = esl_wuss2ct(ss_nopseudo, msa->alen, ct)) != eslOK) ESL_XFAIL(status, errbuf, "Consensus structure string is inconsistent.");
   }

   /* clean up and return */
   *ret_ct = ct;
   free(ss_nopseudo);
   return eslOK;

   ERROR:
   return status;
}

/* Function: count_msa()
 * Synopsis: count observed residue frequencies in <msa>,
 *           both single site and pairwise for annotated
 *           basepairs
 *
 * args:     msa        - alignment to be analyzed
 *           ct         - CT array with annotated basepairs
 *           useweights - boolean to use seq weights in
 *                        msa->wgt to weight counts
 *           ret_abc_ct - [0...msa->alen-1] array of single-site
 *                        counts,
 *           ret_bp_ct  - [0...msa->alen-1] array pairwise resdue
 *                        counts, only for annotqted bp. For a bp,
 *                        the 5'(left) site contains the counts;
 *                        if i is  unpaired or a 3's site,
 *                        ret_bp_ct is unallocated.
 *
 * Returns:  eslOK on success, counts in <ret_abc_ct> and
 *           <ret_bp_ct>
 */

static int count_msa(ESL_MSA *msa, int *ct, int use_weights, double ***ret_abc_ct, double ****ret_bp_ct, char *errbuf)
{
   double  **abc_ct = NULL; /* single-site residue counts */
   double ***bp_ct  = NULL; /* basepair residue counts    */
   int       n;             /* sequence (row) index       */
   int       i,j;           /* position (column) indices  */
   int       a,b;           /* alphabet indices           */
   double    seqwt;         /* weight of current sequence */
   int       status;        /* esl return code            */

   if (! (msa->flags & eslMSA_DIGITAL)) ESL_FAIL(eslEINVAL,    errbuf, "count_msa() contract violation, MSA is not digitized");
   if (use_weights && msa->wgt == NULL) ESL_FAIL(eslEINCOMPAT, errbuf, "count_msa(): use_weights==TRUE but msa->wgt == NULL");

   /* allocate memory for count arrays */
   ESL_ALLOC(abc_ct, sizeof(double *)  * msa->alen);
   ESL_ALLOC(bp_ct,  sizeof(double **) * msa->alen);

   for(i = 0; i < msa->alen; i++) {
      ESL_ALLOC(abc_ct[i], sizeof(double) * (msa->abc->K+1));
      esl_vec_DSet(abc_ct[i], (msa->abc->K+1), 0.);

      /* if we have the 5' end of a basepair, allocate memory for pairwise counts */
      if (ct[(i+1)] > (i+1)) {
         ESL_ALLOC(bp_ct[i], sizeof(double *) * (msa->abc->K+1));
         for (a = 0; a < msa->abc->K+1; a++) {
            ESL_ALLOC(bp_ct[i][a], sizeof(double) * (msa->abc->K+1));
            esl_vec_DSet(bp_ct[i][a], msa->abc->K+1, 0.);
         }
      }
      else {
         bp_ct[i] = NULL;
      }
   }

   /* loop over sequences in MSA */
   for (n = 0; n < msa->nseq; n++) {
      /* determine if we are using sequence weights */
      seqwt = use_weights ? msa->wgt[n] : 1.0;

      /* update appropriate abc count, careful, ax ranges from 1..msa->alen *
       * * (but abc_ct is 0..msa->alen-1)                                   */
      for (i = 0; i < msa->alen; i++) {
         /* count canonical characters */

         if   (msa->ax[n][i+1] < msa->abc->K) a = msa->ax[n][i+1];
         /* count gaps and degenerate characters as gaps */
         else a = msa->abc->K;
         abc_ct[i][a] += seqwt;

         /* handle basepair counts (if this position is a 5' bp position) */
         if (bp_ct[i] != NULL) { /* our flag for whether position (i+1) is an 'i' in an i:j pair where i < j */
            j = ct[i+1] - 1; /* ct is indexed 1..alen */

            if   (msa->ax[n][j+1] < msa->abc->K) b = msa->ax[n][j+1];
            else b =  msa->abc->K;
            bp_ct[i][a][b] += seqwt;
         }
      }
   }

   /* clean up and return */
   *ret_abc_ct = abc_ct;
   *ret_bp_ct  = bp_ct;
   return eslOK;

   ERROR:
      return status;
}

/* Function:  counts_to_potts()
 * Synopsis:  create <potts> using single site and pairwise residue
 *            conts from <msa>
 *
 * args:     msa       - Training alignment
 *           abc_ct    - Single site residue counts in <msa>
 *           bp_ct     - Pairwise residue count for annotated bp in <msa>
 *           ct        - CT-array marking annotated bp in <msa>
 *           ret_potts - POTTS model to return
 *
 * Returns:  eslOK on success, POTTS model in <ret_potts>
 */

static int counts_to_potts(ESL_MSA *msa,  double **abc_ct, double ***bp_ct, int *ct, POTTS **ret_potts)
{
   int     i,j,k,rpos;                                       /* MSA poistion (column) indices */
   int     a,b;                                              /* character indices             */
   int     Kg          = msa->abc->K +1;                     /* size of alphabet w/ gap char  */
   POTTS *potts        =  potts_Create(msa->alen, msa->abc); /* potts model                   */

   /* loop over msa positions */
   for (i = 0; i < msa->alen; i++) {
      rpos = ct[i+1];
      k    = rpos - 1;

      /* if we have an unpaired position, set h_i's */
      if (rpos == 0) {
         for (a = 0; a < Kg; a++){
            potts->h[i][a] = (double) log(abc_ct[i][a] + 1.0);
         }
         esl_vec_DLogNorm(potts->h[i], Kg);
         esl_vec_DLog(potts->h[i], Kg);

         /* set all e_ij's equal */
         for (j = 0; j < msa->alen; j++) {
            esl_vec_DSet(potts->e[i][j], Kg*Kg, 0.0);
         }
      }

      else {
         /* set all h_i's equal */
         esl_vec_DSet(potts->h[i], Kg, 0.0);

         /* set all e_ij's to zero, unless we have an annotated BP */
         for (j = 0; j < msa->alen; j++) {
            esl_vec_DSet(potts->e[i][j], Kg*Kg, 0.0);
            if (j == k) {
               for (a = 0; a < Kg; a++) {
                  for (b = 0; b < Kg; b++) {
                     /* i is 3' end of BP */
                     if (j > i) potts->e[i][j][Kg*a+b] = log(bp_ct[i][a][b] + 1.0);
                     /* i is 5' end of BP */
                     else       potts->e[i][j][Kg*a+b] = log(bp_ct[j][b][a] + 1.0);
                  }
               }

               esl_vec_DLogNorm(potts->e[i][j], Kg*Kg);
               esl_vec_DLog(potts->e[i][j], Kg*Kg);
            }
         }
      }
   }

   *ret_potts = potts;

   /* clean up and return */
   return eslOK;
}
```
