## Supplemental Code S1 for "Remote homology search with hidden Potts models": hpmemit.c

```
/* hpmemit.c */
#include "p7_config.h"

#include <stdio.h>
#include <stdlib.h>
#include <string.h>
#include <math.h>

#include "easel.h"
#include "esl_alphabet.h"
#include "esl_getopts.h"
#include "esl_msa.h"
#include "esl_msafile.h"
#include "esl_random.h"
#include "esl_sq.h"

#include "hmmer.h"

#include "hpm.h"
#include "hpmfile.h"
#include "hpm_scoreops.h"

static ESL_OPTIONS options[] = {
   /* name       type            default env   range  togs  reqs incomp            help                                                                           docgroup */
   { "-h",       eslARG_NONE,    FALSE,  NULL, NULL,  NULL, NULL, NULL,            "help; show brief info on version and usage",                                         0 },
   { "-s",       eslARG_INT,     "0",    NULL, NULL,  NULL, NULL, NULL,            "set random number seed to <n>",                                                      0 },

   /* options controlling sequence emission */
   { "-N",       eslARG_INT,     "1",    NULL, "n>0", NULL, NULL, NULL,            "number of seqs to sample",                                                           1 },
   { "-L",       eslARG_INT,     "100",  NULL, "n>0", NULL, NULL, NULL,            "set expected length from profile to <l> (for N and C state transition probs)",       1 },
   { "--Niter",  eslARG_INT,     "1000", NULL, "n>0", NULL, NULL, NULL,            "number of MCMC iterations per sequence",                                             1 },

   /* Options forcing which alphabet we're working in (normally autodetected) */
   { "--amino",  eslARG_NONE,    FALSE,  NULL, NULL,  NULL, NULL, "--dna,--rna",   "<seqfile> contains protein sequences",                                               2 },
   { "--rna",    eslARG_NONE,    FALSE,  NULL, NULL,  NULL, NULL, "--dna,--amino", "<seqfile> contains RNA sequences",                                                   2 },
   { "--dna",    eslARG_NONE,    FALSE,  NULL, NULL,  NULL, NULL, "--rna,--amino", "<seqfile> contains DNA sequences",                                                   2 },

   /* debugging tools */
   { "-v",       eslARG_NONE,    FALSE,  NULL, NULL,  NULL, NULL, NULL,            "Verbose mode: print info on intermediate emission steps",                            3 },
   { "--burnin", eslARG_OUTFILE, FALSE,  NULL, NULL,  NULL, NULL, NULL,            "(TBD) send csv of Potts hamiltonian vs MCMC iteration (first sequence only) to <f>", 3 },
   { 0,0,0,0,0,0,0,0,0,0 },
};

static char usage[]   = "[-options] <hpmfile> <msaoutfile>";
static char banner[]  = "sample aligned sequence(s) from an HPM";

/* declaration of internal functions */
static void emit_alignment(ESL_GETOPTS *go, FILE *ofp, int outfmt, ESL_RANDOMNESS *rng, HPM *hpm, int v);
int         emit_match_MCMC(ESL_SQ *msq, HPM *hpm, int Niter, ESL_RANDOMNESS *rng, int v);
int         sample_trace(int L, P7_TRACE *tr, ESL_SQ *msq, HPM *hpm,  ESL_RANDOMNESS *rng, int v);
int         emit_inserts(P7_TRACE *tr, ESL_SQ *sq, ESL_SQ *msq, HPM *hpm,  ESL_RANDOMNESS *rng, int v);

static void
cmdline_failure(char *argv0, char *format, ...)
{
   va_list argp;
   va_start(argp, format);
   vfprintf(stderr, format, argp);
   va_end(argp);
   esl_usage(stdout, argv0, usage);
   printf("\nTo see more help on available options, do %s -h\n\n", argv0);
   exit(1);
}

int main(int argc, char *argv[]) {
   ESL_GETOPTS    *go;                                            /* cmd line configuration  */
   ESL_ALPHABET   *abc                    = NULL;                 /* sequence alphabet       */
   ESL_RANDOMNESS *rng                    = NULL;                 /* random number generator */
   char           *hpmfile                = NULL;                 /* input HPM filepath      */
   HPM            *hpm                    = NULL;                 /* input HPM               */
   int             outfmt                 = eslMSAFILE_STOCKHOLM; /* output MSA format       */
   FILE           *ofp                    = NULL;                 /* output MSA filestream   */
   int             v                      = 0;                    /* verbose mode indicator  */
   char            errbuf[eslERRBUFSIZE];                         /* for error messages      */

   /* parse command line */
   go = esl_getopts_Create(options);
   if (esl_opt_ProcessCmdline(go, argc, argv) != eslOK) cmdline_failure(argv[0], "Failed to parse command line: %s\n", go->errbuf);
   if (esl_opt_VerifyConfig(go)               != eslOK) cmdline_failure(argv[0], "Error in app configuration:   %s\n", go->errbuf);

   if (esl_opt_GetBoolean(go, "-h") ) {
      esl_banner(stdout, argv[0], banner);
      esl_usage (stdout, argv[0], usage);
      puts("\n where options are:");
      esl_opt_DisplayHelp(stdout, go, 1, 2, 80);
      puts("\n Emission options:");
      esl_opt_DisplayHelp(stdout, go, 2, 2, 80);
      puts("\n Alphabet options:");
      esl_opt_DisplayHelp(stdout, go, 3, 2, 80);
      puts("\n Debug options:");
      esl_opt_DisplayHelp(stdout, go, 4, 2, 80);
      exit(0);
   }

   /* read arguments */
   if (esl_opt_ArgNumber(go) != 2) cmdline_failure(argv[0], "Incorrect number of command line arguments.\n", go->errbuf);

   hpmfile = esl_opt_GetArg(go, 1);
   if ((ofp = fopen(esl_opt_GetArg(go, 2), "w")) == NULL) esl_fatal("Failed to open output file %s", esl_opt_GetArg(go, 2));

   /* if user has defined an alphabet we define it here */
   if      (esl_opt_GetBoolean(go, "--amino")) abc = esl_alphabet_Create(eslAMINO);
   else if (esl_opt_GetBoolean(go, "--dna"))   abc = esl_alphabet_Create(eslDNA);
   else                                        abc = esl_alphabet_Create(eslRNA);

   /* check for verbose mode */
   if (esl_opt_GetBoolean(go, "-v")) v = 1;

   /* create random number generator */
   rng = esl_randomness_Create(esl_opt_GetInteger(go, "-s"));

   /* open HPM file */
   hpm = hpmfile_Read(hpmfile, abc, errbuf);

   /* emit an alignment! */
   emit_alignment(go, ofp, outfmt, rng, hpm, v);

   /* clean up and return */
   fclose(ofp);
   hpm_Destroy(hpm);
   esl_randomness_Destroy(rng);
   esl_alphabet_Destroy(abc);
   esl_getopts_Destroy(go);
   return 0;
}


static void emit_alignment(ESL_GETOPTS *go, FILE *ofp, int outfmt, ESL_RANDOMNESS *rng, HPM *hpm, int v)
{
   ESL_MSA   *msa       = NULL;                              /* emitted MSA                   */
   ESL_SQ   **sq        = NULL;                              /* emitted seq array             */
   ESL_SQ    *msq       = NULL;                              /* holds match residues          */
   P7_TRACE **tr        = NULL;                              /* trace array for emitted seqs  */
   int        N         = esl_opt_GetInteger(go, "-N");      /* number of seqs to emit        */
   int        L         = esl_opt_GetInteger(go, "-L");      /* target sequence length        */
   int        Niter     = esl_opt_GetInteger(go, "--Niter"); /* # of MCMC iterations per seq  */
   int        optflags  = p7_ALL_CONSENSUS_COLS;             /* output MSA options            */
   int        i,j;                                           /* sequence and position indices */

   if (v) fprintf(stdout, "in emit_alignment()\n");

   /* open burn-in file */
   /* TO BE ADDED */

   /* allocate space for trace and seq arrays */
   if ((tr = malloc(sizeof(P7_TRACE *) * N)) == NULL) esl_fatal("failed to allocate trace array");
   if ((sq = malloc(sizeof(ESL_SQ   *) * N))  == NULL) esl_fatal("failed to allocate seq array");
   msq =  esl_sq_CreateDigital(hpm->abc);

   /* main loop: initialize and generate traces + seqs */
   for (i = 0; i < N; i++)
   {
      if( i % 1000 == 0) fprintf(stdout, "emitting sequence %d\n", i);


      if ((sq[i] = esl_sq_CreateDigital(hpm->abc)) == NULL) esl_fatal("failed to allocate seq");
      if ((tr[i] = p7_trace_Create())              == NULL) esl_fatal("failed to allocate trace");

      /* PASS BURN-IN FILE */
      /* emit match states from Potts prob dist via MCMC */
      emit_match_MCMC(msq, hpm, Niter, rng, v);

      /* generate trace */
      /* pass match sequence to this function */
      sample_trace(L, tr[i], msq, hpm, rng, v);

      /* emit insert characters */
      emit_inserts(tr[i], sq[i], msq, hpm, rng, v);

      /* print sequence info */
      if (v) {
         fprintf(stdout, "elements of sequence i=%d\n", i);
         for (j=0; j < sq[i]->n+1; j++) {
            fprintf(stdout, "\ti: %d, sq[i]: %d\n",
                    j, sq[i]->dsq[j]);
         }
      }

      /* set sequence name */
      if (esl_sq_FormatName(sq[i], "MCMC-sample%d", i+1) != eslOK) esl_fatal("Failed to set sequence name\n");
      esl_sq_Reuse(msq);
   }

   /* write burn-in file */
   /* TO BE ADDED */

   /* Write MSA to outfile */
   p7_tracealign_Seqs(sq, tr, N, hpm->M, optflags, NULL, &msa);
   esl_msafile_Write(ofp, msa, outfmt);

   /* clean up and return */
   for (i = 0; i < N; i++) p7_trace_Destroy(tr[i]);
   for (i = 0; i < N; i++) esl_sq_Destroy(sq[i]);
   esl_sq_Destroy(msq);
   free(tr);
   free(sq);
   esl_msa_Destroy(msa);
   return;
}


/* modify to accept burn-in file */

int emit_match_MCMC(ESL_SQ *sq, HPM *hpm, int Niter, ESL_RANDOMNESS *rng, int v)
{
   int       k;         /* node index                          */
   int       i;         /* sequence index                      */
   int       x;         /* sampled character                   */
   int       n;         /* MCMC iteration index                */
   float     hsc,esc;   /* Potts Hamiltonian params            */
   float     Ei,Ei_tmp; /* Potts Hamiltonian params            */
   float     deltaE;    /* Potts Hamiltonian params            */
   P7_TRACE *tr;        /* faux trace w/ no insert states      */
   ESL_SQ   *sq_tmp;    /* changed seq in algorithm            */
   float     p,r;       /* for accepting/rejecting seq changes */
   int       status;    /* esl return code                     */

   sq_tmp = esl_sq_CreateDigital(hpm->abc);

   /* generate special states of match-only trace */
   if ((tr = p7_trace_Create()) == NULL) esl_fatal("failed to allocate trace");
   if ((status = p7_trace_Append(tr, p7T_S, 0, 0)) != eslOK) goto ERROR;
   if ((status = p7_trace_Append(tr, p7T_N, 0, 0)) != eslOK) goto ERROR;
   if ((status = p7_trace_Append(tr, p7T_B, 0, 0)) != eslOK) goto ERROR;
   if ((status = p7_trace_Append(tr, p7T_G, 0, 0)) != eslOK) goto ERROR;

   /* initialize sequence and add match states to match-only trace*/
   for (k = 1; k < hpm->M+1; k++) {

      /* add match state to trace */
      if ((status = p7_trace_Append(tr, p7T_MG, k, k)) != eslOK) goto ERROR;
      esl_rnd_Deal(rng, 1, hpm->abc->K, &x);

      //fprintf(stdout, "sampled residue: %d\n", x);
      if ((status = esl_sq_XAddResidue(sq, x)) != eslOK) goto ERROR;
   }

   /* add C-terminus special states to trace */
   if ((status = p7_trace_Append(tr, p7T_E, 0, 0)) != eslOK) goto ERROR;
   if ((status = p7_trace_Append(tr, p7T_C, 0, 0)) != eslOK) goto ERROR;
   if ((status = p7_trace_Append(tr, p7T_T, 0, 0)) != eslOK) goto ERROR;

   if (v) p7_trace_Dump(stdout, tr);
   /* score initial match sequence */
   if (v) {
      hpm_scoreops_CalculateHamiltonian(hpm, tr, sq->dsq, &hsc, &esc);
      fprintf(stdout, "initial h score: %.4f\n", hsc);
      fprintf(stdout, "initial e score: %.4f\n", esc);
   }

   /* run Metropolis Algorithm */
   for (n = 0; n < Niter; n++) {
      esl_sq_Copy(sq, sq_tmp);

      if (v) fprintf(stdout, "\titeration n=%d\n", n);

      /* pick a node at random */
      esl_rnd_Deal(rng, 1, hpm->M, &k);

      /* want match states to start counting at 1 */
      k += 1;
      if (v) fprintf(stdout, "\tnode picked: %d\n", k);

      /* pick a new residue x_k at random */
      esl_rnd_Deal(rng, 1, hpm->abc->K+1, &x);
      if (v) fprintf(stdout, "\tresidue picked: %d\n", x);

      /* change residue in temporary sequence */
      sq_tmp->dsq[k] = x;

      if (v) {
         for(i=1;i<hpm->M+1;i++) {
            fprintf(stdout, "\t\ti: %d; sq[i]: %d; sq_tmp[i]: %d\n",
                    i, sq->dsq[i], sq_tmp->dsq[i]);
         }
      }

      /* calculate change in hamiltonian */
      if ((status = hpm_scoreops_CalculateHamiltonianSingleSite(hpm, tr, sq->dsq, k, &Ei)) != eslOK) goto ERROR;
      if ((status = hpm_scoreops_CalculateHamiltonianSingleSite(hpm, tr, sq_tmp->dsq, k, &Ei_tmp)) != eslOK) goto ERROR;
      deltaE = Ei_tmp - Ei;

      if (v) fprintf(stdout, "\tEi: %.4f; Ei_tmp: %.4f; deltaE: %.4f\n",
                     Ei, Ei_tmp, deltaE);

      /* accept/reject change */
      if (deltaE >= 0) {
         if (v) fprintf(stdout, "\tEnergy same/higher, change accepted!\n");
         esl_sq_Reuse(sq);
         esl_sq_Copy(sq_tmp, sq);
      }

      /* if energy is lower, accept w/ prob $e^{\Delta E}$ */
      else {
         p = exp(deltaE);
         if (v) fprintf(stdout, "\tp = %.4f\n", p);
         r = esl_random(rng);
         if (v) fprintf(stdout, "\tr = %.4f\n", r);
         if (p > r) {
            if (v) fprintf(stdout, "\ttrial passed, change accepted!\n");
            esl_sq_Reuse(sq);
            esl_sq_Copy(sq_tmp, sq);
         }
      }

      //if (n % 100 == 0) {
      // hpm_scoreops_CalculateHamiltonian(hpm, tr, sq->dsq, &hsc, &esc);
      // fprintf(stdout, "%d,%.4f\n", n, hsc + esc);
      //}

      esl_sq_Reuse(sq_tmp);
   }

   if (v) {
      hpm_scoreops_CalculateHamiltonian(hpm, tr, sq->dsq, &hsc, &esc);
      fprintf(stdout, "final h score: %.4f\n", hsc);
      fprintf(stdout, "final e score: %.4f\n", esc);
   }

   /* clean up and return */
   esl_sq_Destroy(sq_tmp);
   p7_trace_Destroy(tr);
   return eslOK;

   ERROR:
      return status;
}

int sample_trace(int L, P7_TRACE *tr, ESL_SQ *msq, HPM *hpm, ESL_RANDOMNESS *rng, int v)
{
   int    k            = 0;      /* position in model nodes 1..M                     */
   int    i            = 0;      /* position in sequence 1..L                        */
   char   st           = p7T_N;  /* state type                                       */
   char   prev         = p7T_S;  /* previous state type                              */
   int    showi;                 /* do we emit residue from N/C state                */
   int    kend         = hpm->M; /* predestined end node                             */
   float  ploop,pmove;           /* transition parameters for N and C states         */
   float *xt;                    /* array of special state (N,C) transition porbs    */
   int    nxt          = 2;      /* number of special state transition possibilities */
   int    status;                /* easel return code                                */

   /* allocate memory for special state transition parameters */
   ESL_ALLOC(xt, nxt * sizeof(float));

   if (tr) {
      if ((status = p7_trace_Reuse(tr))               != eslOK) goto ERROR;
      if ((status = p7_trace_Append(tr, p7T_S, 0, 0)) != eslOK) goto ERROR;
      if ((status = p7_trace_Append(tr, p7T_N, 0, 0)) != eslOK) goto ERROR;
   }

   /* set N and C transition parameters */
   pmove = (2.0f) / ( (float) L + 2.0f);
   ploop = 1.0 - pmove;
   xt[0] = ploop;
   xt[1] = pmove;

   esl_rnd_FChoose(rng, xt, nxt);

   if (v) fprintf(stdout, "pmove: %.4f\n", pmove);
   if (v) fprintf(stdout, "ploop: %.4f\n", ploop);

   while (st != p7T_T)
   {
      if (v) fprintf(stdout, "Current state: %d\n", st);
      if (v) fprintf(stdout, "Previous state: %d\n", prev);
      prev = st;

      switch (st) {

         /* case 1: N state */
         case (p7T_N):

            if (v) fprintf(stdout, "Currently in N state!\n");
            if   (esl_rnd_FChoose(rng, xt, nxt) == 0) {
               st = p7T_N;
               i++;
            }

            else {
               st = p7T_G;
            }
            break;

         /* case 2: G state */
         case (p7T_G):
            if (esl_abc_XIsGap(hpm->abc, msq->dsq[k+1])) {
               st = p7T_DG;
            }
            else {
               st = p7T_MG;
               i++;
            }
            break;

         /* case 3: MG */
         case (p7T_MG):
            if (k == kend) {
               st = p7T_E;
            }

            else {
               if (esl_rnd_DChoose(rng, HPM_TMAT(hpm, k), HPM_NTMAT) == 0) {
                  if (esl_abc_XIsGap(hpm->abc, msq->dsq[k+1])) {
                     st = p7T_DG;
                  }
                  else {
                     st = p7T_MG;
                     i++;
                  }
               }
               else {
                  st = p7T_IG;
                  i++;
               }
            }
            break;


         /* case 4: DG */
         case (p7T_DG):
            if (k == kend) {
               st = p7T_E;
            }
            else {
               if (esl_rnd_DChoose(rng, HPM_TMAT(hpm, k), HPM_NTMAT) == 0) {
                  if (esl_abc_XIsGap(hpm->abc, msq->dsq[k+1])) {
                     st = p7T_DG;
                  }
                  else {
                     st = p7T_MG;
                     i++;
                  }
               }
               else {
                  st = p7T_IG;
                  i++;
               }
            }
            break;

         /* case 5: IG */
         case (p7T_IG):
            if (esl_rnd_DChoose(rng, HPM_TINS(hpm, k),HPM_NTINS) == 0) {
               if (esl_abc_XIsGap(hpm->abc, msq->dsq[k+1])) {
                  st = p7T_DG;
               }
               else {
                  st = p7T_MG;
                  i++;
               }
            }
            else {
               st = p7T_IG;
               i++;
            }
            break;

         /* case 6: E */
         case(p7T_E):
            st = p7T_C;
            break;

         /* case 7: C */
         case(p7T_C):
            if (esl_rnd_FChoose(rng, xt, nxt) == 0) {
               st = p7T_C;
               i++;
            }
            else {
               st = p7T_T;
            }
            break;

         default: ESL_XEXCEPTION(eslEINCONCEIVABLE, "impossible state reached during emission");
      }

      /* based on state we just sampled, update k */
      if (st == p7T_MG || st == p7T_DG) k++;

      /* Add state to trace. */
      if (tr) {
         /* N and cstate only emits on trasmission */
         if (st == p7T_N || st == p7T_C) {
            if (prev == st) showi = i;
            else            showi = 0;
         }
         else if (st == p7T_G || st == p7T_E || st == p7T_T || st == p7T_DG) {
            showi = 0;
         }
         else if (st == p7T_MG || st == p7T_IG) {
            showi = i;
         }
         if ((status = p7_trace_Append(tr, st, k, showi)) != eslOK) goto ERROR;
      }
   }

   if (v) p7_trace_Dump(stdout, tr);
   free(xt);
   return eslOK;

   ERROR:
      return status;
}

int emit_inserts(P7_TRACE *tr, ESL_SQ *sq, ESL_SQ *msq, HPM *hpm, ESL_RANDOMNESS *rng, int v)
{
   int z;            /* trace index     */
   int x;            /* sampled residue */
   int prev = p7T_S; /* previous state  */
   int status;       /* esl return code */

   if (v) fprintf(stdout, "in function emit_inserts()\n");
   if (v) p7_trace_Dump(stdout, tr);

   /* loop thru trace */
   for (z = 0; z < tr->N; z++) {

      if (v) fprintf(stdout, "z: %d\t k: %d\t: i %d\t st: %d\n",
                     z, tr->k[z], tr->i[z], tr->st[z]);

      /* handle N and C states */
      if ((tr->st[z] == p7T_N || tr->st[z] == p7T_C) && tr->st[z] == prev) {

         /* choose insert residue */
         x = esl_rnd_DChoose(rng, hpm->ins[tr->k[z]], hpm->abc->K);
         if ((status = esl_sq_XAddResidue(sq, x)) != eslOK) goto ERROR;
         if (v) fprintf(stdout, "\tChosen residue: %d\n", x);
      }

      /* handle normal insert states */
      else if (tr->st[z] == p7T_IG) {

         /* choose insert residue */
         x = esl_rnd_DChoose(rng, hpm->ins[tr->k[z]], hpm->abc->K);
         if ((status = esl_sq_XAddResidue(sq, x)) != eslOK) goto ERROR;
         if (v) fprintf(stdout, "\tChosen residue: %d\n", x);
      }

      else if (tr->st[z] == p7T_MG) {
         if (v) fprintf(stdout, "we have a match state! node: %d, residue: %d\n",
                        tr->k[z], msq->dsq[tr->k[z]]);
         if ((status = esl_sq_XAddResidue(sq, msq->dsq[tr->k[z]])) != eslOK) goto ERROR;
      }
      prev = tr->st[z];
   }

   /* terminate sequence */
   if (sq && (status = esl_sq_XAddResidue(sq, eslDSQ_SENTINEL)) != eslOK) goto ERROR;

   return eslOK;

   ERROR:
      return status;
}
```
