## Supplemental Code S1 for "Remote homology search with hidden Potts models": hpmfile.c

```
#include <string.h>
#include <math.h>

#include "easel.h"
#include "esl_alphabet.h"

#include "hmmer.h"

#include "hpm.h"
#include "hpmfile.h"

/* declaration of internal functions */
static int printprob(FILE *fp, int fieldwidth, float p);

/* Function:  IsInt()
 * Synopsis:  Determine if string is an integer
 *
 * Purpose:   Determines if a string represents an integer using
 *            ASCII values. Can handle negative integers.
 *
 * Args:      str - string of interest
 *
 * Returns:   0 if <str> doesn't represent an int, 1 if it does
 */

int IsInt(char *str){
   int i;
   int n = strlen(str);

   for (i=0; i<n; i++) {

      /* check ASCII value of character */
      /* we allow '-' on the first character only */
      if (i == 0) {
         if ( (str[i] < 48 || str[i] > 57) && (str[i] != 45 || n == 1)) {
            return 0;
         }
      }
      else {
         if (str[i] < 48 || str[i] > 57) {
            return 0;
         }
      }
   }
   return 1;

}

/* Function:  hpmfile_Write()
 * Synopsis:  Write an HPM to a stream.
 *
 * Purpose:   Writes hidden Potts model <hpm> to open stream <fp>.
 *
 * Args:      fp   - open stream (such as <stdout>)
 *            hpm  - alignment to write
 *
 * Returns:   <eslOK> on success.
 *
 * Throws:    <eslEMEM> on allocation error.
 *            <eslEWRITE> on any system write error, such as a filled disk.
 */
int hpmfile_Write(FILE *fp, HPM *hpm) {
   int k,l;            /* model node indices        */
   int x,y;            /* character indices         */
   int status;         /* esl return code           */
   float compo = 0.05; /* fake value for compo line */

   fprintf(fp, ".hpm file\n");
   fprintf(fp, "LENG    %d\n", hpm->M);

   /* model parameters */

   /* denote 1st param section beginning with a '/' */
   fprintf(fp, "/\n");
   fprintf(fp, "HPM     ");

   for (x = 0; x < hpm->abc->K+1; x++) {
      fprintf(fp, "     %c   ", hpm->abc->sym[x]);
   }
   fprintf(fp, "\n");
   fprintf(fp, "        %8s %8s %8s %8s\n",
               "m->m", "m->i", "i->m", "i->i");

   /* need to add avg h_i values in compo line? */
   fprintf(fp, "  COMPO ");

   /* temporary fix to make this look like a .hmm file */
   for (x = 0; x < hpm->abc->K+1; x++) {
      if ( (status = printprob(fp, 8, compo)) != eslOK) return status;
	}
   if (fputc('\n', fp)         < 0) ESL_EXCEPTION_SYS(eslEWRITE, "hpm write failed");

   /* node 0 is special: insert emissions, and B-> transitions */
   if (fputs("        ", fp) < 0) ESL_EXCEPTION_SYS(eslEWRITE, "hpm write failed");
   for (x = 0; x < hpm->abc->K; x++) {
      if ( (status = printprob(fp, 8, hpm->ins[0][x])) != eslOK) return status;
   }

   // no insert emssions for a gap
   if ( (status = printprob(fp, 8, 0)) != eslOK) return status;

   if (fputc('\n', fp)       < 0) ESL_EXCEPTION_SYS(eslEWRITE, "hpm write failed");
   if (fputs("        ", fp) < 0) ESL_EXCEPTION_SYS(eslEWRITE, "hpm write failed");

   for (x = 0; x < hpm->nTransition; x++) {
      if ( (status = printprob(fp, 8, hpm->t[0][x])) != eslOK) return status;
   }

   if (fputc('\n', fp) < 0) ESL_EXCEPTION_SYS(eslEWRITE, "hpm write failed");

   for (k = 1; k < hpm->M+1; k++) {

      /* Line 1: k; hi's */
      if (fprintf(fp, " %6d ", k) < 0) ESL_EXCEPTION_SYS(eslEWRITE, "hpm write failed");
      for (x = 0; x < hpm->abc->K+1; x++) {
         fprintf(fp, " %*.6g", 8, hpm->h[k][x]);
      }
      if (fputc('\n', fp) < 0) ESL_EXCEPTION_SYS(eslEWRITE, "hpm write failed");

      /* Line 2: insert emissions  */
      if (fputs("         ", fp) < 0) ESL_EXCEPTION_SYS(eslEWRITE, "hpm write failed");
      for (x = 0; x < hpm->abc->K; x++) {
         if ( (status = printprob(fp, 8, hpm->ins[k][x])) != eslOK) return status;
      }
      // No insert emissions for gaps
      if ( (status = printprob(fp, 8, 0)) != eslOK) return status;
      if (fputc('\n', fp) < 0) ESL_EXCEPTION_SYS(eslEWRITE, "hpm write failed");

      /* Line 3: transitions */
      if (fputs("        ", fp) < 0) ESL_EXCEPTION_SYS(eslEWRITE, "hpm write failed");
      for (x = 0; x < hpm->nTransition; x++) {
         if ( (status = printprob(fp, 8, hpm->t[k][x])) != eslOK) return status;
      }
      if (fputc('\n', fp) < 0) ESL_EXCEPTION_SYS(eslEWRITE, "hpm write failed");
   }

   /* Write eij's */
   /* denote 2st param section beginning with a '/' */
   fprintf(fp, "/\n");

   for (k = 1; k < hpm->M+1; k++){
      for (l = k+1; l < hpm->M+1; l++) {

         //fprintf(fp, "%d %d\n", k, l);
         if (fprintf(fp, "%3d %3d ", k,l) < 0) ESL_EXCEPTION_SYS(eslEWRITE, "hpm write failed");
         for (x = 0; x < hpm->abc->K+1; x++) {
            fprintf(fp, "     %c   ", hpm->abc->sym[x]);
         }
         if (fputc('\n', fp) < 0) ESL_EXCEPTION_SYS(eslEWRITE, "hpm write failed");

         for (x = 0; x < hpm->abc->K+1; x++) {
            fprintf(fp, "    %c   ", hpm->abc->sym[x]);
            for (y = 0; y < hpm->abc->K+1; y++) {
               fprintf(fp, " %*.6g", 8, hpm->e[k][l][IDX(x,y,hpm->abc->K+1)]);
            }
            if (fputc('\n', fp) < 0) ESL_EXCEPTION_SYS(eslEWRITE, "hpm write failed");
         }
      }
   }

   /* end file with a '//' */
   if (fputs("//\n", fp) < 0) ESL_EXCEPTION_SYS(eslEWRITE, "hpm write failed");
   return eslOK;
}


/* Function:  hpmfile_Read()
 * Synopsis:  Read HPMfrom input
 *
 * Purpose:   Reads hidden Ootts model from <f>, returns it.
 *
 * Args:      f      - input file
 *            abc    - esl alphabet to assign to HPM
 *            errbuf - memory for error messages
 *
 * Returns:   HPM <ret_hpm> on sucess, NULL on failure
 */

HPM *
hpmfile_Read(char *f, ESL_ALPHABET *abc, char *errbuf) {

   ESL_FILEPARSER  *efp           = NULL;  /* easel fileparser object                                 */
   HPM             *ret_hpm       = NULL;  /* hpm to return                                           */
   char            *tok;                   /* file parser token                                       */
   char            *prev_tok      = NULL;  /* file parser token                                       */
   int              tok_count;             /* token count                                             */
   int              section_count = 0;     /* file section count                                      */
   int              M;                     /* number of nodes in model                                */
   int              i             = 0;     /* index for keeping track of match states                 */
   int              j             = 0;     /* another index for keeping track of match states         */
   int              l             = 0;     /* index for counting lines related to a given match state */
   int              a             = 0;     /* index for alphabet                                      */
   int              lc            = 1;     /* line counter                                            */
   int              status;                /* esl return code                                         */

   if (esl_fileparser_Open(f, NULL, &efp) != eslOK)  ESL_XFAIL(eslFAIL, errbuf, "HPM file %s open failed, returned status %s\n",
                                                               f, status);

   while (esl_fileparser_NextLine(efp) == eslOK) {
      lc ++;

      /* for parsing section 0 of the file */
      if (section_count == 0){
         tok_count = 0;
         while (esl_fileparser_GetTokenOnLine(efp, &tok, NULL) == eslOK) {

            /* extract model parameters */
            if (tok_count == 1 && strcmp(prev_tok, "LENG") == 0) M = atoi(tok);

            /*presumably extract more model parameters (e.g, name) once I add them */
            /* if (...) */

            /* see if we've reached the end of section 0 */
            if (tok_count == 0 && strcmp(tok, "/") == 0) {
               /* go ahead and create the hpm object */
               ret_hpm = hpm_Create(M, abc);
               section_count = 1;
            }

            prev_tok = tok;
            tok_count ++;
         }
      }

      /* for parsing section 1 of the model */
      /* should contain insert emissions, transition probs, and h_i's */
      else if (section_count == 1) {
         tok_count = 0;
         while (esl_fileparser_GetTokenOnLine(efp, &tok, NULL) == eslOK) {

            /* see if we've reached the end of section 0 */
            if (tok_count == 0 && strcmp(tok, "/") == 0) {
               section_count = 2;
               break;
            }

            /* skip the user-friendly column label lines */
            else if (strcmp(tok, "HPM") == 0 || strcmp(tok, "m->m") == 0) {
               l = -1;
               break;
            }

            /* parse node zero information */
            /* there are no node-zero hi's */
            else if (tok_count == 0 && strcmp(tok, "COMPO") == 0) {
               break;
            }

            /* read hi's */
            else if (l == 0) {
               if (tok_count > 0) {
                  ret_hpm->h[i][tok_count -1] = atof(tok);
               }
            }

            /* read insert emissions */
            else if (l == 1) {
               if (tok_count < abc->K) {
                  ret_hpm->ins[i][tok_count] = exp(-atof(tok));
               }
            }

            /* read transition probs */
            if (l == 2 ) {
               ret_hpm->t[i][tok_count] = exp(-atof(tok));
            }
            prev_tok = tok;
            tok_count ++;
         }
         if (l < 2) l++;

         /* move onto the next node */
	 else if (l == 2) {
            l = 0;
            i++;
         }
      }

      /* for parsing section 2 of the model */
      /* should contain e_{ij}'s */
      else if (section_count == 2) {
         tok_count = 0;
         prev_tok = NULL;
         while (esl_fileparser_GetTokenOnLine(efp, &tok, NULL) == eslOK) {

            /* check to see if we've reached the last line */
            if ( tok_count == 0 && strcmp(tok, "//") == 0) {
               break;
            }

            /* if the line starts with two integers, those are i and j */
            if ( (tok_count == 1) && (IsInt(tok)) && (IsInt(prev_tok))) {
               i = atoi(prev_tok);
               j = atoi(tok);
               a = -1;
               break;
            }

            else if (tok_count > 0) {
               ret_hpm->e[i][j][IDX(a,tok_count-1,abc->K+1)] = atof(tok);
               ret_hpm->e[j][i][IDX(tok_count-1,a,abc->K+1)] = atof(tok);
            }

            tok_count ++;
            prev_tok = tok;
         }
         a++;
      }
   }

   esl_fileparser_Close(efp);
   return ret_hpm;

   ERROR:
      return NULL;
}

/* function for writing probabilities in loge */
/* handles p = 0  and p = 1 specially         */
/* based on hpmfile.c::printprob()            */
static int
printprob(FILE *fp, int fieldwidth, float p) {
   if      (p == 0.0) { if (fprintf(fp, " %*s",   fieldwidth, "*")      < 0) ESL_EXCEPTION_SYS(eslEWRITE, "hmm write failed"); }
   else if (p == 1.0) { if (fprintf(fp, " %*.5f", fieldwidth, 0.0)      < 0) ESL_EXCEPTION_SYS(eslEWRITE, "hmm write failed"); }
   else               { if (fprintf(fp, " %*.5f", fieldwidth, -logf(p)) < 0) ESL_EXCEPTION_SYS(eslEWRITE, "hmm write failed"); }
   return eslOK;
}
```
