## Supplemental Code S1 for "Remote homology search with hidden Potts models": hpmfile.h

```
/* hpmfile.h */

#ifndef HPMFILE_INCLUDED
#define HPMFILE_INCLUDED

#include <stdio.h>

#include "easel.h"
#include "esl_alphabet.h"
#include "esl_fileparser.h"

#include "hpm.h"

/* hpmfile.c */
extern int  IsInt(char *str);
extern int  hpmfile_Write(FILE *f, HPM *hpm);
HPM        *hpmfile_Read(char *f, ESL_ALPHABET *abc, char *errbuff);
#endif
```
