## Supplemental Code S1 for "Remote homology search with hidden Potts models": hpmscore.c

```
#include <stdio.h>
#include <stdlib.h>
#include <string.h>
#include <math.h>

#include "easel.h"
#include "esl_alphabet.h"
#include "esl_msa.h"
#include "esl_sq.h"
#include "esl_sqio.h"
#include "esl_msafile.h"
#include "esl_getopts.h"
#include "esl_composition.h"

#include "p7_config.h"
#include "hmmer.h"

#include "hpm.h"
#include "hpmfile.h"
#include "hpm_scoreset.h"
#include "potts.h"
#include "pottsfile.h"

struct cfg_s { /* shared configuration in master and workers */
	ESL_ALPHABET	 *abc;
	ESL_MSAFILE   	 *afp;     	/* input ali file						*/
	ESL_MSA			 *msa;     	/* input msa							*/
	P7_HMMFILE		 *hfp;		/* input hmm file						*/
	HPM				 *hpm;		/* hidden potts model				*/
	HPM_SCORESET    *hpm_ss;   /* for storing hpm sores         */

};


static ESL_OPTIONS options[] = {
	/* name         type        default env   range togs  reqs  incomp      help                                                   docgroup */
	{ "-h",         eslARG_NONE,  FALSE, NULL, NULL, NULL,NULL, NULL,            "help; show brief info on version and usage",              1 },

	/* Options forcing which alphabet we're working in (normally autodetected) */
	{ "--amino",  eslARG_NONE,  FALSE,  NULL, NULL, NULL,NULL,"--dna,--rna",    "<seqfile> contains protein sequences",                   2 },
	{ "--dna",    eslARG_NONE,  FALSE, NULL, NULL, NULL,NULL,"--amino,--rna",  "<seqfile> contains DNA sequences",                        2 },
	{ "--rna",    eslARG_NONE,  FALSE, NULL, NULL, NULL,NULL,"--amino,--dna",  "<seqfile> contains RNA sequences",                        2 },
	{ 0,0,0,0,0,0,0,0,0,0 },
};
static char usage[]  = "[-options] <hpmfile> <seqfile> <scoreoutfile>";

static void
cmdline_failure(char *argv0, char *format, ...)
{
	va_list argp;
	va_start(argp, format);
	vfprintf(stderr, format, argp);
	va_end(argp);
	esl_usage(stdout, argv0, usage);
	printf("\nTo see more help on available options, do %s -h\n\n", argv0);
	exit(1);
}

int CalculateHamiltonian(HPM *hpm, P7_TRACE **tr, ESL_MSA *msa, HPM_SCORESET *hpm_ss) {
	int   z;     /* index for trace elements 	               */
	int   y;     /* index for trace elements 	               */
	int   n;     /* sequence index          	               */
	int   i;     /* match state index			               */
	int   j;     /* match state index	    		            */
	int   a;     /* residue index            	               */
	int   b;     /* residue index                            */
	int   idx;   /* index for potts parameters               */
	float E_hi;  /* pseudo-energy contribution from h_i's    */
	float E_eij; /* pseudo-energy contribution from e_ij's   */


	/* get alphabet size */
	int K = msa->abc->K;

	/* copy over sqname */
	hpm_ss->sqname = msa->sqname;

	/* loop over sequences */
	for (n=0; n < msa->nseq; n++) {
		E_hi = 0.0;
		E_eij = 0.0;

		/* print out sequence info */
		// fprintf(stdout, "%d: %d %d %d %s\n", z, tr[n]->L, tr[n]->M, tr[n]->N, hpm_ss->sqname[n]);

		/* loop over trace positions for this seq */
		//fprintf(stdout, "%s\n", msa->sqname[n]);

		for (z = 0; z < tr[n]->N; z++) {
			//fprintf(stdout, "z = %d, state = %d\n", z, tr[n]->st[z]);

			/* check if we are in a match or delete state */
			if (tr[n]->st[z] == 2 || tr[n]->st[z] == 6) {

				i = tr[n]->k[z];

				//fprintf(stdout, "\t %d %d %d\n", i, tr[n]->st[z], tr[n]->k[z]);

				/* we have a match position */
				//if (tr[n]->st[z] == P7T_M ) {
				if (tr[n]->st[z] == 2) {
					a = msa->ax[n][tr[n]->i[z]];

					/* treat non-canonical characters as gaps */
					if (a > K) {
						a = K;
					}
				}

				/* we have a delete position */
				//else if ( tr[n]->st[z] == p7T_D ) {
				else if (tr[n]->st[z] == 6) {
					a = K;
				}
				// fprintf(stdout, "h: i=%d, a=%d, h[i][a] = %.4f \n", i, a,  hpm->h[i][a]);
				E_hi = E_hi + hpm->h[i][a];

				/* now add e_ij terms to pseudo-energy */
				for (y = z+1; y < tr[n]->N; y++) {

					/* check if we are in a match position */
					 if (tr[n]->st[y] == 2 || tr[n]->st[y] == 6) {


						 j = tr[n]->k[y];

						/* we have a match state */
						if (tr[n]->st[y] == 2) {
							b =  msa->ax[n][tr[n]->i[y]];

							/* treat non-canonical characters as gaps */
							 if (b > K) {
								 b = K;
							 }
						}

						/* we have a delete position */
						else if (tr[n]->st[y] == 6) {
							b = K;
						}

						idx = IDX(a,b,msa->abc->K+1);
						E_eij += hpm->e[i][j][idx];

					 }
				}
			}

		}
		hpm_ss->E_hi[n] = E_hi;
		hpm_ss->E_eij[n] = E_eij;
	}


	return eslOK;
}

int CalculateInsertProbs(HPM *hpm, P7_TRACE **tr, ESL_MSA *msa, HPM_SCORESET *hpm_ss) {
	int   z;    /* index for trace elements 	     */
	int   n;    /* sequence index          	     */
	int   i;    /* match state index			        */
	int   a;    /* residue index            	     */
	float lp;   /* log of insert emission probs    */

	/* loop over sequences */
	for (n=0; n < msa->nseq; n++) {
		lp = 0.0;
		//fprintf(stdout, "%s\n", msa->sqname[n]);

		/* loop over trace positions for this seq */
      for (z = 0; z < tr[n]->N; z++) {


			/* we have an insert position */
			if (tr[n]->st[z] == 4 || (tr[n]->st[z] == 8 && tr[n]->i[z] > 0 ) || ((tr[n]->st[z] == 13 && tr[n]->i[z] > 0 )) ) {
				/* previous match state */
				i = tr[n]->k[z];
				/* residue */
				a = msa->ax[n][tr[n]->i[z]];
				lp = lp + log(hpm->ins[i][a]);
			}
		}
		hpm_ss->lp_ins[n] = lp;
		//fprintf(stdout, "%f\n", hpm_ss->lp_ins[n]);

	}


	return eslOK;
}

int CalculateTransitionProbs(HPM *hpm, P7_TRACE **tr, ESL_MSA *msa, HPM_SCORESET *hpm_ss) {
	int   z;            /* index for trace elements 	             */
	int   st;           /* state id index           	             */
	int   stprev;       /* state id index           	             */
	int   n;            /* sequence index            	             */
	int   i;            /* match state index			                */
	int   iprev;        /* match state index			                */
	float lp;           /* log of transition probs                  */
	float lp_NN;        /* log of N->N and C->C transition prob     */
	float lp_NB;        /* log of N->B and C->T transition prob     */
	int   L;            /* sequence length					             */

	/* loop over sequences */
	for (n=0; n < msa->nseq; n++) {
		L = 0;
		lp = 0.0;
		stprev = -1;

		/* loop over alignment columns to determine how many residues are in seq */
		/* is there a better way to do this???? */

		for( i = 1; i < msa->alen+1; i++) {
			if ( esl_abc_XIsGap(msa->abc, msa->ax[n][i]) == 0) {
				L++;
			}
		}

		/* calculate N->N and C->C transition probability, in log space */
	   lp_NN = log(L) - log(L+2);
		/* calculate N->B and C->T transition probability, in log space */
		lp_NB = log(2) - log(L+2);


		/* loop over trace positions for this seq */
      for (z = 0; z < tr[n]->N; z++) {
			/* state type */
			st = tr[n]->st[z];
			/* node in model */
			i = tr[n]->k[z];

			/* handle transitions into N state */
			if (st == 8) {

				/*ignore S->N transitions, they have prob 1 */

				/* handle N->N transitions */
				if (stprev == 8) {
					lp += lp_NN;
				}
			}
			/* handle transitions into B state */
			else if (st == 9) {

				/* handle N->B transitons */
				if (stprev == 8) {
					lp += lp_NB;
				}
				/* no other transitions into B state possible */
			}
			/* handle transitions into hybrid match-del states */
			else if (st == 2 || st == 6) {

				/* handle M->M transitions */
				if (stprev == 2 || stprev == 6) {

					lp += log(hpm->t[iprev][HPM_MM]);
				}
				/* handle I->M transitions */
				else if (stprev == 4) {
					lp += log(hpm->t[iprev][HPM_IM]);
				}
				/* ignore B->M transitions, they have prob 1 */
			}
			/* handle transitions into I states */
			else if (st == 4) {
				/* handle M->I transitions */
				if (stprev == 2 || stprev == 6) {
					lp += log(hpm->t[iprev][HPM_MI]);
				}
				/* handle I->I transitions */
				else if (stprev == 4) {
					lp += log(hpm->t[iprev][HPM_II]);
				}

			}

			/* handle transitions into C state */
			else if (st == 13) {
				if (stprev == 13) {
					lp += lp_NN;
				}
				/* ignore E->C transitions, they have prob 1 */
			}
			/*handle transitions into T state */
			else if (st == 15) {
				if (stprev == 13) {
					lp += lp_NB;
				}
				/* no other possible transitions to T state */
			}

			stprev = st;
			iprev = i;
		}

		/* add this seq's transition prob to score set */
		hpm_ss->lp_trans[n] = lp;
	}


	return eslOK;
}

int CalculateMatchProbsNull(HPM *hpm, P7_TRACE **tr, ESL_MSA *msa, HPM_SCORESET *hpm_ss) {
	int     z;             /* index for trace elements 	      */
	int     st;            /* state id index           	      */
	int     n;				  /* sequence index                    */
	int     a;				  /* residue index                     */
	float   lp;            /* null model log prob               */

	for (n=0; n < msa->nseq; n++) {
		lp = 0;

		/* loop over trace positions for this seq */
      for (z = 0; z < tr[n]->N; z++) {
			/* state type */
			st = tr[n]->st[z];

			if (st == 2) {
				a = msa->ax[n][tr[n]->i[z]];
				/* janky hack: just use 0th state insert emission prob */
				lp = lp + log(hpm->ins[0][a]);
			}
		}
		hpm_ss->lpnull_match[n] = lp;
	}


	return eslOK;


}

int CalculateTransitionProbsNull(ESL_MSA *msa, HPM_SCORESET *hpm_ss) {
	P7_BG  *bg  = NULL;
	int     n;           /* sequence index       */
	int     i;           /* position index       */
	int     L;           /* sequence length      */
	float   lp;          /* null model lot prob  */

	bg = p7_bg_Create(msa->abc);

	for (n=0; n < msa->nseq; n++) {
		/* loop over alignment columns to determine how many residues are in seq */
		/* is there a better way to do this???? */
		L = 0;
		for( i = 1; i < msa->alen+1; i++) {
			if ( esl_abc_XIsGap(msa->abc, msa->ax[n][i]) == 0) L++;
		}

		/* Set the null model's target length models */
		p7_bg_SetLength(bg, L);
		lp = 0.0;
		/* calculate log prob of null transitions */
		p7_bg_NullOne(bg, msa->ax[n], L, &lp);
		/* add above value to scoreset object */
		hpm_ss->lpnull_trans[n] = lp;

	}

	/* clean up */
	p7_bg_Destroy(bg);

	return eslOK;

}


int main(int argc, char *argv[])
{
	ESL_GETOPTS    *go;															/* cmd line configuration 			*/
	struct cfg_s    cfg;  														/* application configuration 		*/
	char           *alifile 					= NULL;						/* alignment file path		 		*/
	char				*hpmfile 					= NULL;						/* hmm file path						*/
	char				*scorefile 	    			= NULL;						/* hpm score output file path	   */
	FILE				*hpm_ss_fp 	    			= NULL;						/* hpm score output file object	*/
	P7_TRACE      **tr                     = NULL;                 /* trace for alignment paths     */
	int            *matassign              = NULL;                 /* array for ali match positions */
	int             i;                                             /* ali position index            */
	int				 status;														/* easel return code 				*/
	char            errbuf[eslERRBUFSIZE];

	/* parse command line */
	go = esl_getopts_Create(options);
	if (esl_opt_ProcessCmdline(go, argc, argv) != eslOK) cmdline_failure(argv[0], "Failed to parse command line: %s\n", go->errbuf);
	if (esl_opt_VerifyConfig(go) 					 != eslOK) cmdline_failure(argv[0], "Error in app configuration:   %s\n", go->errbuf);
	if (esl_opt_ArgNumber(go)						 != 3) 	  cmdline_failure(argv[0], "Incorrect number of command line arguments.\n", go->errbuf);

	hpmfile    = esl_opt_GetArg(go, 1);
	alifile    = esl_opt_GetArg(go, 2);
	scorefile  = esl_opt_GetArg(go, 3);

	/* Set up the configuration structure shared amongst functions here */
	cfg.abc = NULL; /* until we read the seq file below */

	/* Determine alphabet */
	if 		(esl_opt_GetBoolean(go, "--amino"))		cfg.abc = esl_alphabet_Create(eslAMINO);
	else if  (esl_opt_GetBoolean(go, "--dna"))		cfg.abc = esl_alphabet_Create(eslDNA);
	else if  (esl_opt_GetBoolean(go, "--rna"))		cfg.abc = esl_alphabet_Create(eslRNA);


	/* open the msa file */
	/* alphabet should be set in function if not user-specified */
	status = esl_msafile_Open(&(cfg.abc), alifile, NULL, eslMSAFILE_UNKNOWN, NULL, &cfg.afp);
	if (status != eslOK) esl_msafile_OpenFailure(cfg.afp, status);

	/* Read in the MSA */
	status = esl_msafile_Read(cfg.afp, &cfg.msa);
	if (status != eslOK) esl_msafile_ReadFailure(cfg.afp, status);

	/* Allow '.' as gap, too */
	esl_alphabet_SetEquiv(cfg.abc, '.', '-');

	/* read hpm file */
	cfg.hpm = hpmfile_Read(hpmfile, cfg.abc, errbuf);
	//int j;
	//int a;
	//int b;
	/*
	for (i=1; i<cfg.hpm->M+1; i++){
		for (j=i+1; j<cfg.hpm->M+1; j++) {
			for (a=0; a<cfg.abc->K+1; a++) {
				for (b=0; b<cfg.abc->K+1; b++) {
					fprintf(stdout, "i = %d, j= %d, a = %d, b = %d, eij(ab) = %f\n", i,j,a,b,cfg.hpm->e[i][j][IDX(a,b,cfg.abc->K+1)]);
				}
			}
		}
	}
	*/
	/* Set up the score set object */
	cfg.hpm_ss = hpm_scoreset_Create(cfg.msa->nseq);

	/* allocate memory for trace and match arrays */
	ESL_ALLOC(tr, sizeof(P7_TRACE *) * cfg.msa->nseq);
	ESL_ALLOC(matassign, sizeof(int) * (cfg.msa->alen + 1));

	/* extract match states from alignment */
	for (i=0; i < cfg.msa->alen; i++){
		if (cfg.msa->rf[i] != '.') {
			matassign[i+1] = 1;
		}
		else {
			matassign[i+1] = 0;
		}
		//fprintf(stdout, "%d: %c, %d\n", i, cfg.msa->rf[i], matassign[i+1]);
	}

	/* assign traces to msa seqs */
	p7_trace_FauxFromMSA(cfg.msa, matassign, p7_MSA_COORDS, tr);


	/* Calculate hamiltonian for all sequences */
	CalculateHamiltonian(cfg.hpm, tr, cfg.msa, cfg.hpm_ss);
	/* Calculate insert probabilities for all seqs */
	CalculateInsertProbs(cfg.hpm, tr, cfg.msa, cfg.hpm_ss);
	/* Calculate transition probabilities for all seqs */
	CalculateTransitionProbs(cfg.hpm, tr, cfg.msa, cfg.hpm_ss);
	/* Calculate match state null model probabilities for all seqs */
	CalculateMatchProbsNull(cfg.hpm, tr, cfg.msa, cfg.hpm_ss);
	/* calculate null transition probababilities for all sequences */
	CalculateTransitionProbsNull(cfg.msa, cfg.hpm_ss);

	/* write potts scores to outfile */
	if ((hpm_ss_fp = fopen(scorefile, "w")) == NULL) esl_fatal("Failed to open output hpm score setfile %s for writing", scorefile);

	hpm_scoreset_Write(hpm_ss_fp, cfg.hpm_ss);
	fclose(hpm_ss_fp);

	/* print hello world */
	fprintf(stdout, "hello world!\n");

	/* clean up */
	esl_alphabet_Destroy(cfg.abc);
	esl_msa_Destroy(cfg.msa);
	esl_msafile_Close(cfg.afp);
	free(matassign);
	esl_getopts_Destroy(go);

	return 0;

	ERROR:
		return status;

}
```
