## Supplemental Code S1 for "Remote homology search with hidden Potts models": hpmscore_path.c

```
/* hpmscore_path.c */

#include <stdio.h>
#include <stdlib.h>
#include <string.h>

#include "easel.h"
#include "esl_alphabet.h"
#include "esl_getopts.h"
#include "esl_msa.h"
#include "esl_msafile.h"
#include "esl_sq.h"

#include "h4_config.h"
#include "h4_mode.h"
#include "h4_path.h"

#include "hpm.h"
#include "hpmfile.h"
#include "hpm_scoreops.h"
#include "h4_path_hpm.h"

static ESL_OPTIONS options[] = {
   /* name        type        default    env range togs  reqs  incomp            help                                            docgroup */
   { "-h",        eslARG_NONE,    FALSE, NULL, NULL, NULL, NULL, NULL,            "help; show brief info on version and usage",  1 },

   /* Options forcing which alphabet we're working in (normally autodetected) */
   { "--amino",   eslARG_NONE,    FALSE, NULL, NULL, NULL, NULL, "--dna,--rna",   "We are dealing with protein sequences",       2 },
   { "--rna",     eslARG_NONE,    FALSE, NULL, NULL, NULL, NULL, "--amino,--dna", "We are dealing with rna sequences",           2 },
   { "--dna",     eslARG_NONE,    FALSE, NULL, NULL, NULL, NULL, "--amino,--rna", "We are dealing with dna sequences",           2 },

   { 0,0,0,0,0,0,0,0,0,0 },
};

static char banner[] = "Score sequence(s) and path(s) with an HPM";
static char usage[]  = "[-options] <hpmfile> <msafile>";

static void
cmdline_failure(char *argv0, char *format, ...)
{
   va_list argp;
   printf("\nERROR: ");
   va_start(argp, format);
   vfprintf(stderr, format, argp);
   va_end(argp);
   esl_usage(stdout, argv0, usage);
   printf("\nTo see more help on available options, do %s -h\n\n", argv0);
   exit(1);
}

int main(int argc, char *argv[]){
   ESL_GETOPTS      *go        = NULL;
   ESL_ALPHABET     *abc       = NULL;
   char             *hpmfile   = NULL;                       /* input hpm filepath                        */
   char             *msafile   = NULL;                       /* input msa filepath                        */
   ESL_MSAFILE      *afp       = NULL;                       /* oopen input msa file stream               */
   HPM              *hpm       = NULL;                       /* input hpm                                 */
   H4_MODE          *mo        = h4_mode_Create();           /* h4 profile hmm mode                       */
   ESL_MSA          *msa       = NULL;                       /* input msa                                 */
   ESL_SQ          **sq        = NULL;                       /* array of sequences                        */
   H4_PATH         **pi        = NULL;                       /* path array for aligned sequences          */
   int8_t           *matassign = NULL;                       /* MAT state assignments if 1; 1..alen       */
   float             hpmsc;                                  /* Joint hpm score of seq and path, in nats  */
   float             hsc;                                    /* Sum of Potts h_i terms for a seq, path    */
   float             esc;                                    /* Sum of Potts e_ij terms for a seq, path   */
   float             tsc;                                    /* HPM transition score for a seq, path      */
   float             nmsc;                                   /* Null match emission score for a seq, path */
   int               i;                                      /* msa position (column) index               */
   int               n;                                      /* msa sequence (row) index                  */
   int               fmt       = eslMSAFILE_STOCKHOLM;       /* input msa format #sorrynotsorry           */
   int               status;                                 /* easel return code                         */
   char              errbuf[eslERRBUFSIZE];                  /* for error messages                        */

   /* parse command line */
   go = esl_getopts_Create(options);
   if (esl_opt_ProcessCmdline(go, argc, argv) != eslOK) cmdline_failure(argv[0], "Failed to parse command line: %s\n", go->errbuf);
   if (esl_opt_VerifyConfig(go)               != eslOK) cmdline_failure(argv[0], "Error in app configuration:   %s\n", go->errbuf);

   /* output help if requested */
   if (esl_opt_GetBoolean(go, "-h") ) {
      esl_banner(stdout, argv[0], banner);
      esl_usage (stdout, argv[0], usage);
      puts("\n where options are:");
      esl_opt_DisplayHelp(stdout, go, 1, 2, 80);
      puts("\n Alphabet options:");
      esl_opt_DisplayHelp(stdout, go, 2, 2, 80);
      exit(0);
   }

   if      (esl_opt_GetBoolean(go, "--amino")) abc = esl_alphabet_Create(eslAMINO);
   else if (esl_opt_GetBoolean(go, "--dna"))   abc = esl_alphabet_Create(eslDNA);
   else if (esl_opt_GetBoolean(go, "--rna"))   abc = esl_alphabet_Create(eslRNA);

   /* read arguments */
   if (esl_opt_ArgNumber(go) != 2)  cmdline_failure(argv[0], "Incorrect number of command line arguments.\n", go->errbuf);

   hpmfile = esl_opt_GetArg(go, 1);
   msafile = esl_opt_GetArg(go, 2);

   /* read the msa file */
   if ((status = esl_msafile_Open(&abc, msafile, NULL, fmt, NULL, &afp)) != eslOK) esl_msafile_OpenFailure(afp, status);
   if ((status = esl_msafile_Read(afp, &msa))                            != eslOK) esl_fatal ("Failed to read MSA");
   esl_msafile_Close(afp);

   /* allow '-' and '.' as gap */
   esl_alphabet_SetEquiv(abc, '.', '-');

   /* read the hpm file */
   hpm = hpmfile_Read(hpmfile, abc, errbuf);

   /* extract sequences from MSA */
   ESL_ALLOC(sq, sizeof(ESL_SQ *) * msa->nseq);
   for (n = 0; n < msa->nseq; n++) {
      fprintf(stdout, "n: %d\n", n);
      esl_sq_FetchFromMSA(msa, n, &sq[n]);
   }

   if (! (msa->flags & eslMSA_DIGITAL)) esl_fatal("need a digital msa");
   if (msa->rf == NULL)                 esl_fatal("msa lacks an RF line");

   /* extract match states from alignment */
   ESL_ALLOC(matassign, sizeof(int8_t) * (msa->alen + 1));
   for (i=0; i < msa->alen; i++){
      if (msa->rf[i] != '.') matassign[i+1] = 1;
      else                   matassign[i+1] = 0;
   }

   /* generate trace array from msa */
   ESL_ALLOC(pi, sizeof(H4_PATH *) * msa->nseq);
   h4_path_FromMSA(msa, matassign, 0, pi);

   /* set HMM mode to uniglocal  */
   h4_mode_SetUniglocal(mo);

   /* score sequences and paths */
   for (n=0;  n < msa->nseq; n++) {
       /* configure profile and background models using seq length */
       h4_mode_SetLength(mo, sq[n]->n);

       /* score sequence and given path */
       hpm_scoreops_ScorePath(pi[n], sq[n]->dsq, hpm, mo, &hpmsc, &hsc, &esc, &tsc, &nmsc);

       fprintf(stdout, "n: %d, hpmsc: %.2f\n", n, hpmsc);
       fprintf(stdout, "\t\thsc:  %.2f\n", hsc);
       fprintf(stdout, "\t\tesc:  %.2f\n", esc);
       fprintf(stdout, "\t\ttsc:  %.2f\n", tsc);
       fprintf(stdout, "\t\tnmsc:  %.2f\n", nmsc);
   }
   /* clean up and return */
   for (n = 0; n < msa->nseq; n++) {
      esl_sq_Destroy(sq[n]);
      h4_path_Destroy(pi[n]);
   }
   free(sq);
   free(pi);
   free(matassign);
   hpm_Destroy(hpm);
   esl_msa_Destroy(msa);
   esl_alphabet_Destroy(abc);
   h4_mode_Destroy(mo);
   esl_getopts_Destroy(go);
   return 0;

   ERROR:
      return status;

}
```
