## Supplemental Code S1 for "Remote homology search with hidden Potts models": hpmscoreIS.c

```
/* hpmscoreIS.c */
/* Score (and optionally align) sequences with hidden Potts models using importance samplign and HMM's */

#include <stdio.h>
#include <stdlib.h>
#include <string.h>

/* easel includes */
#include "easel.h"
#include "esl_alphabet.h"
#include "esl_getopts.h"
#include "esl_sq.h"
#include "esl_sqio.h"
#include "esl_vectorops.h"

/* h4 nwo includes */
#include "h4_hmmfile.h"
#include "h4_mode.h"
#include "h4_profile.h"
#include "h4_refmx.h"
#include "general.h"
#include "logsum.h"
#include "reference_dp.h"

/* HPMHomology includes */
#include "hpm.h"
#include "hpmfile.h"
#include "hpm_scoreset.h"
#include "hpm_scoreops.h"
#include "h4_path_hpm.h"
#include "h4_pathalign.h"

/* declaration of internal functions */
int   Calculate_IS_scores(HPM *hpm, H4_PROFILE *hmm, ESL_SQ **sq, ESL_RANDOMNESS *rng, HPM_IS_SCORESET *hpm_is_ss, ESL_MSA **msa, char *scoreprogfile, char *aliprogfile, int start, int end, int totseq, int R, int R_batch, int A, int v, int fj);
int   find_jumps(double *pr_unsorted, int r, int R_batch, ESL_SQ *sq, H4_PATH **pi, H4_PROFILE *hmm, H4_MODE *mo, float fsc);
char *strsafe(char *dest, const char *src);

static ESL_OPTIONS options[] = {
   /* name          type            default env   range  togs  reqs            incomp           help                                                 docgroup */
   { "-h",          eslARG_NONE,    FALSE,  NULL, NULL,  NULL, NULL,           NULL,            "help; show brief info on version and usage",               1 },

   /* Options forcing which alphabet we're working in (normally autodetected) */
   { "--amino",     eslARG_NONE,    FALSE,  NULL, NULL,  NULL, NULL,           "--dna,--rna",   "<seqfile> contains protein sequences",                     2 },
   { "--rna",       eslARG_NONE,    FALSE,  NULL, NULL,  NULL, NULL,           "--dna,--amino", "<seqfile> contains RNA sequences",                         2 },
   { "--dna",       eslARG_NONE,    FALSE,  NULL, NULL,  NULL, NULL,           "--rna,--amino", "<seqfile> contains DNA sequences",                         2 },

   /* options for bounding sequences we score */
   { "--seqstart",  eslARG_INT,     "-1",   NULL, NULL,  NULL, NULL,           NULL,            "Start sequence",                                           3 },
   { "--seqend",    eslARG_INT,     "-2",   NULL, NULL,  NULL, NULL,           NULL,            "End sequence",                                             3 },

    /* options for controlling number of alignments sampled per sequence */
   { "-R",          eslARG_INT,     "1000", NULL, "n>0", NULL, NULL,           NULL,            "Number of HMM paths sampled per sequence",                 4 },
   { "--Rbatch",    eslARG_INT,     "1000", NULL, "n>0", NULL, NULL,           NULL,            "Number of HMM paths sampled per batch (for speed up)",     4 },
   { "-s",          eslARG_INT,     "0",    NULL, NULL,  NULL, NULL,           NULL,            "set random number seed to <n>",                            4 },

   /* control of output */
   { "-A",          eslARG_OUTFILE, NULL,   NULL, NULL,  NULL, NULL,           NULL,            "save multiple alignment of all hits to file <s>",          5 },
   { "--scoreprog", eslARG_OUTFILE, NULL,   NULL, NULL,  NULL, NULL,           NULL,            "save .csv with info on IS scoring progress to file <s>",   5 },
   { "--aliprog",   eslARG_OUTFILE, NULL,   NULL, NULL,  NULL, "-A",           NULL,            "save .csv with info on IS alignment prgoress to file <s>", 5 },

   /* debugging tools */
   { "-v",          eslARG_NONE,    FALSE,  NULL, NULL,  NULL, NULL,          NULL,            "Verbose mode: print info on intermediate scoring steps",    5 },
   { "--findjumps", eslARG_NONE,    FALSE,  NULL, NULL,  NULL, "--scoreprog", NULL,            "Output info on sampled paths that cause scoe jumps",        5 },

   { 0, 0, 0, 0, 0, 0, 0, 0, 0, 0 },
};

static char usage[]  = "[-options] <hpmfile> <hmmfile> <seqfile> <scoreoutfile>";
static char banner[] = "Score unaligned sequences with an HPM via importance sampling";

static void
cmdline_failure(char *argv0, char *format, ...) {
   va_list argp;
   printf("\nERROR: ");
   va_start(argp, format);
   vfprintf(stderr, format, argp);
   va_end(argp);
   esl_usage(stdout, argv0, usage);
   printf("\nTo see more help on available options, do %s -h\n\n", argv0);
   exit(1);
}


int main(int argc, char **argv) {

   ESL_GETOPTS      *go            = NULL;                  /* application configuration                         */
   ESL_RANDOMNESS   *rng           = NULL;                  /* random number generator                           */
   ESL_ALPHABET     *abc           = NULL;                  /* biological alphabet                               */
   char             *hpmfile       = NULL;                  /* input HPM filepath                                */
   char             *hmmfile       = NULL;                  /* input HMM filepath                                */
   char             *seqfile       = NULL;                  /* input seq filepath                                */
   char             *scorefile     = NULL;                  /* output score filepath                             */
   char             *scoreprogfile = NULL;                  /* output IS scoring progress filepath (--scoreprog) */
   char             *aliprogfile   = NULL;                  /* output IS alignment progress filepath (--aliprog) */
   H4_HMMFILE       *hmmfp         = NULL;                  /* open input hmm file stream                        */
   ESL_SQFILE       *sqfp          = NULL;                  /* open seq file stream                              */
   H4_PROFILE       *hmm           = NULL;                  /* hmm                                               */
   HPM              *hpm           = NULL;                  /* hpm                                               */
   ESL_SQ          **sq            = NULL;                  /* array of sequences                                */
   HPM_IS_SCORESET  *hpm_is_ss     = NULL;                  /* object w/ sequence scores                         */
   int               format        = eslSQFILE_UNKNOWN;     /* input seq file format                             */
   FILE             *afp           = NULL;                  /* alignment output file (-A)                        */
   ESL_MSA          *msa           = NULL;                  /* alignment output object (-A)                      */
   int               outfmt        = eslMSAFILE_STOCKHOLM;  /* alignment output format (-A)                      */
   FILE             *hpm_is_ss_fp  = NULL;                  /* file stream for scores                            */
   int               n             = 0;                     /* sequence index                                    */
   int               totseq        = 0;                     /* number of seqs in seqfile                         */
   int               nseq          = 0;                     /* number of seqs we deal with                       */
   int               start,end;                             /* start and end seq indices                         */
   int               R;                                     /* number of sampled paths/seq                       */
   int               R_batch;                               /* number of sampled paths/batch                     */
   int               A             = 0;                     /* Boolean for output alignment  (-A)                */
   int               v             = 0;                     /* Boolean for verbose mode (-v)                     */
   int               fj            = 0;                     /* Boolean for finding jumping paths (--findjumps)   */
   int               status;                                /* Easel return code                                 */
   char              errbuf[eslERRBUFSIZE];                 /* for error messages                                */

   /* parse command line */
   go = esl_getopts_Create(options);
   if (esl_opt_ProcessCmdline(go, argc, argv) != eslOK) cmdline_failure(argv[0], "Failed to parse command line: %s\n", go->errbuf);
   if (esl_opt_VerifyConfig(go)               != eslOK) cmdline_failure(argv[0], "Error in app configuration:   %s\n", go->errbuf);

   if (esl_opt_GetBoolean(go, "-h") ) {
      esl_banner(stdout, argv[0], banner);
      esl_usage (stdout, argv[0], usage);
      puts("\n where options are:");
      esl_opt_DisplayHelp(stdout, go, 1, 2, 80);
      puts("\n Alphabet options:");
      esl_opt_DisplayHelp(stdout, go, 2, 2, 80);
      puts("\n Options for bounding sequences we score:");
      esl_opt_DisplayHelp(stdout, go, 3, 2, 80);
      puts("\n Sampling options:");
      esl_opt_DisplayHelp(stdout, go, 4, 2, 80);
      puts("\n MSA Output options:");
      esl_opt_DisplayHelp(stdout, go, 5, 2, 80);
      puts("\n Debug options:");
      esl_opt_DisplayHelp(stdout, go, 6, 2, 80);
      exit(0);
   }

   /* read arguments */
   if (esl_opt_ArgNumber(go) != 4)  cmdline_failure(argv[0], "Incorrect number of command line arguments.\n", go->errbuf);

   hpmfile   =  esl_opt_GetArg(go, 1);
   hmmfile   =  esl_opt_GetArg(go, 2);
   seqfile   =  esl_opt_GetArg(go, 3);
   scorefile =  esl_opt_GetArg(go, 4);

   /* check for verbose mode */
   if (esl_opt_GetBoolean(go, "-v")) v=1;

   /* if output msa requested by user, try to open it */
   if (esl_opt_IsOn(go, "-A")) {
      A = 1;
      if ((afp = fopen(esl_opt_GetString(go, "-A"), "w")) == NULL) esl_fatal("Failed to open alignment file %s for writing\n", esl_opt_GetString(go, "-A"));
   }

   /* if IS intermediate scoring progress csv is requested by user, set variables*/
   if (esl_opt_IsOn(go, "--scoreprog")) scoreprogfile = (esl_opt_GetString(go, "--scoreprog"));

   /* check if we are looking at jumping paths */
   if (esl_opt_IsOn(go, "--findjumps")) fj = 1;

   /* if IS intermediate alignment progress csv is requested by user, set variables*/
   if (esl_opt_IsOn(go, "--aliprog")) aliprogfile = (esl_opt_GetString(go, "--aliprog"));

   /* if user has defined an alphabet we define it here */
   if        (esl_opt_GetBoolean(go, "--amino"))       abc = esl_alphabet_Create(eslAMINO);
   else if   (esl_opt_GetBoolean(go, "--rna"))         abc = esl_alphabet_Create(eslRNA);
   else if   (esl_opt_GetBoolean(go, "--dna"))         abc = esl_alphabet_Create(eslDNA);

   /* check if user has specified the number of samples/sequence */
   R = esl_opt_GetInteger(go, "-R");

   /* check if user has specified the number of samples/batch */
   R_batch = esl_opt_GetInteger(go, "--Rbatch");

   /* create random number generator */
   rng = esl_randomness_Create(esl_opt_GetInteger(go, "-s"));

   /* open the .hmm file */
   if ( h4_hmmfile_Open(hmmfile, NULL, &hmmfp) != eslOK) esl_fatal("couldn't open profile file %s", hmmfile);
   /* read first hmm  from hmm file */
   if ( h4_hmmfile_Read(hmmfp, &abc, &hmm)     != eslOK) esl_fatal("failed to read profile from file %s", hmmfile);
   h4_hmmfile_Close(hmmfp);

   /* open the HPM file */
   hpm = hpmfile_Read(hpmfile, abc, errbuf);

   /* open the sequence file */
   status = esl_sqfile_OpenDigital(abc, seqfile, format, NULL, &sqfp);
   if      (status == eslENOTFOUND) esl_fatal("No such file.");
   else if (status == eslEFORMAT)   esl_fatal("Format unrecognized.");
   else if (status == eslEINVAL)    esl_fatal("Can't autodetect stdin or .gz.");
   else if (status != eslOK)        esl_fatal("Open failed, code %d.", status);

   /* read sequences into array */
   ESL_REALLOC(sq, sizeof(ESL_SQ *));
   sq[n] = esl_sq_CreateDigital(abc);
   while ((status = esl_sqio_Read(sqfp, sq[n+nseq])) == eslOK) {
      nseq++;
      ESL_REALLOC(sq, sizeof(ESL_SQ *) * (nseq+1));
      sq[n+nseq] = esl_sq_CreateDigital(abc);
   }

   /* error handling and cleanup */
   if      (status == eslEFORMAT) esl_fatal("Parse failed (sequence file %s):\n%s\n",
                                            sqfp->filename, esl_sqfile_GetErrorBuf(sqfp));
   else if (status != eslEOF)     esl_fatal("Unexpected error %d reading sequence file %s",
                                            status, sqfp->filename);

   /* if user specified bounds, set them here */
   start = 0;
   end   = nseq;
   if (esl_opt_GetInteger(go, "--seqstart") > -1) start = esl_opt_GetInteger(go, "--seqstart");
   if (esl_opt_GetInteger(go, "--seqend")   > -1)     end = esl_opt_GetInteger(go, "--seqend");

   /* check if end is beyond number of sequences in input file */
   if (end > nseq) end = nseq;

   /* total number of sequences we will be scoring */
   totseq = end-start;
   fprintf(stdout, "totseq: %d\n", totseq);

   /* create scoreset object */
   hpm_is_ss = hpm_is_scoreset_Create(totseq);

   /* initiate logsum magic */
   h4_logsum_Init();

   Calculate_IS_scores(hpm, hmm, sq, rng, hpm_is_ss, &msa, scoreprogfile, aliprogfile, start, end, totseq, R, R_batch, A, v, fj);

   /* write hpm is scores to output csv */
   if ((hpm_is_ss_fp = fopen(scorefile, "w")) == NULL) esl_fatal("Failed to open output hpm IS scoreset file %s for writing", scorefile);
   hpm_is_scoreset_Write(hpm_is_ss_fp, hpm_is_ss);
   fclose(hpm_is_ss_fp);

   /* if output msa requested, write it to output file */
   if (A) {
      esl_msafile_Write(afp, msa, outfmt);
      fclose(afp);
      esl_msa_Destroy(msa);
   }

   /* clean up and return */
   for (n = 0; n < nseq+1; n++) {
      esl_sq_Destroy(sq[n]);
   }
   free(sq);
   esl_sqfile_Close(sqfp);
   h4_profile_Destroy(hmm);
   hpm_Destroy(hpm);
   hpm_is_scoreset_Destroy(hpm_is_ss);
   esl_randomness_Destroy(rng);
   esl_alphabet_Destroy(abc);
   esl_getopts_Destroy(go);

   return 0;

   ERROR:
      return status;
}


/* Function: Calculate_IS_scores()
 *
 * Purpose: Given target sequences <sq>, a query HPM <hpm>, and an h4 HMM
 *          <hmm>, estimate the unnormalized score of each sequence under
 *          the HPM, S^*(x). The importance sampling algorithm is used,
 *          with the HMM serving as the proposal distribution.
 *
 * args:    hpm            - query HPM
 *          hmm            - HMM, used as proposal distributiion
 *          sq             - array of target sequences
 *          rng            - random number generator
 *          hpm_is_ss      - hpm scoreset object, for storing scores
 *          msa            - optional msa with estimated optimal paths under
 *                           the HMM, used with -A (else NULL)
 *          scoreprogfile - optional path to .csv file with intermediate IS
 *                          scores, used with --scoreprogfile (else NULL)
 *          aliprogfile   - optional path to .csv file with intermediate
 *                          estimated optimal path scores, used with
 *                          --aliprogfile (else NULL)
 *          start         - start sequence index in <sq>
 *          end           - end sequence index in <sq>
 *          totseq        - total number of sequences to score
 *          R             - number of paths sampled / sequence
 *          R_batch       - number of paths sampled / batch (for speed)
 *          A             - boolean to see if we are outputting an MSA
 *          v             - boolean for verbose mode (TBD)
 *          fj            - boolean to call find_jumps on "jumping" paths
 *
 * Returns: eslOK on success.
 *
 * */


int Calculate_IS_scores(HPM *hpm, H4_PROFILE *hmm, ESL_SQ **sq, ESL_RANDOMNESS *rng, HPM_IS_SCORESET *hpm_is_ss, ESL_MSA **msa, char *scoreprogfile, char *aliprogfile, int start, int end, int totseq, int R, int R_batch, int A, int v, int fj)
{
   H4_MODE        *mo         = h4_mode_Create();           /* h4 profile hmm mode                           */
   H4_REFMX       *fwd        = h4_refmx_Create(100, 100);  /* forward DP matrix                             */
   ESL_SQ        **sq_dummy;                                /* dummy sequence array for sampled alignments   */
   ESL_SQ        **out_sq      = NULL;                      /* traces used to construct output msa (-A)      */
   H4_PATH       **pi_dummy;                                /* dummy path array for sampled alignments       */
   H4_PATH       **out_pi      = NULL;                      /* paths used to construct output MSA            */
   FILE           *scoreprogfp = NULL;                      /* score progress output file (--scoreprog)      */
   FILE           *aliprogfp   = NULL;                      /* alignment progress output file (--aliprog)    */
   int             i,j;                                     /* sequence indices                              */
   int             b;                                       /* batch index                                   */
   int             r,s;                                     /* alignment sample indices                      */
   int             N_batch;                                 /* total number of batches                       */
   int             nBetter;                                 /* number of better alignments we've found       */
   float           fsc;                                     /* forward partial log-odds score, in bits       */
   float           hmmsc_ld;                                /* hmm log odds S(x,pi), in bits                 */
   float           hpmsc_ld;                                /* hpm unnormalized log odds S*(x, pi), in bits  */
   float           hpmsc_max;                               /* best hpm log odds S*(x, pi) for all paths     */
   float           ldprev;                                  /* previous IS log-odds score                    */
   double          ld;                                      /* importance sampling log odds score S*(x)      */
   double          ls;                                      /* log of importance sampling sum                */
   double         *pr, *pr_unsorted;                        /* terms in importance sampling sum              */
   int             status;                                  /* easel return code                             */
   //char            errbuf[eslERRBUFSIZE];                   /* buffer for easel errors                       */

   /* if score progress file is requested, open it */
   if (scoreprogfile) {
      if ((scoreprogfp = fopen(scoreprogfile, "w")) == NULL) esl_fatal("Failed to open file %s for writing\n", scoreprogfile);
       fprintf(scoreprogfp, "id,iter,hpmis_logodds\n");
   }

    /* if alignment progress file is requested, open it */
   if (aliprogfile) {
      if ((aliprogfp = fopen(aliprogfile, "w")) == NULL) esl_fatal("Failed to open file %s for writing\n", aliprogfile);
      fprintf(aliprogfp, "id,iter,hpmis_logodds\n");
   }

   if (A) {
      /* allocate memory for paths for output MSA */
      ESL_ALLOC(out_pi, sizeof(H4_PATH *) * totseq);
      ESL_ALLOC(out_sq, sizeof(ESL_SQ *) * (totseq));
   }

   /* calculate number of batches */
   N_batch = (R / R_batch) + (R % R_batch != 0);
   fprintf(stdout, "N_batch: %d\n", N_batch);

   /* allocate space for dummy trace  and sequence arrays */
   ESL_ALLOC(sq_dummy, sizeof(ESL_SQ *) * R_batch);
   ESL_ALLOC(pi_dummy, sizeof(H4_PATH *) * R_batch);
   for (r = 0; r < R_batch; r++) {
      sq_dummy[r] = esl_sq_CreateDigital(hmm->abc);
      pi_dummy[r] = h4_path_Create();
   }

   /* allocate array to store terms in importance sampling sum */
   ESL_ALLOC(pr, R*sizeof(double));
   ESL_ALLOC(pr_unsorted, R*sizeof(double));
   esl_vec_DSet(pr, R, 0.0);
   esl_vec_DSet(pr_unsorted, R, 0.0);

   /* set HMM mode to uniglocal  */
   h4_mode_SetUniglocal(mo);

   /* outer loop over sequences */
   for (i=start; i < end; i++) {
      nBetter = 0;
      ldprev = 0.0;

      /* index for score set: must start at 0 */
      j=i-start;

      /* for getting optimal path under hpm */
      if (A) {
         hpmsc_max  = -eslINFINITY;
         out_sq[j] = esl_sq_CreateDigital(hmm->abc);
         esl_sq_Copy(sq[i], out_sq[j]);
      }

      /* set profile length */
      h4_mode_SetLength(mo, sq[i]->n);

      /* run forward algorithm */
      h4_reference_Forward(sq[i]->dsq, sq[i]->n, hmm, mo, fwd, &fsc);

      /* prep for sampling paths */
      esl_vec_DSet(pr, R, 0.0);
      esl_vec_DSet(pr_unsorted, R, 0.0);

      /* middle loop over batches of sampled alignments */
      for (b = 0; b < N_batch; b++) {

         /* inner loop 1: sample alignments from HMM */
         for (s = 0;  s < R_batch; s++) {

            /* calculte overall sample number */
            r = b*R_batch + s;
            /* if we've sampled as much as requested, skip inner loop 1 */
            if (r >= R) goto cleanup;

            /* copy sequence into dummy array */
            esl_sq_Copy(sq[i], sq_dummy[s]);

            /* get stochastic traceback */
            h4_reference_StochasticTrace(rng, NULL, hmm, mo, fwd, pi_dummy[s]);

            //h4_profile_Dump(stdout, hmm);

            /* score path under HMM */
            h4_path_Score(pi_dummy[s], sq[i]->dsq, hmm, mo, &hmmsc_ld);
            pr[r] -= (hmmsc_ld / eslCONST_LOG2R);

            /* score path under HPM */
            hpm_scoreops_ScorePath(pi_dummy[s], sq[i]->dsq, hpm, mo, &hpmsc_ld, NULL, NULL, NULL, NULL);
            pr[r] += hpmsc_ld;

            /* for output alignment , see if we've got a better path */
            if (A && hpmsc_ld > hpmsc_max) {
               hpmsc_max = hpmsc_ld;
               /* if we're tracking alignment progress, print S(x,pi) of new best alignment */
               if (aliprogfile) fprintf(aliprogfp, "%s,%d,%.2f\n", sq[i]->name, r, hpmsc_max);

               /* if we've already created out_pi[j], free it before recreating it */
               if (nBetter > 0) h4_path_Destroy(out_pi[j]);
               out_pi[j] = h4_path_Clone(pi_dummy[s]);
               nBetter++;

            }

         }

         /* handle this batch of path scores */
         /* if requested, track intermediate scoring progress for this batch */
         if (scoreprogfile) {

            /* get IS score up to this point */
            esl_vec_DCopy(pr,r+1, pr_unsorted);
            esl_vec_DSortIncreasing(pr, r+1);
            ls = esl_vec_DLogSum(pr, r+1);
            ld = (fsc / eslCONST_LOG2R) - logf(r+1) + ls;

            /* record to scoreprog file */
            fprintf(stdout, "r: %d, intermediate score: %2f\n", r, ld);
            fprintf(scoreprogfp, "%s,%d,%.2f\n", sq[i]->name, r, ld);

            /* if we have a jump, track it down if requested by caller */
            if ( (r > 1e5) && (ld-ldprev > 0.25) && fj )  find_jumps(pr_unsorted, r, R_batch, sq[i], pi_dummy, hmm, mo, fsc);

            ldprev = ld;
         }


         cleanup: for (s = 0; s < R_batch; s++) {
                     esl_sq_Reuse(sq_dummy[s]);
                     h4_path_Reuse(pi_dummy[s]);
                  }
      }
      /* calculate IS approximation for this sequence */
      /* sort to increase accuracy if LogSum */
      esl_vec_DSortIncreasing(pr, R);

      /* run log sum */
      ls = esl_vec_DLog2Sum(pr, R);
      ld = (fsc / eslCONST_LOG2R) - logf(R) + ls;

      /* add scoring info to scoreset object */
      hpm_is_ss->sqname[j] = sq[i]->name;
      hpm_is_ss->R[j]      = R;
      hpm_is_ss->H[j]      = -1.0;
      hpm_is_ss->fwd[j]    = (fsc - mo->nullsc) / eslCONST_LOG2R;
      hpm_is_ss->is_ld[j]   = ld;

   }

   /* if output msa requested, create it from traces */
   if (A) {
      h4_pathalign_Seqs(out_sq, out_pi, totseq, hmm->M, TRUE, hmm, msa);
   }

   /* clean up and return */
   if (A) {
      for (j=0; j<totseq; j++) {
         esl_sq_Destroy(out_sq[j]);
         h4_path_Destroy(out_pi[j]);
      }
      free(out_sq);
      free(out_pi);
   }
   if (scoreprogfp) fclose(scoreprogfp);
   if (aliprogfp) fclose(aliprogfp);
   for (s = 0; s < R_batch; s++) {
      esl_sq_Destroy(sq_dummy[s]);
      h4_path_Destroy(pi_dummy[s]);
   }
   free(sq_dummy);
   free(pi_dummy);
   free(pr);
   free(pr_unsorted);
   h4_refmx_Destroy(fwd);
   h4_mode_Destroy(mo);
   return eslOK;

   ERROR:
      return status;

}

/* Function: find_jumps()
 *
 * Purpose:  Poor importance sampling often shows sudden "jumps" in bitscore
 *           due to a single sample. Given a set of <R_batch> sampled paths
 *           of sequence <sq> through model <hmm>, find the path(s) that
 *           cause(s) the jump in score. Print S_HMM(x,pi) and \tilde{S}_HPM(x,pi)
 *           to stdout. Also, output the alignment to an output msafile in the
 *           current working directly.
 *
 *           In case this isn't clear, this function is mostly for debugging.
 *           It's only called with the --scoreprog option set.
 *
 *
 * args:    pr_unsorted   - vector containing ratios of bitscores. At least
 *                          the last <R_batch> elements are in the order in
 *                          which they were sampled.
 *          r             - sample index of the **last** path in the batch.
 *          R_batch       - number of sampled paths in this batch
 *          sq            - sequence we are scoring
 *          pi            - <R_batch>-sized path array with batch of sampled
 *                          paths.
 *          hmm           - H4 HMM used for generating paths.
 *          fsc           - forward score of <sq> under HMM, in bits.
 *
 * Returns: eslOK on success.
 *
 * */

int find_jumps(double *pr_unsorted, int r, int R_batch, ESL_SQ *sq, H4_PATH **pi, H4_PROFILE *hmm, H4_MODE *mo, float fsc) {
   double    *pr_sorted;                                  /* vector of log-odds ratios, sorted from largest to smallest */
   float      ls;                                         /* log_sum of pr_sorted                                       */
   float      ld;                                         /* IS log odds score                                          */
   float      ldprev;                                     /* Previous iteration's IS log-odds score                     */
   float      hmmsc_ld;                                   /* log-odds score of a sequence and path given <hmm>          */
   int        s,t;                                        /* IS iteration indices                                       */
   H4_PATH  **out_pi             = NULL;                  /* 1-element path array for outputting path to an MSA         */
   ESL_SQ   **out_sq;                                     /* 1-element seq array for outputting path to an MSA          */
   ESL_MSA   *out_msa            = NULL;                  /* output MSA object                                          */
   FILE      *afp                = NULL;                  /* output alignment file                                      */
   char       out_msa_path[200];                          /* output MSA filepath                                        */
   char       sqname_safe[100];                           /* sequence name with '/' replace with '_'                    */
   int        outfmt             = eslMSAFILE_STOCKHOLM;  /* output MSA format. I am unwavering on this matter.         */
   int        njump              = 0;                     /* keep track of number of jumping paths in this batch        */
   int        status;                                     /* esl return code                                            */

   strsafe(sqname_safe, sq->name);

   /* allocate memory */
   ESL_ALLOC(out_pi, sizeof(H4_PATH *));
   ESL_ALLOC(out_sq, sizeof(ESL_SQ *));
   ESL_ALLOC(pr_sorted, r*sizeof(double));

   /* create the output sequence */
   out_sq[0] = esl_sq_CreateDigital(hmm->abc);
   esl_sq_Copy(sq, out_sq[0]);

   /* figure out what the IS score was before this batch */
   s = r-R_batch;
   /* copy pr_unsorted to pr_unsorted */
   esl_vec_DCopy(pr_unsorted, r-R_batch+1, pr_sorted);
   /* sort pr_sorted for more accurate log summing */
   esl_vec_DSortIncreasing(pr_sorted, s+1);
   ls = esl_vec_DLog2Sum(pr_sorted,s+1);
   /* calcuate full log odds-score */
   ldprev = fsc - log2f(r-R_batch+1) + ls;

   /* now loop through paths and find jump(s) */
   for (s = r-R_batch+1; s<r; s++){

      /* index w/in batch: t = 0,...,R_batch-1 */
      t = s-r+R_batch-1;
      fprintf(stdout, "\tt: %d\n", t);

      /* calculate log-odds score at this iteration */
      esl_vec_DCopy(pr_unsorted,s+1, pr_sorted);
      esl_vec_DSortIncreasing(pr_sorted, s+1);
      ls = esl_vec_DLog2Sum(pr_sorted, s+1);
      ld = fsc - log2f(s+1) + ls;

      /* put path that causes jump in an output MSA file */
      if (fabsf(ld-ldprev) > 0.25) {

         /* calculate HMM score of this seq and sampled path */
         h4_path_Score(pi[t], sq->dsq, hmm, mo, &hmmsc_ld);
         hmmsc_ld /= eslCONST_LOG2R;

         if (njump > 0) {
            h4_path_Destroy(out_pi[0]);
            esl_msa_Destroy(out_msa);
         }

         out_pi[0] = h4_path_Clone(pi[t]);
         h4_pathalign_Seqs(out_sq, out_pi, 1, hmm->M, TRUE, hmm, &out_msa);

         /* create unique output MSA path */
         sprintf(out_msa_path,"%s_path_%d.sto", sqname_safe, s);

         /* write the msa file */
         if ((afp = fopen(out_msa_path, "w")) == NULL) esl_fatal("Failed to open alignment file %s for writing\n", out_msa_path);
         esl_msafile_Write(afp, out_msa, outfmt);
         fclose(afp);

         /* print info about jumping path */
         fprintf(stdout, "\nJumping IS score for sequence %s, path %d\n", sq->name, s);
         fprintf(stdout, "IS score at previous iteration:                  %.2f\n", ldprev);
         fprintf(stdout, "IS score at this iteration:                      %.2f\n", ld);
         fprintf(stdout, "HPM log-odds score for this seq, path:           %.2f\n", pr_unsorted[s] + hmmsc_ld);
         fprintf(stdout, "HMM log-odds score for this seq, path:           %.2f\n",  hmmsc_ld);
         fprintf(stdout, "This path's contribution to IS sum (in bits):    %.2f\n",  pr_unsorted[s]);
         fprintf(stdout, "Log posterior probability of path under HMM:     %.2f\n",  hmmsc_ld - (fsc  / eslCONST_LOG2R) );

         h4_path_DumpAnnotated(stdout, out_pi[0], hmm, mo, sq->dsq);

         /* note that we've seen a jump */
          njump++;
      }

      ldprev = ld;
   }


   /* clean up and return */
   if (njump > 0) {
      h4_path_Destroy(out_pi[0]);
      esl_msa_Destroy(out_msa);
   }

   esl_sq_Destroy(out_sq[0]);
   free(out_pi);
   free(out_sq);
   free(pr_sorted);
   return eslOK;

   ERROR:
      return status;

}

/* Function: strsafe()
 *
 * Purpose:  Some sequence names have '/' characters in them. When creating
 *           output files with the sequence name in the path (as we do in
 *           find_jumps()), this causes issues. Here replace all of the '/'
 *           characters in a string with '_' characters.
 *
 * args:    dest - New string with '_' chars instead of '/'
 *          src  - Original string
 *
 * Returns: dest
 *
 * */

char *strsafe(char *dest, const char *src)
{
   int i;
   for (i=0; src[i] != '\0'; i++) {
      if (src[i] == '/') dest[i] = '_';
      else               dest[i] = src[i];
   }

   dest[i] = '\0';

   return dest;
}
```
