## Supplemental Code S1 for "Remote homology search with hidden Potts models": Makefile

```
#useful directories
ESLDIR = ../../hmmer4/easel
HMMERDIR = ../../hmmer4/src
H4DIR = ../../hmmer4/nwo

#variables that I will run
INCLUDES = -I${HMMERDIR} -I${H4DIR} -I${ESLDIR}
IFLAGS   = -I${ESLDIR} -I${HMMERDIR} -I${H4DIR} -I../src
LFLAGS   = -L${ESLDIR} -L${HMMERDIR} -L${H4DIR} -L../src
DFLAGS   = -DHPM_3MER
LIBS     = -lm
SLIBS    = ${HMMERDIR}/libhmmer.a ${H4DIR}/libhmmer.a ${ESLDIR}/libeasel.a

#compiler information
CC = gcc
CFLAGS = -Wall -g

# targets for make

all: hpm hpmfile hpm_scoreset hpm_trace potts pottsfile hpmscore hpmbuild hmm_scoreset hpm_scoreops hmm_entropy h4_path_hpm h4_pathalign hmmscore_uniglocal hmmalign_uniglocal hpmscoreIS hpmscore_path hpmalign hpmemit hpmbuild_masked

hpm: hpm.c
	${CC} ${CFLAGS} -c -o hpm.o ${IFLAGS} hpm.c

hpmfile: hpmfile.c
	${CC} ${CFLAGS} -c -o hpmfile.o ${IFLAGS} hpmfile.c

hpm_scoreset: hpm_scoreset.c
	${CC} ${CFLAGS} -c -o hpm_scoreset.o ${IFLAGS} hpm_scoreset.c

hpm_scoreops: hpm_scoreops.c
	${CC} ${CFLAGS} -c -o hpm_scoreops.o ${IFLAGS} hpm_scoreops.c

hpm_trace: hpm_trace.c
	${CC} ${CFLAGS} -c -o hpm_trace.o ${IFLAGS} hpm_trace.c

potts: potts.c
	${CC} ${CFLAGS} -c -o potts.o ${IFLAGS} potts.c

pottsfile: pottsfile.c
	${CC} ${CFLAGS} -c -o pottsfile.o ${IFLAGS} pottsfile.c

hmm_scoreset: hmm_scoreset.c
	${CC} ${CFLAGS} -c -o hmm_scoreset.o ${IFLAGS} hmm_scoreset.c

hmm_entropy: hmm_entropy.c
	${CC} ${CFLAGS} -c -o hmm_entropy.o ${IFLAGS} hmm_entropy.c

h4_path_hpm: h4_path_hpm.c
	${CC} ${CFLAGS} -c -o h4_path_hpm.o ${IFLAGS} h4_path_hpm.c

h4_pathalign: h4_pathalign.c
	${CC} ${CFLAGS} -c -o h4_pathalign.o ${IFLAGS} h4_pathalign.c

hpmscore: hpmscore.c
	${CC} ${CFLAGS} -o hpmscore ${IFLAGS} ${LFLAGS} hpmscore.c hpm.o hpmfile.o hpm_scoreset.o potts.o pottsfile.o ${SLIBS} ${LIBS}

hpmbuild: hpmbuild.c
	${CC} ${CFLAGS} -o hpmbuild ${IFLAGS} ${LFLAGS} hpmbuild.c hpm.o hpmfile.o potts.o pottsfile.o ${SLIBS} ${LIBS}

hmmscore_uniglocal: hmmscore_uniglocal.c
	${CC} ${CFLAGS} -o hmmscore_uniglocal ${IFLAGS} ${LFLAGS}  hmmscore_uniglocal.c hmm_scoreset.o ${SLIBS} ${LIBS}

hmmalign_uniglocal: hmmalign_uniglocal.c
	${CC} ${CFLAGS} -o hmmalign_uniglocal ${IFLAGS} ${LFLAGS} hmmalign_uniglocal.c ${SLIBS} ${LIBS}

hpmscoreIS: hpmscoreIS.c
	${CC} ${CFLAGS} -o hpmscoreIS ${IFLAGS} ${LFLAGS} hpmscoreIS.c hpm.o hpmfile.o potts.o pottsfile.o hpm_scoreops.o hpm_scoreset.o h4_path_hpm.o h4_pathalign.o ${SLIBS} ${LIBS}

hpmscore_path: hpmscore_path.c
	${CC} ${CFLAGS} -o hpmscore_path ${IFLAGS} ${LFLAGS} hpmscore_path.c hpm.o hpmfile.o hpm_scoreops.o h4_path_hpm.o ${SLIBS} ${LIBS}

hpmalign: hpmalign.c
	${CC} ${CFLAGS} -o hpmalign ${IFLAGS} ${LFLAGS} hpmalign.c hpm.o hpmfile.o potts.o pottsfile.o hmm_entropy.o hpm_scoreops.o hpm_trace.o ${SLIBS} ${LIBS}

hpmemit: hpmemit.c
	${CC} ${CFLAGS} -o hpmemit ${IFLAGS} ${LFLAGS} hpmemit.c hpm.o hpmfile.o hpm_scoreops.o ${SLIBS} ${LIBS}

hpmbuild_masked: hpmbuild_masked.c
	${CC} ${CFLAGS} -o hpmbuild_masked ${IFLAGS} ${LFLAGS} hpmbuild_masked.c hpm.o hpmfile.o potts.o ${SLIBS} ${LIBS}

3mer:
	 ${CC} ${CFLAGS} -o hpm_3mer ${IFLAGS} ${LFLAGS} hpm.c ${DFLAGS} hpmfile.o hpm_scoreset.o potts.o pottsfile.o ${SLIBS} ${LIBS}
	 ${CC} ${CFLAGS} -o hmm_entropy_3mer ${IFLAGS} ${LFLAGS} hmm_entropy.c ${DFLAGS} ${SLIBS} ${LIBS}

entropy_msa:
	${CC} ${CFLAGS} -o hmm_entropy_msa ${IFLAGS} ${LFLAGS}  hmm_entropy.c  -DENTROPY_MSA ${SLIBS} ${LIBS}

utest:
	 ${CC} ${CFLAGS} -o hmm_entropy_utest ${IFLAGS} ${LFLAGS} hmm_entropy.c -DHMMENTROPY_TESTDRIVE ${SLIBS} ${LIBS}

clean:
	rm -f hpm.o hpmfile.o hpm_scoreset.o hpm_scoreops.o  hpm_trace.o potts.o pottsfile.o hmm_scoreset.o hmm_entropy.o h4_path_hpm.o h4_pathalign.o hpmscore hpmbuild  hmmscore_uniglocal  hmmalign_uniglocal hpmscoreIS hpmscore_path hpmalign  hpmemit hpmbuild_masked hpm_3mer hmm_entropy_3mer hmm_entropy_msa hmm_entropy_utest


.PHONY: all clean
```
