## Supplemental Code S1 for "Remote homology search with hidden Potts models": potts.c

```
/* potts.c */
#include <potts.h>

#include "easel.h"
#include "esl_alphabet.h"
#include "hpm.h"

/* Function:  potts_Create()
 * Synopsis:  Allocate a new <POTTS> model
 *
 * Purpose:   Allocate a <POTTS> model of <M> nodes, for symbol
 *            alphabet <abc>, and return a pointer to it.
 *
 * Note:      Based on p7_hmm_Create()
 *
 * Throws:    NULL on allocation failure.
 */

POTTS *
potts_Create(int L, ESL_ALPHABET *abc)
{
   POTTS    *potts    = NULL;      /* potts model object to return */
   int       Kg       = abc->K+1;  /* alphabet size w/ gap char    */
   int       Kg2      = Kg*Kg;     /* alphabet size squared        */
   int       i,j;                  /* position indices             */
   int       status;               /* esl return code              */

   ESL_ALLOC(potts, sizeof(POTTS));

   potts->L   = L;
   potts->abc = abc;

   /* allocate memory for potts params */
   ESL_ALLOC(potts->h,  sizeof(double  *) * (L));
   ESL_ALLOC(potts->e,  sizeof(double **) * (L));

   for (i=0; i<L; i++){
      ESL_ALLOC(potts->h[i], sizeof(double ) * Kg);
      ESL_ALLOC(potts->e[i], sizeof(double *) * (L));

      for (j=0; j<L; j++) {
         ESL_ALLOC(potts->e[i][j], sizeof(double ) * Kg2);
      }
   }

   return potts;

   ERROR:
      return NULL;
}

/* Function:  potts_Destroy()
 * Synopsis:  Free a <POTTS> object
 *
 * Purpose:   Frees the body of a <POTTS> object
 *
 * Note:      Based on p7_hmm_Destroy()
 *
 * Returns:   (void).
 */

void
potts_Destroy(POTTS *potts)
{
   int i, j; /* position indices */

   if (potts == NULL) return;

   /* free hi array */
   if (potts->h && potts->L) {
      for (i=0; i < potts->L; i++) {
         if (potts->h[i]) free(potts->h[i]);
      }
      free(potts->h);
   }

   /* free eij array */
   if (potts->e && potts->L) {
      for (i=0; i < potts->L; i++) {
         if (potts->e[i]) {
            for (j=0; j < potts->L; j++) {
               if (potts->e[i][j]) free(potts->e[i][j]);
            }
            free(potts->e[i]);
         }
      }
      free(potts->e);
   }
   free(potts);
   return;
}
```
