## Supplemental Code S1 for "Remote homology search with hidden Potts models": potts.h

```
/* potts.h */

#ifndef POTTS_INCLUDED
#define POTTS_INCLUDED

#include <stdio.h>
#include "easel.h"
#include "esl_alphabet.h"


typedef struct potts_s{
   int             L;    /* number of states in model */
   double        **h;    /* site-specific h_i params  */
   double       ***e;    /* coupling e_ij params      */
   ESL_ALPHABET   *abc;  /* alphabet                  */
} POTTS;

/* potts.c */
extern POTTS *potts_Create(int L, ESL_ALPHABET *abc);
extern void   potts_Destroy(POTTS *potts);

#endif
```
