## Supplemental Code S1 for "Remote homology search with hidden Potts models": pottsfile.c

```
/* pottsfile.c */
#include <string.h>
#include <math.h>
#include <errno.h>

#include "easel.h"
#include "esl_alphabet.h"
#include "hmmer.h"

#include "hpmfile.h"
#include "pottsfile.h"
#include "potts.h"

/* Function:  pottsfile_Read()
 * Synopsis:  Read Potts model from input
 *
 * Purpose:   Reads potts model from <f>, returns it.
 *
 * Args:      f      - input file
 *            abc    - esl alphabet to assign to Potts model
 *            errbuf - memory for error messages
 *
 * Returns:   potts model <ret_potts>
 */

POTTS *
pottsfile_Read(char *f, ESL_ALPHABET *abc, char *errbuf){
   ESL_FILEPARSER  *efp           = NULL;      /* easel fileparse object */
   POTTS           *ret_potts     = NULL;      /* potts model to return  */
   char            *tok;                       /* file parser token      */
   char            *prev_tok      = "blargh";  /* file parser token      */
   int              tok_count;                 /* token count            */
   int              i;                         /* position index         */
   int              prev_i;                    /* position index         */
   int              j;                         /* position index         */
   int              a;                         /* alphabet index         */
   int              lc            = 0;         /* line counter           */
   int              section_count = 0;         /* file section counter   */
   int              status;

   if ((status = esl_fileparser_Open(f, NULL, &efp)) != eslOK) ESL_XFAIL(eslFAIL, errbuf,
                                                               "pottsfile_Read(): file open failed w/ status %s\n", status);

   /* loop through one to get model length */
   while (esl_fileparser_NextLine(efp) == eslOK) {
      if (section_count == 0) {
         tok_count = 0;

         while (esl_fileparser_GetTokenOnLine(efp, &tok, NULL) == eslOK) {

            /* if the 2nd character is an integer, we've reached the end of the h_i section */
            if ((tok_count == 1) && (IsInt(tok) > 0) && (IsInt(prev_tok))) {
               section_count = 1;
               ret_potts = potts_Create(prev_i+1, abc);
               break;
            }

            if (tok_count == 0) i = atoi(tok);
            tok_count++;
            prev_tok = tok;
         }
         prev_i = i;
      }
   }

   esl_fileparser_Close(efp);
   fprintf(stdout, "%d\n", ret_potts->L);

   /* loop through again to get potts parameters */
   if ((status = esl_fileparser_Open(f, NULL, &efp)) != eslOK) ESL_XFAIL(eslFAIL, errbuf,
                                                               "pottsfile_Read(): file open failed w/ status %s\n", status);

   while (esl_fileparser_NextLine(efp) == eslOK) {
      /* we are in the h_i section */
      if (lc < ret_potts->L) {
         tok_count = 0;

         while (esl_fileparser_GetTokenOnLine(efp, &tok, NULL) == eslOK) {

            if   (tok_count == 0) i = atoi(tok);
            else                  ret_potts->h[i][tok_count-1] = atof(tok);

            tok_count += 1;
         }
      }

      /* we are in e_ij section */
      else {
         tok_count = 0;
         while (esl_fileparser_GetTokenOnLine(efp, &tok, NULL) == eslOK) {
            /* if the line starts with two integers, those are i and j */
            if ( (tok_count == 1) && (IsInt(tok)) && (IsInt(prev_tok))) {
               i = atoi(prev_tok);
               j = atoi(tok);
               a = -1;
            }

            /* if the token is a float, we have a potts parameter */
            else if (IsInt(tok) == 0) {
               ret_potts->e[i][j][IDX(a,tok_count,abc->K+1)] = atof(tok);
               ret_potts->e[j][i][IDX(tok_count,a,abc->K+1)] = atof(tok);
            }
            tok_count ++;
            prev_tok = tok;
         }
         a++;
      }
      lc++;
   }

   esl_fileparser_Close(efp);
   return ret_potts;

   ERROR:
      return NULL;
};
```
