## Supplemental Code S1 for "Remote homology search with hidden Potts models": pottsfile.h

```
/* pottsfile.h */

#ifndef POTTSFILE_INCLUDED
#define POTTSFILE_INCLUDED

#include <stdio.h>
#include "easel.h"
#include "esl_alphabet.h"
#include "esl_fileparser.h"

#include "potts.h"

/* pottsfile.c */
POTTS *pottsfile_Read(char *f, ESL_ALPHABET *abc, char *errbuff);

#endif
```
